# Cell Cycle Arrest of a ‘Zippering’ Epithelial Cell Cluster Shapes the Face and is Disrupted in Craniofacial Disorders

**DOI:** 10.1101/2025.07.28.667310

**Authors:** T. Qu, B.H. Chacón, L. Faure, M. Losa, R. Hernández-Martínez, K. Robinson, A. Jones, S. Lisgo, J. De Anda, M. Risolino, G. Panagiotakos, E.J. Leslie-Clarkson, I. Adameyko, L. Selleri

## Abstract

Facial features identify individuals, but the mechanisms shaping the human face remain elusive. Orofacial clefting (OFC), the most common craniofacial abnormality, results from failed fusion of the facial prominences that is in part caused by persistence of the cephalic epithelium. Here we uncover the identity, behaviors, and molecular blueprints of a novel craniofacial epithelial population, the Zippering Lambda (ZL), which mediates prominence fusion and is characterized by cell cycle arrest in mouse and human embryos. Remarkably, cell cycle is unleashed in the ZL of *Pbx1/2* and *p63* mutant mice with OFC. Intersection of ZL-enriched genes with human OFC whole-genome sequencing datasets identifies *ZFHX3* variants in affected individuals and cephalic epithelial *Zfhx3* deletion causes murine OFC. ZFHX3 and PBX1 genetically interact and synergistically regulate cell cycle inhibitor genes within a complex in embryonic faces. Collectively, we deconstruct new mechanisms that pattern the face, connecting cell cycle arrest to developmental tissue fusion.

## Introduction

Our faces uniquely shape our identities and forge our connections. Early craniofacial morphogenesis requires fusion of transient swellings called the facial prominences. This process occurs between gestational weeks 4^th^-8^th^ in humans^1,2^ and from embryonic days (E)9.5-E12.5 in mice, exhibiting remarkable similarities between the two species^3–5^. Specifically, during the 4^th^ week of human development (E9.5-E10.5 in mice), the frontonasal prominence (FNP) emerges and divides into the medial (MNP) and lateral nasal (LNP) prominences, which subsequently fuse with the maxillary prominence (MxP) around the 7^th^ week in humans^1,2^ (E11.5 in mice). The prominence cores mainly comprise mesenchymal cells that originate from cranial neural crest (CNC), encased in an epithelial layer of surface ectodermal and neuroectodermal origin^6–8^. In mice, MNP, LNP, and MxP convergence creates an epithelial seam named the lambdoidal junction (λ)^9^. Epithelial cells at the murine λ must be removed to enable seamless merging of the prominence mesenchymal cores from E11.5-E12.5. Prominence fusion shapes the midface, which consists of the nose, upper lip, and primary palate^1^. While CNC-derived mesenchymal cells have been extensively studied for their role in craniofacial morphogenesis^10–19^, previous reports from our^20,21^ and other labs^22–24^ highlighted the essential contribution of the epithelium to this process.

Developmental abnormalities like orofacial clefting (OFC) occur when facial prominences fail to merge. OFC affects approximately 1 in 1000 live births^25^, leading to significant medical challenges in patients and requiring numerous surgical corrections^26^. Our work has demonstrated that mouse embryos with constitutive compound loss-of-function (LOF) of *Pbx* genes, encoding homeodomain transcription factors (TFs)^27^, exhibit fully penetrant OFC. Notably, OFC is also present in all embryos with *Pbx1* conditional loss in the cephalic epithelium on a *Pbx2* or *Pbx3* deficient background^20,21^, underscoring the importance of the epithelium in facial prominence fusion. Interestingly, humans with *PBX1* mutations exhibit a developmental syndrome with facial dysmorphology^28^, and GWAS analysis has revealed an association between *PBX1*/*PBX2* variants and OFC^29^. We previously reported that PBX TFs mediate facial prominence fusion by either activating a WNT-p63-IRF6 regulatory module that results in apoptosis^20^ or regulating epithelial removal at the λ seam through epithelial-to-mesenchymal transition (EMT)^21^. However, only about 50% of epithelial cells are eliminated through apoptosis or EMT at the seam^20,21^, suggesting that the epithelium is heterogenous, and additional cellular behaviors contribute to epithelial removal. Actomyosin contractility^30^ and cell extrusion^31^, crucial in secondary palate fusion, as well as other cellular behaviors yet to be uncovered, may also mediate epithelial cell removal during facial prominence fusion.

Despite the prevalence of human OFC, our knowledge of the underlying cellular and molecular mechanisms is primarily based on mouse models, which have significantly informed craniofacial development^4,32–34^. Nevertheless, deeper mechanistic understanding of facial morphogenesis and congenital abnormalities in our species necessitates studies in humans. Here, we identified a novel cephalic epithelial cell population, which mediates facial prominence fusion, and is arrested in the cell cycle, that we named the Zippering Lambda (ZL) cluster in mouse embryos. Importantly, comparative analysis revealed the presence of a ZL-like population in human embryos. Both *Pbx1/2* and *p63* mutant mice with OFC showed cell cycle release in the ZL epithelium. Intersection of ZL-enriched genes with whole genome sequencing (WGS) datasets from OFC trios further uncovered OFC-associated gene variants, including mutations affecting *ZFHX3*, which encodes a TF that mediates cell cycle arrest^35–37^. Notably, *Zfhx3* deletion in mouse cephalic epithelia resulted in OFC, demonstrating that *Zfhx3* mutations cause this craniofacial disorder in mammals. Lastly, using biochemical, genetic, and –omics approaches, we found that PBX1-ZFHX3 genetically interact during midfacial morphogenesis, and function within a protein complex in the midface to co-regulate cell cycle inhibitor genes.

Our studies uncover a novel cephalic epithelial cell cycle-arrested population at the λ and reveal identical cellular behaviors and transcriptomic signatures in mice and humans during early facial morphogenesis. This research also demonstrates functional involvement of this cell cluster in facial prominence fusion, linking it to a common craniofacial disorder. Together, these investigations highlight novel mechanisms shaping the face.

## Results

### Midface λ epithelium comprises multiple cell clusters with distinct transcriptomes

To dissect the heterogeneity of the cephalic epithelial cells that envelop the facial prominences during their fusion, we conducted single-cell (sc) RNAseq^38–40^. By fluorescence-activated cell sorting (FACS), we purified EpCAM+^41^ epithelial cells from micro-dissected λ tissue of wild-type mouse embryos at developmental stages E9.5, E10.5, and E11.5, encompassing the critical timepoints of facial prominence fusion (Fig. 1A-A’’’; Fig. S1A). Unsupervised clustering^42^ after cell cycle regression and marker gene identification revealed dynamic changes in cell subpopulations across the examined timepoints. By integrating the E9.5 to E11.5 datasets, we detected fourteen unique cell clusters across timepoints, including oral cavity epithelia (OR; *Pitx1+* and *Pitx2+*)^43^; olfactory epithelia (OF; *Sox2+* and *Dlk1+*)^44^; early eye surface epithelia (EE; *Hmx1+* and *Sfrp2+*)^45,46^; prominence surface epithelia 1 and 2 (PS1 and PS2; *Wnt6+* and *Tfap2b+*)^24^; nasolacrimal duct epithelia (ND; *Aldh1a3+* and *Sox9+*)^47,48^; periderm (PD; *Grhl3+* and *Tacstd2+*)^49^; zippering λ epithelia (ZL; *Bambi+* and *Bmp4+*)^50^; neural stem cells (NSC; *Sst+* and *Nrsn1+*)^51^; olfactory neurons (ON; *Tubb3+* and *Neurod1+*)^52^; neural crest cells (NCC; *Sox10+* and *Foxd3+*)^53^; mesenchyme-like cells (ME; *Prrx1+* and *Twist1+*)^54^; forebrain cells (FB; *Fez1+* and *Zic1+*)^55,56^; and blood cells (B; *Hba-x+* and *Hbb-y+*)^57^ (Fig. 1B; Fig. S1B-E). The presence of the latter cell clusters (NCC, ME, FB, B) reflects a relaxed gating strategy during FACS purification to maximize yield of epithelial cells. Notably, the PS cluster comprises PS1 and PS2 subpopulations, grouped based on shared transcriptomes, but separated by cell cycle phases. Specifically, based on enrichment of known markers, we annotated eleven, fourteen, and eleven distinct clusters at E9.5, E10.5, and E11.5, respectively (Fig. 1C).

**Fig. 1:**
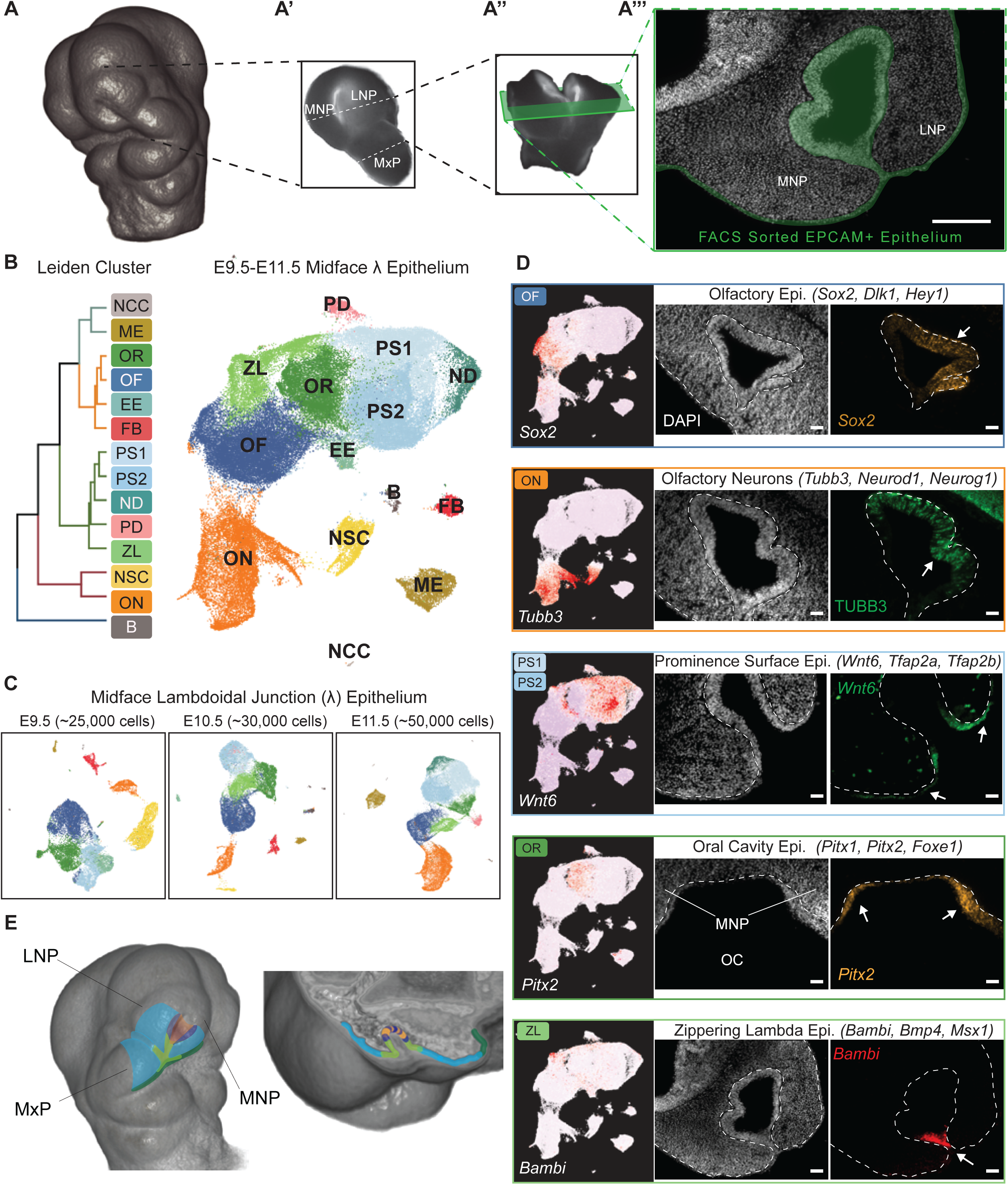
scRNAseq of midface λ epithelium highlights cell heterogeneity: spatial localization of select cell clusters. **(A-A’’’)** Experimental design of epithelial cell isolation from the lambdoidal junction (λ) of mouse embryonic facial prominences at different gestational days (E10.5 shown). **(A)** 3D rendering of an E10.5 mouse embryonic head. **(A’)** Representative microdissection of half-midface comprising medial nasal prominence (MNP), lateral nasal prominence (LNP), and maxillary prominence (MxP). Dashed lines indicating further dissection to λ-centric region. **(A’’)** Dissected λ. Green rectangle highlights cross section. **(A’’’)** Epithelial tissue within λ, isolated via fluorescence activated cell sorting (FACS) for single-cell (sc) RNAseq experiments, highlighted in green on λ midface cryosection. Scale bar: 200 μm. **(B)** Dendrogram and Uniform Manifold Approximation and Projection (UMAP) of 14 combined clusters across three timepoints. NCC, neural crest cell; ME, mesenchyme; OR, oral cavity epithelium; OF, olfactory epithelium; EE, early eye surface epithelium; FB, forebrain; PS1/PS2, prominence surface epithelium; ND, nasolacrimal duct; PD, periderm; ZL, zippering λ epithelium; NSC, neural stem cell; ON, olfactory neuron; B, blood. **(C)** UMAP representing scRNAseq datasets of E9.5-E11.5 wild-type murine λ epithelium. Captured cell numbers indicated. **(D)** Heatmap of normalized gene expression (NGE) for cluster-defining transcripts overlaid on UMAP (left panel) and RNAscope Fluorescent in-situ Hybridization (FISH) validation of the 5 largest clusters across datasets, each characterized by 3 highly enriched transcripts (center and right panels). Representative *in vivo* spatial validation of gene expression; one transcript assayed for each cluster at E11.5. DAPI stain (center) and RNAscope (right). TUBB3 visualized via immunofluorescence (IF). Dashed lines show epithelium-mesenchyme boundaries. Arrows highlight spatial enrichment of gene expression. OC, oral cavity. Scale bars: 50 μm. **(E)** Color-coded rendering of E10.5 murine facial prominences in whole head (left) and λ cross section (right) showing spatial patterns of validated cell clusters. OF, dark blue; PS1/PS2, light blue; ON, orange; OR, dark green; ZL, light green.

To delineate the spatial relationships among EpCAM+ epithelial populations, we validated the expression of one top differentially expressed gene (DEG) from each cluster by either RNAscope fluorescent *in-situ* hybridization (FISH; hereafter RNAscope)^58^ or immunofluorescence (IF) on E10.5 embryos (Fig. 1D; Fig. S1F). OF cells, marked by *Sox2*, are localized to the nasal pit together with TUBB3+ ON cells. *Wnt6+* PS cells, comprising one of the largest clusters, envelop the surface of the three λ prominences (MNP, LNP and MxP), whereas OR cells, highlighted by *Pitx2* expression, flank the oral cavity (Fig. 1D). ND, PD, and EE cells cover the nasolacrimal duct, the prominences’ apical surface, and optic vesicle, respectively (Fig. S1F). Intriguingly, a cell cluster adjoining OF, PS and OR, that we termed the zippering λ (ZL) epithelium, is uniquely positioned at the λ prominences’ tips where fusion occurs (Fig. 1D, E). This cluster is absent at E9.5 before prominence fusion begins, emerges by E10.5 during prominence fusion, and persists through E11.5 when fusion is nearly complete (Fig. 1C; Fig. S1C). Altogether, these findings underscore the heterogeneity and dynamic nature of the λ epithelium involved in facial prominence fusion and suggest potential roles for ZL cells in this process.

### Molecular characterization and *in vivo* lineage tracing of the ZL cell cluster

Considering the murine ZL cell cluster’s distinctive positioning and timing of appearance, we proceeded to explore its molecular signatures. At E9.5, prior to ZL emergence, *Bmp4*, a critical craniofacial regulator associated with OFC^59^, is expressed more broadly within the OR, OF, and PS cell clusters, as shown by normalized gene expression (NGE) overlaid on UMAPs (Fig. 2A; left). RNAscope and Whole Mount *in situ* Hybridization (WISH)^20^ localized these *Bmp4-* expressing cells to the FNP and prospective branchial arch (BA)1 junction (Fig. 2A; right) and to the rostral FNP (Fig. 2A’), respectively. By E10.5, *Bmp4* expression increases at the site of prominence fusion and is enriched in the ZL (Fig. 2B, B’). By E11.5, *Bmp4* expression expands to encompass not only the ZL but also the adjacent PS and OF cell populations, overlaying the MNP and LNP (Fig. S2A). Transcripts encoding members of signaling pathways with known roles in craniofacial development and associated with OFC, including TGF-β^60–62^, FGF^63,64^, and WNT^65–67^, are also enriched in the ZL cluster at E10.5 (Fig. 2C). We previously reported that the three above pathways converge on *Trp63* to orchestrate epithelial apoptosis during midface prominence fusion^20,21,68^ (Fig. 2C). Additionally, ZL cells show enrichment of *Bambi* and *Igfbp5*, inhibitors of TGF-β/BMP and IGF signaling^69–72^, respectively, and reduced expression of *Top2a*, a cell cycle progression marker^73^ (Fig. S2B,C). To interrogate the developmental trajectory of ZL cells in relation to adjacent cell populations in our scRNAseq datasets, we employed RNA velocity^74^, which infers cell trajectory based on the ratio of unspliced and spliced mRNAs (Fig. 2D; Fig. S2D). At E9.5, converging velocity vectors originating from the OR, OF, and PS clusters, point to the emergence of a new cell cluster (Fig. 2D). By E10.5, as fusion commences, the ZL cluster, presaged at E9.5, appears at the convergence of distinct vector trajectories stemming from these adjacent cell populations. This robust vector pattern deriving from OF and OR and directed towards the ZL cluster persists until E11.5 (Fig. 2D; Fig. S2D). CytoTRACE analysis^75^, which predicts cell differentiation states based on transcriptional diversity, corroborated these findings, identifying the ZL as the most differentiated cluster among λ epithelial cells (Fig. 2D’; Fig. S2E). Gene ontology (GO) analysis associated the top 100 DEGs in the ZL cluster with regulation of cell cycle and apoptosis (Fig. 2E). Notably, we previously implicated apoptosis in prominence fusion^20,21^.

**Fig. 2:**
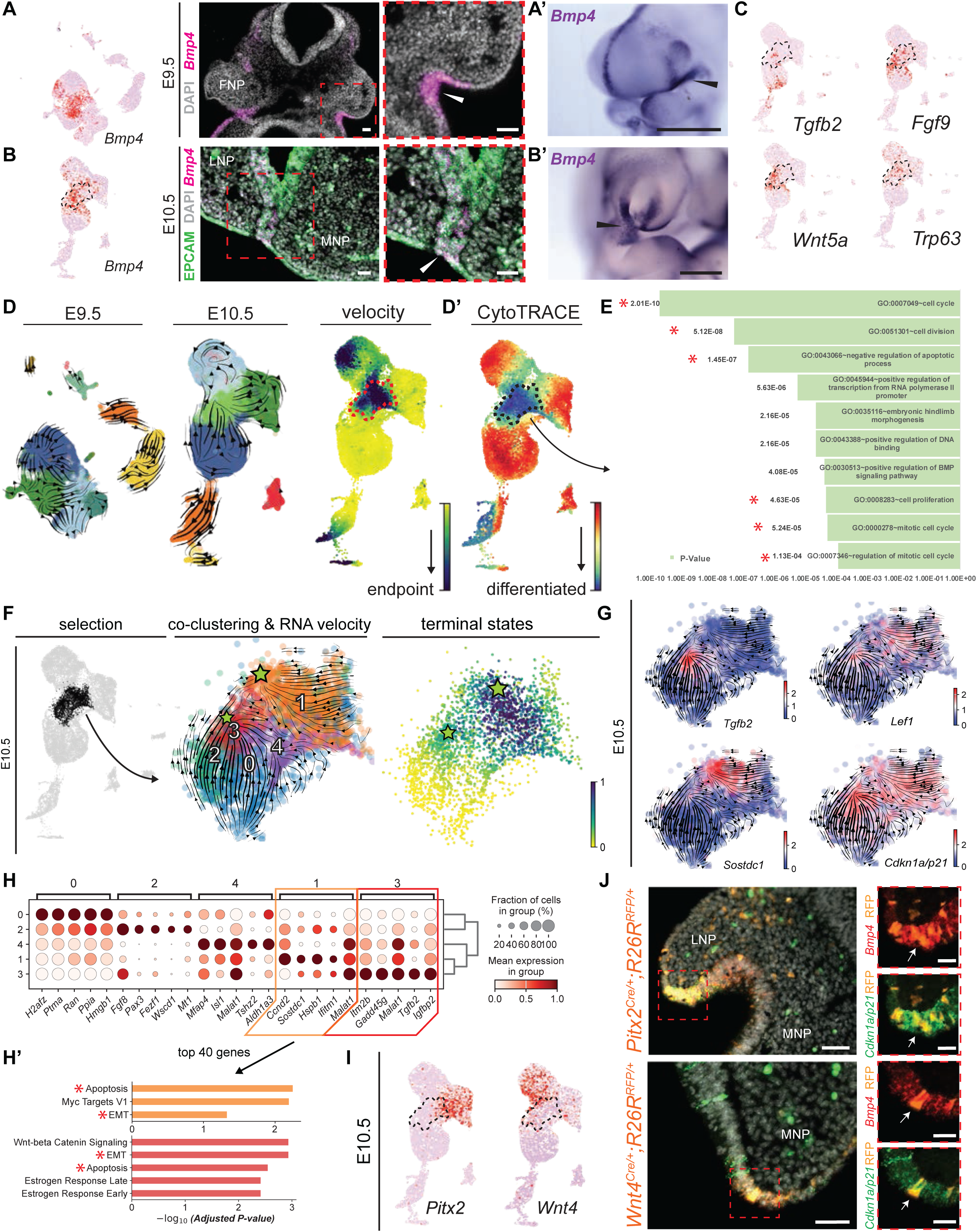
Molecular characterization and *in vivo* cell fate mapping of the ZL epithelial cell cluster. **(A, B)** Overlay of NGE on UMAP shows *Bmp4* expression at E9.5 **(A)** and E10.5 **(B)** (left). Black dashed area indicates ZL cluster boundaries. RNAscope validations in transverse section (E9.5) and coronal section (E10.5; center). Magnified insets (right; red dashed squares). White arrowheads highlight gene expression. Scale bars: 50 μm. **(A’, B’)** Whole Mount *in situ* Hybridization (WISH) validation of *Bmp4* expression at E9.5 (**A’**, sagittal view) and E10.5 (**B’**, frontal view). At E10.5, *Bmp4* expression (black arrowheads) marks ZL cluster at fusion site. Scale bars: 500 μm. **(C)** NGE overlaid on UMAPs highlighting ZL enrichment of transcripts previously associated with midface development and cleft lip/palate at E10.5. **(D)** RNA velocity of E9.5 (vectors, left) and E10.5 (vectors, middle; endpoints, right) scRNAseq datasets. ZL cluster, red dashed area. **(D’)** CytoTRACE computation of E10.5 scRNAseq dataset highlights ZL (black dashed area) amongst the most differentiated cell clusters. **(E)** Gene ontology (GO) biological processes associated with top differentially expressed genes (DEGs) in the ZL cluster. Asterisks indicate terms of interest. **(F)** Reclustering of E10.5 ZL (black-colored area) reveals 5 subclusters (ZL0-4). RNA velocity indicates two terminal endpoints (green stars) in ZL1 and ZL3 corresponding to highly differentiated cell states. **(G)** NGE overlaid on UMAPs for transcripts enriched in terminal endpoints of E10.5 ZL subclusters at E10.5. **(H)** Dot plot showing expression of top 5 DEGs in ZL subclusters. ZL1 (orange box) and ZL3 (red box) comprise RNA velocity-predicted endpoints. Dot sizes represent the proportion of cells within a given population expressing the target gene; color intensities indicate average expression levels. **(H’)** GO terms of top 40 enriched genes in ZL1 and ZL3. Asterisks indicate terms of interest. **(I)** NGE overlaid on UMAPs showing enrichment of *Pitx2* and *Wnt4* in OR (left) and PS (right) cluster, respectively, at E10.5. Black dashed area highlights ZL. **(J)** Genetic lineage tracing validates RNA velocity predictions of ZL cluster origin from OR and PS epithelium. RNAscope validation showing *Bmp4/Cdkn1a* co-expression in RFP positive cells (left). Arrows within insets highlight signal overlap within prominence tips (right, red dashed squares). Scale bars: 50 μm. Inset scale bars: 20 μm.

Further re-clustering of the ZL identified five subclusters (ZL0-4) at E10.5, with ZL1 and ZL3 as endpoints in RNA velocity analysis (Fig. 2F). ZL1 and ZL3 are enriched in transcripts that encode signaling effectors we previously implicated in apoptosis at the λ (e.g., *Tgfb2* and *Lef1*^20,21^), an antagonist of WNT/BMP signaling (*Sostdc1*^76,77^), and a marker of cell cycle arrest (*Cdkn2a/p21*^78,79^) (Fig. 2G). GO analysis of the top 40 DEGs from these subclusters revealed that these genes are highly associated with both EMT and apoptosis, cell behaviors we previously reported to mediate prominence fusion^20,21^ (Fig. 2H, 2H’). Reclustering of the ZL at E11.5 identified the same 5 subclusters as at E10.5. RNA velocity showed that the ZL population converged into ZL3 and ZL4 at this timepoint, maintaining strong expression of *Tgfb2* and *Cdkn1a/p21* with reduced *Lef1* and *Sostdc1* expression. ZL4 was enriched in *Isl1* and depleted in *Pax3*, both implicated in craniofacial development^80,81^ (Fig. S2F, G). Compositional analysis indicated a marked decrease in ZL1 and an increase in ZL3 within the ZL from E10.5 to E11.5, highlighting the dynamic evolution of these subpopulations (Fig. S2H). To validate the RNA velocity predictions of neighboring epithelial populations converging into the ZL, we conducted lineage tracing experiments using *Cre*-recombinase-expressing mouse lines for *Pitx2*^82^ (enriched in OR; Fig2I, left) and *Wnt4*^83^ (enriched in PS; Fig2I, right) crossed with an RFP reporter line^84^. By E10.5, embryos obtained from these crosses displayed distinct RFP+ cells, colocalizing with ZL markers *Bmp4* and *Cdkn1a/p21*, at the prominence tips (Fig. 2J). Taken together, our findings demonstrate that the ZL originates from the adjacent epithelial clusters OR and PS, respectively, is the most differentiated cell cluster, is enriched in genes essential for prominence fusion as well as apoptotic and EMT regulators, and is implicated in cell cycle regulation.

### The majority of ZL cluster cells are arrested in G0/G1 during facial prominence fusion

To elucidate epithelial cell division dynamics at the prominence fusion site, we assigned cell cycle gene expression scores^85^ to E9.5 – E11.5 scRNAseq datasets (Fig. 3A). At E9.5, prior to ZL emergence, all major cell clusters exhibited G2M/S^86,87^ phase activity, indicating their proliferative states. At E10.5 and E11.5, in contrast to the adjacent clusters, the ZL showed absence of G2M/S^86,87^ scoring, suggesting cell cycle arrest. This observation was supported by the ZL exhibiting the lowest S-score among the neighboring clusters at E10.5 (Fig. 3A). Confirming these findings *in vivo*, phosphorylated histone H3 (pHH3) staining, indicative of cells in late-G2/M^88^, was reduced within the ZL *versus* the adjacent mesenchyme and OF cluster at E10.5 (Fig. S3A). Moreover, bromodeoxyuridine (BrdU) incorporation, which marks DNA synthesis during S^89,90^, was not detected in the ZL after a 1-hour labeling period at both E10.5 and E11.5 (Fig. 3B; Fig. S3B). Even after a 7-hour chase, BrdU incorporation in ZL cells remained negligible compared to surrounding cells, underscoring the absence of cell cycle progression (Fig. S3C). Transcripts of *Mki67*, a marker of active G1/S/G2/M phases that is absent in the quiescent G0 and early G1 phases^91^, were significantly reduced in the ZL at both E10.5 and E11.5 (Fig. 3C; left), which was confirmed by decreased KI67 protein levels at E11.5 (Fig. 3C, right). Additionally, *Cdk1* and *Aurkb* transcripts, associated with G2/M^86,87,92^, were downregulated in the ZL (Fig. S3D, top left), and significantly reduced *in vivo* within E10.5 (Fig. 3D; Fig. S3E) and E11.5 (Fig. S3F) prominence tips, both adjoining and apart. Other cell cycle-related genes^86,87,92^, including *Anln* (M), *Pcna* (S)*, Cdc6* (S)*, Cdca8* (G2/M)*, Mcm7* (G1/S)*, Gins2* (S)*, Ccne2* (G1/S)*, Ccnd1* (G1/S)*, Ccna2* (S/G2)*, Ccnb1* (G2/M)*, E2f1* (G1/S), and *Cdc20* (M), were either absent or downregulated in the ZL cluster at E10.5 (Fig. S3D). The collective downregulation of these genes, coupled with the lack of pHH3 and KI67 staining, along with reduced BrdU incorporation, points to cell cycle arrest in the ZL during prominence fusion.

**Fig. 3:**
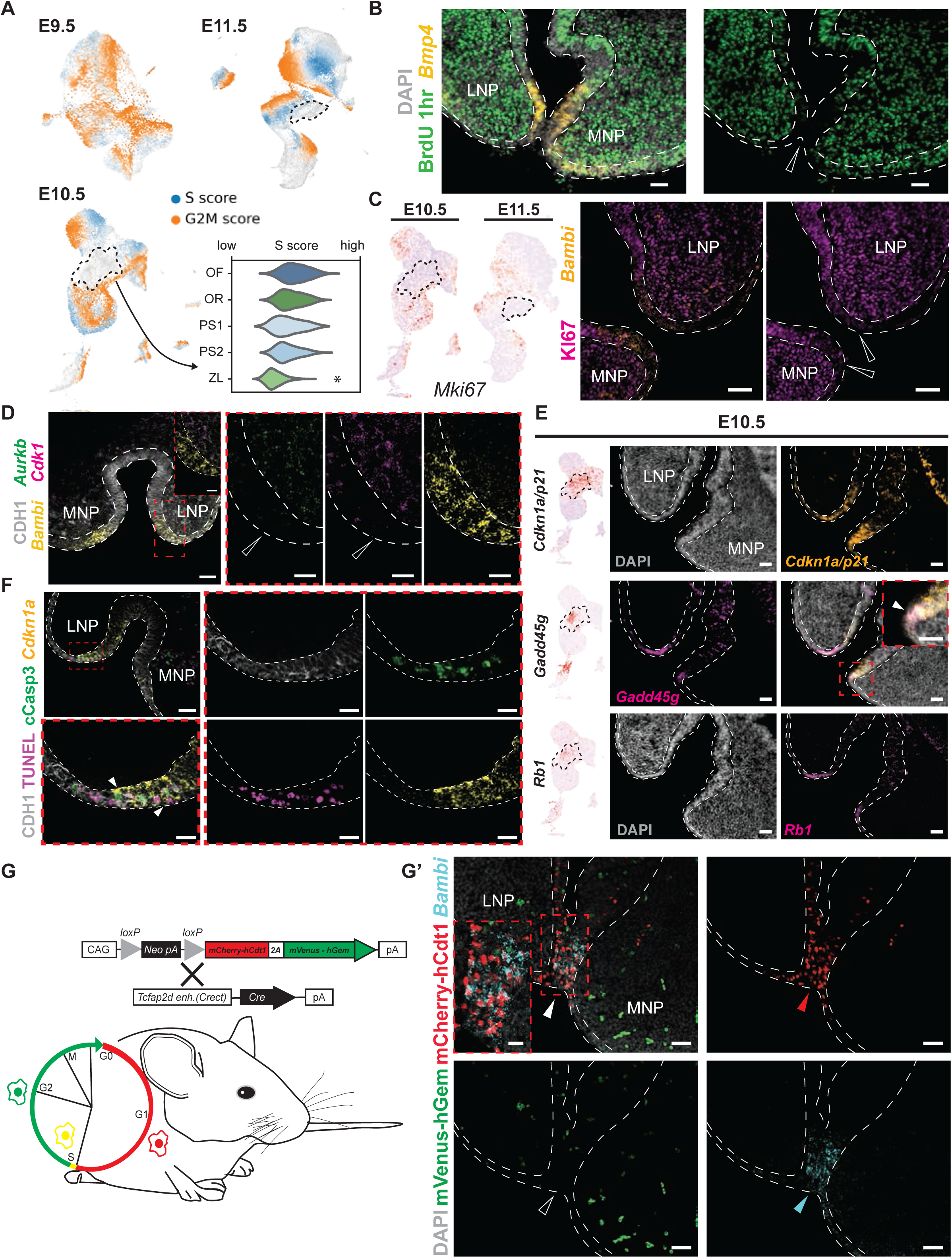
The ZL is devoid of cell cycle progression markers, enriched for cell cycle inhibitors, and arrested in G0/G1. **(A)** Cell cycle phase assignment depicted on UMAP embedding of scRNAseq cell clusters from E9.5–E11.5. Violin plot of scvelo S-phase scoring for ZL and neighboring clusters at E10.5 (t-test * p-value < 0.0001). Black dashed areas indicate ZL cluster boundaries. **(B)** BrdU incorporation revealed with anti-BrdU antibody (green) after 1 hour chase at E11.5. White dashed lines define epithelium boundaries. Scale bars: 50 µm. **(C)** NGE overlaid on UMAP shows reduced transcripts for cell proliferation marker *Mki67* in E10.5 and E11.5 ZL cluster (left). IF reveals absence of KI67 (pink) from murine ZL at E11.5 (right). Empty arrowheads point to reduced signal in ZL. Scale bars: 50 µm. **(D)** RNAscope reveals downregulation of *Cdk1* (pink) and *Aurkb* (yellow) from murine ZL at E10.5. Red dashed squares show magnified insets. Empty arrowheads point to absence of signal in ZL cell cluster. Scale bars: 50 µm. Inset scale bars: 20 µm. **(E)** UMAP of cell cycle arrest gene transcripts (*Cdkn1a/p21*, *Gadd45g*, and *Rb1*) enriched in the ZL cluster as shown by NGE overlaid on UMAPs (left) and RNAscope validations (middle, right) at E10.5. Red dashed square displays magnified inset with merged signal of *Cdkn1a* and *Gadd45g*. Scale bars: 50 µm. **(F)** Overlap of apoptosis (Cleaved Caspase-3 IF, cCasp3; terminal deoxynucleotidyl transferase dUTP nick end labeling, TUNEL) and cell cycle arrest (*Cdkn1a/p21*, RNAscope) markers, at E10.5. Scale bars: 50 µm. Inset scale bars: 20 µm. **(G, G’)** *Fucci2a* mouse system used to selectively label cell cycle phases in the cephalic epithelium **(G)**, highlights accumulation of ZL cells in G0/G1 at E11.5. IF for cell cycle markers (mCherry, G0/G1; mVenus, S/G2/M) demonstrates enrichment of cells in G0/G1 and absence of cells in S/G2/M within the ZL **(G’)**. White dashed lines define epithelium boundaries. Presence or absence/reduction of signal is highlighted by full or empty arrowheads, respectively, at the fusion site. Scale bars: 50 µm. Inset scale bars: 20 µm. *Bmp4, Cdkn1a,* and *Bambi* expression labels ZL cells across panels. CDH1 protein marks epithelial cells in panels D and F.

Cell cycle regulation depends on the synchronized interplay between promoting and inhibitory factors like cyclins and cyclin-dependent kinases^93,94^. The absence of cell cycle progression markers in the ZL prompted explorations of cell cycle inhibitors in this population. Our scRNAseq data revealed that *Cdkn1a/p21*, encoding a cyclin-dependent kinase inhibitor that inhibits the G1/S and G2/M transitions^78,79^, is significantly enriched in the ZL (Fig. S1E). WISH throughout prominence fusion stages visualized *Cdkn1a/p21* spatio-temporal expression dynamics (Fig. S4A). *Cdkn1a/p21* becomes progressively more restricted from the FNP and prospective BA1 junction at E9.5, to marking the adjoining LNP, MNP, and MxP by E10.5, and to labeling only the fusing prominence tips by E11.25 (Fig. S4A). In parallel, *Gadd45g*, encoding a GADD45 family protein implicated in cell cycle arrest^95^, exhibits similar expression dynamics (Fig. S4B). NGE overlaid on UMAPs from scRNAseq datasets confirmed restricted expression of *Cdkn1a/p21* and *Gadd45g* in the ZL at E10.5 and E11.5 (Fig. 3E; Fig. S4C; top two rows, left). RNAscope corroborated the WISH findings, showing enriched expression of both transcripts at the prominence tips (Fig. 3E; Fig. S4C; top two rows, right). Similar expression enrichment in the ZL was confirmed for *Rb1*, encoding a transcriptional corepressor that inhibits cell cycle progression^79,96^, validated by RNAscope at the prominence tips of E10.5 and E11.5 embryos (Fig. 3E; Fig. S4C; bottom row). While detected at negligible levels in the scRNAseq datasets, other cell cycle inhibitor genes, such as *Cdkn2a/p16*^97^*, Cdkn2b/p15*^98^, and *Cdkn1c/p57*^87,97^, are also expressed at the prominence tips (Fig. S4D, E). Instead, *Cdkn1b/p27*^87,97^ transcripts are present across all λ epithelial populations (Fig. S4D, E). Notably, functional roles of cell cycle inhibitors in the ZL are highlighted by the co-localization of *Cdkn1a/p21* transcripts and proteins at the prominence tips (Fig. S4F). Consistent with our previous reports that apoptosis occurs in a subset of epithelial cells at the λ during prominence fusion^20,21^, TUNEL staining^99^ and IF for cleaved-caspase 3^100^ showed that only a subset of *Cdkn1a/p21*+ ZL cells express apoptotic markers, indicating that not all cell cycle-arrested ZL cells undergo apoptosis at E10.5 (Fig. 3F). Collectively, these findings indicate that ZL cells are likely arrested in G0/G1, as demonstrated by lack of proliferative markers and enrichment of cell cycle inhibitors. To orthogonally validate ZL G0/G1 arrest *in vivo*, we employed the *Fucci2a* mouse model, which fluorescently labels cells in different cell cycle phases^101^. To specifically mark cephalic epithelial cells, we crossed *Fucci2a* mice with the *Crect-cre* line^102^ (Fig. 3G). E11.5 embryos showed an accumulation of mCherry+ cells (denoting G0/G1) at the prominence tips (Fig. 3G’), which was already apparent at E10.5 (Fig. S4G). The presence of mCherry+ cells, which colocalized with the ZL marker *Bambi*, and the absence/reduction of mVenus+ cells (denoting G2/M) at E11.5 and E10.5 strongly support that the majority of ZL cells are arrested at G0/G1 (Fig. 3G’, Fig. S4G).

### ZL cell cycle arrest is disrupted in different OFC mouse models

To examine whether the ZL is perturbed in OFC models, we studied mouse embryos with epithelial-specific LOF of *Pbx1* on a *Pbx2*-deficient background (*Crect^Cre/+^;Pbx1^f/f^;Pbx2^+/-^;* named *Pbx1/2* mutants)^20,21,103,104^ or with *p63* constitutive LOF (*p63^-/-^*)^105^. Notably, *p63* is expressed only in epithelial tissues, including cephalic epithelia^20,106^. Both mutations result in OFC (Fig. 4A, B) and/or facial dysmorphology in mice and humans^20,28,29,68,107,108^. scRNAseq analysis of EpCAM+ cells from micro-dissected λ tissue of *Pbx1/2* mutant embryos and littermate controls identified 10 cell clusters at E10.5 (Fig. 4C). Furthermore, RNA velocity suggested a potential disruption in the vector patterns in the mutant embryos from surrounding epithelial clusters, particularly from PS1, toward the ZL (Fig. 4D). Cross-boundary direction correctness^109^, assessing the likelihood of a cell transitioning to a target state based on its current trajectory, pointed to an altered contribution of PS1 to ZL in mutants compared to controls, while OF and OR clusters exhibited similar patterns in both groups (Fig. 4E). Given the limitations of scRNAseq in detecting low-abundance transcripts^110^, we also conducted bulk RNAseq of FACS-purified λ epithelium from E11.5 *Pbx1/2* mutant and control embryos. Principal component analysis distinctly separated the transcriptomic profiles of mutants from controls (Fig. 4F; left), with 718 DEGs identified (Fig. 4F; right). As expected, *Pbx1* expression was downregulated in mutants. Notably, cell cycle inhibitor genes, such as *Cdkn2a/p16*, *Cdkn2b/p15*, *Gadd45g*, *Cdkn1c/p57*, and *Cdkn1b/p27*, were significantly downregulated in mutants (Fig. 4F; bottom). RNA-sieve analysis^111^, which deconvolutes bulk RNA cell samples using scRNAseq expression data, confirmed that the genes significantly dysregulated in mutants were primarily enriched within the ZL, suggesting that perturbation of this epithelial subpopulation is strongly associated with OFC (Fig. 4G).

**Fig. 4:**
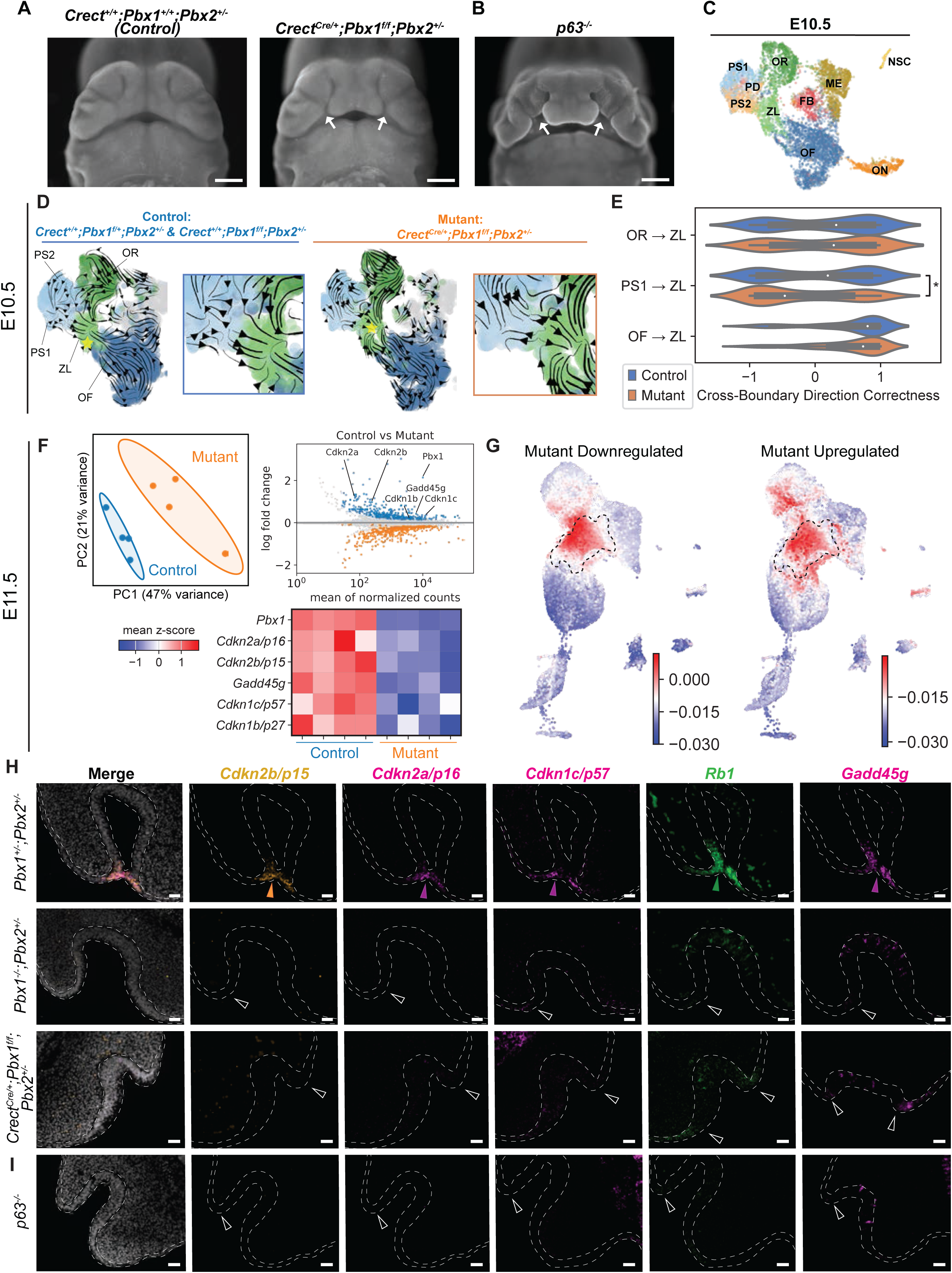
OFC mouse models with loss of *Pbx1/2* exhibit perturbation of the ZL cell cluster and, together with *p63* mutants, show dysregulation of cell cycle arrest. **(A, B)** Gross morphology of E12.5 mouse embryonic midface with epithelial-specific deletion of *Pbx1* on a *Pbx2*-deficient background (*Crect^Cre/+;^Pbx1^f/f^;Pbx2^+/-^;* hereafter *Pbx1/2* mutant) compared to littermate control (*Crect^+/+^;Pbx1^+/+^;Pbx2^+/-^*) **(A)** and *p63* constitutive LOF mutant (*p63^-/-^*) **(B)**. Arrows point to bilateral cleft lip. Scale bar: 500 µm. **(C)** UMAP of combined scRNAseq clusters from E10.5 control and *Pbx1/2* mutant λ epithelium. **(D)** scRNAseq RNA velocity from micro-dissected λ tissue of E10.5 *Pbx1/2* mutant embryos and littermate controls. Epithelial cell clusters comprising OR, OF, PS1/PS2, and ZL highlighted. Insets show boundaries between PS and ZL cluster transitions. **(E)** Cross-boundary direction correctness of RNA velocity predictions. Asterisk indicates statistical significance (p ≤ 0.05, Kolmogorov-Smirnov test). **(F)** Principal component analysis (PCA) of bulk RNAseq datasets from sorted λ epithelium of E11.5 *Pbx1/2* mutants and controls (n=4; top left). DEGs in *Pbx1/2* mutants *versus* controls highlight cell cycle inhibitor transcripts, including *Cdkn2a/p16*, *Cdkn2b/p15*, *Gadd45g*, *Cdkn1c/p57*, and *Cdkn1b/p27* (top right). These transcripts, enriched in the ZL cluster, are downregulated in *Pbx1/2* mutant *versus* control λ epithelium (heatmap, bottom right). **(G)** RNAsieve deconvolution of bulk RNAseq expression data using scRNAseq as a reference shows enrichment of top downregulated (left) and upregulated (right) genes in the ZL of *Pbx1/2* mutant λ epithelium compared to controls. Heatmap overlaid on scRNAseq of E10.5 wild-type λ epithelium. **(H, I)** Cell cycle inhibitor gene transcripts, *Cdkn2b, Cdkn2a, Cdkn1c, Rb1,* and *Gadd45g,* are reduced/absent in the ZL cluster at the fusion site in OFC mouse models, including *Pbx1/2* constitutive loss-of-function (LOF) (*Pbx1^-/-^;Pbx2^+/-^*), *Pbx1/2* mutants **(H)**, and *p63* constitutive LOF mutants (*p63^-/-^)* **(I)** *versus* controls (*Pbx1^+/-^;Pbx2^+/-^*) **(H)**, as shown by RNAscope at E10.5. Presence or absence/reduction of signal highlighted by full or empty arrowheads respectively. All scale bars: 50 µm.

To validate the predicted gene expression changes *in vivo*, RNAscope was performed on E10.5 and E11.5 mouse embryos with *Pbx1/2* constitutive LOF (*Pbx1^-/-^;Pbx2^+/-^*)^20,21,103,104^, as well as conditional *Pbx1/2* mutants (*Crect^Cre/+^;Pbx1^f/f^;Pbx2^+/-^*)^20,21^, and controls (*Pbx1^+/-^;Pbx2^+/-^*). The expression of *Cdkn2b/p15*, *Cdkn2a/p16*, *Cdkn1c/p57*, *Rb1*, and *Gadd45g*, enriched in wild-type ZL cells, was markedly reduced or absent at the prominence tips of all mutant embryos compared to controls, suggesting that cell cycle is unleashed in mutant ZL cells (Fig. 4H). Similar to prominences that are not touching, adjoining prominences of E11.5 *Pbx1/2* mutants also exhibited strikingly decreased expression of cell cycle inhibitors, including *Cdkn2a/p16* and *Cdkn2b/p15* (Fig. S5A). This finding indicates that cell cycle arrest in the ZL occurs independently of physical contact during midface morphogenesis. Similarly, *p63^-/-^*embryos exhibited marked downregulation of the same cell cycle inhibitor-encoding genes at the prominence tips at E10.5 as *Pbx1/2* mutants, pointing to disruption of cell cycle arrest in ZL cells as a general mechanism underlying OFC in two distinct models (Fig. 4I). Surprisingly, *Cdkn1a/p21* expression was unchanged in either *Pbx1/2* or *p63* mutants, highlighting the complexity of cell cycle arrest regulation that is driven by inhibitors with overlapping functions^112^ (Fig. S5B). Based on expression of cluster-defining makers *Bmp4* and *Cdkn1a/p21,* RNAscope/IF confirmed the presence of a ZL population in *Pbx1/2* mutants, characterized by disrupted cell cycle arrest. While PBX1 is absent in the epithelium, as expected, it is still detected in intercalated cells of presumptive olfactory neuronal identity (Fig. S5C). Together, these findings establish that the release of ZL cell cycle arrest in two different OFC models is a critical and likely general mechanism in the pathogenesis of OFC.

### A cell cycle arrested ZL-like population is present in human embryos: association of *ZFHX3* with OFC

Given the similarities in early craniofacial morphogenesis between mice and humans^113^ and the common genetic etiologies of craniofacial defects, we investigated whether a ZL-like cell cycle-arrested population is present in human embryos during facial prominence fusion. In humans, this process occurs between 5-6 weeks post conception, corresponding to Carnegie stages (CS) 15-17^114,115^. In CS16 human embryos, analogous to E10.5 mice^116^, we observed a marked reduction in transcripts and proteins for the cell cycle progression markers *CDK1* and KI67, respectively, at the FNP, LNP, and MxP fusion seam, mirroring our findings in the murine ZL cluster (Fig. 5A, B; Fig. S6A). This reduction coincides with increased expression of the cell cycle inhibitor-encoding genes *CDKN1A/P21* and *GADD45G*, transcripts that defined the ZL cluster in mice (Fig. 5A, B; Fig. S6A). By CS17, human embryos, akin to E11.5 mice, exhibited similar expression patterns as at CS16, including a reduction of KI67 and *CDK1*, along with increased expression of *CDKN1A/P21* and *GADD45G* at the fusion site (Fig. 5C; Fig. S6B). These observations highlight striking similarities between mouse and human midface morphogenesis at the morphological, cellular, and molecular level at early gestation, pointing to the involvement of a ZL-like population in human prominence fusion.

**Fig. 5:**
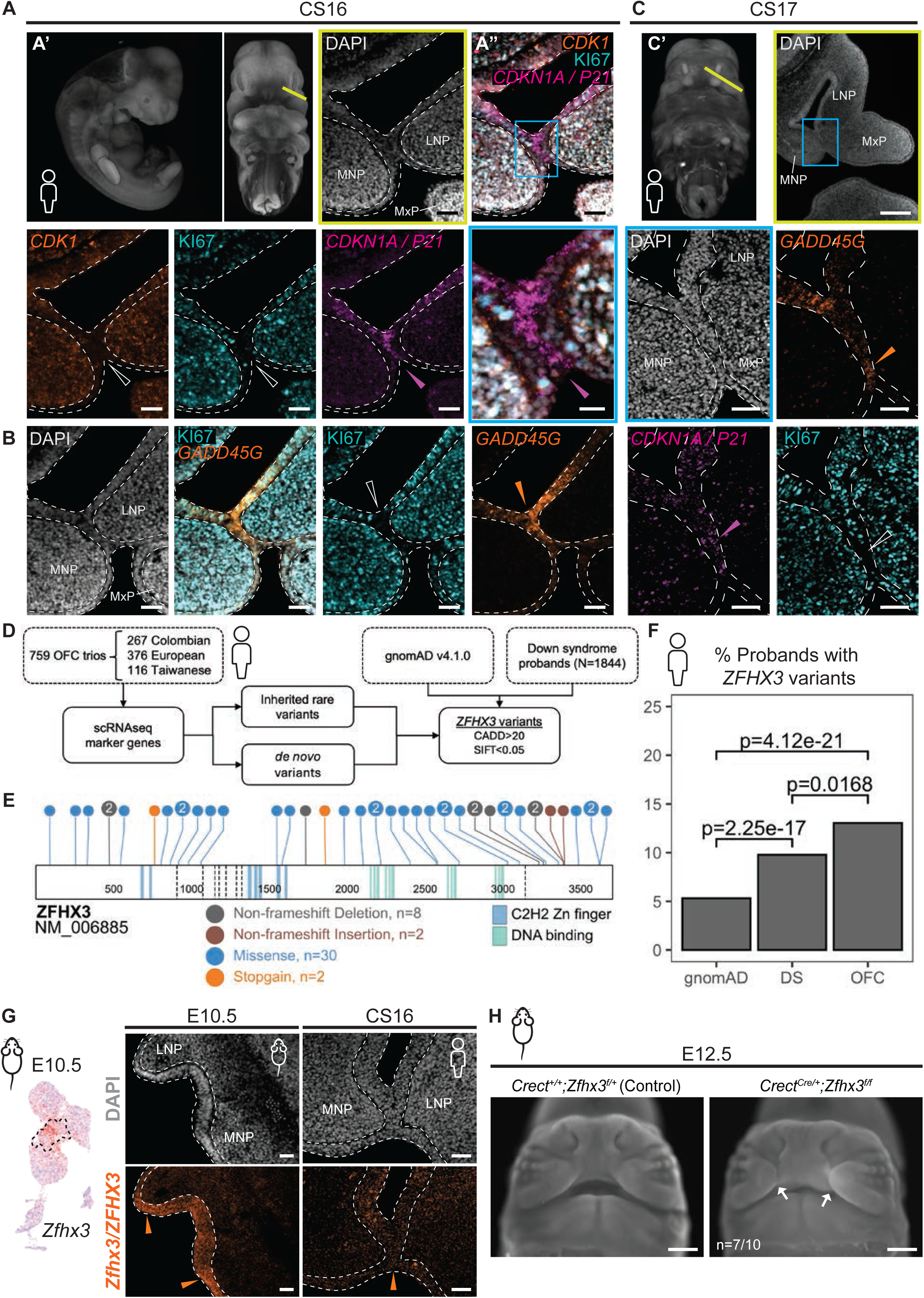
The human embryonic midface comprises an epithelial ‘ZL-like’ cell cycle-arrested population, which is enriched for *ZFHX3*, a gene mutated in OFCs. **(A)** Absence of *CDK1* (RNAscope) and KI67 (IF) at the facial prominence fusion site of CS16 human embryos. Increased expression of cell cycle arrest gene *CDKN1A/P21* (RNAscope). **(A’)** Representative high resolution episcopic microscopy (HREM) image of CS16 embryo in lateral (left) and frontal (right) view^160,161^. Yellow line denotes plane of section (yellow box). **(A’’)** *CDKN1A/P21* expression domain overlaps with reduced *CDK1* and KI67 (magnified inset, blue box). White dashed lines define epithelium boundaries. Presence or absence/reduction of signal highlighted by full or empty arrowheads, respectively. Scale bars: 50 µm. Inset scale bar: 20 µm. **(B)** Negligible KI67 levels (IF) coincide with domains of high *GADD45G* expression (RNAscope) at the fusion seam of the three prominences in CS16 embryos. Scale bars: 50 µm. **(C)** In CS17 human embryos, high *GADD45G* and *CDKN1A/P21* expression overlaps with low KI67 protein at the prominence fusion site. (C’) Representative HREM image of CS17 embryo in frontal view^160,161^. Yellow line indicates plane of section (yellow box, scale bar: 200 µm). Blue box shows MNP, LNP and MxP fusion seam. Scale bars: 50 µm. Whole embryo images provided by HDBR Atlas (https://hdbratlas.org). **(D)** Visualization of workflow for data analysis of 759 OFC trios subjected to Whole Genome Sequencing (WGS) and intersected with scRNAseq datasets. **(E)** Lollipop plot illustrating all rare, predicted damaging *ZFHX3* variants exhibited by OFC probands. **(F)** Bar chart showing significant differences in the percent of probands harboring a rare, predicted damaging *ZFHX3* variant in each category analyzed. OFC=orofacial cleft. DS=Down syndrome. gnomAD=genome Aggregation Database (Chi-Squared test). **(G)** NGE overlaid on UMAP shows strong *Zfhx3* enrichment in the E10.5 murine ZL cluster (black dashed area; left). In mouse (E10.5) and human (CS16) midfacial sections, *Zfhx3/ZFHX3* is highly expressed (RNAscope) in the λ epithelium. Arrowheads highlight prominence tips/fusion site. Scale bars: 50 µm. **(H)** Gross morphology of E12.5 mouse embryonic midface with *Zfhx3* epithelial-specific deletion (*Crect^Cre/+^;Zfhx3^f/f^*) compared to littermate control (*Crect^+/+^;Zfhx3^f/+^*). Arrows point to bilateral cleft lip. Scale bar: 500 µm.

To investigate potential associations between genes enriched in the murine ZL cluster and gene variants linked to human OFC, we intersected our scRNAseq dataset from murine λ epithelium with datasets of whole genome sequencing (WGS) from 759 OFC case-parent trios^117^ (Fig. 5D). Our analyses included both inherited rare variants and *de novo* variants (MAF <0.5%) with protein-altering consequences (*i.e.,* not synonymous). We then focused on variants within constrained genes (pLI > 0.9 and LOEUF < 0.35) from the ZL cluster with high pathogenicity prediction scores (SIFT < 0.05 and CADD > 20). As expected, we found qualifying variants in several genes associated with OFCs, such as *BMP4*^59^ (N=14) and *TP63*^108^ (N=9) (Fig. S6C; Supplemental Table 1). Interestingly, the highest number of variants was found in Zinc-finger homeobox 3 (*ZFHX3* or *ATBF1*; N=42), including one truncating *de novo* variant (Fig. 5E; Fig. S6C). ZFHX3, encoding a transcription factor associated with neuronal^118,119^ and cardiac^120^ function, has been implicated in the control of cell cycle arrest and transactivation of *p21*^35–37^. In humans, previous work highlighted *ZFHX3* as a candidate risk gene for OFC^121^, and its variants were linked to a neurodevelopmental disorder occasionally accompanied by cleft palate^119^. However, a causative role for *ZFHX3* in OFC pathogenesis remained unexplored.

### *ZFHX3/Zfhx3* is required for facial prominence fusion in mammals

As ZFHX3 encodes a large protein comprising 3703 amino acids, we used additional cohorts to determine whether the large number of variants was a function of its size, or if it had biological relevance. We thus compared the percent of OFC probands (N=759) with *ZFHX3* variants to trisomy 21 (T21) probands^122^ (N=1844), and to individuals in gnomAD v.4.1.0^123^, in order to test if mutation burden was higher for OFCs. We found a statistically significant difference between all three groups, where OFC probands carried the highest burden of variants (Fig. 5F) at 13.0% compared to 9.8% in T21 and 5.3% in gnomAD. Given this evidence supporting roles of *ZFHX3* in OFC etiology, we further investigated its function in mouse embryos. *Zfhx3* was enriched in the ZL cluster at both E10.5 and E11.5 (Fig. 5G, left; Fig. S6D, right). Interestingly, *Zfhx3* enrichment was already present at E9.5, before the emergence of the ZL (Fig. S6D; left), at the converging point of the RNA velocity trajectories (see Fig. 2D; Fig. S2E). RNAscope confirmed *Zfhx3/ZFHX3* expression in the λ epithelium during prominence fusion in both E10.5/E11.5 mouse and CS16 human embryos (Fig. 5G right; Fig. S6E). Notably, *Zfhx3* displayed significant enrichment in the ZL terminal endpoints ZL3 and ZL4 at both E10.5 and E11.5, suggesting potential roles in mediating ZL cell dynamics (Fig. S6F). To unequivocally assess the requirement of epithelial *Zfhx3* in prominence fusion, we generated mouse embryos with epithelial-specific *Zfhx3* deletion^124^ (*Crect^Cre/+^;Zfhx3^f/f^*; named *Zfhx3* mutants). By E12.5, post prominence fusion, 7 out of 10 homozygous *Zfhx3* mutants (70%) exhibited OFC, in contrast to their non-cleft heterozygous and littermate controls (Fig. 5H; Fig. S6G). Overall, our findings reveal a ZL-like cell population in humans, sharing transcriptomic signatures in spatial domains overlapping with the murine ZL during prominence fusion. Human *ZFHX3*, identified by WGS as the gene with the highest number of *de novo* variants in OFC patients, is enriched in the murine ZL and human λ epithelium. Lastly, epithelial-specific *Zfhx3* loss in the mouse results in embryonic OFC, establishing its requirement in ZL for mammalian facial prominence fusion.

### ZFHX3 and PBX1 synergistically regulate cell cycle arrest during prominence fusion

Our findings above, coupled with previously reported roles of ZFHX3 in mediating cell cycle inhibition, make this transcription factor an ideal candidate to drive facial prominence fusion by regulating cell cycle arrest. To investigate this possibility, we performed chromatin immunoprecipitation and sequencing (ChIPseq)^125^ for ZFHX3 on E11.5 whole murine λ tissue. ChIPseq analysis identified ZFHX3 binding at promoters of key cell cycle arrest genes that are significantly enriched in the ZL during prominence fusion, including *Rb1* and *Cdkn1c/p57*, as well as a putative *Cdkn2a/p16* regulatory region (Fig. 6A; Fig. S7A). Additionally, ZFHX3 was found to bind its own promoter, a feature common among transcription factors^125–127^ (Fig. S7A). ZFHX3-bound regions were also marked by H3K27ac in the λ epithelium (our dataset) and p300 peaks (GSM1199037; dataset from Attanasio et al. 2013^128^), indicative of active enhancers and promoters^129,130^, and corresponded to open chromatin regions, as shown by ATACseq^131^ (GSE199339; dataset from Van Otterloo et al. 2022^24^). All datasets were obtained from mouse embryonic tissues: ATACseq on E11.5 cephalic surface ectoderm^24^; p300 ChIPseq on E11.5 craniofacial tissue^128^; and our H3K27ac ChIPseq on E11.5 murine λ epithelium. Given the role of epithelial PBX in murine facial prominence fusion^20,21^ and the altered expression of cell cycle gene in *Pbx1/2* mutant murine ZL, we also performed PBX1 ChIPseq on purified E11.5 murine λ epithelium. These experiments revealed overlapping binding of PBX1 and ZFHX3 at *Rb1* and *Cdkn1c/p57* promoters, as well as at a *Cdkn2a/p16* adjacent non-coding region (Fig. 6A; Fig. S7A). Notably, the transcripts for all above genes were significantly downregulated in *Pbx1/2* mutant ZL (see Fig. 4F, H). ZFHX3 also bound both the *Pbx1* and *Pbx3* promoters, while PBX1 did not bind the *Zfhx3* promoter, suggesting a potential regulatory hierarchy whereby ZFHX3 may direct *Pbx* gene transcription (Fig. 6A; Fig. S7A). Analysis of the putative regulatory elements interacting with these transcription factors showed that PBX1 and ZFHX3 predominantly associated with distal elements when bound independently, with 11,915 peaks and 31,391 peaks, respectively. However, regions co-bound by both PBX1 and ZFHX3 (403 peaks) are primarily associated with promoters, indicating selective co-regulation (Fig. 6B). Genomic Regions Enrichment Annotation (GREAT)^132^ analysis revealed that these co-bound sites were linked to cell cycle genes, such as cyclin-dependent kinase regulators, and associated with phenotypes related to cellular senescence, a permanent cell cycle arrested state^133,134^ (Fig. 6C). This association was not evident in regions bound solely by either ZFHX3 or PBX1. PBX1 bound-sites were linked to developmental processes including stem cell maintenance and olfactory placode morphogenesis, while ZFHX3 bound-sites were primarily associated with epigenetic regulation (Fig. 6D). To ensure robustness in our ChIPseq analysis, only peaks reproducibly detected in at least two replicates were included. *De novo* motif enrichment analysis using HOMER^135^ confirmed that several of the top ZFHX3 motifs comprised AT-rich sequences, including a motif closely resembling the previously published ZFHX3 consensus^118,120^ (TATTTAATAAT) (Fig. 6E). Known motif analysis for PBX1 primarily identified binding sites for PBX family members or co-factors (Fig. 6F). Functional validation of PBX1– and ZFHX3-bound regions was performed using luciferase reporter assays in HEK293 cells^20,21,28,136^. Co-transfection of cells with expression constructs for both PBX1 and ZFHX3 significantly increased transactivation of the reporter at the *Rb1*, *Cdkn1c/p57,* and *Pbx3* promoters, compared to transactivation by empty vector, PBX1 alone, or ZFHX3 alone, indicating strong synergistic effects of the two factors on transcriptional regulation (Fig. 6G).

**Fig. 6:**
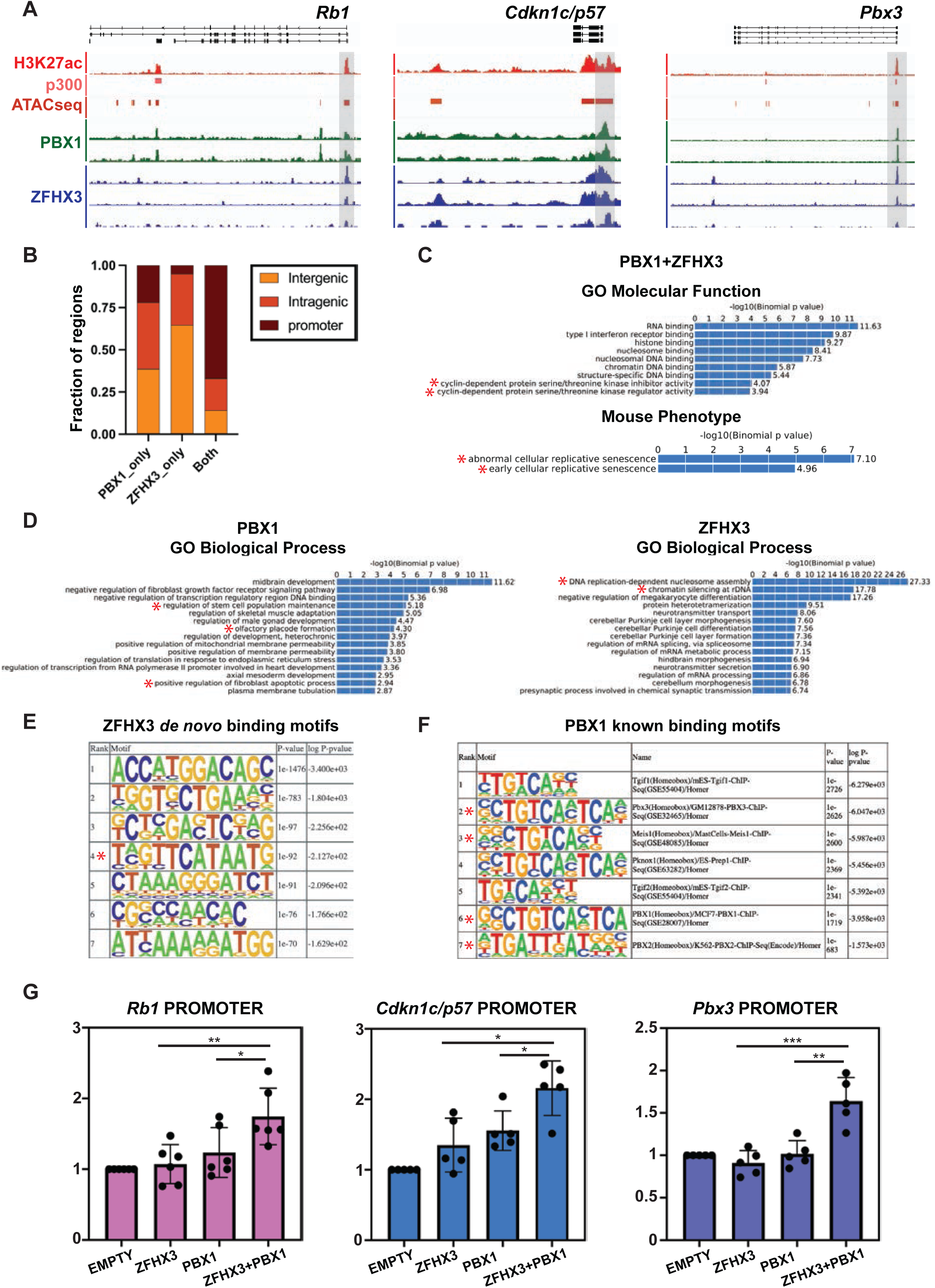
ZFHX3 and PBX1 co-regulate cell cycle arrest genes in the λ epithelium. **(A)** IGV genome browser tracks depicting the *Rb1*, *Cdkn1c/p57* and *Pbx3* loci. ChIPseq tracks: H3K27ac, red; p300 (GSM1199037), pink; PBX1 and ZFHX3, green and blue, respectively. ATACseq track (GSE199339), maroon. Gray bars highlight predicted cis-regulatory elements (CREs) bound by PBX1 and ZFHX3. **(B)** Fractions of PBX1-only, ZFHX3-only, and PBX1-ZFHX3 co-bound peaks (Both) relative to intergenic, intragenic, or gene promoter regions. **(C)** Top enriched molecular functions (top) and mouse phenotypes (bottom) associated with PBX1-ZFHX3 co-bound peaks. X-axes show the −log10 of uncorrected p-values. Asterisks highlight categories linked to cell cycle arrest. p-values from Binomial Tests. **(D)** Top enriched biological processes associated with PBX1 (left) and ZFHX3 (right) peaks, with x-axes displaying the –log10 of uncorrected p-values. Asterisks emphasize biological processes related to prominence fusion for PBX1 and epigenetic regulation for ZFHX3. p-values derived from Binomial Tests. **(E)** Top motifs identified by HOMER *de novo* motif enrichment analysis of ZFHX3-bound genomic regions, with p-values and log p-values shown. Asterisk indicates the motif closely resembling the published consensus ZFHX3 binding motif. **(F)** Known motifs identified by HOMER in genomic regions bound by PBX1; p-values and log p-values displayed. Asterisks denote motifs associated with PBX family members or PBX co-factors. **(G)** Normalized luciferase activity from transient transfections of HEK293T cells with promoter fragments of *Rb1*, *Cdkn1c/p57*, or *Pbx3* linked to a luciferase reporter and co-transfected with either empty vector or expression constructs for ZFHX3 alone, PBX1 alone, or ZFHX3+PBX1. Transactivation normalized relative to the empty vector control and depicted as fold increase. *p < 0.05, **p < 0.01, ***p < 0.001, T-test for paired comparisons. Black dots represent each biological replicate.

### A ZFHX3-PBX1 protein complex at the λ mediates facial prominence fusion in mammals

Co-immunoprecipitation of PBX1 with ZFHX3 protein from E10.5 and E11.5 murine λ tissue confirmed the association of PBX1 and ZFHX3 proteins during prominence fusion (Fig. 7A). To genetically dissect the developmental impact of the PBX1-ZFHX3 complex on facial prominence fusion, using the *CrectCre* deleter we generated compound mutant embryos in the cephalic epithelium that lacked 1) both alleles of *Pbx1* and one allele of *Zfhx3,* as well as 2) one allele each for *Pbx1*, *Pbx2*, and *Zfhx3* (Fig. 7B). Wild-type morphology was observed in embryos with epithelial-specific homozygous loss of *Pbx1* or constitutive loss of *Pbx2* alone, as well as in embryos heterozygous for both *Pbx1* and *Pbx2* as we previously reported^20,21,103^ and in embryos with *Zfhx3* epithelial-specific heterozygous loss (see Fig. S6G). In contrast, OFC was detected in compound mutants lacking both copies of *Pbx1* and one copy of *Zfhx3* in the cephalic epithelium (*Crect^Cre/+^;Pbx1^f/f^;Zfhx3^f/+^*; 1 out of 3 embryos). Further, deleting one allele each for *Pbx1, Pbx2,* and *Zfhx3* in the cephalic epithelium also led to OFC (*Crect^Cre/+^;Pbx1^f/+^;Pbx2^+/-^;Zfhx3^f/+^*; 1 out of 1 embryo) (Fig. 7B). These experiments reveal a genetic interaction of *Pbx1-Zfhx3* in facial prominence fusion. Together, our findings establish a coordinated genetic and regulatory interaction of PBX1 and ZFHX3, primarily at promoters and cis-regulatory modules of genes encoding cell cycle inhibitors. The PBX1-ZFHX3 complex comprises part of a gene regulatory network essential for cell cycle arrest at the ZL epithelial cluster during prominence fusion and midfacial morphogenesis.

**Fig. 7:**
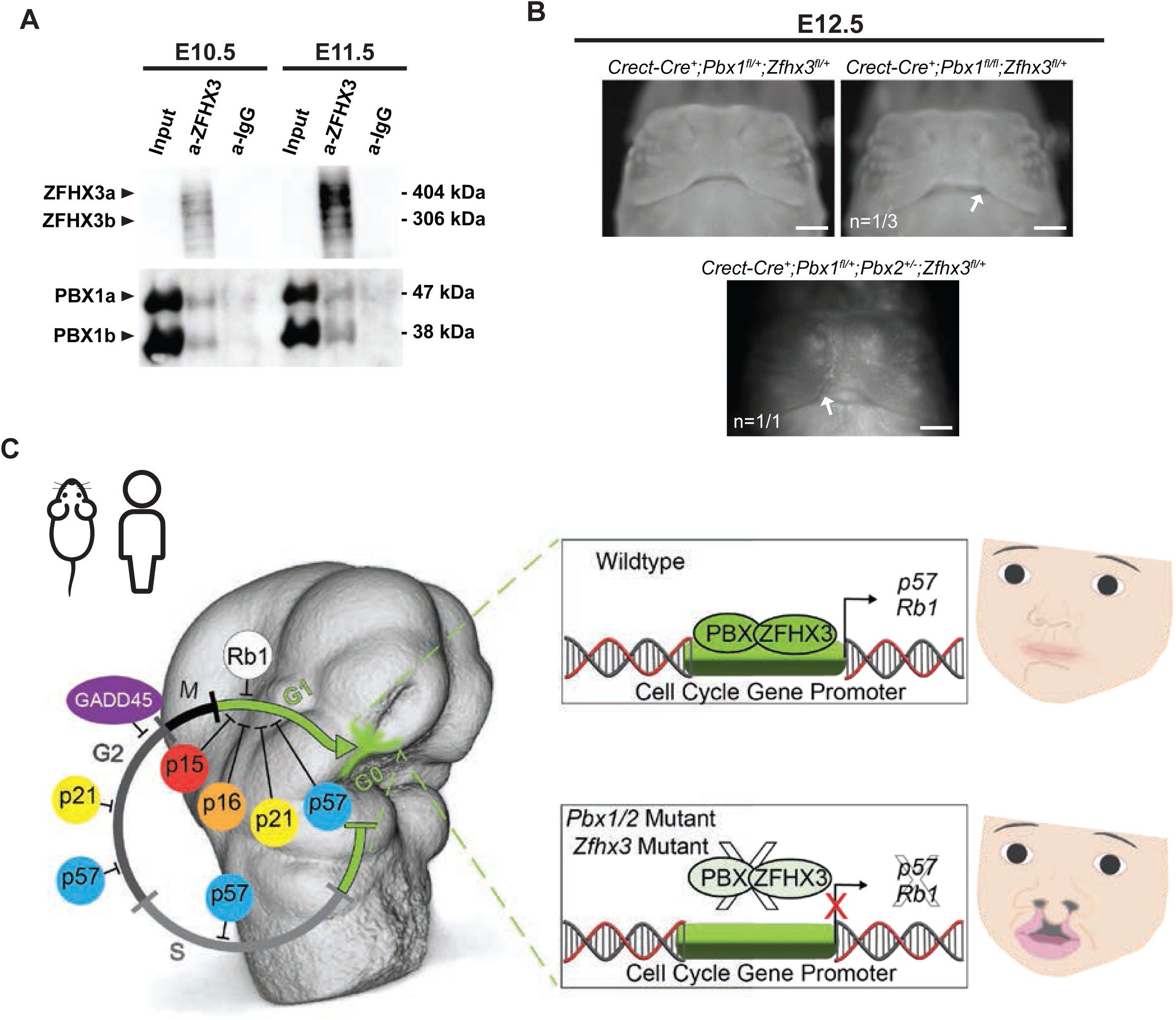
A ZFHX3-PBX1 protein complex at the λ mediates facial prominence fusion in mammals. **(A)** Immunoprecipitation of a PBX1-ZFHX3 protein complex in dissected murine λ tissues at E10.5 (left) and E11.5 (right) using ZFHX3 antibody (n=2 biological replicates). IgG used as negative control. kDa, kilodalton. PBX1a, PBX1b, ZFHX3a, and ZFHX3b are known protein isoforms^27,125,162^. **(B)** Gross morphology of E12.5 mouse embryonic midface with homozygous *Pbx1* and heterozygous *Zfhx3* epithelial-specific deletion (*Crect^Cre/+^;Pbx1^f/f^;Zfhx3^f/+^*) or triple heterozygous deletion of *Pbx1*, *Pbx2*, and *Zfhx3* (*Crect^Cre/+^;Pbx1^f/+^;Pbx2^+/-^;Zfhx3^f/+^*) compared to littermate control (*Crect^Cre/+^;Pbx1^f/+^;Zfhx3^f/+^*). Arrows point to cleft lip. Scale bar: 500 µm. **(C)** Model: a PBX-ZFHX3 complex directly regulates expression of cell cycle inhibitors, specifically *Cdkn1c/p57* and *Rb1*, in the ZL cell cluster of the developing midface. Disruption of this complex due to genetic mutations of *Pbx1/2* or *Zfhx3* results in OFC in mice. In humans, similar disruption of this complex due to *PBX1* mutations^28,29^ or *ZFHX3* mutations^119,121^ results in facial abnormalities and OFC.

## Discussion

Fusion of the craniofacial prominences shapes the human face, which identifies us. Failure of this process leads to OFC, the most common craniofacial birth defect^25^. While most studies of craniofacial development have focused on NC-derived mesenchyme^10–19^, the cephalic epithelium was also recognized as a critical population for proper morphogenesis and fusion of the prominences^20–24,137^. Previous scRNAseq of the entire murine λ tissue, comprising the mesenchyme, epithelium, and periderm, analyzed primarily the larger mesenchymal population^22,137^. By sorting λ epithelial cells, we identified a new cephalic epithelial cell cluster, that we named ZL, located at the λ prominence tips where fusion occurs. The ZL emerged as a distinct and temporally-regulated population with unique spatial location and specific transcriptomic signatures. Lineage tracing experiments provided *in vivo* validations and confirmed the RNA velocity predictions establishing that neighboring epithelial cell clusters converge onto the ZL.

Lack of G2M/S gene expression, accompanied by enrichment of cell cycle inhibitors, along with the use of *Fucci2a* mice, demonstrated that ZL cells are arrested in G0/G1. Notably, cell cycle regulation has been implicated in the development and disease of many tissues and species, relying on the synchronized interplay between factors promoting and inhibiting cell cycle progression^93,94^. Interestingly, a population enriched for *Tgfb2*, as well as the cell cycle inhibitors *Cdkn1c/p57* and *Cdkn2a/p16*, was previously identified at the prominence fusion seam^22^ but was not characterized. While only a subset of the cell cycle-arrested ZL cells undergo apoptosis at E10.5, we cannot exclude that additional cells may become apoptotic at later stages. Alternatively, these cells could transition to other cellular states such as EMT, cell extrusion, senescence, or differentiation^20,21,30,31,72,134,138^, some of which have been previously linked to primary and secondary palate fusion^20,21,30,31^. Indeed, cell cycle arrest can precede apoptosis or senescence, both transiently implicated in tissue remodeling during embryogenesis^139,140^. Consistent with this idea, cell cycle arrest markers, including *Cdkn1a/p21, Cdkn1c/p57*, *Cdkn2b/p15, Cdkn2a/p16*, and *Rb1*, are involved in both processes^140–143^. Further investigations will be needed to elucidate the temporal transition between cell cycle arrest and ensuing cellular behaviors. Intriguingly, in cancer, which is often characterized by uncontrolled proliferation and failure of apoptosis, cell cycle arrest promotes behaviors including extracellular matrix degradation, migration, and invasion^144–149^. Furthermore, in *Drosophila* and *Xenopus*, cell cycle inhibition is required for neurulation, gastrulation, and organ formation^150–153^. Here, we uncover a novel mechanism whereby cell cycle arrest mediates facial prominence and primary palate fusion in mammals, potentially representing a general process underlying diverse developmental tissue fusion events in different species.

Significant downregulation of cell cycle inhibitor genes in the ZL of embryos from two different genetic OFC mouse models also strengthens the notion that disruption of cell cycle arrest is likely a general mechanism underlying failure of facial prominence fusion to give rise to this congenital disorder. The intersection of our murine scRNAseq datasets and human WGS of OFCs, identified a novel transcription factor-encoding gene enriched in the ZL, *ZFHX3*. Notably, *ZFHX3,* which is highly mutated in OFC patients, was previously implicated in *Cdkn1a/p21* transactivation^35–37^, reinforcing the importance of ZL cell cycle arrest in the etiology of OFC. The presence of OFC in mice with cephalic epithelial-specific *Zfhx3* loss validates its essential role in mammalian facial prominence fusion.

Despite their widespread presence and genome-wide binding, PBX1-ZFHX3 co-binding to a subset of ZL-enriched cell cycle arrest genes highlights their combined requirement in cell cycle regulation during prominence fusion. The appearance of OFC in mutant embryos with compound loss of *Pbx* family members on a *Zfhx3* heterozygous background further establishes genetic interaction between these genes at the organismal level. Lastly, transactivation of select cell cycle target gene promoters elicited by PBX1-ZFHX3 co-expression in luciferase assays points to their synergistic regulation of cell cycle, confirmed by the finding that the two proteins form a complex within murine λ tissues. The presence of PBX1-ZFHX3 complexes may partly explain their selective co-binding to cell cycle regulator genes, restricting their otherwise promiscuous binding behavior^125^. Additional, yet unknown, transcription factors may act as co-factors, further enhancing restricted DNA binding and conferring tissue specificity^154^.

Multiple atlases have described genomic landscapes, gene networks, and single-cell transcriptomes of human embryonic tissues and organs^155–159^. However, the links between distinct cell populations, their gene transcripts and behaviors, as well as their *in vivo* functions, in human morphogenesis have remained largely unexplored. Our discovery of a ZL-like epithelial population arrested in the cell cycle in human facial prominences, expressing the same cell cycle regulators as in mice, demonstrates that fusion of the prominences and primary palate share evolutionarily conserved morphogenetic processes and transcriptional mechanisms in mice and humans. Our model proposes that the ZL, a novel, temporally regulated, cell cycle-arrested cephalic epithelial subpopulation is essential for facial prominence fusion in mice and humans and is compromised in craniofacial disorders. Furthermore, ZL-enriched ZFHX3 forms a complex with PBX1 to regulate expression of key cell cycle arrest genes during prominence fusion (Fig. 7C). Together, our studies establish a pivotal role for the ZL epithelial cell cluster in shaping the mammalian face and connect cell cycle arrest to developmental tissue fusion.

## Acknowledgements

We thank R. Aho for all artwork; P. Nolan and R. Dumbell for the ZFHX3 antibody; A. Arjun-McKinney for assistance with initial BrdU experiments; P. Martin for mouse colony maintenance and *in situ* hybridizations not included in the current figures; I. Barozzi, D. Goekbuget, and R. Boileau for guidance in bioinformatic analyses; V. Hermosilla Aguayo for guidance in Co-IP and for generation of probes; J. Zheng for generation of probes; J. Bush for useful discussions; J. Cyster, J. Bush, E. Hutchins and C. Teng for use of microscopes; P. Lupo, S. Sherman, and J. Espinosa for the Down Syndrome WGS data. Work supported by NIH R01 grant R01DE024745 and by UCSF Chancellor recruitment package to LS; R01-DE030342 and R03-DE027103 grants to EJL-C; TRDRP A139592 pre-doctoral fellowship and UCSF ‘Discovery Fellowship’ to TQ; F31DE032561 pre-doctoral fellowship to BC; Austrian Science Fund DOC33-B27 to LF; F31-DE032588 pre-doctoral fellowship to KR; and American Association for Anatomy post-doctoral fellowship to ML. WGS generated through grants: X01-HG010835, X01-HL136465, X01-HL140516, and X01-HL145692. G8.8 antibody developed by A.G. Farr was obtained from the Developmental Studies Hybridoma Bank, created by the NICHD of the NIH and maintained at The University of Iowa. Human embryonic and fetal material and related services provided by the Joint MRC/Wellcome (MR/R006237/1, MR/X008304/1 and 226202/Z/22/Z) Human Developmental Biology Resource (www.hdbr.org). The PFCC (RRID:SCR_018206) assisted the generation of Flow Cytometry data, supported in part by DRC Center Grant NIH P30 DK063720. Sequencing performed at UCSF CAT facility.

## Author Contributions

L.S. designed and supervised the study; T.Q. conducted lineage tracing and BrdU incorporation experiments, characterization of *p63* mutant mice, ChIPseq assays and bioinformatic analysis, and generated *Zfhx3* conditional mutant mice; B.C. performed characterization of epithelial cell populations by RNAscope, WISH, and IF on wild-type and *Pbx1/2* mutant embryos, GO analyses, and liaised with collaborators; L.F. and I.A. performed bioinformatic analysis of sc and bulk RNAseq and contributed critical ideas on cell cycle arrest and cell trajectories; M.L. initiated the project, designed the strategy as well as conducted all scRNAseq and bulk RNAseq of cephalic epithelium on wild-type and *Pbx1/2* mutant embryos and supervised B.C.; R.H-M. conducted experiments on *Fucci2a* mouse embryos, RNAscope and IF assays on wild-type and *Pbx1/2* mutants, genetic interaction experiments between *Pbx1, Pbx2* and *Zfhx3*, Co-IP, and luciferase assays; K.R. and E.L. intersected scRNAseq data from the current study with WGS datasets from human OFC patients and identified *ZFHX3* as a gene enriched in the ZL carrying a high number of variants in OFC patients; A.J. and S.L. conducted RNAscope and IF experiments on human embryos; J.D-A. assisted with RNAscope, IF assays, and mouse colony maintenance; M.L., M.R. and G.P. provided substantial technical guidance and critical intellectual input; B.C. assembled initial version of figures 1-5. T.Q. revised the initial figures including additional panels and composed figures 6-7. L.F. contributed multiple figure panels supporting the bioinformatic analyses. B.C. drafted preliminary parts of Results section. T.Q., R.H-M., and L.S. wrote the manuscript, then critically reviewed by all co-authors.

## Supplementary Figure Legends

**Fig. S1:** Further characterization of cell clusters identified by scRNAseq from E9.5 – E11.5. **(A)** FACS gating strategy to isolate live EPCAM+ epithelial population from microdissected and dissociated midface λ tissue. Cell viability determined via selection of DAPI-cells (blue box, left). DAPI-EpCAM+ cells (green box, right) averaged 6%-10% of sorted λ tissue depending on tissue stage. **(B)** Normalized Gene Expression (NGE) of *Epcam* overlaid on UMAP for combined E9.5-E11.5 datasets. Clusters expressing *Epcam* (black dashed border) were further analyzed. **(C)** Relative proportion of clusters by gestational day. Clusters highly expressing *Epcam* are boxed in red. **(D)** Distinct embryonic timepoints displayed on combined UMAP from E9.5-E11.5 datasets. **(E)** Dot plot showing expression of the top 4 differentially expressed genes (DEGs) defining individual clusters. Dot sizes represent the proportion of cells within a given population expressing the target gene; color intensities indicate average expression levels. Five broad cluster categories were defined based on cell type-defining gene expression and/or spatial validation. **(F)** RNAscope validation of the ND, PD, and EE cell clusters. For ND and PD clusters, NGE heatmaps (left) are overlaid on combined UMAP from E9.5-E11.5 datasets. As the EE cluster is primarily found at E9.5 NGE heatmaps (left, bottom) are overlaid on the corresponding E9.5 UMAP. Cluster defining genes were validated by RNAscope at E11.5 for ND/PD, and at E9.5 for EE (magnified inset for *Aldh1a3* expression in ND, red dashed box). Scale bars: 100 (ND, PD) and 50 (EE) µm, respectively. Inset scale bar: 20 µm. e, eye; OV, optical vesicle; BA1, branchial arch 1; MxP, maxillary prominence. White arrows point to gene expression.

**Fig. S2:** Additional molecular characterization and bioinformatic predictions for cell trajectories of ZL epithelial cell cluster. **(A)** Heatmap of *Bmp4* NGE overlaid on UMAP for E11.5 wild-type λ epithelium showing *Bmp4* expression beyond the ZL boundaries (black dashed area, left). RNAscope confirms expanded *Bmp4* expression beyond the ZL by E11.5 (center). Scale bar: 50 µm. WISH reveals extensive *Bmp4* expression throughout the midface (right). Scale bar: 1 mm. **(B)** Heatmaps of additional key DEGs (enriched *Bambi, Igfbp5* in red; absent *Top2a* in blue) overlaid on UMAPs in the E10.5 ZL cluster. **(C)** RNAscope shows strong expression of ZL marker genes *Bambi* and *Igfbp5* at the prominence fusion site in both E10.5 coronal and transverse sections. The mitotic division marker *Top2a* is depleted at the ZL. Presence or absence/reduction of signal is highlighted by full or empty arrowheads, respectively, at the prominence tips or fusion site. Scale bars: 50 µm. **(D)** RNA velocity analysis of E11.5 scRNAseq datasets. **(E)** CytoTRACE analysis of combined E9.5-E11.5 scRNAseq datasets. **(F)** Further analysis of E11.5 ZL cluster (black highlighted area, left) reveals 5 subclusters (ZL0-4). RNA velocity analysis indicates two terminal endpoints (green stars) in ZL3 and ZL4 (center). Terminal differentiation states are visualized in a heatmap overlay on UMAP (right). **(G)** Select DEGs in E11.5 ZL subclusters shows high expression of *Tgfb2*, *Cdkn1a/p21*, and *Isl1* in terminal states (top), with notable absence of *Lef1*, *Sostdc1*, and *Pax3* in these states (bottom). **(H)** Changes in ZL subcluster proportions between E10.5 and E11.5.

**Fig. S3:** Reduction/absence of additional cell cycle progression gene transcripts in ZL epithelial cell cluster. **(A)** Immunofluorescence (IF) staining for phosphorylated Histone H3 (pHH3) and RNAscope for *Fgf9* at E10.5. White dashed lines define epithelium boundaries. Scale bars: 50 µm. **(B,C)** IF for BrdU shows a lack of BrdU incorporation at the E10.5 fusion site at both 1-hour **(B)** and 7-hour **(C)** BrdU chase. Scale bars: 50 µm. **(D)** Heatmaps of NGE of additional cell cycle genes overlaid on E10.5 UMAPs show reduced expression at the ZL (black dashed area). **(E, F)** RNAscope of *Cdk1* **(E, F)** and *Aurkb* **(E)** in domains of high *Bambi* expression at E10.5 **(E)** and E11.5 **(F)**. Absence/reduction of signal highlighted by empty arrowheads at the prominence tips or fusion site. Red dashed squares show magnified insets (right). White dashed lines define epithelium boundaries. *Bambi* expression labels the ZL cells. Scale bar: 50µm. Inset scale bars: 20µm.

**Fig. S4:** Further characterization of cell cycle arrest in ZL epithelial cell cluster. **(A, B)** WISH of cell cycle arrest gene transcripts. *Cdkn1a/p21* **(A)** and *Gadd45g* **(B)** expression from E9.5 – E11.5. Images shown in sagittal (E9.5 – E10.25) and frontal views (E10.5 – E11.5). Scale bars: 200 µm. Sections of E10.25 and E11.5 WISH embryos (right) highlight localized gene expression at the prominence fusion site. Scale bars: 50 µm. Arrows indicate domains of high expression within the λ. **(C)** UMAP of transcripts for cell cycle arrest genes (*Cdkn1a/p21*, *Gadd45g*, and *Rb1*) are enriched in the ZL cluster as shown by NGE overlaid on UMAPs (left) and RNAscope validations (middle, right) at E11.5. Scale bars: 50 µm. **(D)** NGE of cell cycle inhibitor genes overlaid on E10.5 λ epithelial cluster UMAPs reveal negligible expression of *Cdkn2a/p16* and *Cdkn2b/p15*, in addition to minimal levels of *Cdkn1c/p57*, across clusters. *Cdkn1b/p27* shows consistent expression throughout all clusters. Black dashed area highlights ZL cluster. **(E)** RNAscope reveals localized expression of *Cdkn2a/p16* and *Cdkn2b/p15* at the prominence fusion site of E10.5 mouse embryo, with clear overlap indicated within magnified insets (red dashed squares). *Cdkn1c/p57* is also expressed at the fusion site. *Cdkn1b/p27* is expressed throughout the midface. Arrowheads point to gene expression. Scale bars: 50µm. Inset scale bar: 20µm. **(F)** Co-localization of p21 protein, detected by IF, and *Cdkn1a* gene transcript (encoding p21), revealed by RNAscope, at the λ fusion site at E10.5. Red dashed square shows magnified inset. Arrowheads highlight co-localized signal. Dashed lines define epithelium boundary in all panels. Scale bars: 50 µm. Inset scale bars: 20 µm. **(G)** *Fucci2a* mouse system adapted to selectively label cell cycle phases in the mouse cephalic epithelium *in vivo*. IF for cell cycle defining markers (mCherry, G0/G1; mVenus, S/G2/M) demonstrates an enrichment of G0/G1 phase and an absence of S/G2/M phase cells within the ZL population at E10.5. IF for CDH1 marks epithelial cells and RNAscope for *Bambi* expression marks ZL cells. White dashed lines define epithelium boundaries. Presence or absence/reduction of signal highlighted by full or empty arrowheads, respectively. Scale bars: 50 µm. Inset scale bar: 20 µm.

**Fig. S5:** Downregulation of select cell cycle arrest markers within the ZL in a mouse model of OFC. **(A)** RNAscope shows significant reduction of *Cdkn2a/p16* and *Cdkn2b/p15* expression at the prominence tips in E11.5 *Pbx1/2* mutant (right) vs control (left) embryos. Presence or absence/reduction of signal highlighted by full or empty arrowheads, respectively, at the prominence tips. White dashed lines define epithelium boundaries. All scale bars = 50µm. **(B)** RNAscope of E10.5 mouse embryos showing *Cdkn1a/p21* expression in *Pbx1/2* constitutive LOF (*Pbx1^-/-^;Pbx2^+/-^*); *Pbx1/2* mutant; *p63* constitutive LOF; and control (*Pbx1^+/-^;Pbx2^+/-^*) prominence tips. **(C)** IF demonstrates the absence of PBX1 protein in the λ epithelium of E10.5 *Pbx1/2* mutant (right), with persisting expression in presumptive olfactory neurons embedded within the epithelium. RNAscope reveals expression of *Cdkn1a/p21* and *Bmp4* at the prominence tips in both control (left) and *Pbx1/2* mutant (right) embryos.

**Fig. S6:** *Zfhx3/ZFHX3* is enriched in the ZL epithelium of mouse and human embryos and *ZFHX3* carries numerous variants in OFC patients. **(A)** Transverse section of midface showing λ (blue box) of CS16 human embryo. Scale bar: 100 µm. RNAscope shows enriched expression of *GADD45G* (orange) at the prominence fusion site of CS16 human embryo, while IF reveals striking reduction of KI67 (blue). Scale bar: 50 µm. Magnified inset (red box) further reinforces that the *GADD45G* expression domain overlaps with reduced KI67. Inset scale bar: 20 µm. Presence or absence/reduction of signal highlighted by full or empty arrowheads, respectively, at the prominence tips or fusion site. White dashed lines define epithelium boundaries. **(B)** Representative HREM image of human CS17 embryo in lateral view^160,161^. RNAscope at CS17 reveals the absence of *CDK1* expression at the prominence fusion site. Scale bar: 50 µm. **(C)** Table listing the top 13 genes within the ZL cluster exhibiting the highest number of *de novo* variants along with observed variant types. **(D)** NGE overlaid on UMAPs highlights strong enrichment of murine *Zfhx3* in a subset of cells at E9.5 (left), prior to the emergence of ZL, and within the ZL cluster (black dashed area) at E11.5 (right). **(E)** RNAscope on sections from mouse (E11.5) and human (CS16) embryos shows that *Zfhx3/ZFHX3* is highly expressed in the λ epithelium. Scale bar: 50µm. **(F)** NGE of murine *Zfhx3* overlaid on UMAPs of ZL subclusters analyzed by RNA velocity highlights its enrichment in terminal endpoints (yellow stars) at E10.5 (left) and E11.5 (right). Violin plot of normalized *Zfhx3* expression across individual subclusters at these stages is also included for each time point. **(G)** Gross morphology of E12.5 mouse embryonic midface with epithelial-specific heterozygous deletion of *Zfhx3* (*Crect^Cre/+^;Zfhx3^f/+^*). Scale bar: 500 µm.

**Fig. S7:** Further characterization of ZFHX3-PBX1 collaborative DNA binding during prominence fusion. **(A)** IGV genome browser tracks showing the *Zfhx3*, *Cdkn2a/p16* and *Pbx1* loci. ChIPseq tracks: H3K27ac, red; p300 (GSM1199037), pink; PBX1 and ZFHX3, green and blue, respectively. ATACseq track (GSE199339), maroon. Gray bars highlight predicted cis-regulatory elements (CREs) bound by PBX1 and ZFHX3.

**Supplemental Table 1:** All rare, predicted pathogenic variants in constrained genes from the ZL.

## Materials and Methods

### Ethics statement and approval of animal research

All animal experiments adhered to national laws and received approval from local regulatory bodies and the Institutional Animal Care and Use Committees (IACUC) at UCSF. IACUC guidelines covered aspects of housing, husbandry, and welfare for experiments involving mice and embryos.

### Generation of mouse embryos

Mouse embryos were generated using previously described mutant alleles and genotyping methods for conditional and constitutive alleles of *Zfhx3*^1^*, Pbx1*^2–4^*, Pbx2*^5,6^*, p63*^7,8^, and transgenes such as *CrectCre*^9^*, Pitx2Cre*^10^*, Wnt4Cre*^11^*, Ai14(RCL-tdT)-D*^12^, and *Fucci2a*^13^. In *Fucci2a* mice, G0/G1 phase cells are marked by mCherry-hCdt1, which accumulates during G1 and is degraded at the transition to S. Cells in S/G2/M are labeled by mVenus-hGem, which builds up during the S/G2/M phases and rapidly degrades before cytokinesis. Wild-type (WT) Swiss Webster mice over 6 weeks of age, sourced from Charles River Laboratories, were bred via natural timed mating to generate WT embryos. Following euthanasia of the pregnant females, embryos were collected between gestational days E9.5 to E12.5, with E0.5 defined as noon on the day a vaginal plug was detected. Mouse embryonic stages are indicated in figures and/or figure legends. Genotyping of mice and embryos was performed before proceeding with all the analyses and experiments. Both male and female embryos were analyzed without sex discrimination, as the observed phenotypes in *Pbx1/2* and *p63* mutants are fully penetrant across all specimens.

### Human Embryos

Human embryonic tissues were obtained from the Human Developmental Biology Resource (HDBR) following voluntary pregnancy terminations, with written informed consent from each donor. This study received ethical approval from the Newcastle and North Tyneside 1 National Health Service (NHS) Health Authority Joint Ethics Committee (approval number: 08/H0906/21+5). Human embryo samples at CS16 and CS17 (n=2 for each stage) were fixed with 4% paraformaldehyde (PFA) and then embedded in paraffin. Antigen retrieval was performed according to the manufacturer’s guidelines for the RNAscope Multiplex Fluorescent Reagent Kit v2 (Advanced Cell Diagnostics)^14^. Subsequent incubation and detection steps followed the RNAscope probe manufacturer’s manual. Immediately after completing the RNAscope Multiplex assay, immunofluorescence was conducted using immunohistochemistry (IHC) methods recommended by ACD Bio-Techne.

### Gross Morphology

Mouse embryos were fixed in 4% PFA at 4°C overnight, dehydrated using a methanol/PBS gradient up to 100% methanol, and stored at –20°C. For bleaching, embryos were treated with Dent’s bleach (4:1:1 Methanol:DMSO:Hydrogen peroxide) for 2 hours in the dark at room temperature, followed by a 15-minute rinse in 100% methanol. Embryos were then rehydrated through a decreasing methanol/PBT series ending in PBT (0.1% Triton in 1X PBS). Embryos were stained with DAPI (1:20000) overnight at 4°C.

### Cell Isolation and Fluorescence Activated Cell Sorting

Mouse embryonic litters were dissected in ice-cold PBS, collected and staged based on somite count and limb morphology on designated days of development. Number of embryos used for these experiments are listed in the “Statistics and data reproducibility” section below. At E9.5, embryonic faces were isolated from the heads by dissecting the frontonasal and maxillary prominences, carefully avoiding the forebrain and developing eye placode. At E10.5 and E11.5, entire midfaces were dissected to include the MNP, LNP, and MxP, while avoiding eye and brain tissues. The lambdoidal junction was specifically isolated using surgical scalpels to dissect the upper halves of the LNP and MNP at the horizontal midline of the nasal pits, and the lower half of the MxP at the horizontal midpoint, ensuring the lambdoidal junction remained intact without separating the individual prominences. Swiss Webster WT embryos were pooled before dissociation into single cell suspensions for scRNAseq analysis. Dissected tissue samples were incubated in a fresh cocktail of 1X Liberase TL (Roche cat: 05401020001) and 1X DNAseI Grade II (Roche cat: 10104159001) at 37°C without agitation for 10-13 minutes, with gentle pipetting every 5 minutes to facilitate tissue breakdown. The enzymatic reaction was stopped with ice-cold PBS, and cells were then centrifuged at 300 RCF for 10 minutes and resuspended in ice-cold cell staining buffer (0.5% Bovine Serum Albumin, 2mM EDTA in PBS). Cells were stained with 1.25μL of anti-EpCAM-APC (Invitrogen cat: 17-5791-82) antibody for 15 minutes at 4°C in the dark, followed by centrifugation and resuspension in FACS buffer (5% FBS, 5mM EDTA in PBS). To identify dead cells, DAPI was added to the cell suspension at a 1:1000 ratio. Cells were kept on ice and sorted using a Sony SH800S sorter, with gating strategies employed to exclude doublets and dead cells. DAPI negative, EpCAM positive cells were collected.

### scRNAseq experiments

#### Generation of count matrices, QC and filtering

Live cells from pooled lambdoidal junctions were loaded into a well for single-cell capture using the Chromium Single-Cell 3′ Reagent Kit V3 (10X Genomics). Libraries were prepared with the same kit, with each sample uniquely barcoded using an i7 index. Pooled libraries were then sequenced on an Illumina NovaSeq sequencer. Initial processing of raw sequencing reads was conducted using the Cell Ranger v2.2.0 pipeline from 10X Genomics, which involved demultiplexing and aligning to the mouse genome (mm10 1.2.0). The main subsequent analyses were primarily performed using pagoda2 R package^15^. Additional filtering was performed based on gene/molecule dependency (pagoda2, gene.vs.molecule.cell.filter), with cells having more than 1×10^5^ transcripts, less than 1000 transcripts or more than 10% of mitochondrial proportion being filtered out. Genes with less than 10 total transcripts overall were discarded.

#### FAC sorted WT epithelial cells

Datasets from E9.5, E10.5, and E11.5 developmental time points were combined as a single pagoda2 object, using the timepoints as the covariate for batch correction. The gene expression variance from the filtered batch-corrected count matrix was adjusted (pagoda2, k=10) to extract over-dispersed genes. PCA was performed on the over-dispersed genes (pagoda2, nPcs=50, maxit=1000). A nearest neighbour graph was calculated from the PC space using 100 neighbors and cosine distances (pagoda2, k=100, centred), and UMAP visualisation^16^ (UMAP python, n_neighbors = 30, min_dist = 0.5) was generated from PCA space. Clusters were then identified on the KNN graph using Leiden algorithm^17^ (resolution=0.5). Finally, UMAP embedding was generated using the UMAP python package. Separate timepoints were processed similarly, at the differences of the following parameters: 50 nearest neighbors for E11.5 and 30 nearest neighbors for E9.5 and E10.5. Pathway overdispersion analysis was performed to generate biological aspects^18^. For each timepoint, cell cycle was scored using a list of known cell cycle markers, and CytoTRACE^19^ values were calculated using the raw filtered count matrices.

#### Pbx1/2 mutant epithelial cells

E10.5 datasets from mutant and WT were combined. A single pagoda2 object was generated, using the condition (WT/mutant) as covariate for batch correction. The filtered overview count matrix gene expression variance was adjusted (pagoda2, k=10) to extract over-dispersed genes, and PCA was performed on the over-dispersed genes (pagoda2, nPcs=50, maxit=1000). A nearest neighbour graph was calculated from the PC space using 100 neighbors and cosine distances (pagoda2, k=100, centred, cosine distance), and UMAP visualisation (UMAP python, n_neighbors = 30, min_dist = 0.5) were generated in PCA space. Clusters were then identified on the KNN graph using Leiden algorithm (pagoda2, resolution=0.5). Finally, UMAP embedding was generated using the UMAP python package. To specifically analyze epithelial cells, clusters expressing *Epcam* were only selected, and subsequently reanalysed using the same parameters as previously. For consistency, Leiden cluster labels from FACS-purified WT datasets (see previous section) were assigned to the WT/Mutant cells, using the conos package^20^.

#### RNA Velocity analysis

For all datasets, spliced and unspliced matrices were generated with the kallisto^21^ BUS tool using fastq files as input. The generated matrices were subset using the cell barcodes list extracted from the filtering process of the previous analyses mentioned. UMAP coordinates and Leiden clustering labels were transferred to the newly generated data. Main downstream analyses were performed using the scvelo package^22^. Genes with less than 20 total counts or less than 10 unspliced counts were discarded, and only the 2000 top variable genes were kept. For the WT datasets, velocity was computed for each timepoint separately, using stochastic model and default parameters from scvelo. For the mutant E10.5 dataset, RNA velocity was computed separately for each condition, with varying number of neighbors to account for the difference of total number of cells captured (20 and 15 neighbors for WT and mutant respectively). RNA Velocity was estimated using the dynamical model for higher precision. To estimate possible changes in fate transitions between WT and mutant at E10.5, Cross-Boundary Direction Correctness (CBDir) measurement was employed, a measurement initially developed to benchmark RNA velocity estimation tools^23^. Briefly, this measurement estimates how likely it is that a cell will develop into a specific target cell based on its current velocity. The velocity direction of the cell should match the direction of its development towards the target cell. As we know the cluster-wise transitions, we estimated CBDir of cells from a progenitor cluster that bounds the progeny cluster (in other words the nearest neighbors of that cluster). To assess any deviation of CBDir in mutant, Kolmogorov-Smirnov test was performed between the two estimations.

### Bulk RNAseq

As described above (see section titled: Cell Isolation and Fluorescence Activated Cell Sorting), epithelial cells were sorted using FACS (Sony SH800S), collecting 5,000-10,000 live EpCAM positive and negative cells for each embryo directly into separate tubes containing RLT plus lysis buffer (Qiagen, cat: 1053393).

After genotyping of individual embryos, RNA from four biological replicates per genotype was collected, with each replicate comprising 3-4 pooled embryos. Number of embryos used for these experiments are listed in the “Statistics and data reproducibility” section below. RNA extraction was performed using the RNeasy Plus Micro Kit (Qiagen, cat: 74034), and RNA quantity was assessed using the Qubit RNA HD Assay Kit (Invitrogen, cat: Q32852). RNA quality was evaluated using an RNA 6000 Pico kit (Agilent, cat: 5067-1513) on a 2100 Bioanalyzer (Agilent). Only RNA samples with RIN>9 were used for library preparation. PolyA mRNAs were captured using the NEBNext Poly(A) mRNA Magnetic Isolation Module (NEB, cat: E7490). RNA sequencing libraries were prepared from 100ng of RNA using the NEBNext Ultra™ II RNA Library Prep Kit for Illumina (NEB, cat: E7775). Library size and quality were verified using an Agilent 2100 Bioanalyzer with the High Sensitivity DNA kit (Agilent, cat: 5067-4626). Library DNA concentration was determined with the QuBit dsDNA HS Assay kit (Invitrogen, cat: Q32854). Libraries were sequenced on the Illumina HiSeq 4000, generating 50 base pair (bp) single-end reads. RNAseq fastq files from all samples were aligned using STAR^24^ with GENCODE vM27 genome annotation. Feature counting was done using featureCounts^25^. Processing of count matrix, which includes variance stabilization, PCA and differential expression analysis, was done using the Deseq2 R package^26^. Genes with a fold change ≥1.2 or ≤-1.2 and an FDR ≤0.05 were identified as differentially expressed genes (DEGs).

### ChIPseq

Swiss Webster WT murine embryonic midfaces were dissected from mouse embryos at E11.5 and immediately crosslinked for 10 minutes in 1% formaldehyde (Electron Microscopy Sciences, cat: 15710). Number of embryos used for these experiments are listed in the “Statistics and data reproducibility” section below. For ChIPseq experiments on isolated epithelium cells, dissected midfaces were dissociated into single-cell suspensions following the same protocol as for FACS, and immediately crosslinked for 10 minutes in 1% paraformaldehyde. For H3K27ac datasets, after crosslinking, epithelium and mesenchyme cells were sorted by FACS (Sony SH800S) as described above. For PBX1 datasets, the epithelial population was sorted using the MACS (Miltenyi Biotec) system with CD326 (EpCAM; cat: 130-061-101) microbeads. ChIP assays were conducted as previously reported^27^. Briefly, nuclei were isolated from the crosslinked cells. After disruption of the nuclear membrane, chromatin was sonicated using a Diagenode biorupter to achieve 150–400 bp DNA fragments. After a pre-clearing step, chromatin was incubated overnight with specific antibodies (3-5 µg) at 4°C, followed by a 30-minute incubation with Dynabeads protein A (for PBX1, ZFHX3, H3K27ac ChIPseq; Invitrogen cat: 10002D) to immunoprecipitate specific chromatin complexes. IP and input DNA were purified using the MicroChIP DiaPure kit (Diagenode, cat: C03040001). Antibodies used included PBX1 (Cell Signaling, cat: 4342), H3K27ac (Abcam, cat: 4729) and ZFHX3 (MBL, cat: PD011 and PD010). Following ChIP, DNA libraries were constructed using the MicroPlex Library Preparation Kit v2 (H3K27ac; Diagenode cat: C05010012) or v3 (PBX1, ZFHX3; Diagenode cat: C05010001) and sequenced on an Illumina HiSeq4000 (H3K27ac) or NovaSeqX (PBX1, ZFHX3) to generate 50 bp single-end or paired-end reads, respectively. ChIPseq analysis was conducted using the nf-core/chipseq bioinformatic pipeline^28,29^ (workflow container 2.0.0), aligned to the mm10 release of the mouse genome (Dec. 2011, GRCm38) using Bowtie2^30^ under default parameters. Genomic Regions Enrichment of Annotations Tool (GREAT) analysis identified gene ontology terms associated with genomic regions bound by ZFHX3. Bedtools^31^ intersect was used to identify regions co-bound by ZFHX3 and PBX1. Hypergeometric Optimization of Motif EnRichment (HOMER^32^; v5.1) was utilized for enrichment analysis of known transcription-factor-binding sites and *de novo* motif discovery.

### Whole-Mount In Situ Hybridization

Dissected embryos were fixed overnight at 4°C in 4% PFA, then dehydrated through a methanol/PBS gradient to 100% methanol and stored at –20°C for up to one year. To prevent probe trapping in the head regions of embryos older than E10.5, holes were punched from the back through to the hindbrain. Whole-mount *in situ* hybridization was performed as previously described^27,33,34^. Briefly, embryos were rehydrated and pretreated with Proteinase K (duration of Proteinase K dependent on the stage of the embryos), then hybridized overnight at 70°C with either sense or antisense riboprobes at a final concentration of 1 µg/ml in an incubation buffer containing 50% formamide, 5× SSC, 50 µg/ml yeast RNA, 1% SDS, 50 µg/ml heparin, and 0.1% CHAPS detergent (ThermoFisher Scientific, cat: 28299). Post-hybridization, embryos were washed at 70°C through a series of SSC solutions (5× SSC and 2× SSC, three times each for 30 minutes, and once each in 0.2× SSC and 0.1× SSC for 30 minutes, respectively). After a brief rinse in Tris-buffered saline with 0.1% Tween (TBST), embryos were incubated in 10% blocking reagent (ThermoFisher Scientific, cat: R37620). Positive signals were detected using an alkaline phosphatase (AP)-conjugated anti-digoxigenin antibody (Roche Diagnostics, cat: 11093274910) at a 1:5000 dilution. Following washes in TBST, embryos were incubated in NBT/BCIP in NTMT buffer (Roche Diagnostics, cat: 11681451001) according to the manufacturer’s instructions until color development was complete. Positive hybridization was visualized by a purple (NBT/BCIP) signal. To ensure reproducibility, at least three embryos for each genotype and developmental stage were analyzed.

### RNAscope and Immunofluorescence on mouse embryonic sections

Mouse embryos were collected and fixed in 4% PFA solution either overnight at 4°C or for 2 hours at room temperature with gentle agitation. Fixed embryos were then washed with PBS and allowed to sink in a 30% sucrose solution before being embedded in 100% Neg-50 frozen cryoblocks and stored at –80°C. Frozen blocks were sectioned at 14 μm thickness, air-dried at RT for 1 hour, and subsequently stored at – 80°C. Slides containing transverse and/or coronal midface sections were thawed and washed with 1X PBS to remove excess freezing medium before use. These slides were assayed using the RNAscope Multiplex Fluorescent Reagent Kit V2 (Advanced Cell Diagnostics, cat: 323100), following modified manufacturer instructions. Specifically, antigen retrieval steps were omitted, and protease treatment was limited to a 10-minute exposure to Protease Plus to prevent tissue damage. Probe mixes were hybridized for 2 hours at 40°C in a HybEZ II Oven (Advanced Cell Diagnostics). The appropriate HRP channels were developed with Opal 520, TSA Cy3 Plus, and Cy5 Plus (PerkinElmer, cat: FP1487001KT, NEL744E001KT and NEL745E001KT) dyes. Following this, DAPI staining and mounting with ProLong Gold (Invitrogen, cat: P36930) were performed. For sections requiring immunofluorescence in conjunction with RNAscope, immunofluorescence assays were carried out after the final RNAscope step, which involved blocking the last HRP channel. An antigen blocking solution containing 5% Normal Donkey Serum and 0.1% Tween-20 in 1X PBS was applied to the slides for 45 minutes. Subsequently, primary antibodies against TUBB3 (1:200, Abcam cat: ab18207), RFP mCherry (1:300, Cell Signaling cat: 43590), GFP-Venus (1:200, Nacalai cat: GF090R), E-cadherin (1:100, R&D systems cat: AF748), pHH3 (1:500, Millipore Sigma cat: 06-570), KI67 (1:200, Abcam cat: ab15580), cleaved-Caspase3 (1:200, Cell Signaling cat:9661), EpCAM (1:200, DSHB cat: G8.8), and P21 (1:100, Millipore Sigma cat: MABE1816-100UG) were diluted in an antibody buffer (a 1:5 dilution of blocking solution to PBS-0.1% Tween-20 solution) and applied to the slides overnight at 4°C.The following day, primary antibodies were washed away, and the slides were washed three times for 15 minutes each with antibody buffer. Alexa Fluor secondary antibodies (1:500, Invitrogen), diluted in antibody buffer along with DAPI (1:1000), were applied to the slides for 2 hours at room temperature. Following additional washes to remove the secondary antibody, slides were flicked dry and mounted using ProLong Gold with coverslips.

### Thymidine Analogue Incorporation and TUNEL Assays

Bromodeoxyuridine (BrdU; Invitrogen cat: B23151) was diluted in saline solution (0.9%) to a final concentration of 1mg/ml. Pregnant Swiss Webster dams were anesthetized using isoflurane gas inhalation and injected intraperitoneally with BrdU at a dose of 100μl/10g body weight. Following the injection, the dams were observed and allowed to recover. Pregnant dams were then sacrificed at either 1-or 7-hour post-injection, and embryos were collected, fixed, and cryo-embedded as previously described. For sections from BrdU-injected embryos, slides were treated with a 2M HCl solution for 30 minutes at 37°C, followed by washing and incubating with blocking buffer (10% Normal Goat Serum in PBS with 0.3% Triton X) for 1 hour at room temperature. Rat anti-BrdU antibody (1:500, Abcam ab6326) was diluted in antibody buffer and incubated overnight. The following day, slides were washed, and anti-Rat secondary antibodies (1:500; Invitrogen cat: 21208), along with a DAPI counterstain (1:1000), were applied for incubation before final washes and mounting with Prolong Gold antifade. The TUNEL assay was performed on cryosections following the manufacturer’s guidelines for the In Situ Cell Death Detection Kit (Millipore Sigma, cat: 12156792910).

### Microscopy of whole embryos or embryonic sections

Whole Mount *in situ* hybridization embryos were imaged using a Leica M20F5A dissecting microscope with a 1x/0.03 lens and variable zoom settings. Images were processed and exported using LAS-X software package. Embryos for gross morphology were imaged using the Axio Imager.Z2 scope at 0.25X magnification using the DAPI filter. Sections were imaged using a Zeiss Axio Observer.Z1, Zeiss Axiovert 200M, or confocal microscope Zeiss LSM 900 microscope with 5x, 10x, or 20x/0.8 objective. For sections imaged on the confocal microscope, several z-slices were acquired. Images were post-processed using ZEN 3.6 or ZEN 2.3 Pro. Adjustments were made to reduce background autofluorescence of the tissues by adjusting the absolute black and absolute white levels using a Best Fit adjustment where necessary. Sections were imaged using the same settings for each experiment.

### Whole-genome sequencing (WGS) of OFC trios and Down syndrome patients

#### OFC trio samples

Case-parent trios with whole genome sequencing data originating from three ancestral groups were used for this study. The first set consists of 376 trios of European ancestry that were recruited from sites in the United States, Argentina, Turkey, Hungary, and Spain; the second is a set of 267 trios from Medellin, Colombia; and the third is a set of 116 trios from Taiwan. In 93.8% of European trios, 96.5% of Taiwanese trios, and 100% of the Colombian trios, parents were unaffected. Because OFC etiology is likely to be multifactorial, parent phenotype status was not considered in this analysis. By proband OFC type, there were 88 cleft lip only (CL; 8 Colombian, 80 European), 613 cleft lip and palate (CLP; 259 Colombian, 238 European, 116 Taiwanese), and 58 cleft palate (CP; all European). Subject recruitment and phenotypic assessment occurred at regional treatment centers. Each site’s institutional review board (IRB) and the IRBs of the affiliated US coordinating institutions (HRPO #03-0871, IRB#HSC-MS-03-090, IRB#970405, IRB#200109094, and IRB#200109094) provided ethical approval and oversight.

#### Whole genome sequencing and variant calling

The case-parent OFC trios used in this study were sequenced as part of the Gabriella Miller Kids First (GMKF) Research Consortium. Sequencing was performed on blood samples when available (the majority of samples) and saliva when blood was not obtainable. Whole genome sequencing for European samples was carried out by the McDonnell Genome Institute (MGI) at the Washington University School of Medicine (St. Louis, MO) followed by alignment to hg38 and variant calling at the GMKF Data Resource Center at the Children’s Hospital of Philadelphia. Sequencing for Colombian and Taiwanese samples was carried out by the Broad Institute, with alignment to hg38 and variant calling by GATK pipelines^35–37^. Additional details on alignment and workflow used to harmonize these datasets have been published in Mukhopadhyay et al^38^.

#### Quality control

The WGS data for all case-parent trios was subjected to several quality metrics. Individual samples were evaluated for missingness, Mendelian error rate, and average read depth, and were removed if these values were greater than three standard deviations from the mean. Additionally, samples with transition/transversion (Ts/Tv), exonic Ts/Tv, silent/replacement, or heterozygotes/homozygotes ratios outside of the expected values were removed. Due to trios within the Colombian cohort having higher rates of consanguinity than other groups, lower heterozygotes/homozygotes ratios were allowed. Identity by descent as tested in PLINK (version 1.90b53) was used to confirm familial relationships, and sex of the samples was confirmed by X chromosome heterozygosity.

#### Down syndrome samples

Paired-end WGS was completed by the Broad Institute as part of funded NIH INCLUDE and Gabriella Miller Kids First projects on an Illumina HiSeqX with a target read depth of ∼30x coverage. The data were aligned to hg38 and harmonized by the Kids First Data Research Center via Cavatica using a custom pipeline based on GATK best practices^37^. Quality control (QC) was performed on 2,394 samples and 98,049,632 variants using VCFtools v0.1.13. Variant calls with genotype quality (GQ) < 20 or depth (DP) < 10 were set to missing. Variants were then filtered to remove flagged sites (filter flag not equal to “PASS”), multiallelic sites, monomorphic sites (minor allele count (MAC) < 1) and sites with > 3% missingness, leaving 74,841,842 variants. Samples were dropped for quality metrics outside of three standard deviations from the mean for missingness, theta, transition/transversion (Ts/Tv) ratio, silent/replacement ratio, and heterozygous/homozygous ratio. Sex and family relationships were confirmed by X chromosome heterozygosity and identity-by-descent analyses in PLINK v1.9. The final analysis dataset consisted of 2,344 samples. Variants were then excluded from analysis if they had MAF < 5%, deviated from Hardy-Weinberg (p-value < 1×10−7), or had site missingness > 5%. Full details of sequencing and variant calling are found in Feldman, et. al^39^.

#### Identification of de novo variants

Mendelian errors were called in trios using PLINK (version 1.90b53) and bcftools (v1.9). These mendelian errors then underwent further filtering to yield high quality de novo mutation calls including filtering for passing variants, bi-allelic variants only, a minor allele count (MAC) = 1, genotype quality (GQ) ≥ 20, and depth (DP) ≥ 10 using VCFtools (version 0.1.13). Furthermore, we filtered variants on the basis of allele balance (AB), with de novo calls requiring an AB ratio ≥0.30 and ≤0.70 in the proband and an AB ratio < 0.05 in the parents. Annotation of high confidence DNMs was completed with ANNOVAR (version 201707). Following annotation, de novo variants in coding regions were selected based on functional classification (“exonic” and/or “splicing”) and frequency (MAF < 0.3% across all of gnomAD v3.1.2)

#### De novo variant enrichment

Enrichment of de novo variants (DNs) in 759 OFC trios, 6,430 ASD probands, and 2,179 unaffected siblings was statistically analyzed using the ‘DenovolyzeR’ package (version 0.2.0) in R. Enrichment is tested by determining if the number of observed variants in a dataset is greater than what is expected based on mutational models described by Samocha, et al^40^. The functions ‘DenovolyzeByClass’ and ‘DenovolyzeByGene’ were used to test for an excess of DNMs both globally and per gene, respectively; however, use of the function ‘includeGenes’ was implemented to restrict testing to the gene sets derived from single cell RNA sequencing only. When testing enrichment per cluster, results were considered significant at p<0.0083 (Bonferroni correction 0.05/6 clusters).

#### Rare variants in top scRNAseq genes

The top 150 genes by absolute Z-score of each scRNAseq cluster were converted from Mus musculus to Homo sapiens orthologues using g:Profiler (Raudvere et al., 2019), and coding variants in orthologous genes were then extracted for 876 families with OFCs. Synonymous variants were first removed so that only protein-altering variants remained, and variants were then filtered for minor allele frequency (MAF) of <0.5% in any population in both gnomAD v2.1.1 and v3.1.2. Qualitatively, we evaluated variants based on multiple pathogenicity predictors. We excluded variants with CADD scores <20 and SIFT scores >0.05 and prioritized constrained genes (LOEUF <0.35 and pLI >0.9). For *ZFHX3*, we visualized predicted damaging variants in OFC probands with ProteinPaint^41^. The percent of probands harboring *ZFHX3* variants in OFC, DS, and healthy cohorts were compared using a Chi-Squared test.

### Gene Ontology Analysis

The top 100 differentially expressed genes ordered by Z-score absolute values from E10.5 λ epithelium scRNAseq datasets were used as input into the NIH Database for Annotation, Visualization, and Integrated Discovery (DAVID) functional annotation tool. 100 gene DAVID ID matches to *M. musculus* were charted using the GOTERM_BP_DIRECT category and ranked by P-value.

### Luciferase assays

The vector constructs for the gene fragment sequences listed below were synthesized by TWIST Bioscience and cloned into the KpnI/HindIII sites of the pNL3.2 (Nluc/minP) vector (Promega, cat: N1041). After cloning, colonies were screened, selected, and sequenced to verify correct insertion. HEK293T cells were cultured in DMEM containing 4.5 g/L glucose, L-glutamine, and sodium pyruvate (Corning, cat: 10-013-CV) supplemented with 10% fetal bovine serum (Cytiva Hyclone, cat: SH30007203) and 1% penicillin-streptomycin (Hyclone, cat: SV30079). The cells were maintained in a humidified chamber with 5% CO2 at 37°C. For co-activation assays, approximately 15,000 HEK293T cells were transfected for 6 hours with 100 µl of OPTIMEM (Gibco, cat: 31985070), 2 µl of Lipofectamine 2000 (Invitrogen, cat: 11668027), and 5 ng of pGL4.54 (luc2/TK) (Promega, cat: E506A) plus the following vector cocktails: 1) Control: 200 ng of empty vector (pCDNA3.0) + 50 ng of empty pNL3.2 (Nluc/minP), 2) ZFHX3: 100 ng of ZFHX3 overexpression vector (OV) + 100 ng of empty vector + 50 ng of the targeted sequence, 3) PBX1: 100 ng of PBX1 OV + 100 ng of empty vector + 50 ng of the targeted sequence, and 4) PBX1+ZFHX3: 100 ng of PBX1 OV + 100 ng of ZFHX3 OV + 50 ng of the targeted sequence. The cocktail mix was added directly to the growth media. After 6 hours, transfection media/cocktail mix was replaced with fresh media, and the cells were incubated for an additional 48 hours. Transfections were performed in 24-well plates, with each condition tested in triplicate. After 54 hours, cells were harvested and analyzed using the Nano-Glo Dual-Luciferase Reporter (Promega, cat: N1610). Briefly, cells were washed with 1X PBS and lysed with 100 µl of 1X PLB lysis buffer (Promega, cat: PRE1941) for 20 minutes at room temperature. The cell lysate was collected and centrifuged for 10 minutes at room temperature. Then, 20 µl of the supernatant was transferred to a 96-well white flat-bottom plate (Costar). The Nano-Glo Dual-Luciferase Reporter Assay was performed using an automated sample injector system (Promega). First, 80 µl of ONE-Glo EX Reagent was injected, and the pGL4.54 (luc2/TK) signal was measured.

Subsequently, 80 µl of Nano-DLR Stop&Glo Reagent was injected, and after 5 minutes, the pNL3.2 (Nluc/minP) readout was recorded. The ratio of pGL4.54 (luc2/TK) to pNL3.2 (Nluc/minP) signal for each sample was calculated. Triplicates for each condition were averaged, and results were compared to the normalized control readout. Data are represented as a fold increase and T-tests were performed for paired comparisons.

#### Genome coordinates of the fragments cloned

Cdkn1c promoter: 442pb-chr7:143,460,804-143,461,362

Pbx3 promoter-729pb –chr2:34,372,272-34,373,000

Rb1 promoter: 295pb-chr14:73,325,702-73,325,988.

### Immunoprecipitation

Approximately 60 midfaces of Swiss-Webster embryos at either E10.5 or E11.5 were lysed in 500 µl of lysis buffer (50 mM Tris-HCl, pH 7.4, 150 mM NaCl, 2.5 mM MgCl2, 1 mM EDTA, 10% glycerol, 0.2% sodium deoxycholate, 1% Triton X-100) containing phosphatase inhibitors (cocktails #2 and #3, Sigma cat: P5726 and P0044) and protease inhibitors (Sigma, cat: P8340). The supernatant was collected, and protein concentration was measured using the Pierce BCA Protein Assay Kit (Thermo, cat: 23235). For each sample, 800 µg of total protein in 500 µl was used. To pre-clear the samples, 100 µl of Dynabeads Protein-G (Invitrogen, cat: 10003D) were first washed twice with 1X PBS and then twice with wash solution (50 mM Tris-HCl, pH 7.4, 200 mM NaCl). Subsequently, 100 µl of the washed Dynabeads were added to 800 µg of protein supernatant per sample and incubated for 2 hours at 4°C with rotation. The supernatant was then collected, and each sample was incubated overnight at 4°C with rotation with either 5 µg/µl of ZFHX3 antibody (R&D Systems AF7384) or 5 µg/µl of normal sheep IgG (R&D Systems 5-001-A). The following day, the supernatant was discarded, and the Dynabeads were extensively washed with washing solution (50 mM Tris-HCl, pH 7.4, 200 mM NaCl, 10% glycerol, 1% Triton X-100) on ice. Protein complexes were eluted from the Dynabeads by incubating for 15 minutes at 95°C in 80 µl of 4x NuPage Buffer (Invitrogen, cat: NP007) containing 355 mM β-mercaptoethanol (Sigma, cat: 444203). Thirty microliters of each sample were loaded onto a 4-12% polyacrylamide gel (Invitrogen, cat: NP0322) and resolved using a standard Western blot protocol. Following transfer, the protein membrane was incubated overnight at 4°C with a PBX1 antibody (Cell Signaling cat: 4342, 1.4 µg/µl, customized) diluted 1:5000 in 5% blocking reagent/TBST. For signal development, the membrane was incubated with TidyBlot (BioRad, cat: STAR209) diluted 1:200 in blocking reagent/TBST for 1 hour at room temperature, followed by an HRP assay. The signal was developed using SuperSignal substrate (Thermo, cat: 34075).

## Statistics and data reproducibility

Statistical methods were not employed to predetermine sample sizes but followed established field standards. Data were included in the analyses unless there were rare instances of clear technical failures. All omics-datasets adhered to ENCODE guidelines, which require at least two biological replicates for each experiment. scRNAseq required large numbers of WT embryos to isolate λ epithelium through FACS purification. 2 biological replicates were conducted at E9.5 (n=7-14 embryos each; 25k cells) and E10.5 (n=7-14 embryos each; 30k cells), and 3 biological replicates for E11.5 (n=7-14 embryos each; 30k cells). In contrast, scRNAseq of *Pbx1/2* mutants was conducted on entire midfaces due to limitations in obtaining mutant embryos because of complex genetics (n=2 embryos used for mutants and WTs respectively; 30k cells). For bulk RNAseq, purified λ epithelium was used in 4 biological replicates (n=2-3 embryos each). For all ChIPseq, 10-12 embryonic midfaces per replicate (minimum 2 independent biological replicates per condition) were pooled for experiments using whole midfaces (ZFHX3 and H3K27ac assays) and 50-60 embryos for experiments using purified λ epithelium (PBX1 assays). The high reproducibility and quality of these replicates led to statistically significant peak detections. For whole-mount *in situ* hybridization (WISH), RNAScope FISH, gross morphology, immunoprecipitation, and immunofluorescence, at least three independent biological replicates were examined for each genotype or developmental stage. Embryos for these analyses were sourced from various females and underwent a minimum of two completely independent experiments.

### List of Primers Used

Primers to generate WISH riboprobes:

*Cdkn1a/p21* F: CTCTTCCCCATCTTCGGCC

*Cdkn1a/p21* R: GAGACGCTTACAATCTGAGTGG

*Gadd45g* F: CCGATGAAGAAGATGAGGGCG

*Gadd45g* R: TGAAAGAGCAGTGCAGTCGG

*Bmp4* WISH probe was provided by Ian C. Welsh (see Welsh and O’Brian 2009)

### List of Probes Used

*<u>Catalog numbers of RNAscope probes (Advanced Cell Diagnostics)</u>: Gadd45g* (803431-C2), *Cdkn2a* (411011-C3), *Cdkn2b* (458341-C2), *Bambi* (523071), *Pbx1* (435171), *Foxn3* (586011), *Bmp4* (401301-C2), *Cdkn1c* (458331-C2), *Pitx2* (412841-C2), *Aurkb* (461761), *Zfhx3* (803471), *Rb1* (486191-C3), *Bambi* (523071-C3), *Igfbp5* (425731-C2), *Cdk1* (476081-C2), *Cdkn1b* (499991), *Itm2b* (491791), *Tfap2b* (536371-C3), *Sox2* (401041-C3), *Hey1* (319021-C3), *Dlk1* (405971-C2), *Fgf9* (499811), *Wnt6* (401111).

---

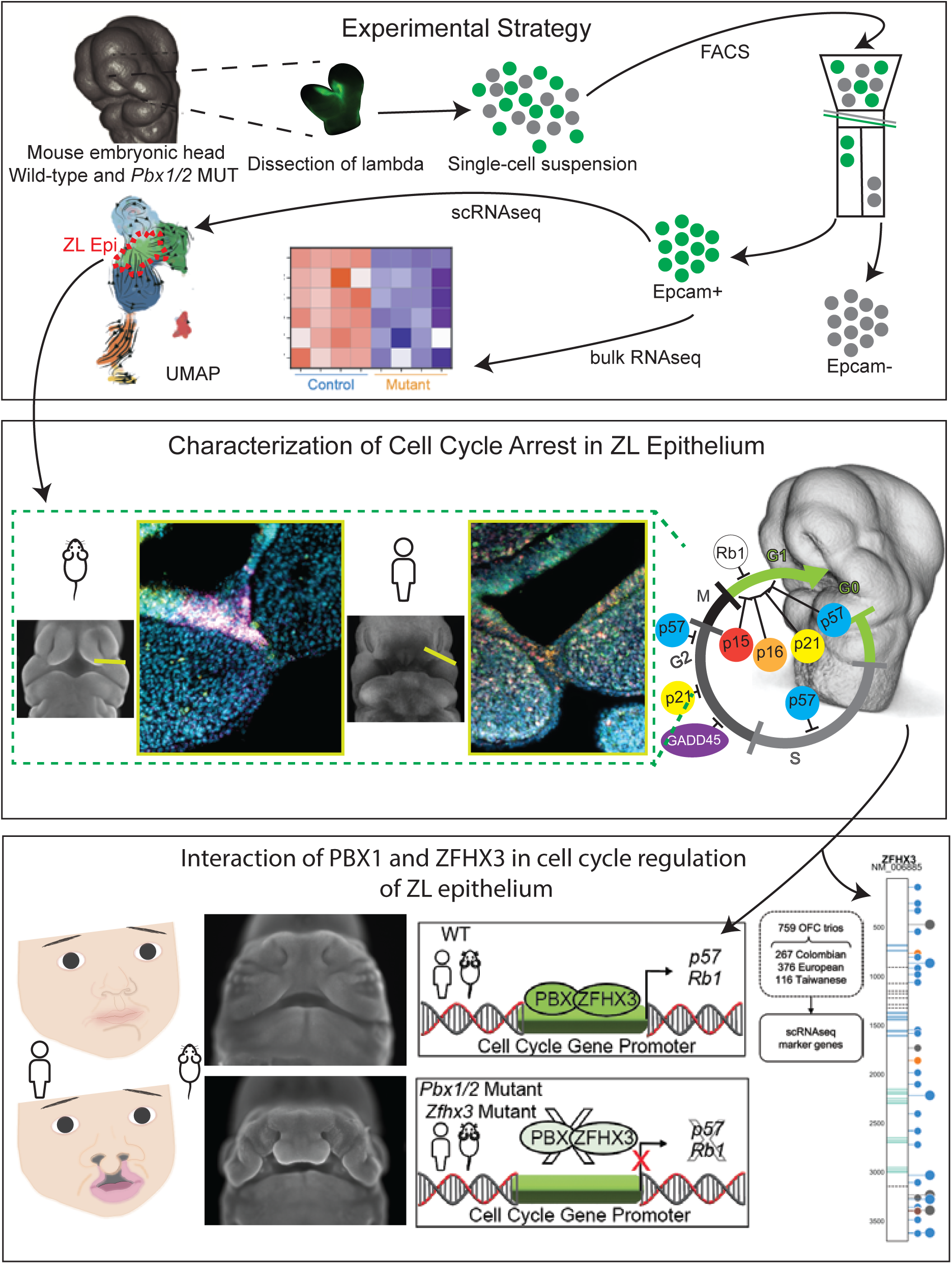

