## Supplementary figures and images for "Cell Cycle Arrest of a ‘Zippering’ Epithelial Cell Cluster Shapes the Face and is Disrupted in Craniofacial Disorders"

### Supplemental Figure 1

Fig.S1: Further characterization of cell clusters identified by scRNAseq from E9.5 - E11.5.

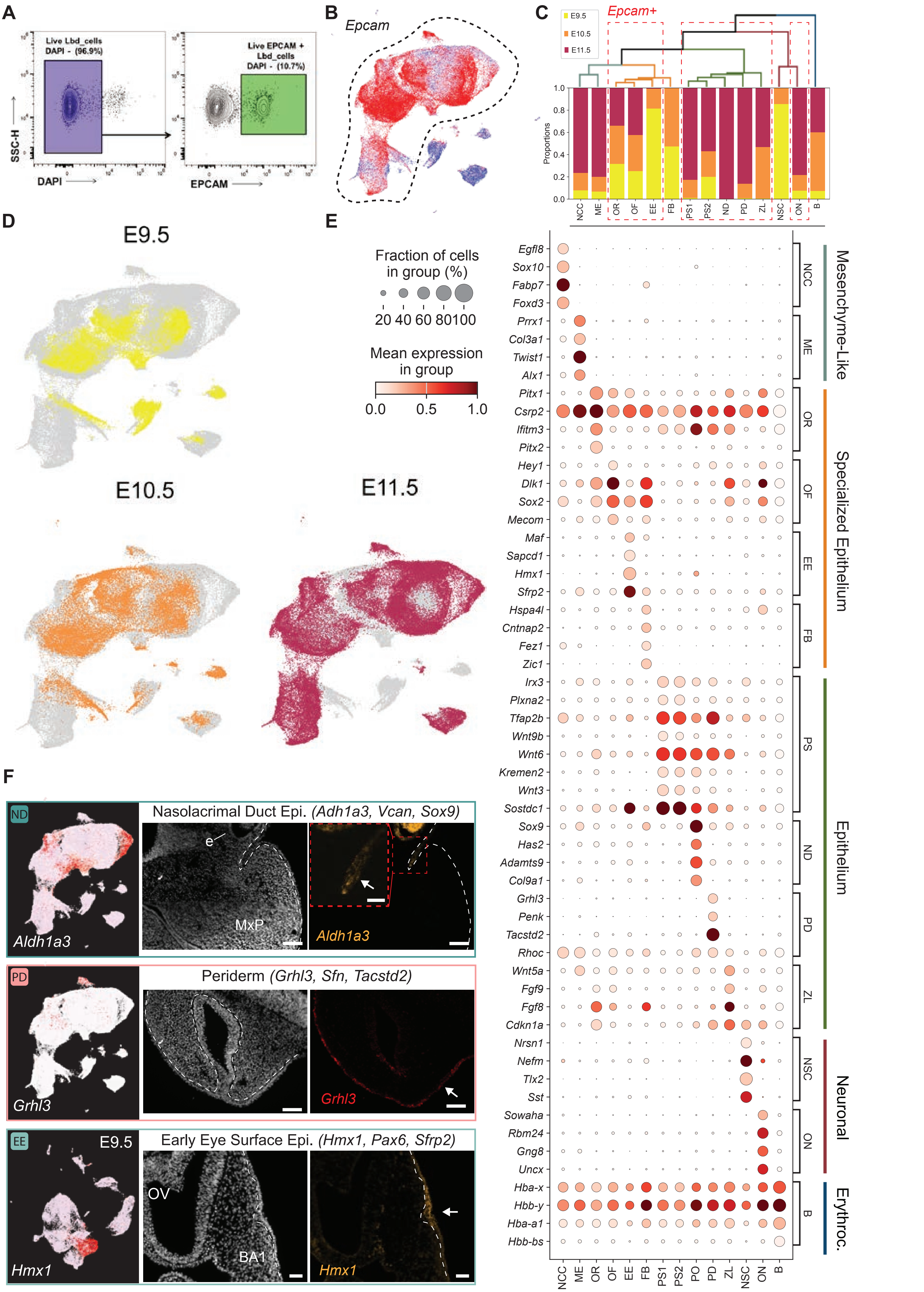

### Supplemental Figure 4

Fig.S4: Further characterization of cell cycle arrest in ZL epithelial cell cluster.

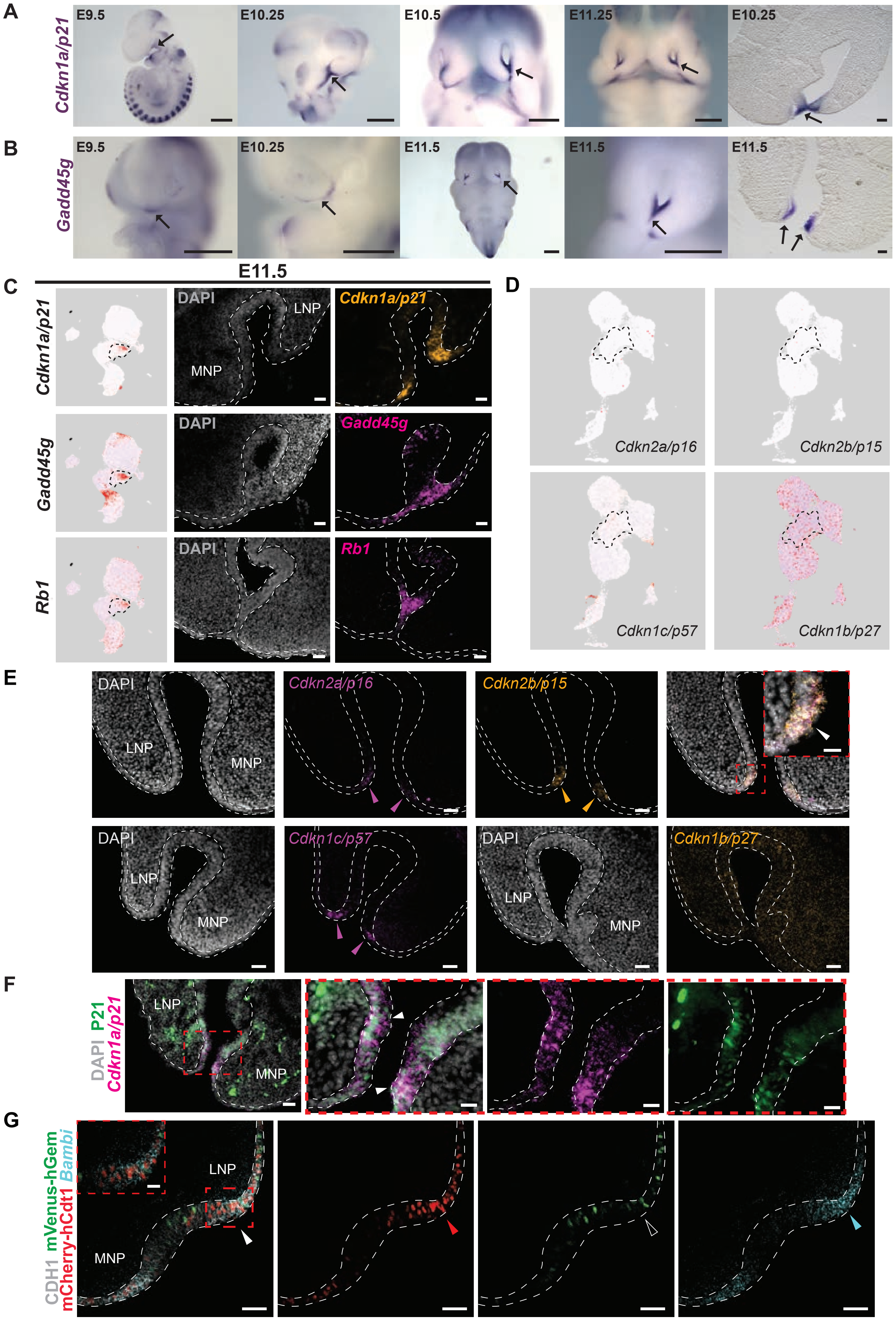

### Supplemental Figure 5

Fig.S5: Downregulation of select cell cycle arrest markers within the ZL in a mouse model of OFC.

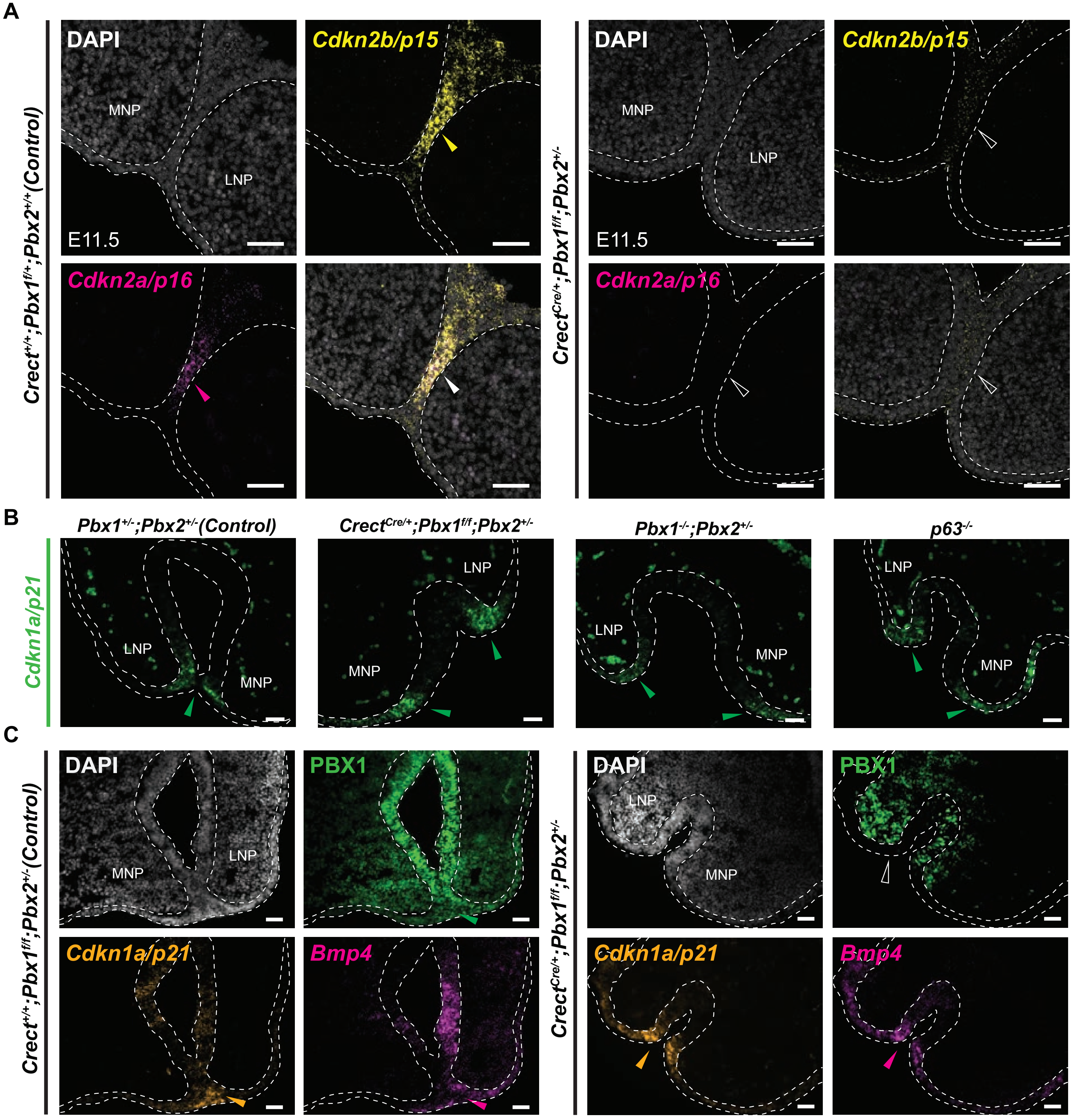
