## Supplemental Figure 2 for "Cell Cycle Arrest of a ‘Zippering’ Epithelial Cell Cluster Shapes the Face and is Disrupted in Craniofacial Disorders"

Fig.S2: Additional molecular characterization and bioinformatic predictions for cell trajectories of ZL epithelial cell cluster.

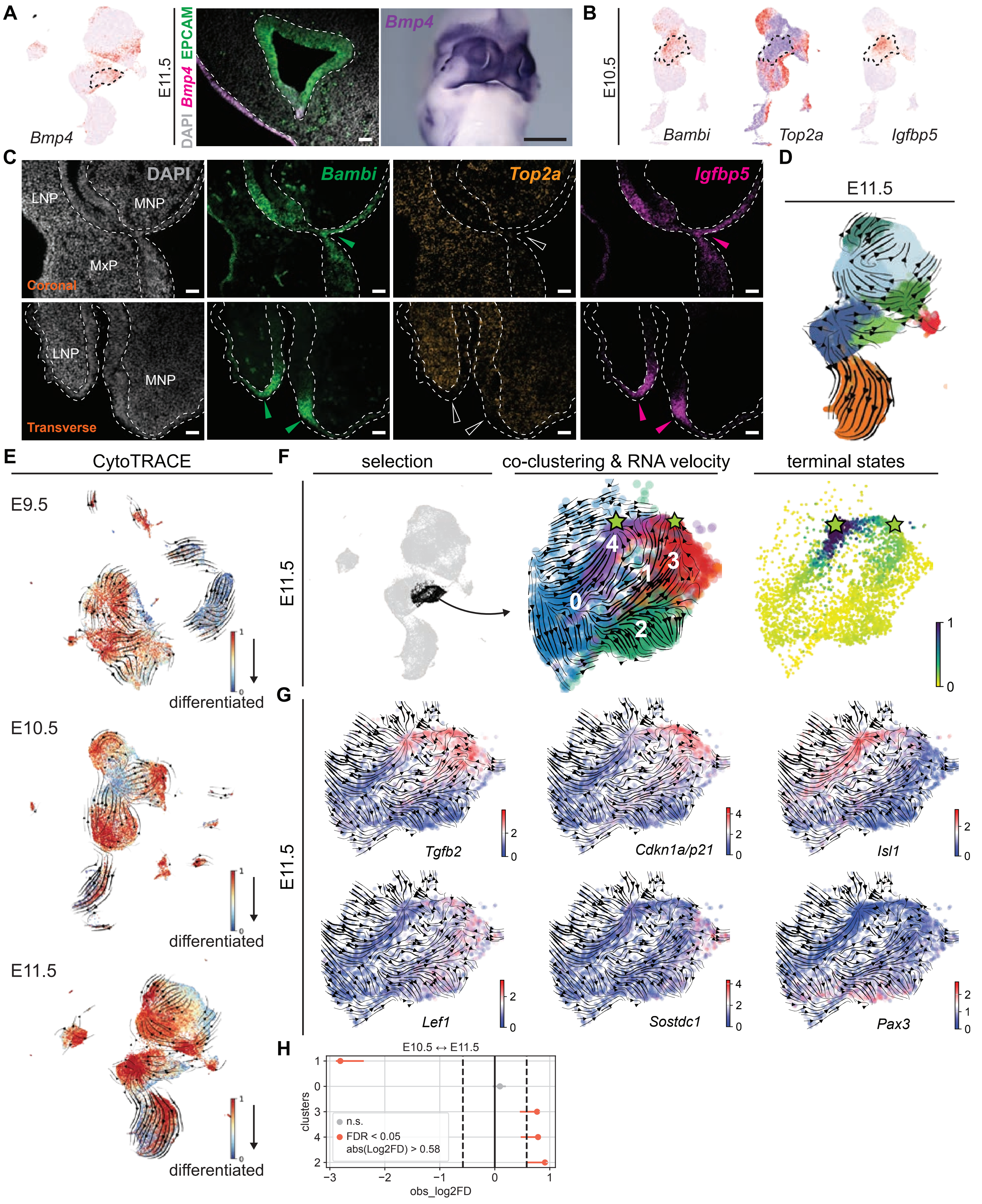
