## Supplemental Figure 3 for "Cell Cycle Arrest of a ‘Zippering’ Epithelial Cell Cluster Shapes the Face and is Disrupted in Craniofacial Disorders"

Fig.S3: Reduction/absence of additional cell cycle progression gene transcripts in ZL epithelial cell cluster.

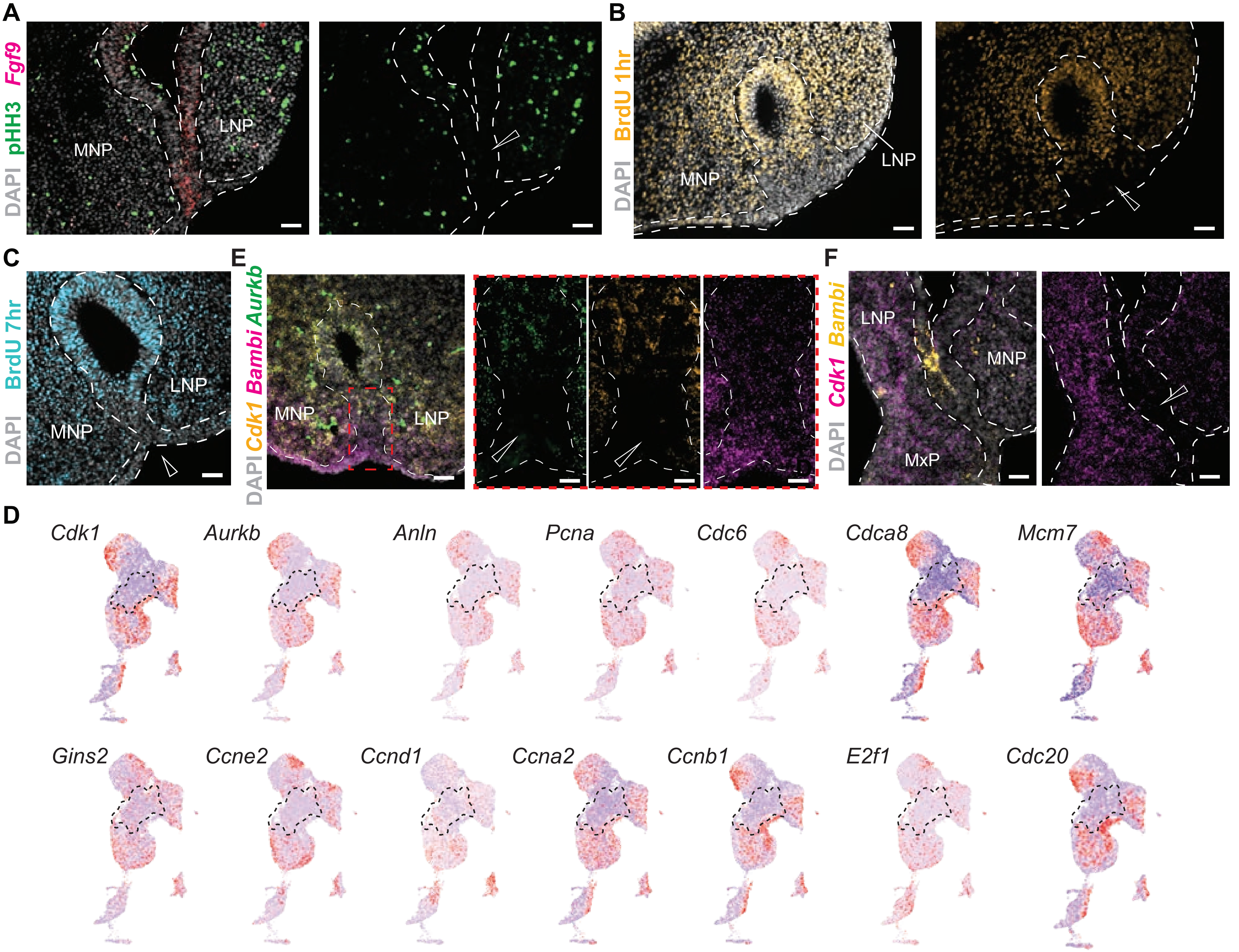
