## Supplemental Figure 6 for "Cell Cycle Arrest of a ‘Zippering’ Epithelial Cell Cluster Shapes the Face and is Disrupted in Craniofacial Disorders"

Fig.S6: *Zfhx3/ZFHX3* is enriched in the ZL epithelium of mouse and human embryos and *ZFHX3* carries numerous variants in OFC patients.

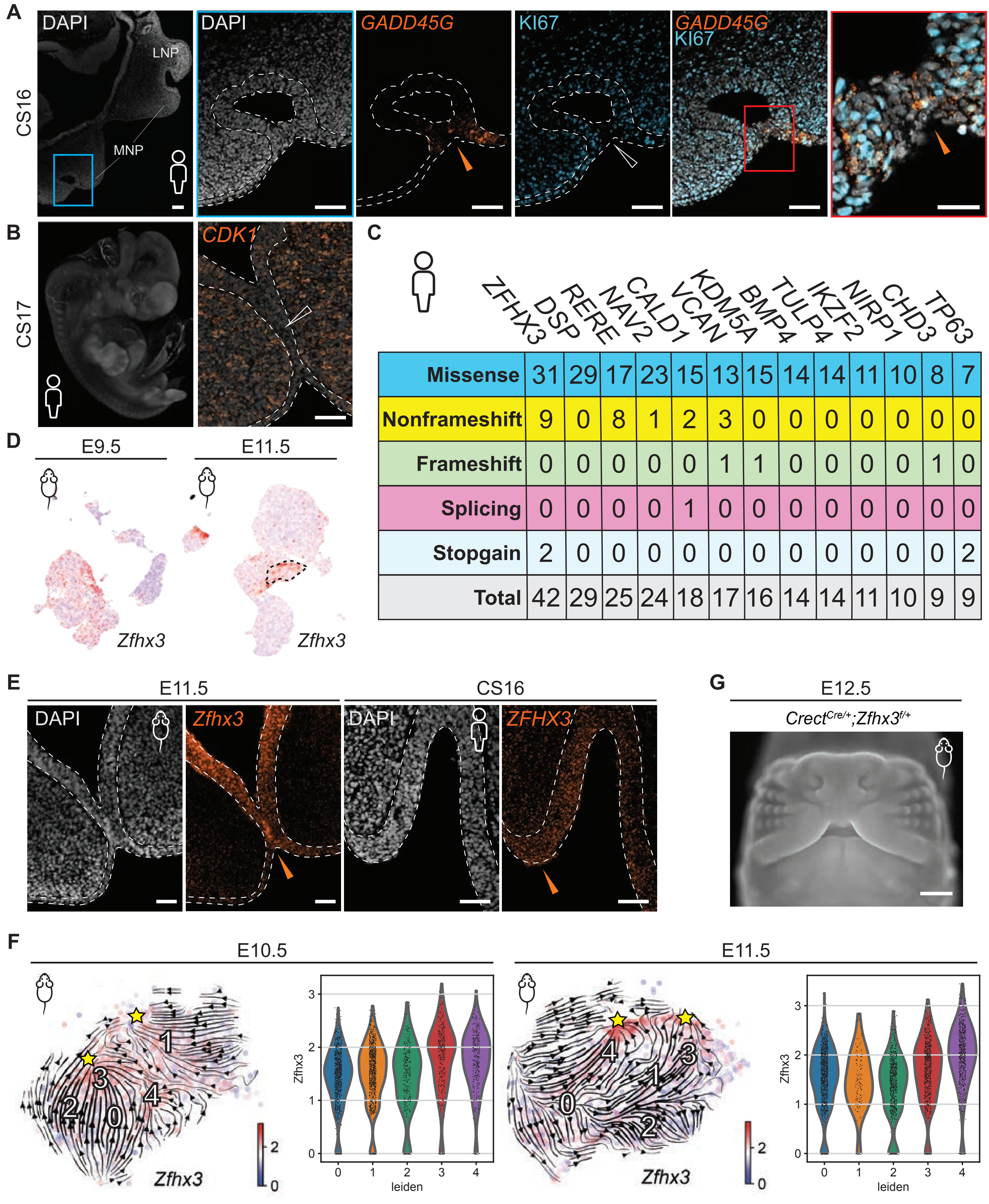


| | ZFHX3 | DSP | RERE | NAV2 | CALD1 | VCAN | KDM5A | BMP4 | TULP4 | IKZF2 | NIRP1 | CHD3 | TP63 |
|---|---|---|---|---|---|---|---|---|---|---|---|---|---|
| Missense | 31 | 29 | 17 | 23 | 15 | 13 | 15 | 14 | 14 | 11 | 10 | 8 | 7 |
| Nonframeshift | 9 | 0 | 8 | 1 | 2 | 3 | 0 | 0 | 0 | 0 | 0 | 0 | 0 |
| Frameshift | 0 | 0 | 0 | 0 | 0 | 1 | 1 | 0 | 0 | 0 | 0 | 1 | 0 |
| Splicing | 0 | 0 | 0 | 0 | 1 | 0 | 0 | 0 | 0 | 0 | 0 | 0 | 0 |
| Stopgain | 2 | 0 | 0 | 0 | 0 | 0 | 0 | 0 | 0 | 0 | 0 | 0 | 2 |
| Total | 42 | 29 | 25 | 24 | 18 | 17 | 16 | 14 | 14 | 11 | 10 | 9 | 9 |
