## Supplemental Figure 7 for "Cell Cycle Arrest of a ‘Zippering’ Epithelial Cell Cluster Shapes the Face and is Disrupted in Craniofacial Disorders"

Fig.S7: Further characterization of ZFHX3-PBX1 collaborative DNA binding during prominence fusion.

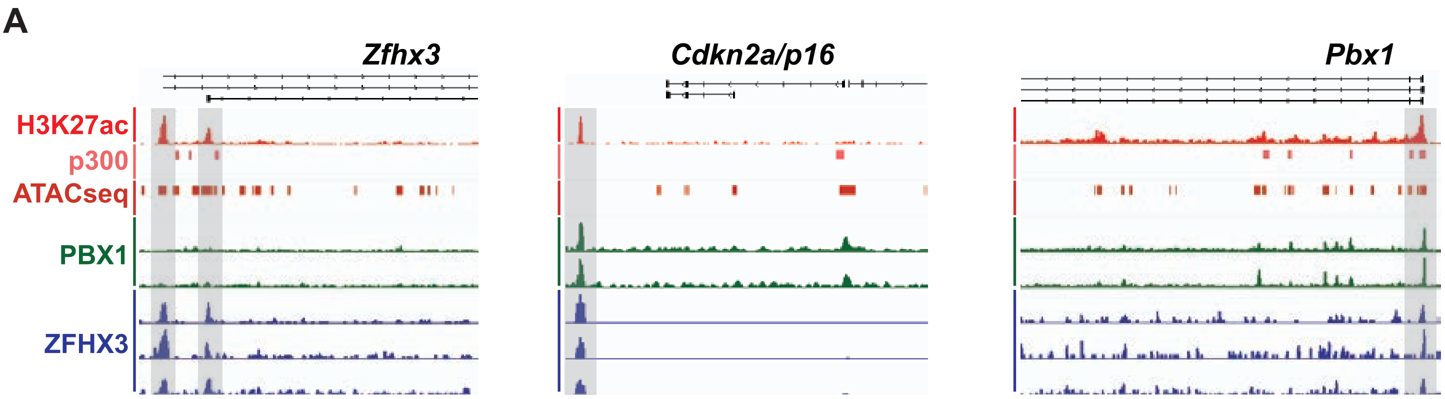
