## Supplemental Table 1 for "Cell Cycle Arrest of a ‘Zippering’ Epithelial Cell Cluster Shapes the Face and is Disrupted in Craniofacial Disorders"

| CHROM | POS | REF | ALT | CHR_POS_RE | Func.refGene | Gene.refGen |
|---|---|---|---|---|---|---|
| chr16 | 72797104 | TAGAG | T | chr16_72797 | exonic | ZFHX3 |
| chr7 | 50782497 | G | A | chr7_507824 | exonic | GRB10 |
| chr2 | 213147743 | T | C | chr2_213147 | exonic | IKZF2 |
| chr2 | 213007730 | C | A | chr2_213007 | exonic | IKZF2 |
| chr17 | 7894218 | G | A | chr17_78942 | exonic | CHD3 |
| chr1 | 8656255 | G | A | chr1_865625 | exonic | RERE |
| chr17 | 7903881 | C | T | chr17_79038 | exonic | CHD3 |
| chr7 | 134960013 | G | A | chr7_134960 | exonic | CALD1 |
| chr11 | 20103261 | C | T | chr11_20103 | exonic | NAV2 |
| chr19 | 58393144 | G | A | chr19_58393 | exonic | RPS5 |
| chr11 | 19939678 | A | C | chr11_19939 | exonic | NAV2 |
| chr2 | 213147743 | T | C | chr2_213147 | exonic | IKZF2 |
| chr1 | 8359852 | T | G | chr1_835985 | exonic | RERE |
| chr12 | 310904 | G | C | chr12_31090 | exonic | KDM5A |
| chr14 | 53953389 | A | C | chr14_53953 | exonic | BMP4 |
| chr17 | 72121467 | A | G | chr17_72121 | exonic | SOX9 |
| chr17 | 7911497 | A | C | chr17_79114 | exonic | CHD3 |
| chr19 | 3979331 | T | C | chr19_39793 | exonic | EEF2 |
| chr11 | 19939678 | A | C | chr11_19939 | exonic | NAV2 |
| chr1 | 8360358 | G | A | chr1_836035 | exonic | RERE |
| chr3 | 189890833 | C | T | chr3_189890 | exonic | TP63 |
| chr14 | 53956759 | C | T | chr14_53956 | exonic | BMP4 |
| chr16 | 72957864 | C | A | chr16_72957 | exonic | ZFHX3 |
| chr11 | 20097604 | A | C | chr11_20097 | exonic | NAV2 |
| chr3 | 55479451 | A | T | chr3_554794 | exonic | WNT5A |
| chr11 | 19948893 | C | T | chr11_19948 | exonic | NAV2 |
| chr3 | 55474467 | T | C | chr3_554744 | exonic | WNT5A |
| chr19 | 17861345 | C | T | chr19_17861 | exonic | RPL18A |
| chr2 | 172100590 | G | A | chr2_172100 | exonic | DLX2 |
| chr12 | 292794 | C | T | chr12_29279 | exonic | KDM5A |
| chr11 | 19939678 | A | C | chr11_19939 | exonic | NAV2 |
| chr2 | 172102224 | C | G | chr2_172102 | exonic | DLX2 |
| chr16 | 72950938 | C | T | chr16_72950 | exonic | ZFHX3 |
| chr19 | 58393144 | G | A | chr19_58393 | exonic | RPS5 |
| chr21 | 14965353 | C | T | chr21_14965 | exonic | NRIP1 |
| chr21 | 14964894 | G | A | chr21_14964 | exonic | NRIP1 |
| chr1 | 8362695 | C | G | chr1_836269 | exonic | RERE |
| chr10 | 61902261 | T | A | chr10_61902 | exonic | ARID5B |
| chr12 | 328993 | G | A | chr12_32899 | exonic | KDM5A |
| chr3 | 189631577 | G | A | chr3_189631 | exonic | TP63 |
| chr11 | 20045262 | G | A | chr11_20045 | exonic | NAV2 |
| chr2 | 221443552 | C | T | chr2_221443 | exonic | EPHA4 |

| | | | | | | |
|---|---|---|---|---|---|---|
| chr2 | 213147743 | T | C | chr2_213147 | exonic | IKZF2 |
| chr5 | 83539132 | A | T | chr5_835391 | exonic | VCAN |
| chr6 | 7576386 | G | A | chr6_757638 | exonic | DSP |
| chr6 | 7563752 | T | C | chr6_756375 | exonic | DSP |
| chr3 | 189890833 | C | T | chr3_189890 | exonic | TP63 |
| chr16 | 72957859 | C | A | chr16_72957 | exonic | ZFHX3 |
| chr5 | 132262725 | GGGC | G | chr5_132262 | exonic | PDLIM4 |
| chr7 | 134933250 | G | A | chr7_134933 | exonic | CALD1 |
| chr14 | 53951951 | G | C | chr14_53951 | exonic | BMP4 |
| chr1 | 8365860 | C | T | chr1_836586 | exonic | RERE |
| chr12 | 333510 | C | T | chr12_33351 | exonic | KDM5A |
| chr19 | 3981365 | C | T | chr19_39813 | exonic | EEF2 |
| chr5 | 83521209 | A | G | chr5_835212 | exonic | VCAN |
| chr7 | 50782526 | A | G | chr7_507825 | exonic | GRB10 |
| chr5 | 83520237 | C | T | chr5_835202 | exonic | VCAN |
| chr14 | 99257563 | G | A | chr14_99257 | exonic | BCL11B |
| chr6 | 7567376 | C | A | chr6_756737 | exonic | DSP |
| chr21 | 14964958 | C | A | chr21_14964 | exonic | NRIP1 |
| chr5 | 83540664 | T | C | chr5_835406 | exonic | VCAN |
| chr6 | 7581408 | G | A | chr6_758140 | exonic | DSP |
| chr6 | 7585729 | C | G | chr6_758572 | exonic | DSP |
| chr16 | 72788109 | T | TTGC | chr16_72788 | exonic | ZFHX3 |
| chr2 | 213147743 | T | C | chr2_213147 | exonic | IKZF2 |
| chr12 | 352218 | T | C | chr12_35221 | exonic | KDM5A |
| chr6 | 7583533 | A | G | chr6_758353 | exonic | DSP |
| chr11 | 19939678 | A | C | chr11_19939 | exonic | NAV2 |
| chr12 | 297184 | T | C | chr12_29718 | exonic | KDM5A |
| chr19 | 3982291 | C | T | chr19_39822 | exonic | EEF2 |
| chr7 | 134933133 | G | A | chr7_134933 | exonic | CALD1 |
| chr7 | 50732369 | C | G | chr7_507323 | splicing | GRB10 |
| chr5 | 83580408 | C | T | chr5_835804 | exonic | VCAN |
| chr6 | 44249822 | C | T | chr6_442498 | exonic | HSP90AB1 |
| chr6 | 7568461 | T | C | chr6_756846 | exonic | DSP |
| chr11 | 20103261 | C | T | chr11_20103 | exonic | NAV2 |
| chr2 | 213147743 | T | C | chr2_213147 | exonic | IKZF2 |
| chr2 | 213147743 | T | C | chr2_213147 | exonic | IKZF2 |
| chr16 | 72794944 | T | A | chr16_72794 | exonic | ZFHX3 |
| chr11 | 20044267 | G | T | chr11_20044 | exonic | NAV2 |
| chr5 | 68280645 | C | T | chr5_682806 | exonic | PIK3R1 |
| chr5 | 83542260 | A | G | chr5_835422 | exonic | VCAN |
| chr6 | 7583569 | A | G | chr6_758356 | exonic | DSP |
| chr17 | 72121467 | A | G | chr17_72121 | exonic | SOX9 |
| chr6 | 44249473 | C | T | chr6_442494 | exonic | HSP90AB1 |

| | | | | | | |
|---|---|---|---|---|---|---|
| chr11 | 19892487 | G | A | chr11_19892 | exonic | NAV2 |
| chr5 | 128084233 | C | CGCT | chr5_128084 | exonic | SLC12A2 |
| chr11 | 19713850 | A | G | chr11_19713 | exonic | NAV2 |
| chr14 | 53952059 | C | A | chr14_53952 | exonic | BMP4 |
| chr14 | 53951951 | G | C | chr14_53951 | exonic | BMP4 |
| chr6 | 158503376 | G | A | chr6_158503 | exonic | TULP4 |
| chr1 | 8358536 | C | G | chr1_835853 | exonic | RERE |
| chr2 | 213147743 | T | C | chr2_213147 | exonic | IKZF2 |
| chr7 | 134928798 | G | A | chr7_134928 | exonic | CALD1 |
| chr17 | 72124155 | C | T | chr17_72124 | exonic | SOX9 |
| chr1 | 8359782 | GCGCTCC | G | chr1_835978 | exonic | RERE |
| chr17 | 7894218 | G | A | chr17_78942 | exonic | CHD3 |
| chr14 | 53950654 | G | A | chr14_53950 | exonic | BMP4 |
| chr14 | 53956801 | C | T | chr14_53956 | exonic | BMP4 |
| chr7 | 50782497 | G | A | chr7_507824 | exonic | GRB10 |
| chr11 | 20103261 | C | T | chr11_20103 | exonic | NAV2 |
| chr11 | 19879928 | AAGC | A | chr11_19879 | exonic | NAV2 |
| chr6 | 7580602 | A | C | chr6_758060 | exonic | DSP |
| chr17 | 7893852 | C | T | chr17_78938 | exonic | CHD3 |
| chr1 | 164792621 | A | AGCG | chr1_164792 | exonic | PBX1 |
| chr6 | 7575307 | G | T | chr6_757530 | exonic | DSP |
| chr6 | 158503216 | T | C | chr6_158503 | exonic | TULP4 |
| chr7 | 50782518 | G | GCGCGTGGA | chr7_507825 | exonic | GRB10 |
| chr2 | 213147743 | T | C | chr2_213147 | exonic | IKZF2 |
| chr11 | 19939678 | A | C | chr11_19939 | exonic | NAV2 |
| chr16 | 72793368 | G | A | chr16_72793 | exonic | ZFHX3 |
| chr19 | 58393144 | G | A | chr19_58393 | exonic | RPS5 |
| chr1 | 8359817 | CCTT | C | chr1_835981 | exonic | RERE |
| chr5 | 132257745 | C | T | chr5_132257 | exonic | PDLIM4 |
| chr5 | 128178614 | C | T | chr5_128178 | exonic | SLC12A2 |
| chr6 | 158504006 | G | A | chr6_158504 | exonic | TULP4 |
| chr21 | 14966872 | T | C | chr21_14966 | exonic | NRIP1 |
| chr16 | 72958515 | G | A | chr16_72958 | exonic | ZFHX3 |
| chr7 | 50674484 | G | A | chr7_506744 | exonic | GRB10 |
| chr6 | 7581408 | G | A | chr6_758140 | exonic | DSP |
| chr6 | 7558121 | G | C | chr6_755812 | exonic | DSP |
| chr6 | 7559311 | G | A | chr6_755931 | exonic | DSP |
| chr5 | 68226704 | C | T | chr5_682267 | exonic | PIK3R1 |
| chr6 | 7579956 | C | T | chr6_757995 | exonic | DSP |
| chr14 | 53951861 | T | C | chr14_53951 | exonic | BMP4 |
| chr16 | 72788110 | TGCTGCTGCT | T | chr16_72788 | exonic | ZFHX3 |
| chr16 | 72788498 | CCTT | C | chr16_72788 | exonic | ZFHX3 |
| chr6 | 158504006 | G | A | chr6_158504 | exonic | TULP4 |

| | | | | | | |
|---|---|---|---|---|---|---|
| chr14 | 53953371 | G | A | chr14_53953 | exonic | BMP4 |
| chr2 | 213147743 | T | C | chr2_213147 | exonic | IKZF2 |
| chr6 | 7581408 | G | A | chr6_758140 | exonic | DSP |
| chr19 | 3981371 | C | T | chr19_39813 | exonic | EEF2 |
| chr5 | 68273401 | C | A | chr5_682734 | exonic | PIK3R1 |
| chr6 | 7585798 | C | T | chr6_758579 | exonic | DSP |
| chr6 | 7584143 | C | G | chr6_758414 | exonic | DSP |
| chr22 | 39318520 | G | A | chr22_39318 | exonic | RPL3 |
| chr16 | 72950744 | C | T | chr16_72950 | exonic | ZFHX3 |
| chr7 | 134950516 | T | C | chr7_134950 | splicing | CALD1 |
| chr3 | 189869371 | C | T | chr3_189869 | exonic | TP63 |
| chr11 | 20045321 | C | T | chr11_20045 | exonic | NAV2 |
| chr16 | 72796036 | T | C | chr16_72796 | exonic | ZFHX3 |
| chr17 | 7889055 | AG | A | chr17_78890 | exonic | CHD3 |
| chr2 | 221429970 | G | A | chr2_221429 | exonic | EPHA4 |
| chr6 | 158503446 | T | A | chr6_158503 | exonic | TULP4 |
| chr3 | 189868672 | A | G | chr3_189868 | exonic | TP63 |
| chr5 | 83537051 | GAATCTA | G | chr5_835370 | exonic | VCAN |
| chr19 | 58393144 | G | A | chr19_58393 | exonic | RPS5 |
| chr5 | 83541698 | G | A | chr5_835416 | exonic | VCAN |
| chr5 | 128134228 | G | A | chr5_128134 | exonic | SLC12A2 |
| chr6 | 7584143 | C | G | chr6_758414 | exonic | DSP |
| chr7 | 50782508 | G | T | chr7_507825 | exonic | GRB10 |
| chr16 | 72793589 | G | C | chr16_72793 | exonic | ZFHX3 |
| chr22 | 39315448 | C | G | chr22_39315 | exonic | RPL3 |
| chr8 | 73293606 | C | G | chr8_732936 | exonic | RPL7 |
| chr21 | 14964949 | G | C | chr21_14964 | exonic | NRIP1 |
| chr5 | 83520059 | A | G | chr5_835200 | exonic | VCAN |
| chr5 | 83540483 | G | C | chr5_835404 | exonic | VCAN |
| chr16 | 72794529 | G | A | chr16_72794 | exonic | ZFHX3 |
| chr7 | 134933133 | G | A | chr7_134933 | exonic | CALD1 |
| chr6 | 7584143 | C | G | chr6_758414 | exonic | DSP |
| chr7 | 50616274 | CATG | C | chr7_506162 | exonic | GRB10 |
| chr1 | 8361767 | G | A | chr1_836176 | exonic | RERE |
| chr12 | 354122 | G | A | chr12_35412 | exonic | KDM5A |
| chr11 | 20118173 | C | G | chr11_20118 | exonic | NAV2 |
| chr16 | 72957552 | T | C | chr16_72957 | exonic | ZFHX3 |
| chr21 | 14966047 | G | A | chr21_14966 | exonic | NRIP1 |
| chr19 | 3980644 | CGGA | C | chr19_39806 | exonic | EEF2 |
| chr7 | 134933133 | G | A | chr7_134933 | exonic | CALD1 |
| chr19 | 3982979 | A | G | chr19_39829 | exonic | EEF2 |
| chr21 | 14964763 | A | C | chr21_14964 | exonic | NRIP1 |
| chr9 | 36210654 | T | C | chr9_362106 | exonic | CLTA |

| | | | | | | |
|---|---|---|---|---|---|---|
| chr5 | 68273398 | C | T | chr5_682733 | exonic | PIK3R1 |
| chr6 | 7585571 | A | G | chr6_758557 | exonic | DSP |
| chr16 | 72788089 | G | GGCTGCTGCT | chr16_72788 | exonic | ZFHX3 |
| chr12 | 307786 | CT | C | chr12_30778 | exonic | KDM5A |
| chr21 | 14964949 | G | C | chr21_14964 | exonic | NRIP1 |
| chr11 | 19351007 | G | A | chr11_19351 | exonic | NAV2 |
| chr16 | 72796377 | A | G | chr16_72796 | exonic | ZFHX3 |
| chr5 | 128134228 | G | A | chr5_128134 | exonic | SLC12A2 |
| chr21 | 14967333 | G | A | chr21_14967 | exonic | NRIP1 |
| chr1 | 8359782 | GCGCTCCCGC | G | chr1_835978 | exonic | RERE |
| chr9 | 36191078 | G | C | chr9_361910 | exonic | CLTA |
| chr16 | 72788452 | G | A | chr16_72788 | exonic | ZFHX3 |
| chr6 | 44249514 | A | C | chr6_442495 | exonic | HSP90AB1 |
| chr6 | 44249514 | A | C | chr6_442495 | exonic | HSP90AB1 |
| chr19 | 3980644 | CGGA | C | chr19_39806 | exonic | EEF2 |
| chr16 | 72797486 | TTGCTGTTGC | T | chr16_72797 | exonic | ZFHX3 |
| chr12 | 328984 | G | A | chr12_32898 | exonic | KDM5A |
| chr15 | 36892252 | A | G | chr15_36892 | exonic | MEIS2 |
| chr10 | 62000151 | G | A | chr10_62000 | exonic | ARID5B |
| chr5 | 83540365 | AAGC | A | chr5_835403 | exonic | VCAN |
| chr16 | 72793589 | G | C | chr16_72793 | exonic | ZFHX3 |
| chr16 | 72797935 | C | A | chr16_72797 | exonic | ZFHX3 |
| chr12 | 310904 | G | C | chr12_31090 | exonic | KDM5A |
| chr14 | 99175335 | C | T | chr14_99175 | exonic | BCL11B |
| chr16 | 72794967 | A | G | chr16_72794 | exonic | ZFHX3 |
| chr16 | 72798037 | C | T | chr16_72798 | exonic | ZFHX3 |
| chr5 | 83520054 | CAA | C | chr5_835200 | exonic | VCAN |
| chr9 | 36191078 | G | C | chr9_361910 | exonic | CLTA |
| chr16 | 72788210 | A | T | chr16_72788 | exonic | ZFHX3 |
| chr2 | 213147743 | T | C | chr2_213147 | exonic | IKZF2 |
| chr7 | 134933133 | G | A | chr7_134933 | exonic | CALD1 |
| chr16 | 72788452 | G | A | chr16_72788 | exonic | ZFHX3 |
| chr7 | 134933133 | G | A | chr7_134933 | exonic | CALD1 |
| chr6 | 7581408 | G | A | chr6_758140 | exonic | DSP |
| chr6 | 158503301 | T | C | chr6_158503 | exonic | TULP4 |
| chr17 | 7911496 | A | G | chr17_79114 | exonic | CHD3 |
| chr11 | 20045262 | G | A | chr11_20045 | exonic | NAV2 |
| chr12 | 310904 | G | C | chr12_31090 | exonic | KDM5A |
| chr6 | 7576386 | G | A | chr6_757638 | exonic | DSP |
| chr19 | 3979402 | G | A | chr19_39794 | exonic | EEF2 |
| chr1 | 8359782 | GCGCTCCCGC | G | chr1_835978 | exonic | RERE |
| chr11 | 20103701 | C | A | chr11_20103 | exonic | NAV2 |
| chr12 | 354122 | G | A | chr12_35412 | exonic | KDM5A |

| | | | | | | |
|---|---|---|---|---|---|---|
| chr7 | 134935733 | CAAG | C | chr7_134935 | exonic | CALD1 |
| chr7 | 134933133 | G | A | chr7_134933 | exonic | CALD1 |
| chr12 | 310904 | G | C | chr12_31090 | exonic | KDM5A |
| chr12 | 354122 | G | A | chr12_35412 | exonic | KDM5A |
| chr21 | 14964949 | G | C | chr21_14964 | exonic | NRIP1 |
| chr12 | 354122 | G | A | chr12_35412 | exonic | KDM5A |
| chr6 | 158503404 | C | A | chr6_158503 | exonic | TULP4 |
| chr16 | 72796735 | G | T | chr16_72796 | exonic | ZFHX3 |
| chr1 | 8361046 | AGGGATGCG | A | chr1_836104 | exonic | RERE |
| chr16 | 72957552 | T | C | chr16_72957 | exonic | ZFHX3 |
| chr11 | 20045262 | G | A | chr11_20045 | exonic | NAV2 |
| chr5 | 83541015 | C | A | chr5_835410 | exonic | VCAN |
| chr1 | 8359782 | GCGCTCCCGC | G | chr1_835978 | exonic | RERE |
| chr5 | 83519745 | A | G | chr5_835197 | exonic | VCAN |
| chr1 | 8361767 | G | A | chr1_836176 | exonic | RERE |
| chr12 | 366037 | C | A | chr12_36603 | exonic | KDM5A |
| chr6 | 158503472 | C | T | chr6_158503 | exonic | TULP4 |
| chr14 | 99175206 | G | GCTC | chr14_99175 | exonic | BCL11B |
| chr11 | 19842873 | A | G | chr11_19842 | exonic | NAV2 |
| chr16 | 72959380 | C | T | chr16_72959 | exonic | ZFHX3 |
| chr6 | 158503472 | C | T | chr6_158503 | exonic | TULP4 |
| chr16 | 72787423 | G | T | chr16_72787 | exonic | ZFHX3 |
| chr16 | 72788584 | TGCTGCTGCT | T | chr16_72788 | exonic | ZFHX3 |
| chr6 | 158479781 | C | T | chr6_158479 | exonic | TULP4 |
| chr16 | 72958731 | GCCTCTTCCT | G | chr16_72958 | exonic | ZFHX3 |
| chr14 | 53950774 | C | T | chr14_53950 | exonic | BMP4 |
| chr10 | 62092093 | A | G | chr10_62092 | exonic | ARID5B |
| chr3 | 189869350 | C | T | chr3_189869 | exonic | TP63 |
| chr1 | 8358610 | TCTCCCGCTC | T | chr1_835861 | exonic | RERE |
| chr6 | 7579791 | G | A | chr6_757979 | exonic | DSP |
| chr1 | 8360116 | C | G | chr1_836011 | exonic | RERE |
| chr6 | 7559288 | G | T | chr6_755928 | exonic | DSP |
| chr16 | 72787819 | G | A | chr16_72787 | exonic | ZFHX3 |
| chr5 | 68226685 | G | A | chr5_682266 | exonic | PIK3R1 |
| chr7 | 134941128 | AAAG | A | chr7_134941 | exonic | CALD1 |
| chr5 | 83522015 | G | A | chr5_835220 | exonic | VCAN |
| chr6 | 7559288 | G | T | chr6_755928 | exonic | DSP |
| chr6 | 73518386 | G | A | chr6_735183 | exonic | EEF1A1 |
| chr11 | 19934013 | G | A | chr11_19934 | exonic | NAV2 |
| chr7 | 134933563 | C | T | chr7_134933 | exonic | CALD1 |
| chr16 | 72959157 | C | T | chr16_72959 | exonic | ZFHX3 |
| chr6 | 158504059 | G | A | chr6_158504 | exonic | TULP4 |
| chr1 | 8364141 | G | A | chr1_836414 | exonic | RERE |

| | | | | | | |
|---|---|---|---|---|---|---|
| chr16 | 72788584 | TGCTGCTGCT | T | chr16_72788 | exonic | ZFHX3 |
| chr14 | 53950774 | C | T | chr14_53950 | exonic | BMP4 |
| chr17 | 7894438 | G | A | chr17_78944 | exonic | CHD3 |
| chr3 | 189894266 | G | C | chr3_189894 | exonic | TP63 |
| chr6 | 7582879 | C | T | chr6_758287 | exonic | DSP |
| chr16 | 72950503 | T | G | chr16_72950 | exonic | ZFHX3 |
| chr14 | 53950334 | G | A | chr14_53950 | exonic | BMP4 |
| chr16 | 72958731 | GCCTCTTCCT | G | chr16_72958 | exonic | ZFHX3 |
| chr1 | 8360256 | G | A | chr1_836025 | exonic | RERE |
| chr3 | 189880092 | C | T | chr3_189880 | exonic | TP63 |
| chr16 | 72796036 | T | C | chr16_72796 | exonic | ZFHX3 |
| chr5 | 83542242 | T | A | chr5_835422 | exonic | VCAN |
| chr17 | 7899356 | G | A | chr17_78993 | exonic | CHD3 |
| chr22 | 39313226 | G | A | chr22_39313 | exonic | RPL3 |
| chr6 | 7579954 | G | C | chr6_757995 | exonic | DSP |
| chr16 | 72795474 | G | A | chr16_72795 | exonic | ZFHX3 |
| chr7 | 134928897 | A | G | chr7_134928 | exonic | CALD1 |
| chr6 | 158502637 | C | T | chr6_158502 | exonic | TULP4 |
| chr10 | 62085838 | C | T | chr10_62085 | exonic | ARID5B |
| chr7 | 134928897 | A | G | chr7_134928 | exonic | CALD1 |
| chr16 | 72787402 | G | A | chr16_72787 | exonic | ZFHX3 |
| chr6 | 7559282 | G | A | chr6_755928 | exonic | DSP |
| chr16 | 72787423 | G | T | chr16_72787 | exonic | ZFHX3 |
| chr7 | 134928897 | A | G | chr7_134928 | exonic | CALD1 |
| chr1 | 8361033 | G | A | chr1_836103 | exonic | RERE |
| chr5 | 83537170 | TGAAGAA | T | chr5_835371 | exonic | VCAN |
| chr6 | 158502637 | C | T | chr6_158502 | exonic | TULP4 |
| chr1 | 8358481 | C | T | chr1_835848 | exonic | RERE |
| chr6 | 158503472 | C | T | chr6_158503 | exonic | TULP4 |
| chr16 | 72959877 | C | T | chr16_72959 | exonic | ZFHX3 |
| chr1 | 8360670 | T | A | chr1_836067 | exonic | RERE |
| chr16 | 72957733 | G | A | chr16_72957 | exonic | ZFHX3 |
| chr14 | 53950334 | G | A | chr14_53950 | exonic | BMP4 |
| chr7 | 134928897 | A | G | chr7_134928 | exonic | CALD1 |
| chr1 | 8358481 | C | T | chr1_835848 | exonic | RERE |
| chr11 | 19934013 | G | A | chr11_19934 | exonic | NAV2 |
| chr2 | 216694652 | G | A | chr2_216694 | exonic | IGFBP5 |
| chr1 | 8656232 | CTCTCGGTCC | C | chr1_865623 | exonic | RERE |
| chr6 | 7580212 | G | T | chr6_758021 | exonic | DSP |
| chr2 | 221563982 | C | G | chr2_221563 | exonic | EPHA4 |
| chr14 | 53950453 | C | T | chr14_53950 | exonic | BMP4 |
| chr3 | 189894266 | G | C | chr3_189894 | exonic | TP63 |
| chr7 | 134928897 | A | G | chr7_134928 | exonic | CALD1 |

| | | | | | | |
|---|---|---|---|---|---|---|
| chr10 | 62050980 | G | A | chr10_62050 | exonic | ARID5B |
| chr1 | 8358481 | C | T | chr1_835848 | exonic | RERE |
| chr1 | 8362685 | T | C | chr1_836268 | exonic | RERE |

| GeneDetail.re | ExonicFunc.re | AAChange.ref | loeuf | pLI | SIFT_score | SIFT_convert |
|---|---|---|---|---|---|---|
| . | stopgain | ZFHX3:NM_0 | 0.136 | 1 | NA | . |
| NM_0013508 | nonsynonym | GRB10:NM_0 | 0.335 | 0.939 | NA | . |
| . | nonsynonym | IKZF2:NM_00 | 0.286 | 0.987 | 0.043 | 0.913 |
| . | nonsynonym | IKZF2:NM_00 | 0.286 | 0.987 | 0.01 | 0.575 |
| . | nonsynonym | CHD3:NM_00 | 0.147 | 1 | 0.025 | 0.473 |
| . | nonsynonym | RERE:NM_01 | 0.12 | 1 | 0 | 0.913 |
| . | nonsynonym | CHD3:NM_00 | 0.147 | 1 | 0.001 | 0.785 |
| . | nonsynonym | CALD1:NM_0 | 0.258 | 0.999 | 0.021 | 0.913 |
| . | nonsynonym | NAV2:NM_18 | 0.247 | 1 | 0.002 | 0.913 |
| dist=220 | nonsynonym | RPS5:NM_00 | 0.322 | 0.948 | 0.043 | 0.414 |
| . | nonsynonym | NAV2:NM_18 | 0.247 | 1 | 0.005 | 0.785 |
| . | nonsynonym | IKZF2:NM_00 | 0.286 | 0.987 | 0.043 | 0.913 |
| . | nonsynonym | RERE:NM_01 | 0.12 | 1 | 0.003 | 0.722 |
| . | nonsynonym | KDM5A:NM_ | 0.163 | 1 | 0 | 0.913 |
| NM_0013479 | nonsynonym | BMP4:NM_00 | 0.334 | 0.956 | NA | . |
| dist=674 | nonsynonym | SOX9:NM_00 | 0.168 | 0.998 | 0.022 | 0.486 |
| . | nonsynonym | CHD3:NM_00 | 0.147 | 1 | 0.007 | 0.599 |
| . | nonsynonym | EEF2:NM_00 | 0.172 | 1 | 0.01 | 0.565 |
| . | nonsynonym | NAV2:NM_18 | 0.247 | 1 | 0.005 | 0.785 |
| . | nonsynonym | RERE:NM_01 | 0.12 | 1 | 0.001 | 0.785 |
| . | nonsynonym | TP63:NM_00 | 0.267 | 0.997 | 0.034 | 0.444 |
| NM_001202: | nonsynonym | BMP4:NM_00 | 0.334 | 0.956 | NA | . |
| . | nonsynonym | ZFHX3:NM_0 | 0.136 | 1 | 0.008 | 0.586 |
| . | nonsynonym | NAV2:NM_18 | 0.247 | 1 | 0.013 | 0.722 |
| . | stopgain | WNT5A:NM_ | 0.269 | 0.987 | NA | . |
| . | nonsynonym | NAV2:NM_18 | 0.247 | 1 | 0 | 0.913 |
| . | nonsynonym | WNT5A:NM_ | 0.269 | 0.987 | 0.002 | 0.722 |
| . | nonsynonym | RPL18A:NM_ | 0.243 | 0.982 | 0.039 | 0.425 |
| . | nonsynonym | DLX2:NM_00 | 0.254 | 0.978 | 0.006 | 0.614 |
| . | nonsynonym | KDM5A:NM_ | 0.163 | 1 | 0.05 | 0.396 |
| . | nonsynonym | NAV2:NM_18 | 0.247 | 1 | 0.005 | 0.785 |
| dist=782 | nonsynonym | DLX2:NM_00 | 0.254 | 0.978 | 0.004 | 0.654 |
| . | nonsynonym | ZFHX3:NM_0 | 0.136 | 1 | 0.026 | 0.913 |
| dist=220 | nonsynonym | RPS5:NM_00 | 0.322 | 0.948 | 0.043 | 0.414 |
| . | nonsynonym | NRIP1:NM_00 | 0.279 | 0.994 | 0.008 | 0.586 |
| . | nonsynonym | NRIP1:NM_00 | 0.279 | 0.994 | 0.004 | 0.654 |
| . | nonsynonym | RERE:NM_01 | 0.12 | 1 | 0.025 | 0.722 |
| . | nonsynonym | ARID5B:NM_ | 0.11 | 1 | 0.011 | 0.555 |
| . | nonsynonym | KDM5A:NM_ | 0.163 | 1 | 0.014 | 0.532 |
| . | nonsynonym | TP63:NM_00 | 0.267 | 0.997 | 0.004 | 0.682 |
| . | nonsynonym | NAV2:NM_18 | 0.247 | 1 | 0.002 | 0.722 |
| . | nonsynonym | EPHA4:NM_0 | 0.175 | 1 | 0.02 | 0.507 |

| | | | | | | |
|---|---|---|---|---|---|---|
| . | nonsynonymo | IKZF2:NM_00 | 0.286 | 0.987 | 0.043 | 0.913 |
| . | nonsynonymo | VCAN:NM_00 | 0.199 | 1 | 0.006 | 0.654 |
| . | nonsynonymo | DSP:NM_001 | 0.26 | 1 | 0.005 | 0.654 |
| . | nonsynonymo | DSP:NM_001 | 0.26 | 1 | 0.004 | 0.654 |
| . | nonsynonymo | TP63:NM_00 | 0.267 | 0.997 | 0.034 | 0.444 |
| . | stopgain | ZFHX3:NM_0 | 0.136 | 1 | NA | . |
| . | nonframeshif | PDLIM4:NM_ | 0.242 | 0.983 | NA | . |
| . | nonsynonymo | CALD1:NM_0 | 0.258 | 0.999 | 0.004 | 0.654 |
| . | nonsynonymo | BMP4:NM_00 | 0.334 | 0.956 | 0.002 | 0.785 |
| NM_0010426 | nonsynonymo | RERE:NM_01 | 0.12 | 1 | 0.025 | 0.482 |
| . | nonsynonymo | KDM5A:NM_ | 0.163 | 1 | 0.031 | 0.45 |
| . | nonsynonymo | EEF2:NM_001 | 0.172 | 1 | 0.039 | 0.425 |
| . | nonsynonymo | VCAN:NM_00 | 0.199 | 1 | 0 | 0.913 |
| NM_0013508 | nonsynonymo | GRB10:NM_0 | 0.335 | 0.939 | NA | . |
| . | nonsynonymo | VCAN:NM_00 | 0.199 | 1 | 0.001 | 0.785 |
| . | nonsynonymo | BCL11B:NM_ | 0.282 | 0.989 | 0.001 | 0.785 |
| . | nonsynonymo | DSP:NM_001 | 0.26 | 1 | 0 | 0.913 |
| . | nonsynonymo | NRIP1:NM_00 | 0.279 | 0.994 | 0.018 | 0.507 |
| . | nonsynonymo | VCAN:NM_00 | 0.199 | 1 | 0 | 0.913 |
| . | nonsynonymo | DSP:NM_004 | 0.26 | 1 | 0.025 | 0.473 |
| . | nonsynonymo | DSP:NM_001 | 0.26 | 1 | 0 | 0.913 |
| . | nonframeshif | ZFHX3:NM_0 | 0.136 | 1 | NA | . |
| . | nonsynonymo | IKZF2:NM_00 | 0.286 | 0.987 | 0.043 | 0.913 |
| . | nonsynonymo | KDM5A:NM_ | 0.163 | 1 | 0.019 | 0.501 |
| . | nonsynonymo | DSP:NM_001 | 0.26 | 1 | 0.017 | 0.525 |
| . | nonsynonymo | NAV2:NM_18 | 0.247 | 1 | 0.005 | 0.785 |
| . | nonsynonymo | KDM5A:NM_ | 0.163 | 1 | 0.016 | 0.519 |
| dist=216 | nonsynonymo | EEF2:NM_001 | 0.172 | 1 | 0.009 | 0.575 |
| . | nonsynonymo | CALD1:NM_0 | 0.258 | 0.999 | 0 | 0.913 |
| NM_0013508 | . | . | 0.335 | 0.939 | NA | . |
| . | nonsynonymo | VCAN:NM_00 | 0.199 | 1 | 0 | 0.913 |
| . | nonsynonymo | HSP90AB1:NM | 0.213 | 0.999 | 0.03 | 0.454 |
| . | nonsynonymo | DSP:NM_001 | 0.26 | 1 | 0.013 | 0.565 |
| . | nonsynonymo | NAV2:NM_18 | 0.247 | 1 | 0.002 | 0.913 |
| . | nonsynonymo | IKZF2:NM_00 | 0.286 | 0.987 | 0.043 | 0.913 |
| . | nonsynonymo | IKZF2:NM_00 | 0.286 | 0.987 | 0.043 | 0.913 |
| . | nonsynonymo | ZFHX3:NM_0 | 0.136 | 1 | 0.008 | 0.586 |
| . | nonsynonymo | NAV2:NM_18 | 0.247 | 1 | 0 | 0.913 |
| . | nonsynonymo | PIK3R1:NM_1 | 0.228 | 1 | 0.001 | 0.785 |
| . | nonsynonymo | VCAN:NM_00 | 0.199 | 1 | 0 | 0.913 |
| . | nonsynonymo | DSP:NM_001 | 0.26 | 1 | 0.007 | 0.599 |
| dist=674 | nonsynonymo | SOX9:NM_00 | 0.168 | 0.998 | 0.022 | 0.486 |
| . | nonsynonymo | HSP90AB1:NM | 0.213 | 0.999 | 0.01 | 0.565 |

| | | | | | | |
|---|---|---|---|---|---|---|
| . | nonsynonym | NAV2:NM_18 | 0.247 | 1 | 0.006 | 0.632 |
| . | nonframeshif | SLC12A2:NM_ | 0.31 | 0.962 NA | . | |
| . | nonsynonym | NAV2:NM_18 | 0.247 | 1 | 0.008 | 0.586 |
| NM_001347 | nonsynonym | BMP4:NM_00 | 0.334 | 0.956 | 0.032 | 0.722 |
| . | nonsynonym | BMP4:NM_00 | 0.334 | 0.956 | 0.002 | 0.785 |
| . | nonsynonym | TULP4:NM_0 | 0.133 | 1 | 0 | 0.913 |
| . | nonsynonym | RERE:NM_01 | 0.12 | 1 | 0.041 | 0.419 |
| . | nonsynonym | IKZF2:NM_00 | 0.286 | 0.987 | 0.043 | 0.913 |
| . | nonsynonym | CALD1:NM_0 | 0.258 | 0.999 | 0.004 | 0.913 |
| . | nonsynonym | SOX9:NM_00 | 0.168 | 0.998 | 0.001 | 0.785 |
| . | nonframeshif | RERE:NM_01 | 0.12 | 1 NA | . | |
| . | nonsynonym | CHD3:NM_00 | 0.147 | 1 | 0.025 | 0.473 |
| . | nonsynonym | BMP4:NM_00 | 0.334 | 0.956 | 0 | 0.913 |
| NM_001202: | nonsynonym | BMP4:NM_00 | 0.334 | 0.956 NA | . | |
| NM_001350 | nonsynonym | GRB10:NM_0 | 0.335 | 0.939 NA | . | |
| . | nonsynonym | NAV2:NM_18 | 0.247 | 1 | 0.002 | 0.913 |
| . | nonframeshif | NAV2:NM_18 | 0.247 | 1 NA | . | |
| . | nonsynonym | DSP:NM_004 | 0.26 | 1 | 0.027 | 0.465 |
| . | nonsynonym | CHD3:NM_00 | 0.147 | 1 | 0.004 | 0.654 |
| . | nonframeshif | PBX1:NM_00 | 0.255 | 0.995 NA | . | |
| . | nonsynonym | DSP:NM_001 | 0.26 | 1 | 0.001 | 0.785 |
| . | nonsynonym | TULP4:NM_0 | 0.133 | 1 | 0 | 0.913 |
| NM_001350 | frameshift in | GRB10:NM_0 | 0.335 | 0.939 NA | . | |
| . | nonsynonym | IKZF2:NM_00 | 0.286 | 0.987 | 0.043 | 0.913 |
| . | nonsynonym | NAV2:NM_18 | 0.247 | 1 | 0.005 | 0.785 |
| . | nonsynonym | ZFHX3:NM_0 | 0.136 | 1 | 0.003 | 0.722 |
| dist=220 | nonsynonym | RPS5:NM_00 | 0.322 | 0.948 | 0.043 | 0.414 |
| . | nonframeshif | RERE:NM_01 | 0.12 | 1 NA | . | |
| . | nonsynonym | PDLIM4:NM_ | 0.242 | 0.983 | 0.007 | 0.654 |
| . | nonsynonym | SLC12A2:NM_ | 0.31 | 0.962 | 0.046 | 0.406 |
| . | nonsynonym | TULP4:NM_0 | 0.133 | 1 | 0.042 | 0.416 |
| . | nonsynonym | NRIP1:NM_00 | 0.279 | 0.994 | 0 | 0.913 |
| . | nonsynonym | ZFHX3:NM_0 | 0.136 | 1 | 0.042 | 0.416 |
| . | nonsynonym | GRB10:NM_0 | 0.335 | 0.939 | 0.02 | 0.539 |
| . | nonsynonym | DSP:NM_004 | 0.26 | 1 | 0.025 | 0.473 |
| . | nonsynonym | DSP:NM_001 | 0.26 | 1 | 0.006 | 0.614 |
| . | nonsynonym | DSP:NM_001 | 0.26 | 1 | 0.027 | 0.465 |
| . | nonsynonym | PIK3R1:NM_1 | 0.228 | 1 | 0.002 | 0.722 |
| . | nonsynonym | DSP:NM_001 | 0.26 | 1 | 0.013 | 0.539 |
| . | nonsynonym | BMP4:NM_00 | 0.334 | 0.956 | 0.045 | 0.913 |
| . | nonframeshif | ZFHX3:NM_0 | 0.136 | 1 NA | . | |
| . | nonframeshif | ZFHX3:NM_0 | 0.136 | 1 NA | . | |
| . | nonsynonym | TULP4:NM_0 | 0.133 | 1 | 0.042 | 0.416 |

| | | | | | | |
|---|---|---|---|---|---|---|
| NM_0013479 | nonsynonym | BMP4:NM_00 | 0.334 | 0.956 | NA | . |
| . | nonsynonym | IKZF2:NM_00 | 0.286 | 0.987 | 0.043 | 0.913 |
| . | nonsynonym | DSP:NM_004 | 0.26 | 1 | 0.025 | 0.473 |
| . | nonsynonym | EEF2:NM_00 | 0.172 | 1 | 0.016 | 0.519 |
| . | nonsynonym | PIK3R1:NM_1 | 0.228 | 1 | 0.035 | 0.586 |
| . | nonsynonym | DSP:NM_001 | 0.26 | 1 | 0 | 0.913 |
| . | nonsynonym | DSP:NM_001 | 0.26 | 1 | 0 | 0.913 |
| dist=532 | nonsynonym | RPL3:NM_00 | 0.239 | 0.994 | 0 | 0.913 |
| . | nonsynonym | ZFHX3:NM_0 | 0.136 | 1 | 0.001 | 0.785 |
| NM_004342: | . | . | 0.258 | 0.999 | NA | . |
| . | stopgain | TP63:NM_00 | 0.267 | 0.997 | NA | . |
| . | nonsynonym | NAV2:NM_18 | 0.247 | 1 | 0 | 0.913 |
| . | nonsynonym | ZFHX3:NM_0 | 0.136 | 1 | 0.032 | 0.45 |
| . | frameshift de | CHD3:NM_00 | 0.147 | 1 | NA | . |
| . | nonsynonym | EPHA4:NM_0 | 0.175 | 1 | 0.049 | 0.406 |
| . | nonsynonym | TULP4:NM_0 | 0.133 | 1 | 0 | 0.913 |
| . | nonsynonym | TP63:NM_00 | 0.267 | 0.997 | 0.04 | 0.486 |
| . | nonframeshif | VCAN:NM_00 | 0.199 | 1 | NA | . |
| dist=220 | nonsynonym | RPS5:NM_00 | 0.322 | 0.948 | 0.043 | 0.414 |
| . | nonsynonym | VCAN:NM_00 | 0.199 | 1 | 0.009 | 0.575 |
| . | nonsynonym | SLC12A2:NM_ | 0.31 | 0.962 | 0.003 | 0.682 |
| . | nonsynonym | DSP:NM_001 | 0.26 | 1 | 0 | 0.913 |
| NM_0013508 | nonsynonym | GRB10:NM_0 | 0.335 | 0.939 | NA | . |
| . | nonsynonym | ZFHX3:NM_0 | 0.136 | 1 | 0.002 | 0.722 |
| dist=141 | nonsynonym | RPL3:NM_00 | 0.239 | 0.994 | 0.001 | 0.785 |
| dist=996 | nonsynonym | RPL7:NM_00 | 0.181 | 0.996 | 0.001 | 0.785 |
| . | nonsynonym | NRIP1:NM_0 | 0.279 | 0.994 | 0.008 | 0.586 |
| . | nonsynonym | VCAN:NM_00 | 0.199 | 1 | 0.005 | 0.682 |
| . | nonsynonym | VCAN:NM_00 | 0.199 | 1 | 0.001 | 0.785 |
| . | nonsynonym | ZFHX3:NM_0 | 0.136 | 1 | 0.029 | 0.458 |
| . | nonsynonym | CALD1:NM_0 | 0.258 | 0.999 | 0 | 0.913 |
| . | nonsynonym | DSP:NM_001 | 0.26 | 1 | 0 | 0.913 |
| . | nonframeshif | GRB10:NM_0 | 0.335 | 0.939 | NA | . |
| . | nonsynonym | RERE:NM_01 | 0.12 | 1 | 0.001 | 0.913 |
| . | nonsynonym | KDM5A:NM_ | 0.163 | 1 | 0.043 | 0.414 |
| . | nonsynonym | NAV2:NM_18 | 0.247 | 1 | 0.006 | 0.654 |
| . | nonsynonym | ZFHX3:NM_0 | 0.136 | 1 | 0.003 | 0.682 |
| . | nonsynonym | NRIP1:NM_0 | 0.279 | 0.994 | 0 | 0.913 |
| . | nonframeshif | EEF2:NM_00 | 0.172 | 1 | NA | . |
| . | nonsynonym | CALD1:NM_0 | 0.258 | 0.999 | 0 | 0.913 |
| dist=407 | nonsynonym | EEF2:NM_00 | 0.172 | 1 | 0.001 | 0.785 |
| . | nonsynonym | NRIP1:NM_0 | 0.279 | 0.994 | 0.01 | 0.565 |
| . | nonsynonym | CLTA:NM_00 | 0.323 | 0.963 | 0.01 | 0.565 |

| | | | | | | |
|---|---|---|---|---|---|---|
| . | nonsynonymo | PIK3R1:NM_1 | 0.228 | 1 | 0.033 | 0.654 |
| . | nonsynonymo | DSP:NM_001 | 0.26 | 1 | 0.001 | 0.785 |
| . | nonframeshif | ZFHX3:NM_0 | 0.136 | 1 | NA | . |
| . | frameshift de | KDM5A:NM_ | 0.163 | 1 | NA | . |
| . | nonsynonymo | NRIP1:NM_0 | 0.279 | 0.994 | 0.008 | 0.586 |
| . | nonsynonymo | NAV2:NM_00 | 0.247 | 1 | 0.004 | 0.654 |
| . | nonsynonymo | ZFHX3:NM_0 | 0.136 | 1 | 0 | 0.913 |
| . | nonsynonymo | SLC12A2:NM_ | 0.31 | 0.962 | 0.003 | 0.682 |
| . | nonsynonymo | NRIP1:NM_0 | 0.279 | 0.994 | 0.004 | 0.654 |
| . | nonframeshif | RERE:NM_01 | 0.12 | 1 | NA | . |
| NM_0013112 | nonsynonymo | CLTA:NM_00 | 0.323 | 0.963 | 0.004 | 0.654 |
| . | nonsynonymo | ZFHX3:NM_0 | 0.136 | 1 | 0.006 | 0.632 |
| . | nonsynonymo | HSP90AB1:NI | 0.213 | 0.999 | 0.009 | 0.575 |
| . | nonsynonymo | HSP90AB1:NI | 0.213 | 0.999 | 0.009 | 0.575 |
| . | nonframeshif | EEF2:NM_00 | 0.172 | 1 | NA | . |
| . | nonframeshif | ZFHX3:NM_0 | 0.136 | 1 | NA | . |
| . | nonsynonymo | KDM5A:NM_ | 0.163 | 1 | 0 | 0.913 |
| NM_170674: | nonsynonymo | MEIS2:NM_0 | 0.184 | 0.999 | 0.004 | 0.654 |
| . | nonsynonymo | ARID5B:NM_ | 0.11 | 1 | 0.012 | 0.547 |
| . | nonframeshif | VCAN:NM_00 | 0.199 | 1 | NA | . |
| . | nonsynonymo | ZFHX3:NM_0 | 0.136 | 1 | 0.002 | 0.722 |
| . | nonsynonymo | ZFHX3:NM_0 | 0.136 | 1 | 0.025 | 0.486 |
| . | nonsynonymo | KDM5A:NM_ | 0.163 | 1 | 0 | 0.913 |
| . | nonsynonymo | BCL11B:NM_ | 0.282 | 0.989 | 0.008 | 0.599 |
| . | nonsynonymo | ZFHX3:NM_0 | 0.136 | 1 | 0.001 | 0.785 |
| . | nonsynonymo | ZFHX3:NM_0 | 0.136 | 1 | 0.038 | 0.428 |
| . | frameshift de | VCAN:NM_00 | 0.199 | 1 | NA | . |
| NM_0013112 | nonsynonymo | CLTA:NM_00 | 0.323 | 0.963 | 0.004 | 0.654 |
| . | nonsynonymo | ZFHX3:NM_0 | 0.136 | 1 | 0.015 | 0.525 |
| . | nonsynonymo | IKZF2:NM_00 | 0.286 | 0.987 | 0.043 | 0.913 |
| . | nonsynonymo | CALD1:NM_0 | 0.258 | 0.999 | 0 | 0.913 |
| . | nonsynonymo | ZFHX3:NM_0 | 0.136 | 1 | 0.006 | 0.632 |
| . | nonsynonymo | CALD1:NM_0 | 0.258 | 0.999 | 0 | 0.913 |
| . | nonsynonymo | DSP:NM_004 | 0.26 | 1 | 0.025 | 0.473 |
| . | nonsynonymo | TULP4:NM_0 | 0.133 | 1 | 0.001 | 0.785 |
| . | nonsynonymo | CHD3:NM_00 | 0.147 | 1 | 0.006 | 0.614 |
| . | nonsynonymo | NAV2:NM_18 | 0.247 | 1 | 0.002 | 0.722 |
| . | nonsynonymo | KDM5A:NM_ | 0.163 | 1 | 0 | 0.913 |
| . | nonsynonymo | DSP:NM_001 | 0.26 | 1 | 0.005 | 0.654 |
| . | nonsynonymo | EEF2:NM_00 | 0.172 | 1 | 0 | 0.913 |
| . | nonframeshif | RERE:NM_01 | 0.12 | 1 | NA | . |
| . | nonsynonymo | NAV2:NM_18 | 0.247 | 1 | 0.002 | 0.785 |
| . | nonsynonymo | KDM5A:NM_ | 0.163 | 1 | 0.043 | 0.414 |

| | | | | | | |
|---|---|---|---|---|---|---|
| . | nonframeshif | CALD1:NM_0 | 0.258 | 0.999 | NA | . |
| . | nonsynonym | CALD1:NM_0 | 0.258 | 0.999 | 0 | 0.913 |
| . | nonsynonym | KDM5A:NM_ | 0.163 | 1 | 0 | 0.913 |
| . | nonsynonym | KDM5A:NM_ | 0.163 | 1 | 0.043 | 0.414 |
| . | nonsynonym | NRIP1:NM_0 | 0.279 | 0.994 | 0.008 | 0.586 |
| . | nonsynonym | KDM5A:NM_ | 0.163 | 1 | 0.043 | 0.414 |
| . | nonsynonym | TULP4:NM_0 | 0.133 | 1 | 0.002 | 0.722 |
| . | nonsynonym | ZFHX3:NM_0 | 0.136 | 1 | 0.044 | 0.411 |
| . | nonframeshif | RERE:NM_01 | 0.12 | 1 | NA | . |
| . | nonsynonym | ZFHX3:NM_0 | 0.136 | 1 | 0.003 | 0.682 |
| . | nonsynonym | NAV2:NM_18 | 0.247 | 1 | 0.002 | 0.722 |
| . | nonsynonym | VCAN:NM_00 | 0.199 | 1 | 0.003 | 0.682 |
| . | nonframeshif | RERE:NM_01 | 0.12 | 1 | NA | . |
| . | nonsynonym | VCAN:NM_00 | 0.199 | 1 | 0 | 0.913 |
| . | nonsynonym | RERE:NM_01 | 0.12 | 1 | 0.001 | 0.913 |
| . | nonsynonym | KDM5A:NM_ | 0.163 | 1 | 0.002 | 0.722 |
| . | nonsynonym | TULP4:NM_0 | 0.133 | 1 | 0 | 0.913 |
| . | nonframeshif | BCL11B:NM_ | 0.282 | 0.989 | NA | . |
| . | nonsynonym | NAV2:NM_00 | 0.247 | 1 | 0.004 | 0.682 |
| . | nonsynonym | ZFHX3:NM_0 | 0.136 | 1 | 0.019 | 0.501 |
| . | nonsynonym | TULP4:NM_0 | 0.133 | 1 | 0 | 0.913 |
| . | nonsynonym | ZFHX3:NM_0 | 0.136 | 1 | 0.001 | 0.785 |
| . | nonframeshif | ZFHX3:NM_0 | 0.136 | 1 | NA | . |
| . | nonsynonym | TULP4:NM_0 | 0.133 | 1 | 0.001 | 0.785 |
| . | nonframeshif | ZFHX3:NM_0 | 0.136 | 1 | NA | . |
| . | nonsynonym | BMP4:NM_00 | 0.334 | 0.956 | 0.004 | 0.682 |
| . | nonsynonym | ARID5B:NM_ | 0.11 | 1 | 0.001 | 0.785 |
| . | stopgain | TP63:NM_00 | 0.267 | 0.997 | NA | . |
| . | nonframeshif | RERE:NM_00 | 0.12 | 1 | NA | . |
| . | nonsynonym | DSP:NM_001 | 0.26 | 1 | 0.027 | 0.465 |
| . | nonsynonym | RERE:NM_00 | 0.12 | 1 | 0.001 | 0.785 |
| . | nonsynonym | DSP:NM_001 | 0.26 | 1 | 0.003 | 0.682 |
| . | nonsynonym | ZFHX3:NM_0 | 0.136 | 1 | 0 | 0.913 |
| . | nonsynonym | PIK3R1:NM_1 | 0.228 | 1 | 0.004 | 0.654 |
| . | nonframeshif | CALD1:NM_0 | 0.258 | 0.999 | NA | . |
| . | nonsynonym | VCAN:NM_00 | 0.199 | 1 | 0.006 | 0.614 |
| . | nonsynonym | DSP:NM_001 | 0.26 | 1 | 0.003 | 0.682 |
| . | nonsynonym | EEF1A1:NM_ | 0.293 | 0.979 | 0.001 | 0.785 |
| . | nonsynonym | NAV2:NM_00 | 0.247 | 1 | 0.003 | 0.722 |
| . | nonsynonym | CALD1:NM_0 | 0.258 | 0.999 | 0 | 0.913 |
| . | nonsynonym | ZFHX3:NM_0 | 0.136 | 1 | 0.003 | 0.682 |
| . | nonsynonym | TULP4:NM_0 | 0.133 | 1 | 0.006 | 0.614 |
| NM_001042 | nonsynonym | RERE:NM_01 | 0.12 | 1 | 0.001 | 0.785 |

| | | | | | | |
|---|---|---|---|---|---|---|
| . | nonframeshif | ZFHX3:NM_0( | 0.136 | 1 | NA | . |
| . | nonsynonym( | BMP4:NM_0( | 0.334 | 0.956 | 0.004 | 0.682 |
| . | nonsynonym( | CHD3:NM_0( | 0.147 | 1 | 0.046 | 0.406 |
| NM_0011149 | nonsynonym( | TP63:NM_00 | 0.267 | 0.997 | 0.001 | 0.785 |
| . | nonsynonym( | DSP:NM_001 | 0.26 | 1 | 0.001 | 0.785 |
| . | nonsynonym( | ZFHX3:NM_0( | 0.136 | 1 | 0.019 | 0.501 |
| . | nonsynonym( | BMP4:NM_0( | 0.334 | 0.956 | 0 | 0.913 |
| . | nonframeshif | ZFHX3:NM_0( | 0.136 | 1 | NA | . |
| . | nonsynonym( | RERE:NM_00 | 0.12 | 1 | 0.03 | 0.454 |
| . | nonsynonym( | TP63:NM_00 | 0.267 | 0.997 | 0.008 | 0.614 |
| . | nonsynonym( | ZFHX3:NM_0( | 0.136 | 1 | 0.032 | 0.45 |
| . | nonsynonym( | VCAN:NM_0( | 0.199 | 1 | 0.003 | 0.682 |
| . | nonsynonym( | CHD3:NM_0( | 0.147 | 1 | 0.013 | 0.539 |
| dist=593 | nonsynonym( | RPL3:NM_00( | 0.239 | 0.994 | 0.045 | 0.422 |
| . | nonsynonym( | DSP:NM_001 | 0.26 | 1 | 0.002 | 0.722 |
| . | nonsynonym( | ZFHX3:NM_0( | 0.136 | 1 | 0.005 | 0.654 |
| . | nonsynonym( | CALD1:NM_0 | 0.258 | 0.999 | 0 | 0.913 |
| . | nonsynonym( | TULP4:NM_0 | 0.133 | 1 | 0 | 0.913 |
| . | nonsynonym( | ARID5B:NM_( | 0.11 | 1 | 0.003 | 0.682 |
| . | nonsynonym( | CALD1:NM_0 | 0.258 | 0.999 | 0 | 0.913 |
| . | nonsynonym( | ZFHX3:NM_0( | 0.136 | 1 | 0.003 | 0.682 |
| . | nonsynonym( | DSP:NM_001 | 0.26 | 1 | 0.014 | 0.532 |
| . | nonsynonym( | ZFHX3:NM_0( | 0.136 | 1 | 0.001 | 0.785 |
| . | nonsynonym( | CALD1:NM_0 | 0.258 | 0.999 | 0 | 0.913 |
| . | nonsynonym( | RERE:NM_00 | 0.12 | 1 | 0.001 | 0.785 |
| . | nonframeshif | VCAN:NM_0( | 0.199 | 1 | NA | . |
| . | nonsynonym( | TULP4:NM_0 | 0.133 | 1 | 0 | 0.913 |
| . | nonsynonym( | RERE:NM_00 | 0.12 | 1 | 0.005 | 0.632 |
| . | nonsynonym( | TULP4:NM_0 | 0.133 | 1 | 0 | 0.913 |
| . | nonsynonym( | ZFHX3:NM_0( | 0.136 | 1 | 0.008 | 0.586 |
| . | nonsynonym( | RERE:NM_00 | 0.12 | 1 | 0 | 0.913 |
| . | nonsynonym( | ZFHX3:NM_0( | 0.136 | 1 | 0 | 0.913 |
| . | nonsynonym( | BMP4:NM_0( | 0.334 | 0.956 | 0 | 0.913 |
| . | nonsynonym( | CALD1:NM_0 | 0.258 | 0.999 | 0 | 0.913 |
| . | nonsynonym( | RERE:NM_00 | 0.12 | 1 | 0.005 | 0.632 |
| . | nonsynonym( | NAV2:NM_0( | 0.247 | 1 | 0.003 | 0.722 |
| . | nonsynonym( | IGFBP5:NM_( | 0.292 | 0.963 | 0.048 | 0.401 |
| . | nonframeshif | RERE:NM_01 | 0.12 | 1 | NA | . |
| . | nonsynonym( | DSP:NM_001 | 0.26 | 1 | 0.015 | 0.525 |
| . | nonsynonym( | EPHA4:NM_0 | 0.175 | 1 | 0 | 0.913 |
| . | nonsynonym( | BMP4:NM_0( | 0.334 | 0.956 | 0.05 | 0.403 |
| NM_0011149 | nonsynonym( | TP63:NM_00 | 0.267 | 0.997 | 0.001 | 0.785 |
| . | nonsynonym( | CALD1:NM_0 | 0.258 | 0.999 | 0 | 0.913 |

| | | | | | | |
|---|---|---|---|---|---|---|
| . | nonsynonymo | ARID5B:NM_( | 0.11 | 1 | 0.002 | 0.785 |
| . | nonsynonymo | RERE:NM_00 | 0.12 | 1 | 0.005 | 0.632 |
| . | nonsynonymo | RERE:NM_00 | 0.12 | 1 | 0.031 | 0.913 |

| SIFT_pred | SIFT4G_score | SIFT4G_conve | SIFT4G_pred | Polyphen2_H | Polyphen2_H | Polyphen2_H |
|---|---|---|---|---|---|---|
| . | . | . | . | . | . | . |
| . | . | . | . | . | . | . |
| D | 0.156 | 0.928 | T | 0.998 | 0.732 | D |
| D | 0.037 | 0.517 | D | 0.999 | 0.779 | D |
| D | 0.025 | 0.562 | D | 0.986 | 0.615 | D |
| D | 0.001 | 0.834 | D | 1 | 0.906 | D |
| D | 0.002 | 0.794 | D | 1 | 0.906 | D |
| D | 0.174 | 0.597 | T | 1 | 0.906 | D |
| D | 0 | 0.928 | D | 1 | 0.906 | D |
| D | 0.184 | 0.297 | T | 0.018 | 0.178 | B |
| D | 0.088 | 0.423 | T | 0.989 | 0.628 | D |
| D | 0.156 | 0.928 | T | 0.998 | 0.732 | D |
| D | 0.098 | 0.445 | T | 0.917 | 0.507 | P |
| D | 0.001 | 0.834 | D | 1 | 0.906 | D |
| . | . | . | . | . | . | . |
| D | 0.078 | 0.423 | T | 0.041 | 0.214 | B |
| D | 0.161 | 0.35 | T | 0.241 | 0.309 | B |
| D | 0.561 | 0.091 | T | 0.944 | 0.532 | P |
| D | 0.088 | 0.423 | T | 0.989 | 0.628 | D |
| D | 0.018 | 0.597 | D | 0.816 | 0.553 | P |
| D | 0.042 | 0.562 | D | 1 | 0.906 | D |
| . | . | . | . | . | . | . |
| D | 0.007 | 0.692 | D | 0.993 | 0.656 | D |
| D | 0.005 | 0.742 | D | 0.813 | 0.467 | P |
| . | . | . | . | . | . | . |
| D | 0.021 | 0.794 | D | 1 | 0.906 | D |
| D | 0.007 | 0.692 | D | 0.879 | 0.483 | P |
| D | 0.077 | 0.424 | T | 0.098 | 0.256 | B |
| D | 0.068 | 0.441 | T | 0.267 | 0.317 | B |
| D | 0.102 | 0.385 | T | 0.999 | 0.779 | D |
| D | 0.088 | 0.423 | T | 0.989 | 0.628 | D |
| D | 0.037 | 0.517 | D | 1 | 0.906 | D |
| D | 0.215 | 0.928 | T | 1 | 0.906 | D |
| D | 0.184 | 0.297 | T | 0.018 | 0.178 | B |
| D | 0.027 | 0.553 | D | 0.935 | 0.523 | P |
| D | 0.018 | 0.597 | D | 0.465 | 0.364 | P |
| D | 0.099 | 0.624 | T | 0.999 | 0.906 | D |
| D | 0.005 | 0.722 | D | 0.977 | 0.585 | D |
| D | 0.024 | 0.566 | D | 0.999 | 0.779 | D |
| D | 0.019 | 0.668 | D | 0.995 | 0.707 | D |
| D | 0.075 | 0.428 | T | 0.985 | 0.732 | D |
| D | 0.145 | 0.331 | T | 0.979 | 0.59 | D |

| | | | | | | |
|---|---|---|---|---|---|---|
| D | 0.156 | 0.928 | T | 0.998 | 0.732 | D |
| D | 0.069 | 0.439 | T | 0.255 | 0.341 | B |
| D | 0.005 | 0.722 | D | 0.993 | 0.656 | D |
| D | 0.023 | 0.581 | D | 0.487 | 0.369 | P |
| D | 0.042 | 0.562 | D | 1 | 0.906 | D |
| . | . | . | . | . | . | . |
| . | . | . | . | . | . | . |
| D | 0.57 | 0.226 | T | 0.007 | 0.267 | B |
| D | 0.036 | 0.722 | D | 1 | 0.906 | D |
| D | 0.029 | 0.545 | D | 0.999 | 0.779 | D |
| D | 0.057 | 0.464 | T | 0.936 | 0.524 | P |
| D | 0.101 | 0.386 | T | 0.117 | 0.265 | B |
| D | 0.127 | 0.562 | T | 0.996 | 0.779 | D |
| . | . | . | . | . | . | . |
| D | 0.093 | 0.398 | T | 0.995 | 0.675 | D |
| D | 0.029 | 0.679 | D | 0.977 | 0.906 | D |
| D | 0 | 0.928 | D | 1 | 0.906 | D |
| D | 0.764 | 0.046 | T | 0.937 | 0.524 | P |
| D | 0.151 | 0.324 | T | 0.998 | 0.732 | D |
| D | 0.129 | 0.348 | T | 0.962 | 0.556 | D |
| D | 0 | 0.928 | D | 0.999 | 0.779 | D |
| . | . | . | . | . | . | . |
| D | 0.156 | 0.928 | T | 0.998 | 0.732 | D |
| D | 0.035 | 0.524 | D | 0.23 | 0.306 | B |
| D | 0.058 | 0.462 | T | 0.895 | 0.492 | P |
| D | 0.088 | 0.423 | T | 0.989 | 0.628 | D |
| D | 0.507 | 0.109 | T | 0.993 | 0.656 | D |
| D | 0.489 | 0.116 | T | 0.543 | 0.379 | P |
| D | 0.017 | 0.648 | D | 1 | 0.906 | D |
| . | . | . | . | . | . | . |
| D | 0.002 | 0.928 | D | 1 | 0.906 | D |
| D | 0.097 | 0.392 | T | 0.014 | 0.169 | B |
| D | 0.43 | 0.21 | T | 0.626 | 0.401 | P |
| D | 0 | 0.928 | D | 1 | 0.906 | D |
| D | 0.156 | 0.928 | T | 0.998 | 0.732 | D |
| D | 0.156 | 0.928 | T | 0.998 | 0.732 | D |
| D | 0.173 | 0.542 | T | 0.997 | 0.707 | D |
| D | 0.009 | 0.706 | D | 0.946 | 0.779 | P |
| D | 0.002 | 0.794 | D | 0.906 | 0.5 | P |
| D | 0.042 | 0.502 | D | 1 | 0.906 | D |
| D | 0.38 | 0.282 | T | 0.816 | 0.456 | P |
| D | 0.078 | 0.423 | T | 0.041 | 0.214 | B |
| D | 0.039 | 0.511 | D | 1 | 0.906 | D |

| | | | | | | |
|---|---|---|---|---|---|---|
| D | 0.104 | 0.383 | T | 1 | 0.906 | D |
| . | . | . | . | . | . | . |
| D | 0.069 | 0.439 | T | 0.994 | 0.665 | D |
| D | 0.147 | 0.592 | T | 0.999 | 0.779 | D |
| D | 0.036 | 0.722 | D | 1 | 0.906 | D |
| D | 0.018 | 0.597 | D | 0.999 | 0.779 | D |
| D | 0.168 | 0.316 | T | 0.56 | 0.383 | P |
| D | 0.156 | 0.928 | T | 0.998 | 0.732 | D |
| D | 0.007 | 0.928 | D | 1 | 0.906 | D |
| D | 0.003 | 0.765 | D | 1 | 0.906 | D |
| . | . | . | . | . | . | . |
| D | 0.025 | 0.562 | D | 0.986 | 0.615 | D |
| D | 0.001 | 0.834 | D | 1 | 0.906 | D |
| . | . | . | . | . | . | . |
| . | . | . | . | . | . | . |
| D | 0 | 0.928 | D | 1 | 0.906 | D |
| . | . | . | . | . | . | . |
| D | 0.539 | 0.098 | T | 0.976 | 0.583 | D |
| D | 0.003 | 0.765 | D | 0.954 | 0.545 | P |
| . | . | . | . | . | . | . |
| D | 0.01 | 0.657 | D | 1 | 0.906 | D |
| D | 0.377 | 0.161 | T | 0.999 | 0.779 | D |
| . | . | . | . | . | . | . |
| D | 0.156 | 0.928 | T | 0.998 | 0.732 | D |
| D | 0.088 | 0.423 | T | 0.989 | 0.628 | D |
| D | 0.007 | 0.692 | D | 0.255 | 0.313 | B |
| D | 0.184 | 0.297 | T | 0.018 | 0.178 | B |
| . | . | . | . | . | . | . |
| D | 0.012 | 0.639 | D | 0.008 | 0.487 | B |
| D | 0.042 | 0.502 | D | 0.999 | 0.906 | D |
| D | 0.011 | 0.648 | D | 1 | 0.906 | D |
| D | 0.233 | 0.249 | T | 0.994 | 0.665 | D |
| D | 0.25 | 0.237 | T | 0.897 | 0.494 | P |
| D | 0.012 | 0.639 | D | 0.086 | 0.267 | B |
| D | 0.129 | 0.348 | T | 0.962 | 0.556 | D |
| D | 0.039 | 0.531 | D | 0.998 | 0.732 | D |
| D | 0.078 | 0.423 | T | 1 | 0.906 | D |
| D | 0.003 | 0.765 | D | 1 | 0.906 | D |
| D | 0.016 | 0.61 | D | 0.987 | 0.619 | D |
| D | 0.087 | 0.648 | T | 1 | 0.906 | D |
| . | . | . | . | . | . | . |
| . | . | . | . | . | . | . |
| D | 0.011 | 0.648 | D | 1 | 0.906 | D |

| | | | | | | |
|---|---|---|---|---|---|---|
| . | . | . | . | . | . | . |
| D | 0.156 | 0.928 | T | 0.998 | 0.732 | D |
| D | 0.129 | 0.348 | T | 0.962 | 0.556 | D |
| D | 0.039 | 0.511 | D | 0.012 | 0.163 | B |
| D | 0.156 | 0.928 | T | 0.504 | 0.372 | P |
| D | 0.002 | 0.834 | D | 1 | 0.906 | D |
| D | 0 | 0.928 | D | 0.993 | 0.656 | D |
| D | 0.056 | 0.466 | T | 1 | 0.906 | D |
| D | 0.002 | 0.794 | D | 1 | 0.906 | D |
| . | . | . | . | . | . | . |
| . | . | . | . | . | . | . |
| D | 0.002 | 0.794 | D | 1 | 0.906 | D |
| D | 0.654 | 0.199 | T | 0.062 | 0.233 | B |
| . | . | . | . | . | . | . |
| D | 0.114 | 0.369 | T | 0.33 | 0.332 | B |
| D | 0.141 | 0.334 | T | 0.999 | 0.779 | D |
| D | 0.042 | 0.566 | D | 0.912 | 0.535 | P |
| . | . | . | . | . | . | . |
| D | 0.184 | 0.297 | T | 0.018 | 0.178 | B |
| D | 0.12 | 0.36 | T | 0.386 | 0.375 | B |
| D | 0.007 | 0.706 | D | 0.999 | 0.779 | D |
| D | 0 | 0.928 | D | 0.993 | 0.656 | D |
| . | . | . | . | . | . | . |
| D | 0.008 | 0.679 | D | 0.93 | 0.518 | P |
| D | 0.001 | 0.834 | D | 0.885 | 0.517 | P |
| D | 0.026 | 0.558 | D | 0.057 | 0.229 | B |
| D | 0.017 | 0.603 | D | 0.712 | 0.421 | P |
| D | 0.071 | 0.435 | T | 0.953 | 0.574 | P |
| D | 0.012 | 0.639 | D | 1 | 0.906 | D |
| D | 0.002 | 0.794 | D | 1 | 0.906 | D |
| D | 0.017 | 0.648 | D | 1 | 0.906 | D |
| D | 0 | 0.928 | D | 0.993 | 0.656 | D |
| . | . | . | . | . | . | . |
| D | 0.009 | 0.692 | D | 1 | 0.906 | D |
| D | 0.055 | 0.469 | T | 0.741 | 0.43 | P |
| D | 0.002 | 0.794 | D | 1 | 0.906 | D |
| D | 0.074 | 0.43 | T | 1 | 0.906 | D |
| D | 0.015 | 0.616 | D | 1 | 0.906 | D |
| . | . | . | . | . | . | . |
| D | 0.017 | 0.648 | D | 1 | 0.906 | D |
| D | 0.004 | 0.742 | D | 0.991 | 0.641 | D |
| D | 0.016 | 0.61 | D | 0.977 | 0.585 | D |
| D | 0 | 0.928 | D | 0.997 | 0.707 | D |

| | | | | | | |
|---|---|---|---|---|---|---|
| D | 0.03 | 0.928 | D | 0.923 | 0.512 | P |
| D | 0.01 | 0.657 | D | 1 | 0.906 | D |
| . | . | . | . | . | . | . |
| . | . | . | . | . | . | . |
| D | 0.017 | 0.603 | D | 0.712 | 0.421 | P |
| D | 0.175 | 0.299 | T | 1 | 0.906 | D |
| D | 0.015 | 0.616 | D | 0.967 | 0.564 | D |
| D | 0.007 | 0.706 | D | 0.999 | 0.779 | D |
| D | 0.005 | 0.722 | D | 0.992 | 0.647 | D |
| . | . | . | . | . | . | . |
| D | 0.088 | 0.406 | T | 1 | 0.906 | D |
| D | 0.21 | 0.291 | T | 0.524 | 0.376 | P |
| D | 0 | 0.928 | D | 1 | 0.906 | D |
| D | 0 | 0.928 | D | 1 | 0.906 | D |
| . | . | . | . | . | . | . |
| . | . | . | . | . | . | . |
| D | 0 | 0.928 | D | 1 | 0.906 | D |
| D | 0.131 | 0.558 | T | 0.049 | 0.222 | B |
| D | 0.007 | 0.692 | D | 0.996 | 0.688 | D |
| . | . | . | . | . | . | . |
| D | 0.008 | 0.679 | D | 0.93 | 0.518 | P |
| D | 0.078 | 0.423 | T | 0.999 | 0.779 | D |
| D | 0.001 | 0.834 | D | 1 | 0.906 | D |
| D | 0.255 | 0.234 | T | 0.993 | 0.906 | D |
| D | 0.015 | 0.616 | D | 0.421 | 0.354 | B |
| D | 0.034 | 0.527 | D | 0.939 | 0.526 | P |
| . | . | . | . | . | . | . |
| D | 0.088 | 0.406 | T | 1 | 0.906 | D |
| D | 0.474 | 0.491 | T | 0.608 | 0.395 | P |
| D | 0.156 | 0.928 | T | 0.998 | 0.732 | D |
| D | 0.017 | 0.648 | D | 1 | 0.906 | D |
| D | 0.21 | 0.291 | T | 0.524 | 0.376 | P |
| D | 0.017 | 0.648 | D | 1 | 0.906 | D |
| D | 0.129 | 0.348 | T | 0.962 | 0.556 | D |
| D | 0.354 | 0.173 | T | 0.991 | 0.641 | D |
| D | 0.108 | 0.409 | T | 0.062 | 0.233 | B |
| D | 0.075 | 0.428 | T | 0.985 | 0.732 | D |
| D | 0.001 | 0.834 | D | 1 | 0.906 | D |
| D | 0.005 | 0.722 | D | 0.993 | 0.656 | D |
| D | 0.062 | 0.453 | T | 0.985 | 0.611 | D |
| . | . | . | . | . | . | . |
| D | 0.002 | 0.794 | D | 0.999 | 0.906 | D |
| D | 0.055 | 0.469 | T | 0.741 | 0.43 | P |

| | | | | | | |
|---|---|---|---|---|---|---|
| . | . | . | . | . | . | . |
| D | 0.017 | 0.648 | D | 1 | 0.906 | D |
| D | 0.001 | 0.834 | D | 1 | 0.906 | D |
| D | 0.055 | 0.469 | T | 0.741 | 0.43 | P |
| D | 0.017 | 0.603 | D | 0.712 | 0.421 | P |
| D | 0.055 | 0.469 | T | 0.741 | 0.43 | P |
| D | 0.187 | 0.288 | T | 0.112 | 0.263 | B |
| D | 0.071 | 0.435 | T | 0.997 | 0.707 | D |
| . | . | . | . | . | . | . |
| D | 0.074 | 0.43 | T | 1 | 0.906 | D |
| D | 0.075 | 0.428 | T | 0.985 | 0.732 | D |
| D | 0.009 | 0.668 | D | 0.976 | 0.615 | D |
| . | . | . | . | . | . | . |
| D | 0.158 | 0.316 | T | 1 | 0.906 | D |
| D | 0.009 | 0.692 | D | 1 | 0.906 | D |
| D | 0.031 | 0.597 | D | 0.69 | 0.416 | P |
| D | 0.005 | 0.722 | D | 1 | 0.906 | D |
| . | . | . | . | . | . | . |
| D | 0.011 | 0.657 | D | 0.961 | 0.554 | D |
| D | 0.174 | 0.3 | T | 1 | 0.906 | D |
| D | 0.005 | 0.722 | D | 1 | 0.906 | D |
| D | 0.002 | 0.794 | D | 0.98 | 0.594 | D |
| . | . | . | . | . | . | . |
| D | 0.004 | 0.742 | D | 0.999 | 0.779 | D |
| . | . | . | . | . | . | . |
| D | 0.014 | 0.624 | D | 1 | 0.906 | D |
| D | 0.023 | 0.657 | D | 0.877 | 0.482 | P |
| . | . | . | . | . | . | . |
| . | . | . | . | . | . | . |
| D | 0.63 | 0.072 | T | 0.276 | 0.319 | B |
| D | 0.012 | 0.668 | D | 0.999 | 0.906 | D |
| D | 0.053 | 0.497 | T | 0.993 | 0.656 | D |
| D | 0.001 | 0.834 | D | 0.995 | 0.675 | D |
| D | 0.007 | 0.692 | D | 0.96 | 0.553 | D |
| . | . | . | . | . | . | . |
| D | 0.064 | 0.449 | T | 0.999 | 0.779 | D |
| D | 0.053 | 0.497 | T | 0.993 | 0.656 | D |
| D | 0.003 | 0.765 | D | 1 | 0.906 | D |
| D | 0.096 | 0.398 | T | 1 | 0.906 | D |
| D | 0.065 | 0.447 | T | 0.003 | 0.112 | B |
| D | 0.005 | 0.722 | D | 1 | 0.906 | D |
| D | 0.002 | 0.794 | D | 0.999 | 0.779 | D |
| D | 0.016 | 0.794 | D | 1 | 0.906 | D |

| | | | | | | |
|---|---|---|---|---|---|---|
| . | . | . | . | . | . | . |
| D | 0.014 | 0.624 | D | 1 | 0.906 | D |
| D | 0.193 | 0.282 | T | 0.998 | 0.779 | D |
| D | 0.002 | 0.794 | D | 0.997 | 0.707 | D |
| D | 0.022 | 0.576 | D | 0.999 | 0.779 | D |
| D | 0.072 | 0.61 | T | 0.992 | 0.647 | D |
| D | 0.001 | 0.834 | D | 0.999 | 0.779 | D |
| . | . | . | . | . | . | . |
| D | 0.065 | 0.447 | T | 0.026 | 0.194 | B |
| D | 0.137 | 0.339 | T | 0.997 | 0.707 | D |
| D | 0.654 | 0.199 | T | 0.062 | 0.233 | B |
| D | 0.104 | 0.382 | T | 0.999 | 0.779 | D |
| D | 0 | 0.928 | D | 1 | 0.906 | D |
| D | 0.125 | 0.353 | T | 0.075 | 0.242 | B |
| D | 0.007 | 0.692 | D | 0.948 | 0.536 | P |
| D | 0.012 | 0.639 | D | 0.34 | 0.334 | B |
| D | 0.638 | 0.449 | T | 0.958 | 0.675 | D |
| D | 0.007 | 0.692 | D | 1 | 0.906 | D |
| D | 0.139 | 0.361 | T | 0.203 | 0.334 | B |
| D | 0.638 | 0.449 | T | 0.958 | 0.675 | D |
| D | 0.146 | 0.329 | T | 0.001 | 0.075 | B |
| D | 0.442 | 0.133 | T | 0.997 | 0.707 | D |
| D | 0.002 | 0.794 | D | 0.98 | 0.594 | D |
| D | 0.638 | 0.449 | T | 0.958 | 0.675 | D |
| D | 0.005 | 0.742 | D | 0.95 | 0.539 | P |
| . | . | . | . | . | . | . |
| D | 0.007 | 0.692 | D | 1 | 0.906 | D |
| D | 0.022 | 0.576 | D | 0.999 | 0.779 | D |
| D | 0.005 | 0.722 | D | 1 | 0.906 | D |
| D | 0.113 | 0.369 | T | 0.985 | 0.611 | D |
| D | 0.118 | 0.39 | T | 0.999 | 0.906 | D |
| D | 0 | 0.928 | D | 1 | 0.906 | D |
| D | 0.001 | 0.834 | D | 0.999 | 0.779 | D |
| D | 0.638 | 0.449 | T | 0.958 | 0.675 | D |
| D | 0.022 | 0.576 | D | 0.999 | 0.779 | D |
| D | 0.096 | 0.398 | T | 1 | 0.906 | D |
| D | 0.009 | 0.668 | D | 0.074 | 0.241 | B |
| . | . | . | . | . | . | . |
| D | 0.013 | 0.631 | D | 0.067 | 0.237 | B |
| D | 0.001 | 0.834 | D | 1 | 0.906 | D |
| D | 0.154 | 0.32 | T | 0.995 | 0.675 | D |
| D | 0.002 | 0.794 | D | 0.997 | 0.707 | D |
| D | 0.638 | 0.449 | T | 0.958 | 0.675 | D |

| | | | | | | |
|---|---|---|---|---|---|---|
| D | 0.051 | 0.558 | T | 0.089 | 0.536 | B |
| D | 0.022 | 0.576 | D | 0.999 | 0.779 | D |
| D | 0.085 | 0.473 | T | 0.997 | 0.779 | D |

| Polyphen2_H | Polyphen2_H | Polyphen2_H | LRT_score | LRT_converte | LRT_pred | MutationTast |
|---|---|---|---|---|---|---|
| . | . | . | . | . | . | . |
| . | . | . | . | . | . | . |
| 0.978 | 0.741 | D | 0 | 0.843 | D | 0.999 |
| 0.996 | 0.845 | D | 0 | 0.843 | D | 1 |
| 0.579 | 0.502 | P | 0 | 0.45 | D | 1 |
| 0.982 | 0.755 | D | . | . | . | 1 |
| 0.96 | 0.703 | D | 0.001 | 0.413 | D | 1 |
| 0.992 | 0.886 | D | 0 | 0.537 | D | 1 |
| 0.999 | 0.924 | D | 0 | 0.629 | D | 1 |
| 0.067 | 0.275 | B | 0 | 0.843 | D | 1 |
| 0.796 | 0.58 | P | 0 | 0.629 | D | 1 |
| 0.978 | 0.741 | D | 0 | 0.843 | D | 0.999 |
| 0.677 | 0.534 | P | . | . | . | 0.999 |
| 0.999 | 0.924 | D | 0 | 0.629 | D | 1 |
| . | . | . | . | . | . | . |
| 0.062 | 0.269 | B | 0 | 0.843 | U | 1 |
| 0.086 | 0.295 | B | 0.01 | 0.301 | N | 0.991 |
| 0.523 | 0.483 | P | 0 | 0.843 | D | 1 |
| 0.796 | 0.58 | P | 0 | 0.629 | D | 1 |
| 0.146 | 0.46 | B | . | . | . | 1 |
| 0.994 | 0.821 | D | 0 | 0.843 | D | 1 |
| . | . | . | . | . | . | . |
| 0.925 | 0.66 | D | 0 | 0.491 | D | 1 |
| 0.495 | 0.498 | P | 0 | 0.843 | D | 1 |
| . | . | . | 0 | 0.629 | D | 1 |
| 0.999 | 0.924 | D | 0 | 0.629 | D | 1 |
| 0.801 | 0.582 | P | 0 | 0.843 | D | 1 |
| 0.067 | 0.275 | B | 0.019 | 0.273 | U | 1 |
| 0.047 | 0.248 | B | 0.033 | 0.25 | N | 0.907 |
| 0.985 | 0.765 | D | 0 | 0.843 | D | 1 |
| 0.796 | 0.58 | P | 0 | 0.629 | D | 1 |
| 1 | 0.974 | D | 0 | 0.843 | D | 1 |
| 0.996 | 0.845 | D | 0 | 0.457 | D | 1 |
| 0.067 | 0.275 | B | 0 | 0.843 | D | 1 |
| 0.455 | 0.463 | P | 0 | 0.629 | D | 0.981 |
| 0.23 | 0.382 | B | 0.111 | 0.194 | N | 0.991 |
| 0.997 | 0.886 | D | . | . | . | 1 |
| 0.89 | 0.632 | P | 0 | 0.843 | D | 1 |
| 0.872 | 0.619 | P | 0 | 0.843 | D | 1 |
| 0.964 | 0.761 | D | 0 | 0.843 | D | 1 |
| 0.435 | 0.628 | B | 0 | 0.629 | D | 1 |
| 0.273 | 0.4 | B | 0 | 0.843 | D | 1 |

| | | | | | | |
|---|---|---|---|---|---|---|
| 0.978 | 0.741 | D | 0 | 0.843 | D | 0.999 |
| 0.053 | 0.257 | B | 0.882 | 0.087 | N | 1 |
| 0.72 | 0.549 | P | 0 | 0.629 | D | 1 |
| 0.093 | 0.301 | B | 0.001 | 0.421 | D | 0.99 |
| 0.994 | 0.821 | D | 0 | 0.843 | D | 1 |
| . | . | . | 0 | 0.486 | D | 1 |
| . | . | . | . | . | . | . |
| 0.005 | 0.211 | B | 0.036 | 0.245 | N | 1 |
| 0.995 | 0.832 | D | 0 | 0.843 | D | 1 |
| 0.912 | 0.648 | D | . | . | . | 1 |
| 0.767 | 0.567 | P | 0 | 0.629 | D | 1 |
| 0.112 | 0.316 | B | 0 | 0.843 | D | 1 |
| 0.877 | 0.703 | P | 0.031 | 0.252 | N | 0.999 |
| . | . | . | . | . | . | . |
| 0.829 | 0.596 | P | 0.026 | 0.259 | N | 0.991 |
| 0.46 | 0.863 | P | 0.022 | 0.267 | N | 1 |
| 0.996 | 0.845 | D | 0 | 0.629 | D | 1 |
| 0.628 | 0.517 | P | 0.743 | 0.098 | N | 0.961 |
| 0.873 | 0.676 | P | 0.035 | 0.247 | N | 1 |
| 0.173 | 0.355 | B | 0 | 0.491 | D | 1 |
| 0.991 | 0.797 | D | 0 | 0.559 | D | 1 |
| . | . | . | . | . | . | . |
| 0.978 | 0.741 | D | 0 | 0.843 | D | 0.999 |
| 0.032 | 0.221 | B | 0 | 0.843 | D | 1 |
| 0.351 | 0.428 | B | 0 | 0.629 | D | 0.89 |
| 0.796 | 0.58 | P | 0 | 0.629 | D | 1 |
| 0.971 | 0.724 | D | 0 | 0.513 | D | 0.999 |
| 0.121 | 0.323 | B | 0 | 0.843 | D | 1 |
| 0.998 | 0.924 | D | 0 | 0.537 | D | 0.999 |
| . | . | . | . | . | . | . |
| 0.997 | 0.863 | D | 0.003 | 0.361 | N | 0.987 |
| 0.007 | 0.13 | B | 0.009 | 0.307 | U | 0.992 |
| 0.162 | 0.348 | B | 0 | 0.559 | D | 0.997 |
| 0.999 | 0.924 | D | 0 | 0.629 | D | 1 |
| 0.978 | 0.741 | D | 0 | 0.843 | D | 0.999 |
| 0.978 | 0.741 | D | 0 | 0.843 | D | 0.999 |
| 0.707 | 0.544 | P | 0 | 0.629 | D | 1 |
| 0.462 | 0.697 | P | 0 | 0.629 | D | 1 |
| 0.756 | 0.563 | P | 0 | 0.843 | D | 0.994 |
| 0.991 | 0.797 | D | 0 | 0.537 | D | 0.977 |
| 0.432 | 0.454 | B | 0 | 0.629 | D | 0.999 |
| 0.062 | 0.269 | B | 0 | 0.843 | U | 1 |
| 0.873 | 0.62 | P | 0.001 | 0.425 | U | 1 |

| | | | | | | | | | |
|---|---|---|---|---|---|---|---|---|---|
| | 0.995 | 0.832 | D | | 0 | 0.629 | D | | 1 |
| . | . | . | | . | . | . | | . | |
| | 0.892 | 0.633 | P | | 0 | 0.481 | D | | 0.965 |
| | 0.974 | 0.732 | D | | 0 | 0.843 | D | | 1 |
| | 0.995 | 0.832 | D | | 0 | 0.843 | D | | 1 |
| | 0.997 | 0.863 | D | | 0 | 0.843 | D | | 1 |
| | 0.56 | 0.496 | P | . | . | . | | | 1 |
| | 0.978 | 0.741 | D | | 0 | 0.843 | D | | 0.999 |
| | 0.996 | 0.886 | D | | 0.001 | 0.411 | D | | 0.936 |
| | 0.998 | 0.886 | D | | 0 | 0.843 | D | | 1 |
| . | . | . | | . | . | . | | . | |
| | 0.579 | 0.502 | P | | 0 | 0.45 | D | | 1 |
| | 0.994 | 0.821 | D | | 0 | 0.843 | D | | 1 |
| . | . | . | | . | . | . | | . | |
| . | . | . | | . | . | . | | . | |
| | 0.999 | 0.924 | D | | 0 | 0.629 | D | | 1 |
| . | . | . | | . | . | . | | . | |
| | 0.696 | 0.54 | P | | 0 | 0.497 | D | | 1 |
| | 0.276 | 0.401 | B | | 0 | 0.445 | D | | 0.998 |
| . | . | . | | . | . | . | | . | |
| | 0.95 | 0.688 | D | | 0 | 0.629 | D | | 1 |
| | 0.997 | 0.863 | D | | 0 | 0.629 | D | | 1 |
| . | . | . | | . | . | . | | . | |
| | 0.978 | 0.741 | D | | 0 | 0.843 | D | | 0.999 |
| | 0.796 | 0.58 | P | | 0 | 0.629 | D | | 1 |
| | 0.057 | 0.263 | B | | 0 | 0.497 | D | | 1 |
| | 0.067 | 0.275 | B | | 0 | 0.843 | D | | 1 |
| . | . | . | | . | . | . | | . | |
| | 0.038 | 0.537 | B | | 0.074 | 0.213 | N | | 0.999 |
| | 0.949 | 0.738 | D | | 0 | 0.523 | D | | 1 |
| | 0.999 | 0.924 | D | | 0 | 0.843 | D | | 0.993 |
| | 0.919 | 0.655 | D | | 0 | 0.843 | D | | 1 |
| | 0.229 | 0.382 | B | | 0.001 | 0.434 | D | | 0.999 |
| | 0.003 | 0.181 | B | | 0 | 0.629 | D | | 1 |
| | 0.173 | 0.355 | B | | 0 | 0.491 | D | | 1 |
| | 0.991 | 0.797 | D | | 0 | 0.497 | D | | 1 |
| | 0.999 | 0.924 | D | | 0 | 0.629 | D | | 1 |
| | 0.992 | 0.804 | D | | 0 | 0.843 | D | | 1 |
| | 0.823 | 0.592 | P | | 0 | 0.491 | D | | 1 |
| | 0.999 | 0.924 | D | | 0 | 0.843 | D | | 1 |
| . | . | . | | . | . | . | | . | |
| . | . | . | | . | . | . | | . | |
| | 0.999 | 0.924 | D | | 0 | 0.843 | D | | 0.993 |

| | | | | | | |
|---|---|---|---|---|---|---|
| . | . | . | . | . | . | . |
| 0.978 | 0.741 | D | 0 | 0.843 | D | 0.999 |
| 0.173 | 0.355 | B | 0 | 0.491 | D | 1 |
| 0.016 | 0.177 | B | 0 | 0.843 | D | 1 |
| 0.338 | 0.423 | B | 0 | 0.486 | D | 1 |
| 0.981 | 0.752 | D | 0 | 0.559 | D | 0.996 |
| 0.92 | 0.655 | D | 0 | 0.629 | D | 1 |
| 0.899 | 0.638 | P | 0 | 0.843 | D | 1 |
| 0.998 | 0.886 | D | 0 | 0.505 | D | 1 |
| . | . | . | . | . | . | 1 |
| . | . | . | 0 | 0.843 | D | 1 |
| 0.999 | 0.924 | D | 0 | 0.629 | D | 1 |
| 0.036 | 0.229 | B | 0 | 0.505 | D | 1 |
| . | . | . | . | . | . | . |
| 0.046 | 0.247 | B | 0.04 | 0.241 | N | 1 |
| 0.994 | 0.821 | D | 0 | 0.843 | D | 0.846 |
| 0.889 | 0.685 | P | 0 | 0.843 | D | 1 |
| . | . | . | . | . | . | . |
| 0.067 | 0.275 | B | 0 | 0.843 | D | 1 |
| 0.075 | 0.346 | B | 0.098 | 0.2 | N | 0.776 |
| 0.921 | 0.656 | D | 0 | 0.629 | D | 1 |
| 0.92 | 0.655 | D | 0 | 0.629 | D | 1 |
| . | . | . | . | . | . | . |
| 0.564 | 0.497 | P | 0 | 0.513 | D | 0.999 |
| 0.367 | 0.591 | B | 0 | 0.843 | D | 1 |
| 0.151 | 0.342 | B | . | . | . | 0.691 |
| 0.138 | 0.334 | B | 0.06 | 0.223 | N | 0.81 |
| 0.315 | 0.524 | B | 0.049 | 0.232 | N | 0.599 |
| 0.973 | 0.729 | D | 0 | 0.629 | D | 0.668 |
| 0.994 | 0.821 | D | 0 | 0.497 | D | 1 |
| 0.998 | 0.924 | D | 0 | 0.537 | D | 0.999 |
| 0.92 | 0.655 | D | 0 | 0.629 | D | 1 |
| . | . | . | . | . | . | . |
| 0.998 | 0.924 | D | . | . | . | 1 |
| 0.491 | 0.474 | P | 0 | 0.629 | D | 1 |
| 0.971 | 0.724 | D | 0 | 0.629 | D | 1 |
| 0.999 | 0.924 | D | 0 | 0.454 | D | 1 |
| 0.999 | 0.924 | D | 0 | 0.629 | D | 1 |
| . | . | . | . | . | . | . |
| 0.998 | 0.924 | D | 0 | 0.537 | D | 0.999 |
| 0.991 | 0.797 | D | 0 | 0.843 | D | 1 |
| 0.787 | 0.576 | P | 0 | 0.523 | D | 0.998 |
| 0.995 | 0.832 | D | 0 | 0.559 | D | 1 |

| | | | | | | |
|---|---|---|---|---|---|---|
| 0.729 | 0.552 | P | 0 | 0.843 | D | 1 |
| 0.999 | 0.924 | D | 0 | 0.629 | D | 1 |
| . | . | . | . | . | . | . |
| . | . | . | . | . | . | . |
| 0.138 | 0.334 | B | 0.06 | 0.223 | N | 0.81 |
| 0.999 | 0.924 | D | . | . | . | 0.998 |
| 0.916 | 0.652 | D | 0 | 0.629 | D | 1 |
| 0.921 | 0.656 | D | 0 | 0.629 | D | 1 |
| 0.629 | 0.517 | P | 0 | 0.523 | D | 1 |
| . | . | . | . | . | . | . |
| 1 | 0.974 | D | 0 | 0.629 | D | 0.999 |
| 0.017 | 0.181 | B | 0 | 0.497 | D | 1 |
| 1 | 0.974 | D | 0 | 0.843 | U | 1 |
| 1 | 0.974 | D | 0 | 0.843 | U | 1 |
| . | . | . | . | . | . | . |
| . | . | . | . | . | . | . |
| 0.985 | 0.765 | D | 0 | 0.843 | D | 1 |
| 0.013 | 0.193 | B | 0 | 0.559 | D | 1 |
| 0.836 | 0.599 | P | 0 | 0.843 | D | 1 |
| . | . | . | . | . | . | . |
| 0.564 | 0.497 | P | 0 | 0.513 | D | 0.999 |
| 0.995 | 0.832 | D | 0 | 0.513 | D | 1 |
| 0.999 | 0.924 | D | 0 | 0.629 | D | 1 |
| 0.701 | 0.924 | P | 0 | 0.629 | D | 1 |
| 0.073 | 0.281 | B | 0 | 0.629 | D | 1 |
| 0.746 | 0.559 | P | 0 | 0.45 | D | 1 |
| . | . | . | . | . | . | . |
| 1 | 0.974 | D | 0 | 0.629 | D | 0.999 |
| 0.046 | 0.247 | B | 0 | 0.497 | D | 1 |
| 0.978 | 0.741 | D | 0 | 0.843 | D | 0.999 |
| 0.998 | 0.924 | D | 0 | 0.537 | D | 0.999 |
| 0.017 | 0.181 | B | 0 | 0.497 | D | 1 |
| 0.998 | 0.924 | D | 0 | 0.537 | D | 0.999 |
| 0.173 | 0.355 | B | 0 | 0.491 | D | 1 |
| 0.801 | 0.582 | P | 0 | 0.537 | D | 1 |
| 0.031 | 0.219 | B | 0.01 | 0.301 | N | 0.999 |
| 0.435 | 0.628 | B | 0 | 0.629 | D | 1 |
| 0.999 | 0.924 | D | 0 | 0.629 | D | 1 |
| 0.72 | 0.549 | P | 0 | 0.629 | D | 1 |
| 0.338 | 0.423 | B | 0 | 0.843 | D | 1 |
| . | . | . | . | . | . | . |
| 0.993 | 0.863 | D | 0 | 0.843 | D | 1 |
| 0.491 | 0.474 | P | 0 | 0.629 | D | 1 |

| | | | | | | |
|---|---|---|---|---|---|---|
| . | . | . | . | . | . | . |
| 0.998 | 0.924 | D | 0 | 0.537 | D | 0.999 |
| 0.999 | 0.924 | D | 0 | 0.629 | D | 1 |
| 0.491 | 0.474 | P | 0 | 0.629 | D | 1 |
| 0.138 | 0.334 | B | 0.06 | 0.223 | N | 0.81 |
| 0.491 | 0.474 | P | 0 | 0.629 | D | 1 |
| 0.047 | 0.248 | B | 0.001 | 0.39 | N | 1 |
| 0.991 | 0.797 | D | 0.001 | 0.432 | D | 1 |
| . | . | . | . | . | . | . |
| 0.999 | 0.924 | D | 0 | 0.454 | D | 1 |
| 0.435 | 0.628 | B | 0 | 0.629 | D | 1 |
| 0.601 | 0.508 | P | 0.056 | 0.226 | N | 1 |
| . | . | . | . | . | . | . |
| 0.994 | 0.821 | D | 0.047 | 0.234 | N | 0.894 |
| 0.998 | 0.924 | D | . | . | . | 1 |
| 0.372 | 0.436 | B | 0 | 0.843 | D | 1 |
| 0.998 | 0.886 | D | 0 | 0.843 | D | 1 |
| . | . | . | . | . | . | . |
| 0.996 | 0.845 | D | 0 | 0.629 | D | 1 |
| 0.987 | 0.775 | D | 0.001 | 0.438 | D | 1 |
| 0.998 | 0.886 | D | 0 | 0.843 | D | 1 |
| 0.669 | 0.531 | P | 0 | 0.497 | D | 1 |
| . | . | . | . | . | . | . |
| 0.887 | 0.667 | P | 0.001 | 0.431 | D | 1 |
| . | . | . | . | . | . | . |
| 0.993 | 0.811 | D | 0 | 0.843 | D | 1 |
| 0.417 | 0.45 | B | 0.157 | 0.178 | N | 0.998 |
| . | . | . | 0.004 | 0.34 | N | 1 |
| . | . | . | . | . | . | . |
| 0.066 | 0.274 | B | 0 | 0.537 | D | 0.997 |
| 0.984 | 0.761 | D | . | . | . | 1 |
| 0.982 | 0.755 | D | 0 | 0.629 | D | 0.998 |
| 0.979 | 0.745 | D | 0 | 0.477 | D | 1 |
| 0.802 | 0.583 | P | 0 | 0.843 | D | 1 |
| . | . | . | . | . | . | . |
| 0.78 | 0.573 | P | 0.003 | 0.36 | N | 0.601 |
| 0.982 | 0.755 | D | 0 | 0.629 | D | 0.998 |
| 1 | 0.974 | D | 0 | 0.629 | U | 1 |
| 0.993 | 0.811 | D | 0 | 0.559 | D | 1 |
| 0.012 | 0.16 | B | 0.063 | 0.22 | N | 1 |
| 1 | 0.974 | D | 0 | 0.497 | D | 1 |
| 0.957 | 0.699 | D | 0 | 0.629 | D | 1 |
| 0.996 | 0.863 | D | . | . | . | 1 |

| | | | | | | |
|---|---|---|---|---|---|---|
| . | . | . | . | . | . | . |
| 0.993 | 0.811 | D | 0 | 0.843 | D | 1 |
| 0.856 | 0.61 | P | 0 | 0.469 | D | 1 |
| 0.981 | 0.752 | D | 0 | 0.843 | D | 1 |
| 0.799 | 0.581 | P | 0 | 0.629 | D | 1 |
| 0.963 | 0.708 | D | 0 | 0.537 | D | 1 |
| 0.985 | 0.765 | D | 0 | 0.843 | D | 1 |
| . | . | . | . | . | . | . |
| 0.017 | 0.181 | B | . | . | . | 0.574 |
| 0.986 | 0.769 | D | . | . | . | 1 |
| 0.036 | 0.229 | B | 0 | 0.505 | D | 1 |
| 0.957 | 0.699 | D | 0 | 0.537 | D | 1 |
| 1 | 0.974 | D | 0 | 0.466 | D | 1 |
| 0.009 | 0.143 | B | 0 | 0.843 | D | 1 |
| 0.666 | 0.53 | P | 0 | 0.629 | D | 0.936 |
| 0.172 | 0.354 | B | 0.024 | 0.264 | N | 0.965 |
| 0.827 | 0.633 | P | 0 | 0.481 | D | 0.637 |
| 0.997 | 0.863 | D | 0 | 0.843 | D | 1 |
| 0.055 | 0.331 | B | 0.286 | 0.148 | N | 0.993 |
| 0.827 | 0.633 | P | 0 | 0.481 | D | 0.637 |
| 0 | 0.014 | B | 0.012 | 0.292 | N | 0.724 |
| 0.968 | 0.717 | D | 0 | 0.559 | D | 0.993 |
| 0.669 | 0.531 | P | 0 | 0.497 | D | 1 |
| 0.827 | 0.633 | P | 0 | 0.481 | D | 0.637 |
| 0.314 | 0.447 | B | . | . | . | 0.984 |
| . | . | . | . | . | . | . |
| 0.997 | 0.863 | D | 0 | 0.843 | D | 1 |
| 0.941 | 0.677 | D | . | . | . | 1 |
| 0.998 | 0.886 | D | 0 | 0.843 | D | 1 |
| 0.709 | 0.545 | P | 0.001 | 0.434 | D | 0.982 |
| 0.954 | 0.804 | D | . | . | . | 1 |
| 0.997 | 0.863 | D | 0.014 | 0.287 | N | 1 |
| 0.985 | 0.765 | D | 0 | 0.843 | D | 1 |
| 0.827 | 0.633 | P | 0 | 0.481 | D | 0.637 |
| 0.941 | 0.677 | D | . | . | . | 1 |
| 0.993 | 0.811 | D | 0 | 0.559 | D | 1 |
| 0.053 | 0.257 | B | 0 | 0.843 | D | 1 |
| . | . | . | . | . | . | . |
| 0.011 | 0.155 | B | 0 | 0.629 | D | 0.996 |
| 0.999 | 0.924 | D | 0 | 0.843 | D | 1 |
| 0.937 | 0.673 | D | 0 | 0.843 | D | 1 |
| 0.981 | 0.752 | D | 0 | 0.843 | D | 1 |
| 0.827 | 0.633 | P | 0 | 0.481 | D | 0.637 |

| | | | | | | | |
|---|---|---|---|---|---|---|---|
| 0.023 | 0.444 | B | | 0.006 | 0.325 | N | 1 |
| 0.941 | 0.677 | D | . | . | . | | 1 |
| 0.992 | 0.832 | D | . | . | . | | 1 |

| MutationTast | MutationTast | MutationAsse | MutationAsse | MutationAsse | FATHMM_sco | FATHMM_cor |
|---|---|---|---|---|---|---|
| . | . | . | . | . | . | . |
| . | . | . | . | . | . | . |
| 0.464 | D | 2.405 | 0.696 | M | 3.18 | 0.599 |
| 0.81 | D | 2.765 | 0.808 | M | 3.08 | 0.085 |
| 0.508 | D | 2.14 | 0.599 | M | -2.77 | 0.909 |
| 0.482 | D | 0.345 | 0.112 | N | 0.59 | 0.539 |
| 0.479 | D | 2.235 | 0.632 | M | -3.81 | 0.957 |
| 0.51 | D | 1.54 | 0.389 | L | 0.42 | 0.569 |
| 0.81 | D | 3.12 | 0.879 | M | -2.47 | 0.89 |
| 0.81 | D | 1.685 | 0.433 | L | . | . |
| 0.588 | D | 2.35 | 0.675 | M | 2.41 | 0.158 |
| 0.464 | D | 2.405 | 0.696 | M | 3.18 | 0.599 |
| 0.81 | D | 1.645 | 0.42 | L | -0.7 | 0.728 |
| 0.81 | D | 2.98 | 0.855 | M | 0.54 | 0.549 |
| . | . | . | . | . | . | . |
| 0.508 | D | 2.32 | 0.664 | M | -1.05 | 0.767 |
| 0.419 | D | 0.55 | 0.145 | N | -2.7 | 0.905 |
| 0.81 | D | 3.7 | 0.946 | H | 1.1 | 0.391 |
| 0.588 | D | 2.35 | 0.675 | M | 2.41 | 0.158 |
| 0.81 | D | 2.005 | 0.546 | M | 0.76 | 0.52 |
| 0.81 | D | 2.085 | 0.577 | M | -1.99 | 0.853 |
| . | . | . | . | . | . | . |
| 0.81 | D | 2.33 | 0.668 | M | -0.91 | 0.75 |
| 0.81 | D | 1.935 | 0.518 | L | 1.44 | 0.872 |
| 0.81 | A | . | . | . | . | . |
| 0.81 | D | 2.695 | 0.789 | M | 1.06 | 0.398 |
| 0.81 | D | 3.45 | 0.922 | M | -1.32 | 0.798 |
| 0.09 | N | 2.49 | 0.726 | M | . | . |
| 0.81 | D | 0 | 0.065 | N | -2.75 | 0.908 |
| 0.81 | D | 2.38 | 0.686 | M | 0.06 | 0.618 |
| 0.588 | D | 2.35 | 0.675 | M | 2.41 | 0.158 |
| 0.489 | D | 2.66 | 0.779 | M | -2.69 | 0.93 |
| 0.81 | D | 1.995 | 0.541 | M | 0.71 | 0.512 |
| 0.81 | D | 1.685 | 0.433 | L | . | . |
| 0.398 | D | 2.485 | 0.724 | M | 2.82 | 0.109 |
| 0.413 | D | 2.3 | 0.657 | M | 2.87 | 0.104 |
| 0.81 | D | 2.555 | 0.746 | M | 3.38 | 0.058 |
| 0.588 | D | 2.175 | 0.61 | M | 0.52 | 0.553 |
| 0.588 | D | 2.215 | 0.625 | M | -0.64 | 0.721 |
| 0.588 | D | 1.245 | 0.314 | L | -6.3 | 0.998 |
| 0.495 | D | 2.515 | 0.733 | M | 0.84 | 0.475 |
| 0.81 | D | 3.135 | 0.882 | M | 2.45 | 0.15 |

| | | | | | | |
|---|---|---|---|---|---|---|
| 0.464 | D | 2.405 | 0.696 | M | 3.18 | 0.599 |
| 0.284 | N | 0.895 | 0.224 | L | -1.9 | 0.848 |
| 0.502 | D | 1.7 | 0.438 | L | 1.3 | 0.356 |
| 0.411 | D | 1.355 | 0.338 | L | -3.15 | 0.929 |
| 0.81 | D | 2.085 | 0.577 | M | -1.99 | 0.853 |
| 0.81 | A | . | . | . | . | . |
| . | . | . | . | . | . | . |
| 0.09 | N | 2.28 | 0.649 | M | -0.03 | 0.631 |
| 0.588 | A | 3.15 | 0.884 | M | -0.38 | 0.69 |
| 0.81 | D | 1.04 | 0.262 | L | 0.81 | 0.485 |
| 0.548 | D | 1.845 | 0.487 | L | -0.69 | 0.727 |
| 0.81 | D | 1.24 | 0.31 | L | -0.84 | 0.743 |
| 0.455 | D | 1.975 | 0.535 | M | 1.6 | 0.286 |
| . | . | . | . | . | . | . |
| 0.413 | D | 1.935 | 0.518 | L | 1.7 | 0.269 |
| 0.81 | D | 2.3 | 0.657 | M | 1.53 | 0.304 |
| 0.81 | D | 2.7 | 0.79 | M | -3.22 | 0.934 |
| 0.382 | D | 2.38 | 0.686 | M | 2.96 | 0.095 |
| 0.81 | D | 2.585 | 0.756 | M | 0.98 | 0.421 |
| 0.81 | D | 1.39 | 0.349 | L | -0.85 | 0.744 |
| 0.588 | D | 1.61 | 0.411 | L | -0.54 | 0.738 |
| . | . | . | . | . | . | . |
| 0.464 | D | 2.405 | 0.696 | M | 3.18 | 0.599 |
| 0.548 | D | 1.135 | 0.29 | L | -1.89 | 0.846 |
| 0.359 | D | 2.685 | 0.786 | M | -0.43 | 0.697 |
| 0.588 | D | 2.35 | 0.675 | M | 2.41 | 0.158 |
| 0.455 | D | 1.1 | 0.28 | L | -1.96 | 0.851 |
| 0.81 | D | 1.545 | 0.391 | L | 1.45 | 0.325 |
| 0.465 | D | 2.73 | 0.798 | M | 0.72 | 0.51 |
| . | . | . | . | . | . | . |
| 0.499 | D | 2.015 | 0.55 | M | -2.42 | 0.886 |
| 0.415 | D | 2.085 | 0.577 | M | 2.16 | 0.192 |
| 0.434 | D | 1.995 | 0.541 | M | -3.47 | 0.945 |
| 0.81 | D | 3.12 | 0.879 | M | -2.47 | 0.89 |
| 0.464 | D | 2.405 | 0.696 | M | 3.18 | 0.599 |
| 0.464 | D | 2.405 | 0.696 | M | 3.18 | 0.599 |
| 0.81 | D | 0.55 | 0.145 | N | -0.84 | 0.743 |
| 0.81 | D | 1.59 | 0.403 | L | 1.21 | 0.549 |
| 0.421 | D | 1.5 | 0.378 | L | 2.01 | 0.213 |
| 0.81 | D | 1.935 | 0.518 | L | -2.04 | 0.859 |
| 0.457 | D | 1.895 | 0.504 | L | -0.31 | 0.68 |
| 0.508 | D | 2.32 | 0.664 | M | -1.05 | 0.767 |
| 0.81 | D | 3.11 | 0.878 | M | 0.57 | 0.543 |

| | | | | | | |
|---|---|---|---|---|---|---|
| 0.81 | D | 2.045 | 0.56 | M | 1.42 | 0.332 |
| . | . | . | . | . | . | . |
| 0.81 | D | 0.345 | 0.112 | N | 0.24 | 0.596 |
| 0.81 | D | 2.58 | 0.754 | M | -0.22 | 0.665 |
| 0.588 | A | 3.15 | 0.884 | M | -0.38 | 0.69 |
| 0.81 | D | 2.175 | 0.61 | M | 1.92 | 0.231 |
| 0.81 | D | 1.72 | 0.444 | L | 0.72 | 0.51 |
| 0.464 | D | 2.405 | 0.696 | M | 3.18 | 0.599 |
| 0.371 | D | 2.765 | 0.808 | M | -0.15 | 0.652 |
| 0.81 | D | 3.3 | 0.905 | M | -5 | 0.985 |
| . | . | . | . | . | . | . |
| 0.508 | D | 2.14 | 0.599 | M | -2.77 | 0.909 |
| 0.81 | D | 3.46 | 0.923 | M | -0.37 | 0.689 |
| . | . | . | . | . | . | . |
| . | . | . | . | . | . | . |
| 0.81 | D | 3.12 | 0.879 | M | -2.47 | 0.89 |
| . | . | . | . | . | . | . |
| 0.81 | D | 1.845 | 0.487 | L | -2.85 | 0.914 |
| 0.448 | D | 0.895 | 0.224 | L | -2.71 | 0.906 |
| . | . | . | . | . | . | . |
| 0.81 | D | 1.7 | 0.438 | L | -0.7 | 0.764 |
| 0.81 | D | 2.175 | 0.61 | M | 1.88 | 0.239 |
| . | . | . | . | . | . | . |
| 0.464 | D | 2.405 | 0.696 | M | 3.18 | 0.599 |
| 0.588 | D | 2.35 | 0.675 | M | 2.41 | 0.158 |
| 0.81 | D | 1.24 | 0.31 | L | -0.9 | 0.749 |
| 0.81 | D | 1.685 | 0.433 | L | . | . |
| . | . | . | . | . | . | . |
| 0.468 | D | 2.05 | 0.565 | M | 2.64 | 0.219 |
| 0.81 | D | 3.265 | 0.9 | M | -2.02 | 0.855 |
| 0.81 | D | 0.28 | 0.1 | N | 0.04 | 0.621 |
| 0.588 | D | 2.67 | 0.782 | M | 2.19 | 0.187 |
| 0.81 | D | 0 | 0.065 | N | -0.95 | 0.754 |
| 0.548 | D | 1.415 | 0.356 | L | -1.68 | 0.829 |
| 0.81 | D | 1.39 | 0.349 | L | -0.85 | 0.744 |
| 0.487 | D | 0.695 | 0.18 | N | -0.9 | 0.792 |
| 0.499 | D | 0.345 | 0.112 | N | -0.5 | 0.743 |
| 0.81 | D | 1.655 | 0.424 | L | 0.84 | 0.475 |
| 0.81 | D | 2.12 | 0.588 | M | -3.46 | 0.945 |
| 0.81 | D | 3.39 | 0.916 | M | -0.21 | 0.663 |
| . | . | . | . | . | . | . |
| . | . | . | . | . | . | . |
| 0.81 | D | 0.28 | 0.1 | N | 0.04 | 0.621 |

| | | | | | | |
|---|---|---|---|---|---|---|
| . | . | . | . | . | . | . |
| 0.464 | D | 2.405 | 0.696 | M | 3.18 | 0.599 |
| 0.81 | D | 1.39 | 0.349 | L | -0.85 | 0.744 |
| 0.81 | D | 2.475 | 0.719 | M | 1.52 | 0.307 |
| 0.529 | D | 2.545 | 0.743 | M | -0.37 | 0.726 |
| 0.43 | D | 2.275 | 0.646 | M | -0.94 | 0.785 |
| 0.81 | D | 3.52 | 0.93 | H | -1.29 | 0.795 |
| 0.81 | D | 3.535 | 0.931 | H | 0.89 | 0.456 |
| 0.81 | D | 2.65 | 0.776 | M | -0.27 | 0.674 |
| 0.81 | D | . | . | . | . | . |
| 0.81 | A | . | . | . | . | . |
| 0.81 | D | 2.445 | 0.709 | M | 0.54 | 0.549 |
| 0.529 | D | 1.735 | 0.449 | L | -2.87 | 0.915 |
| . | . | . | . | . | . | . |
| 0.09 | N | 1.125 | 0.288 | L | 0.04 | 0.621 |
| 0.81 | D | 2.045 | 0.56 | M | 1.94 | 0.227 |
| 0.81 | D | 2.24 | 0.634 | M | -6.09 | 0.997 |
| . | . | . | . | . | . | . |
| 0.81 | D | 1.685 | 0.433 | L | . | . |
| 0.81 | D | 1.935 | 0.518 | L | 2.21 | 0.184 |
| 0.588 | D | 2.43 | 0.705 | M | -5.36 | 0.99 |
| 0.81 | D | 3.52 | 0.93 | H | -1.29 | 0.795 |
| . | . | . | . | . | . | . |
| 0.479 | D | 0 | 0.065 | N | -0.79 | 0.737 |
| 0.81 | D | 1.73 | 0.447 | L | 1.83 | 0.248 |
| 0.81 | D | 2.955 | 0.85 | M | . | . |
| 0.29 | N | 2.08 | 0.574 | M | 2.98 | 0.094 |
| 0.81 | D | 1.895 | 0.504 | L | 2.05 | 0.207 |
| 0.81 | D | 2.645 | 0.774 | M | 0.69 | 0.517 |
| 0.81 | D | 1.935 | 0.518 | L | 1.64 | 0.278 |
| 0.465 | D | 2.73 | 0.798 | M | 0.72 | 0.51 |
| 0.81 | D | 3.52 | 0.93 | H | -1.29 | 0.795 |
| . | . | . | . | . | . | . |
| 0.81 | D | 2.645 | 0.774 | M | -0.01 | 0.628 |
| 0.588 | D | 1.005 | 0.252 | L | 0.42 | 0.569 |
| 0.81 | D | 2.75 | 0.804 | M | 1.36 | 0.345 |
| 0.81 | D | 2.215 | 0.625 | M | -0.93 | 0.752 |
| 0.81 | D | 2.67 | 0.782 | M | 1.58 | 0.291 |
| . | . | . | . | . | . | . |
| 0.465 | D | 2.73 | 0.798 | M | 0.72 | 0.51 |
| 0.81 | D | 3.425 | 0.92 | M | -1.07 | 0.769 |
| 0.446 | D | 2.43 | 0.705 | M | 2.66 | 0.126 |
| 0.81 | D | 1.7 | 0.438 | L | . | . |

| | | | | | | |
|---|---|---|---|---|---|---|
| 0.588 | D | 2.515 | 0.733 | M | -0.6 | 0.767 |
| 0.51 | D | 2.22 | 0.629 | M | -0.36 | 0.688 |
| . | . | . | . | . | . | . |
| . | . | . | . | . | . | . |
| 0.29 | N | 2.08 | 0.574 | M | 2.98 | 0.094 |
| 0.441 | D | . | . | . | 1.7 | 0.269 |
| 0.81 | D | 1.845 | 0.487 | L | -0.87 | 0.746 |
| 0.588 | D | 2.43 | 0.705 | M | -5.36 | 0.99 |
| 0.516 | D | 1.61 | 0.411 | L | 2.98 | 0.094 |
| . | . | . | . | . | . | . |
| 0.455 | D | 0.69 | 0.17 | N | . | . |
| 0.81 | D | 1.15 | 0.293 | L | -0.83 | 0.742 |
| 0.81 | D | 5 | 0.999 | H | 0.5 | 0.556 |
| 0.81 | D | 5 | 0.999 | H | 0.5 | 0.556 |
| . | . | . | . | . | . | . |
| . | . | . | . | . | . | . |
| 0.81 | D | 2.9 | 0.839 | M | -0.56 | 0.712 |
| 0.81 | D | 1.59 | 0.403 | L | -2.01 | 0.862 |
| 0.81 | D | 2.375 | 0.684 | M | 0.17 | 0.605 |
| . | . | . | . | . | . | . |
| 0.479 | D | 0 | 0.065 | N | -0.79 | 0.737 |
| 0.81 | D | 2.255 | 0.642 | M | -0.86 | 0.745 |
| 0.81 | D | 2.98 | 0.855 | M | 0.54 | 0.549 |
| 0.588 | D | 2.275 | 0.646 | M | 2.64 | 0.128 |
| 0.81 | D | 0.695 | 0.18 | N | -0.83 | 0.742 |
| 0.81 | D | 1.95 | 0.525 | M | 1.63 | 0.28 |
| . | . | . | . | . | . | . |
| 0.455 | D | 0.69 | 0.17 | N | . | . |
| 0.529 | D | 2.07 | 0.568 | M | -1.04 | 0.766 |
| 0.464 | D | 2.405 | 0.696 | M | 3.18 | 0.599 |
| 0.465 | D | 2.73 | 0.798 | M | 0.72 | 0.51 |
| 0.81 | D | 1.15 | 0.293 | L | -0.83 | 0.742 |
| 0.465 | D | 2.73 | 0.798 | M | 0.72 | 0.51 |
| 0.81 | D | 1.39 | 0.349 | L | -0.85 | 0.744 |
| 0.81 | D | 1.87 | 0.496 | L | 1.78 | 0.257 |
| 0.457 | D | 0.55 | 0.145 | N | -2.66 | 0.903 |
| 0.495 | D | 2.515 | 0.733 | M | 0.84 | 0.475 |
| 0.81 | D | 2.98 | 0.855 | M | 0.54 | 0.549 |
| 0.502 | D | 1.7 | 0.438 | L | 1.3 | 0.356 |
| 0.81 | D | 3.085 | 0.874 | M | -0.67 | 0.725 |
| . | . | . | . | . | . | . |
| 0.81 | D | 3.035 | 0.865 | M | 1.07 | 0.396 |
| 0.588 | D | 1.005 | 0.252 | L | 0.42 | 0.569 |

| | | | | | | |
|---|---|---|---|---|---|---|
| . | . | . | . | . | . | . |
| 0.465 | D | 2.73 | 0.798 | M | 0.72 | 0.51 |
| 0.81 | D | 2.98 | 0.855 | M | 0.54 | 0.549 |
| 0.588 | D | 1.005 | 0.252 | L | 0.42 | 0.569 |
| 0.29 | N | 2.08 | 0.574 | M | 2.98 | 0.094 |
| 0.588 | D | 1.005 | 0.252 | L | 0.42 | 0.569 |
| 0.81 | D | 1.79 | 0.468 | L | 1.95 | 0.225 |
| 0.81 | D | 2.125 | 0.59 | M | -1.05 | 0.767 |
| . | . | . | . | . | . | . |
| 0.81 | D | 2.215 | 0.625 | M | -0.93 | 0.752 |
| 0.495 | D | 2.515 | 0.733 | M | 0.84 | 0.475 |
| 0.81 | D | 2.43 | 0.705 | M | 1.31 | 0.354 |
| . | . | . | . | . | . | . |
| 0.81 | D | 2.095 | 0.581 | M | -2.19 | 0.88 |
| 0.81 | D | 2.645 | 0.774 | M | -0.01 | 0.628 |
| 0.496 | D | 1.5 | 0.378 | L | -0.01 | 0.628 |
| 0.81 | D | 2.175 | 0.61 | M | 2.03 | 0.21 |
| . | . | . | . | . | . | . |
| 0.588 | D | 1.535 | 0.389 | L | -3.54 | 0.948 |
| 0.81 | D | 1.39 | 0.349 | L | -0.94 | 0.753 |
| 0.81 | D | 2.175 | 0.61 | M | 2.03 | 0.21 |
| 0.529 | D | 2.045 | 0.56 | M | -1.43 | 0.807 |
| . | . | . | . | . | . | . |
| 0.588 | D | 1.955 | 0.529 | M | 2.26 | 0.176 |
| . | . | . | . | . | . | . |
| 0.588 | D | 3.11 | 0.878 | M | -0.29 | 0.677 |
| 0.45 | D | 1.78 | 0.462 | L | 0.42 | 0.574 |
| 0.81 | A | . | . | . | . | . |
| . | . | . | . | . | . | . |
| 0.81 | D | 1.445 | 0.364 | L | -2.79 | 0.91 |
| 0.81 | D | 2.79 | 0.814 | M | -0.8 | 0.738 |
| 0.44 | D | 0.695 | 0.18 | N | -0.65 | 0.759 |
| 0.81 | D | 2.36 | 0.679 | M | -2.38 | 0.883 |
| 0.81 | D | 2.025 | 0.554 | M | 3.24 | 0.068 |
| . | . | . | . | . | . | . |
| 0.325 | D | 2.43 | 0.705 | M | -2.03 | 0.859 |
| 0.44 | D | 0.695 | 0.18 | N | -0.65 | 0.759 |
| 0.81 | D | . | . | . | 0.73 | 0.507 |
| 0.537 | D | 2.395 | 0.692 | M | 2.72 | 0.119 |
| 0.588 | D | 2.32 | 0.664 | M | 1.22 | 0.371 |
| 0.81 | D | 1.975 | 0.535 | M | -1.77 | 0.837 |
| 0.81 | D | 1.645 | 0.42 | L | -4.17 | 0.969 |
| 0.81 | D | 1.905 | 0.509 | L | 0.67 | 0.522 |

| | | | | | | |
|---|---|---|---|---|---|---|
| . | . | . | . | . | . | . |
| 0.588 | D | 3.11 | 0.878 | M | -0.29 | 0.677 |
| 0.524 | D | 1.585 | 0.399 | L | -3.48 | 0.946 |
| 0.81 | D | 1.24 | 0.31 | L | -1.98 | 0.852 |
| 0.548 | D | 1.61 | 0.411 | L | -0.02 | 0.629 |
| 0.588 | D | 2.045 | 0.56 | M | 0.41 | 0.571 |
| 0.588 | D | 3.665 | 0.943 | H | -0.15 | 0.652 |
| . | . | . | . | . | . | . |
| 0.323 | D | 0.805 | 0.202 | L | 0.9 | 0.456 |
| 0.81 | D | . | . | . | -6.44 | 0.997 |
| 0.529 | D | 1.735 | 0.449 | L | -2.87 | 0.915 |
| 0.81 | D | 1.895 | 0.504 | L | 1.72 | 0.266 |
| 0.81 | D | 2.505 | 0.729 | M | -3.25 | 0.935 |
| 0.81 | D | 2.415 | 0.698 | M | 1.28 | 0.36 |
| 0.81 | D | 2.285 | 0.652 | M | -3.37 | 0.941 |
| 0.385 | D | 1.245 | 0.314 | L | -0.97 | 0.757 |
| 0.328 | D | 2.73 | 0.798 | M | 0.78 | 0.494 |
| 0.81 | D | 1.975 | 0.535 | M | -0.63 | 0.72 |
| 0.238 | N | 0.805 | 0.202 | L | 2.57 | 0.136 |
| 0.328 | D | 2.73 | 0.798 | M | 0.78 | 0.494 |
| 0.31 | N | 0.785 | 0.195 | N | -1.5 | 0.813 |
| 0.419 | D | 0.695 | 0.18 | N | -0.62 | 0.758 |
| 0.529 | D | 2.045 | 0.56 | M | -1.43 | 0.807 |
| 0.328 | D | 2.73 | 0.798 | M | 0.78 | 0.494 |
| 0.81 | D | 1.905 | 0.509 | L | 0.92 | 0.449 |
| . | . | . | . | . | . | . |
| 0.81 | D | 1.975 | 0.535 | M | -0.63 | 0.72 |
| 0.81 | D | 2.62 | 0.767 | M | 0.5 | 0.556 |
| 0.81 | D | 2.175 | 0.61 | M | 2.03 | 0.21 |
| 0.81 | D | 2.445 | 0.709 | M | -1.03 | 0.764 |
| 0.81 | D | 2.565 | 0.75 | M | 0.8 | 0.488 |
| 0.81 | D | 2.42 | 0.7 | M | 0.95 | 0.433 |
| 0.588 | D | 3.665 | 0.943 | H | -0.15 | 0.652 |
| 0.328 | D | 2.73 | 0.798 | M | 0.78 | 0.494 |
| 0.81 | D | 2.62 | 0.767 | M | 0.5 | 0.556 |
| 0.537 | D | 2.395 | 0.692 | M | 2.72 | 0.119 |
| 0.529 | D | 2.49 | 0.726 | M | -0.04 | 0.632 |
| . | . | . | . | . | . | . |
| 0.81 | D | 1.905 | 0.509 | L | -3.41 | 0.943 |
| 0.81 | D | 3.135 | 0.882 | M | -0.96 | 0.756 |
| 0.588 | D | 2.125 | 0.59 | M | -0.15 | 0.652 |
| 0.81 | D | 1.24 | 0.31 | L | -1.98 | 0.852 |
| 0.328 | D | 2.73 | 0.798 | M | 0.78 | 0.494 |

| | | | | | | |
|---|---|---|---|---|---|---|
| 0.524 | D | 1.39 | 0.349 | L | 2.26 | 0.176 |
| 0.81 | D | 2.62 | 0.767 | M | 0.5 | 0.556 |
| 0.81 | D | 2.77 | 0.809 | M | 3.45 | 0.053 |

| FATHMM_pr | PROVEAN_sc | PROVEAN_co | PROVEAN_pr | VEST4_score | VEST4_ranks | MetaSVM_sc |
|---|---|---|---|---|---|---|
| . | . | . | . | . | . | . |
| . | . | . | . | . | . | . |
| T | -2.34 | 0.823 | N | 0.504 | 0.714 | -0.532 |
| T | -6.19 | 0.903 | D | 0.588 | 0.661 | -0.873 |
| D | -2.14 | 0.498 | N | 0.303 | 0.342 | 0.365 |
| T | -1.37 | 0.34 | N | 0.429 | 0.635 | -0.71 |
| D | -5.15 | 0.838 | D | 0.478 | 0.513 | 0.946 |
| T | -3.37 | 0.667 | D | 0.425 | 0.479 | -0.307 |
| D | -5.86 | 0.886 | D | 0.866 | 0.889 | 0.78 |
| . | -0.46 | 0.15 | N | 0.227 | 0.347 | -0.719 |
| T | -2.45 | 0.689 | N | 0.774 | 0.806 | -1.156 |
| T | -2.34 | 0.823 | N | 0.504 | 0.714 | -0.532 |
| T | -2.82 | 0.614 | D | 0.55 | 0.577 | -0.45 |
| T | -4.25 | 0.766 | D | 0.882 | 0.88 | -0.237 |
| . | . | . | . | . | . | . |
| T | -1.96 | 0.454 | N | 0.345 | 0.386 | -0.378 |
| D | -1.27 | 0.358 | N | 0.272 | 0.308 | -0.3 |
| T | -2.21 | 0.497 | N | 0.693 | 0.698 | -0.687 |
| T | -2.45 | 0.689 | N | 0.774 | 0.806 | -1.156 |
| T | -3.49 | 0.712 | D | 0.654 | 0.683 | -0.924 |
| D | -1.65 | 0.395 | N | 0.757 | 0.774 | 0.328 |
| . | . | . | . | . | . | . |
| T | -5.21 | 0.838 | D | 0.557 | 0.582 | 0.003 |
| T | -3.11 | 0.685 | D | 0.877 | 0.875 | -1.058 |
| . | . | . | . | 0.91 | 0.912 | . |
| T | -3.97 | 0.757 | D | 0.796 | 0.898 | -0.783 |
| T | -4 | 0.747 | D | 0.906 | 0.909 | 0.636 |
| . | -1.66 | 0.397 | N | 0.355 | 0.449 | -0.909 |
| D | -0.72 | 0.204 | N | 0.294 | 0.333 | -0.101 |
| T | -1.77 | 0.418 | N | 0.561 | 0.585 | 0.481 |
| T | -2.45 | 0.689 | N | 0.774 | 0.806 | -1.156 |
| D | -3.47 | 0.689 | D | 0.54 | 0.618 | 0.705 |
| T | -2.46 | 0.537 | N | 0.626 | 0.641 | -0.633 |
| . | -0.46 | 0.15 | N | 0.227 | 0.347 | -0.719 |
| T | -2.82 | 0.595 | D | 0.622 | 0.637 | -1.131 |
| T | -2.74 | 0.582 | D | 0.258 | 0.292 | -1.088 |
| T | -2.37 | 0.582 | N | 0.645 | 0.671 | -1.175 |
| T | -2 | 0.461 | N | 0.746 | 0.746 | -0.521 |
| T | -4.17 | 0.755 | D | 0.72 | 0.722 | -0.116 |
| D | -0.23 | 0.107 | N | 0.746 | 0.746 | 1.057 |
| T | -2.21 | 0.586 | N | 0.567 | 0.591 | -0.611 |
| T | -1.82 | 0.428 | N | 0.199 | 0.255 | -0.882 |

| | | | | | | |
|---|---|---|---|---|---|---|
| T | -2.34 | 0.823 | N | 0.504 | 0.714 | -0.532 |
| D | -1.09 | 0.283 | N | 0.105 | 0.142 | -0.591 |
| T | -3.56 | 0.689 | D | 0.46 | 0.524 | -1.021 |
| D | -1.4 | 0.399 | N | 0.569 | 0.628 | 0.292 |
| D | -1.65 | 0.395 | N | 0.757 | 0.774 | 0.328 |
| . | . | . | . | 0.651 | 0.662 | . |
| . | . | . | . | . | . | . |
| T | -1.73 | 0.467 | N | 0.295 | 0.355 | -1.01 |
| T | -1.87 | 0.602 | N | 0.916 | 0.919 | 0.095 |
| T | -1.63 | 0.391 | N | 0.719 | 0.721 | -0.948 |
| T | -1.38 | 0.364 | N | 0.494 | 0.527 | -0.228 |
| T | -2.71 | 0.578 | D | 0.444 | 0.482 | -0.503 |
| T | -2.43 | 0.602 | N | 0.337 | 0.542 | -0.806 |
| . | . | . | . | . | . | . |
| T | -2.76 | 0.586 | D | 0.095 | 0.133 | -0.991 |
| T | -6.81 | 0.936 | D | 0.861 | 0.936 | -0.811 |
| D | -4.89 | 0.814 | D | 0.878 | 0.876 | 0.937 |
| T | -2.93 | 0.613 | D | 0.686 | 0.692 | -1.077 |
| T | -1.13 | 0.308 | N | 0.516 | 0.551 | -0.894 |
| T | -1.58 | 0.382 | N | 0.713 | 0.716 | -0.875 |
| T | -2.77 | 0.636 | D | 0.284 | 0.35 | -0.17 |
| . | . | . | . | . | . | . |
| T | -2.34 | 0.823 | N | 0.504 | 0.714 | -0.532 |
| D | -4.56 | 0.786 | D | 0.636 | 0.649 | 0.108 |
| T | -2.48 | 0.541 | N | 0.689 | 0.695 | -0.878 |
| T | -2.45 | 0.689 | N | 0.774 | 0.806 | -1.156 |
| D | -0.77 | 0.214 | N | 0.371 | 0.413 | 0.181 |
| T | -0.79 | 0.219 | N | 0.815 | 0.811 | -1.102 |
| T | -3.19 | 0.662 | D | 0.634 | 0.666 | -0.518 |
| . | . | . | . | . | . | . |
| D | -5.06 | 0.828 | D | 0.235 | 0.654 | 0.483 |
| T | -2.57 | 0.555 | D | 0.64 | 0.652 | -1.061 |
| D | -3.11 | 0.637 | D | 0.742 | 0.742 | 0.323 |
| D | -5.86 | 0.886 | D | 0.866 | 0.889 | 0.78 |
| T | -2.34 | 0.823 | N | 0.504 | 0.714 | -0.532 |
| T | -2.34 | 0.823 | N | 0.504 | 0.714 | -0.532 |
| T | -3.23 | 0.651 | D | 0.543 | 0.715 | -0.25 |
| T | -1.46 | 0.51 | N | 0.672 | 0.785 | -0.718 |
| T | -1.02 | 0.268 | N | 0.298 | 0.454 | -1.108 |
| D | -3.43 | 0.689 | D | 0.746 | 0.77 | 0.269 |
| T | -1.31 | 0.37 | N | 0.626 | 0.842 | -0.827 |
| T | -1.96 | 0.454 | N | 0.345 | 0.386 | -0.378 |
| T | -3.27 | 0.655 | D | 0.456 | 0.493 | -0.423 |

| | | | | | | |
|---|---|---|---|---|---|---|
| T | -1.35 | 0.336 | N | 0.666 | 0.776 | -1.002 |
| . | . | . | . | . | . | . |
| T | -0.07 | 0.08 | N | 0.732 | 0.769 | -0.944 |
| T | -2.87 | 0.778 | D | 0.81 | 0.808 | -0.005 |
| T | -1.87 | 0.602 | N | 0.916 | 0.919 | 0.095 |
| T | -4.96 | 0.82 | D | 0.853 | 0.849 | -0.942 |
| T | -2.01 | 0.475 | N | 0.432 | 0.471 | -0.818 |
| T | -2.34 | 0.823 | N | 0.504 | 0.714 | -0.532 |
| T | -2.72 | 0.741 | D | 0.753 | 0.776 | -0.384 |
| D | -6.53 | 0.917 | D | 0.481 | 0.516 | 1.095 |
| . | . | . | . | . | . | . |
| D | -2.14 | 0.498 | N | 0.303 | 0.342 | 0.365 |
| T | -4.82 | 0.809 | D | 0.747 | 0.762 | 0.252 |
| . | . | . | . | . | . | . |
| . | . | . | . | . | . | . |
| D | -5.86 | 0.886 | D | 0.866 | 0.889 | 0.78 |
| . | . | . | . | . | . | . |
| D | -0.67 | 0.193 | N | 0.52 | 0.55 | 0.062 |
| D | -3.44 | 0.685 | D | 0.395 | 0.445 | -0.923 |
| . | . | . | . | . | . | . |
| T | -4.14 | 0.752 | D | 0.865 | 0.862 | -0.001 |
| T | -1.93 | 0.449 | N | 0.547 | 0.573 | -0.954 |
| . | . | . | . | . | . | . |
| T | -2.34 | 0.823 | N | 0.504 | 0.714 | -0.532 |
| T | -2.45 | 0.689 | N | 0.774 | 0.806 | -1.156 |
| T | -1.54 | 0.488 | N | 0.737 | 0.737 | -0.649 |
| . | -0.46 | 0.15 | N | 0.227 | 0.347 | -0.719 |
| . | . | . | . | . | . | . |
| T | -2.94 | 0.683 | D | 0.305 | 0.344 | -1.06 |
| D | -1.98 | 0.458 | N | 0.506 | 0.538 | 0.584 |
| T | -2.33 | 0.516 | N | 0.663 | 0.672 | -0.63 |
| T | -1 | 0.264 | N | 0.548 | 0.574 | -1.044 |
| T | -2.68 | 0.573 | D | 0.798 | 0.794 | -0.56 |
| D | -3.78 | 0.737 | D | 0.214 | 0.245 | -0.824 |
| T | -1.58 | 0.382 | N | 0.713 | 0.716 | -0.875 |
| T | -1.61 | 0.387 | N | 0.549 | 0.596 | -0.13 |
| T | -0.48 | 0.195 | N | 0.755 | 0.754 | -0.605 |
| T | -1.86 | 0.435 | N | 0.674 | 0.769 | -0.57 |
| D | -2.76 | 0.586 | D | 0.749 | 0.748 | 0.737 |
| T | -4.02 | 0.916 | D | 0.853 | 0.86 | 0.047 |
| . | . | . | . | . | . | . |
| . | . | . | . | . | . | . |
| T | -2.33 | 0.516 | N | 0.663 | 0.672 | -0.63 |

| | | | | | | |
|---|---|---|---|---|---|---|
| . | . | . | . | . | . | . |
| T | -2.34 | 0.823 | N | 0.504 | 0.714 | -0.532 |
| T | -1.58 | 0.382 | N | 0.713 | 0.716 | -0.875 |
| T | -3.3 | 0.659 | D | 0.855 | 0.851 | -0.989 |
| T | -2.22 | 0.63 | N | 0.493 | 0.527 | -0.682 |
| T | -4.07 | 0.779 | D | 0.708 | 0.749 | -0.276 |
| T | -3.49 | 0.682 | D | 0.706 | 0.739 | 0.569 |
| T | -6.41 | 0.914 | D | 0.737 | 0.737 | 0.379 |
| T | -6.83 | 0.93 | D | 0.84 | 0.836 | 0.033 |
| . | . | . | . | . | . | . |
| . | . | . | . | 0.986 | 0.998 | . |
| T | -5.15 | 0.902 | D | 0.792 | 0.93 | -0.551 |
| D | -0.6 | 0.185 | N | 0.408 | 0.449 | 0.093 |
| . | . | . | . | . | . | . |
| T | -2.22 | 0.498 | N | 0.35 | 0.392 | -0.89 |
| T | -2.84 | 0.599 | D | 0.569 | 0.592 | -1.06 |
| D | -1.62 | 0.405 | N | 0.695 | 0.7 | 1.067 |
| . | . | . | . | . | . | . |
| . | -0.46 | 0.15 | N | 0.227 | 0.347 | -0.719 |
| T | -0.99 | 0.281 | N | 0.179 | 0.193 | -1.125 |
| D | -1.97 | 0.456 | N | 0.517 | 0.571 | 1.097 |
| T | -3.49 | 0.682 | D | 0.706 | 0.739 | 0.569 |
| . | . | . | . | . | . | . |
| T | -1.68 | 0.445 | N | 0.482 | 0.576 | -0.37 |
| T | -2.23 | 0.5 | N | 0.704 | 0.708 | -0.744 |
| . | -0.72 | 0.204 | N | 0.23 | 0.259 | -0.697 |
| T | -3.03 | 0.627 | D | 0.204 | 0.226 | -1.124 |
| T | -1.56 | 0.378 | N | 0.168 | 0.349 | -0.882 |
| T | -1.43 | 0.352 | N | 0.382 | 0.423 | -0.754 |
| T | -3.6 | 0.694 | D | 0.728 | 0.729 | -0.823 |
| T | -3.19 | 0.662 | D | 0.634 | 0.666 | -0.518 |
| T | -3.49 | 0.682 | D | 0.706 | 0.739 | 0.569 |
| . | . | . | . | . | . | . |
| T | -4.18 | 0.755 | D | 0.635 | 0.82 | -0.267 |
| T | -3.43 | 0.674 | D | 0.397 | 0.438 | -0.811 |
| T | -3.18 | 0.655 | D | 0.542 | 0.592 | -0.662 |
| T | -6.71 | 0.925 | D | 0.879 | 0.877 | 0.124 |
| T | -4 | 0.741 | D | 0.695 | 0.7 | -0.902 |
| . | . | . | . | . | . | . |
| T | -3.19 | 0.662 | D | 0.634 | 0.666 | -0.518 |
| T | -3.13 | 0.639 | D | 0.847 | 0.843 | 0.449 |
| T | -6.61 | 0.921 | D | 0.685 | 0.691 | -1.164 |
| . | -0.36 | 0.174 | N | 0.864 | 0.866 | -0.367 |

| | | | | | | |
|---|---|---|---|---|---|---|
| T | -0.91 | 0.573 | N | 0.554 | 0.579 | -0.291 |
| T | -7.14 | 0.95 | D | 0.825 | 0.826 | -0.384 |
| . | . | . | . | . | . | . |
| . | . | . | . | . | . | . |
| T | -3.03 | 0.627 | D | 0.204 | 0.226 | -1.124 |
| T | -0.54 | 0.166 | N | 0.602 | 0.62 | -0.916 |
| T | -4.45 | 0.777 | D | 0.843 | 0.839 | 0.102 |
| D | -1.97 | 0.456 | N | 0.517 | 0.571 | 1.097 |
| T | -1.9 | 0.443 | N | 0.395 | 0.436 | -1.222 |
| . | . | . | . | . | . | . |
| . | -1.62 | 0.441 | N | 0.447 | 0.491 | -0.469 |
| T | -1.73 | 0.484 | N | 0.518 | 0.548 | -0.576 |
| T | -4.2 | 0.757 | D | 0.782 | 0.78 | -0.147 |
| T | -4.2 | 0.757 | D | 0.782 | 0.78 | -0.147 |
| . | . | . | . | . | . | . |
| . | . | . | . | . | . | . |
| T | -7.48 | 0.95 | D | 0.742 | 0.742 | 0.257 |
| D | -0.76 | 0.212 | N | 0.357 | 0.399 | 0.191 |
| T | -2.48 | 0.541 | N | 0.922 | 0.925 | -0.248 |
| . | . | . | . | . | . | . |
| T | -1.68 | 0.445 | N | 0.482 | 0.576 | -0.37 |
| T | -2.73 | 0.639 | D | 0.813 | 0.891 | -0.17 |
| T | -4.25 | 0.766 | D | 0.882 | 0.88 | -0.237 |
| T | -2.9 | 0.608 | D | 0.625 | 0.724 | -1.097 |
| T | -4.14 | 0.762 | D | 0.892 | 0.891 | -0.622 |
| T | -0.9 | 0.242 | N | 0.527 | 0.577 | -1.053 |
| . | . | . | . | . | . | . |
| . | -1.62 | 0.441 | N | 0.447 | 0.491 | -0.469 |
| T | -2.09 | 0.494 | N | 0.233 | 0.267 | -0.529 |
| T | -2.34 | 0.823 | N | 0.504 | 0.714 | -0.532 |
| T | -3.19 | 0.662 | D | 0.634 | 0.666 | -0.518 |
| T | -1.73 | 0.484 | N | 0.518 | 0.548 | -0.576 |
| T | -3.19 | 0.662 | D | 0.634 | 0.666 | -0.518 |
| T | -1.58 | 0.382 | N | 0.713 | 0.716 | -0.875 |
| T | -1.87 | 0.437 | N | 0.532 | 0.561 | -1.06 |
| D | -0.99 | 0.264 | N | 0.329 | 0.37 | -0.165 |
| T | -2.21 | 0.586 | N | 0.567 | 0.591 | -0.611 |
| T | -4.25 | 0.766 | D | 0.882 | 0.88 | -0.237 |
| T | -3.56 | 0.689 | D | 0.46 | 0.524 | -1.021 |
| T | -3.64 | 0.698 | D | 0.795 | 0.791 | 0.069 |
| . | . | . | . | . | . | . |
| T | -5.52 | 0.871 | D | 0.825 | 0.848 | -0.686 |
| T | -3.43 | 0.674 | D | 0.397 | 0.438 | -0.811 |

| | | | | | | |
|---|---|---|---|---|---|---|
| . | . | . | . | . | . | . |
| T | -3.19 | 0.662 | D | 0.634 | 0.666 | -0.518 |
| T | -4.25 | 0.766 | D | 0.882 | 0.88 | -0.237 |
| T | -3.43 | 0.674 | D | 0.397 | 0.438 | -0.811 |
| T | -3.03 | 0.627 | D | 0.204 | 0.226 | -1.124 |
| T | -3.43 | 0.674 | D | 0.397 | 0.438 | -0.811 |
| T | -2 | 0.461 | N | 0.375 | 0.417 | -1.093 |
| T | -1.34 | 0.346 | N | 0.488 | 0.522 | 0.073 |
| . | . | . | . | . | . | . |
| T | -6.71 | 0.925 | D | 0.879 | 0.877 | 0.124 |
| T | -2.21 | 0.586 | N | 0.567 | 0.591 | -0.611 |
| T | -1.69 | 0.452 | N | 0.268 | 0.396 | -0.953 |
| . | . | . | . | . | . | . |
| D | -2.05 | 0.47 | N | 0.192 | 0.323 | 0.525 |
| T | -4.18 | 0.755 | D | 0.635 | 0.82 | -0.267 |
| T | -4.4 | 0.821 | D | 0.632 | 0.646 | -0.484 |
| T | -4.68 | 0.797 | D | 0.677 | 0.684 | -1.02 |
| . | . | . | . | . | . | . |
| D | -2.55 | 0.566 | D | 0.51 | 0.662 | 0.85 |
| T | -0.75 | 0.21 | N | 0.534 | 0.562 | 0.075 |
| T | -4.68 | 0.797 | D | 0.677 | 0.684 | -1.02 |
| T | -4.61 | 0.814 | D | 0.423 | 0.587 | 0.028 |
| . | . | . | . | . | . | . |
| T | -5.57 | 0.865 | D | 0.895 | 0.895 | -1.118 |
| . | . | . | . | . | . | . |
| T | -2.87 | 0.603 | D | 0.738 | 0.782 | 0.078 |
| T | -0.61 | 0.236 | N | 0.253 | 0.414 | -0.725 |
| . | . | . | . | 0.978 | 0.991 | . |
| . | . | . | . | . | . | . |
| D | -1.25 | 0.316 | N | 0.773 | 0.77 | -0.255 |
| T | -3.33 | 0.724 | D | 0.971 | 0.996 | 0.269 |
| T | -2.23 | 0.5 | N | 0.781 | 0.778 | -0.238 |
| D | -4.2 | 0.769 | D | 0.782 | 0.779 | 0.751 |
| T | -1.74 | 0.412 | N | 0.822 | 0.863 | -1.214 |
| . | . | . | . | . | . | . |
| D | -1.06 | 0.277 | N | 0.346 | 0.387 | -0.13 |
| T | -2.23 | 0.5 | N | 0.781 | 0.778 | -0.238 |
| T | -5.71 | 0.875 | D | 0.9 | 0.9 | 0.335 |
| T | -1.33 | 0.34 | N | 0.423 | 0.579 | -1.144 |
| T | -0.93 | 0.249 | N | 0.276 | 0.313 | -1.088 |
| D | -5.93 | 0.89 | D | 0.841 | 0.837 | 0.459 |
| D | -2.7 | 0.576 | D | 0.852 | 0.848 | 1.019 |
| T | -4.75 | 0.803 | D | 0.741 | 0.789 | -0.682 |

| | | | | | | |
|---|---|---|---|---|---|---|
| . | . | . | . | . | . | . |
| T | -2.87 | 0.603 | D | 0.738 | 0.782 | 0.078 |
| D | -1.92 | 0.47 | N | 0.255 | 0.288 | -0.591 |
| D | -2.14 | 0.485 | N | 0.589 | 0.783 | 0.171 |
| T | -3.13 | 0.639 | D | 0.655 | 0.665 | -0.624 |
| T | -4.71 | 0.804 | D | 0.739 | 0.787 | -0.718 |
| T | -6.65 | 0.923 | D | 0.582 | 0.605 | -0.009 |
| . | . | . | . | . | . | . |
| T | -1.27 | 0.374 | N | 0.116 | 0.138 | -1.022 |
| D | -1.08 | 0.281 | N | 0.5 | 0.533 | 1.048 |
| D | -0.6 | 0.185 | N | 0.408 | 0.449 | 0.093 |
| T | -2.79 | 0.591 | D | 0.637 | 0.656 | -1.015 |
| D | -6.67 | 0.923 | D | 0.888 | 0.891 | 0.93 |
| T | -4.7 | 0.825 | D | 0.682 | 0.69 | -0.999 |
| D | -3.71 | 0.707 | D | 0.788 | 0.785 | 0.636 |
| T | -2.69 | 0.574 | D | 0.183 | 0.294 | -0.678 |
| T | -2.34 | 0.772 | N | 0.625 | 0.641 | -0.821 |
| T | -3.28 | 0.656 | D | 0.742 | 0.742 | -0.193 |
| T | -1.36 | 0.338 | N | 0.249 | 0.281 | -1.044 |
| T | -2.34 | 0.772 | N | 0.625 | 0.641 | -0.821 |
| D | -3.02 | 0.672 | D | 0.077 | 0.059 | -0.638 |
| T | -0.4 | 0.138 | N | 0.771 | 0.797 | -0.495 |
| T | -4.61 | 0.814 | D | 0.423 | 0.587 | 0.028 |
| T | -2.34 | 0.772 | N | 0.625 | 0.641 | -0.821 |
| T | -1.2 | 0.306 | N | 0.26 | 0.353 | -0.958 |
| . | . | . | . | . | . | . |
| T | -3.28 | 0.656 | D | 0.742 | 0.742 | -0.193 |
| T | -2.43 | 0.568 | N | 0.708 | 0.756 | -0.437 |
| T | -4.68 | 0.797 | D | 0.677 | 0.684 | -1.02 |
| T | -1.79 | 0.422 | N | 0.388 | 0.429 | -0.096 |
| T | -1.4 | 0.433 | N | 0.526 | 0.657 | -0.685 |
| T | -7.45 | 0.949 | D | 0.855 | 0.851 | -0.561 |
| T | -6.65 | 0.923 | D | 0.582 | 0.605 | -0.009 |
| T | -2.34 | 0.772 | N | 0.625 | 0.641 | -0.821 |
| T | -2.43 | 0.568 | N | 0.708 | 0.756 | -0.437 |
| T | -1.33 | 0.34 | N | 0.423 | 0.579 | -1.144 |
| T | -4.6 | 0.82 | D | 0.29 | 0.328 | -0.849 |
| . | . | . | . | . | . | . |
| D | -3.68 | 0.703 | D | 0.667 | 0.676 | -0.219 |
| T | -8.43 | 0.972 | D | 0.891 | 0.934 | 0.665 |
| T | -3.11 | 0.637 | D | 0.77 | 0.792 | -0.394 |
| D | -2.14 | 0.485 | N | 0.589 | 0.783 | 0.171 |
| T | -2.34 | 0.772 | N | 0.625 | 0.641 | -0.821 |

| | | | | | | |
|---|---|---|---|---|---|---|
| T | -1.34 | 0.36 | N | 0.254 | 0.287 | -1.188 |
| T | -2.43 | 0.568 | N | 0.708 | 0.756 | -0.437 |
| T | -2.81 | 0.594 | D | 0.806 | 0.802 | -1.165 |

| MetaSVM_ra | MetaSVM_pr | MetaLR_scor | MetaLR_rank | MetaLR_pred | MetaRNN_sc | MetaRNN_ra |
|---|---|---|---|---|---|---|
| . | . | . | . | . | . | . |
| . | . | . | . | . | . | . |
| 0.675 | T | 0.24 | 0.608 | T | 0.009 | 0.002 |
| 0.503 | T | 0.163 | 0.5 | T | 0.505 | 0.617 |
| 0.886 | D | 0.639 | 0.874 | D | 0.005 | 0.001 |
| 0.599 | T | 0.202 | 0.559 | T | 0.25 | 0.424 |
| 0.963 | D | 0.892 | 0.964 | D | 0.409 | 0.561 |
| 0.748 | T | 0.34 | 0.706 | T | 0.722 | 0.737 |
| 0.942 | D | 0.838 | 0.946 | D | 0.083 | 0.138 |
| 0.595 | T | 0.227 | 0.591 | T | 0.021 | 0.005 |
| 0.009 | T | 0.074 | 0.3 | T | 0.06 | 0.073 |
| 0.675 | T | 0.24 | 0.608 | T | 0.009 | 0.002 |
| 0.704 | T | 0.343 | 0.709 | T | 0.393 | 0.551 |
| 0.767 | T | 0.389 | 0.744 | T | 0.075 | 0.115 |
| . | . | . | . | . | . | . |
| 0.727 | T | 0.403 | 0.754 | T | 0.009 | 0.002 |
| 0.75 | T | 0.472 | 0.797 | T | 0.185 | 0.339 |
| 0.611 | T | 0.175 | 0.518 | T | 0.663 | 0.702 |
| 0.009 | T | 0.074 | 0.3 | T | 0.06 | 0.073 |
| 0.45 | T | 0.136 | 0.451 | T | 0.128 | 0.244 |
| 0.88 | D | 0.699 | 0.896 | D | 0.819 | 0.811 |
| . | . | . | . | . | . | . |
| 0.823 | D | 0.488 | 0.805 | T | 0.628 | 0.684 |
| 0.124 | T | 0.115 | 0.408 | T | 0.51 | 0.62 |
| . | . | . | . | . | . | . |
| 0.561 | T | 0.218 | 0.58 | T | 0.757 | 0.761 |
| 0.924 | D | 0.703 | 0.898 | D | 0.91 | 0.904 |
| 0.47 | T | 0.135 | 0.449 | T | 0.237 | 0.407 |
| 0.801 | T | 0.474 | 0.798 | T | 0.278 | 0.453 |
| 0.903 | D | 0.716 | 0.902 | D | 0.537 | 0.635 |
| 0.009 | T | 0.074 | 0.3 | T | 0.06 | 0.073 |
| 0.932 | D | 0.848 | 0.949 | D | 0.015 | 0.003 |
| 0.635 | T | 0.269 | 0.641 | T | 0.136 | 0.259 |
| 0.595 | T | 0.227 | 0.591 | T | 0.021 | 0.005 |
| 0.017 | T | 0.036 | 0.155 | T | 0.012 | 0.003 |
| 0.058 | T | 0.032 | 0.139 | T | 0.171 | 0.318 |
| 0.005 | T | 0.052 | 0.219 | T | 0.409 | 0.561 |
| 0.679 | T | 0.288 | 0.66 | T | 0.621 | 0.68 |
| 0.797 | T | 0.454 | 0.787 | T | 0.011 | 0.002 |
| 0.983 | D | 0.981 | 0.994 | D | 0.66 | 0.701 |
| 0.644 | T | 0.258 | 0.629 | T | 0.069 | 0.098 |
| 0.496 | T | 0.08 | 0.316 | T | 0.324 | 0.497 |

| | | | | | | |
|---|---|---|---|---|---|---|
| 0.675 | T | 0.24 | 0.608 | T | 0.009 | 0.002 |
| 0.652 | T | 0.397 | 0.749 | T | 0.168 | 0.314 |
| 0.233 | T | 0.103 | 0.38 | T | 0.013 | 0.003 |
| 0.874 | D | 0.581 | 0.849 | D | 0.688 | 0.717 |
| 0.88 | D | 0.699 | 0.896 | D | 0.819 | 0.811 |
| . | . | . | . | . | . | . |
| . | . | . | . | . | . | . |
| 0.27 | T | 0.17 | 0.51 | T | 0.156 | 0.293 |
| 0.841 | D | 0.51 | 0.816 | D | 0.72 | 0.736 |
| 0.413 | T | 0.193 | 0.546 | T | 0.461 | 0.593 |
| 0.769 | T | 0.396 | 0.749 | T | 0.402 | 0.557 |
| 0.685 | T | 0.335 | 0.702 | T | 0.314 | 0.488 |
| 0.548 | T | 0.142 | 0.462 | T | 0.161 | 0.302 |
| . | . | . | . | . | . | . |
| 0.322 | T | 0.118 | 0.415 | T | 0.045 | 0.036 |
| 0.545 | T | 0.2 | 0.556 | T | 0.693 | 0.72 |
| 0.962 | D | 0.885 | 0.962 | D | 0.769 | 0.77 |
| 0.08 | T | 0.035 | 0.153 | T | 0.358 | 0.525 |
| 0.486 | T | 0.17 | 0.51 | T | 0.469 | 0.598 |
| 0.501 | T | 0.14 | 0.46 | T | 0.082 | 0.134 |
| 0.784 | T | 0.46 | 0.79 | T | 0.14 | 0.266 |
| . | . | . | . | . | . | . |
| 0.675 | T | 0.24 | 0.608 | T | 0.009 | 0.002 |
| 0.843 | D | 0.518 | 0.82 | D | 0.07 | 0.1 |
| 0.499 | T | 0.175 | 0.519 | T | 0.322 | 0.495 |
| 0.009 | T | 0.074 | 0.3 | T | 0.06 | 0.073 |
| 0.856 | D | 0.605 | 0.86 | D | 0.334 | 0.506 |
| 0.039 | T | 0.049 | 0.209 | T | 0.523 | 0.627 |
| 0.68 | T | 0.237 | 0.604 | T | 0.01 | 0.002 |
| . | . | . | . | . | . | . |
| 0.903 | D | 0.677 | 0.888 | D | 0.312 | 0.487 |
| 0.116 | T | 0.049 | 0.21 | T | 0.526 | 0.629 |
| 0.879 | D | 0.665 | 0.884 | D | 0.553 | 0.644 |
| 0.942 | D | 0.838 | 0.946 | D | 0.083 | 0.138 |
| 0.675 | T | 0.24 | 0.608 | T | 0.009 | 0.002 |
| 0.675 | T | 0.24 | 0.608 | T | 0.009 | 0.002 |
| 0.764 | T | 0.374 | 0.733 | T | 0.43 | 0.574 |
| 0.595 | T | 0.266 | 0.637 | T | 0.226 | 0.394 |
| 0.032 | T | 0.071 | 0.288 | T | 0.38 | 0.542 |
| 0.871 | D | 0.64 | 0.874 | D | 0.794 | 0.789 |
| 0.535 | T | 0.221 | 0.584 | T | 0.476 | 0.601 |
| 0.727 | T | 0.403 | 0.754 | T | 0.009 | 0.002 |
| 0.713 | T | 0.255 | 0.625 | T | 0.398 | 0.554 |

| | | | | | | |
|---|---|---|---|---|---|---|
| 0.293 | T | 0.166 | 0.504 | T | 0.485 | 0.606 |
| . | . | . | . | . | . | . |
| 0.42 | T | 0.171 | 0.513 | T | 0.022 | 0.005 |
| 0.822 | T | 0.485 | 0.804 | T | 0.817 | 0.809 |
| 0.841 | D | 0.51 | 0.816 | D | 0.72 | 0.736 |
| 0.424 | T | 0.131 | 0.441 | T | 0.737 | 0.747 |
| 0.541 | T | 0.211 | 0.572 | T | 0.338 | 0.509 |
| 0.675 | T | 0.24 | 0.608 | T | 0.009 | 0.002 |
| 0.725 | T | 0.455 | 0.787 | T | 0.699 | 0.723 |
| 0.994 | D | 0.968 | 0.99 | D | 0.877 | 0.87 |
| . | . | . | . | . | . | . |
| 0.886 | D | 0.639 | 0.874 | D | 0.005 | 0.001 |
| 0.868 | D | 0.55 | 0.835 | D | 0.881 | 0.874 |
| . | . | . | . | . | . | . |
| . | . | . | . | . | . | . |
| 0.942 | D | 0.838 | 0.946 | D | 0.083 | 0.138 |
| . | . | . | . | . | . | . |
| 0.835 | D | 0.584 | 0.851 | D | 0.516 | 0.624 |
| 0.451 | T | 0.165 | 0.502 | T | 0.229 | 0.399 |
| . | . | . | . | . | . | . |
| 0.822 | T | 0.514 | 0.818 | D | 0.767 | 0.768 |
| 0.402 | T | 0.115 | 0.408 | T | 0.464 | 0.594 |
| . | . | . | . | . | . | . |
| 0.675 | T | 0.24 | 0.608 | T | 0.009 | 0.002 |
| 0.009 | T | 0.074 | 0.3 | T | 0.06 | 0.073 |
| 0.627 | T | 0.268 | 0.639 | T | 0.241 | 0.412 |
| 0.595 | T | 0.227 | 0.591 | T | 0.021 | 0.005 |
| . | . | . | . | . | . | . |
| 0.117 | T | 0.086 | 0.334 | T | 0.05 | 0.047 |
| 0.917 | D | 0.744 | 0.913 | D | 0.448 | 0.585 |
| 0.636 | T | 0.281 | 0.653 | T | 0.371 | 0.535 |
| 0.161 | T | 0.12 | 0.419 | T | 0.03 | 0.011 |
| 0.664 | T | 0.243 | 0.612 | T | 0.07 | 0.101 |
| 0.537 | T | 0.312 | 0.682 | T | 0.085 | 0.142 |
| 0.501 | T | 0.14 | 0.46 | T | 0.082 | 0.134 |
| 0.794 | T | 0.438 | 0.777 | T | 0.5 | 0.615 |
| 0.646 | T | 0.318 | 0.688 | T | 0.705 | 0.727 |
| 0.66 | T | 0.279 | 0.65 | T | 0.81 | 0.803 |
| 0.936 | D | 0.808 | 0.935 | D | 0.775 | 0.774 |
| 0.832 | D | 0.487 | 0.805 | T | 0.082 | 0.134 |
| . | . | . | . | . | . | . |
| . | . | . | . | . | . | . |
| 0.636 | T | 0.281 | 0.653 | T | 0.371 | 0.535 |

| | | | | | | |
|---|---|---|---|---|---|---|
| . | . | . | . | . | . | . |
| 0.675 | T | 0.24 | 0.608 | T | 0.009 | 0.002 |
| 0.501 | T | 0.14 | 0.46 | T | 0.082 | 0.134 |
| 0.328 | T | 0.11 | 0.395 | T | 0.68 | 0.712 |
| 0.613 | T | 0.29 | 0.661 | T | 0.212 | 0.377 |
| 0.757 | T | 0.421 | 0.766 | T | 0.801 | 0.795 |
| 0.915 | D | 0.746 | 0.913 | D | 0.164 | 0.307 |
| 0.888 | D | 0.575 | 0.847 | D | 0.227 | 0.395 |
| 0.829 | D | 0.492 | 0.807 | T | 0.813 | 0.806 |
| . | . | . | . | . | . | . |
| . | . | . | . | . | . | . |
| 0.668 | T | 0.309 | 0.679 | T | 0.691 | 0.718 |
| 0.841 | D | 0.65 | 0.878 | D | 0.039 | 0.024 |
| . | . | . | . | . | . | . |
| 0.489 | T | 0.157 | 0.49 | T | 0.052 | 0.051 |
| 0.119 | T | 0.075 | 0.301 | T | 0.3 | 0.475 |
| 0.985 | D | 0.983 | 0.995 | D | 0.682 | 0.713 |
| . | . | . | . | . | . | . |
| 0.595 | T | 0.227 | 0.591 | T | 0.021 | 0.005 |
| 0.02 | T | 0.045 | 0.193 | T | 0.135 | 0.257 |
| 0.995 | D | 0.972 | 0.991 | D | 0.701 | 0.724 |
| 0.915 | D | 0.746 | 0.913 | D | 0.164 | 0.307 |
| . | . | . | . | . | . | . |
| 0.73 | T | 0.327 | 0.695 | T | 0.285 | 0.461 |
| 0.583 | T | 0.166 | 0.504 | T | 0.5 | 0.615 |
| 0.606 | T | 0.233 | 0.599 | T | 0.017 | 0.004 |
| 0.021 | T | 0.023 | 0.099 | T | 0.039 | 0.023 |
| 0.496 | T | 0.083 | 0.325 | T | 0.093 | 0.165 |
| 0.577 | T | 0.277 | 0.648 | T | 0.093 | 0.164 |
| 0.538 | T | 0.197 | 0.551 | T | 0.471 | 0.599 |
| 0.68 | T | 0.237 | 0.604 | T | 0.01 | 0.002 |
| 0.915 | D | 0.746 | 0.913 | D | 0.164 | 0.307 |
| . | . | . | . | . | . | . |
| 0.759 | T | 0.424 | 0.768 | T | 0.458 | 0.591 |
| 0.545 | T | 0.167 | 0.506 | T | 0.007 | 0.002 |
| 0.622 | T | 0.241 | 0.609 | T | 0.314 | 0.488 |
| 0.846 | D | 0.537 | 0.829 | D | 0.022 | 0.006 |
| 0.478 | T | 0.146 | 0.471 | T | 0.125 | 0.238 |
| . | . | . | . | . | . | . |
| 0.68 | T | 0.237 | 0.604 | T | 0.01 | 0.002 |
| 0.898 | D | 0.65 | 0.878 | D | 0.891 | 0.885 |
| 0.007 | T | 0.062 | 0.257 | T | 0.045 | 0.035 |
| 0.731 | T | 0.379 | 0.736 | T | 0.587 | 0.661 |

| | | | | | | |
|---|---|---|---|---|---|---|
| 0.752 | T | 0.501 | 0.811 | D | 0.259 | 0.434 |
| 0.725 | T | 0.344 | 0.709 | T | 0.883 | 0.876 |
| . | . | . | . | . | . | . |
| . | . | . | . | . | . | . |
| 0.021 | T | 0.023 | 0.099 | T | 0.039 | 0.023 |
| 0.461 | T | 0.183 | 0.531 | T | 0.531 | 0.632 |
| 0.842 | D | 0.523 | 0.822 | D | 0.45 | 0.586 |
| 0.995 | D | 0.972 | 0.991 | D | 0.701 | 0.724 |
| 0 | T | 0.046 | 0.196 | T | 0.342 | 0.512 |
| . | . | . | . | . | . | . |
| 0.698 | T | 0.378 | 0.736 | T | 0.081 | 0.133 |
| 0.658 | T | 0.241 | 0.61 | T | 0.037 | 0.021 |
| 0.79 | T | 0.446 | 0.782 | T | 0.784 | 0.781 |
| 0.79 | T | 0.446 | 0.782 | T | 0.784 | 0.781 |
| . | . | . | . | . | . | . |
| . | . | . | . | . | . | . |
| 0.869 | D | 0.583 | 0.85 | D | 0.832 | 0.824 |
| 0.858 | D | 0.618 | 0.865 | D | 0.202 | 0.363 |
| 0.764 | T | 0.403 | 0.754 | T | 0.763 | 0.765 |
| . | . | . | . | . | . | . |
| 0.73 | T | 0.327 | 0.695 | T | 0.285 | 0.461 |
| 0.784 | T | 0.491 | 0.807 | T | 0.27 | 0.445 |
| 0.767 | T | 0.389 | 0.744 | T | 0.075 | 0.115 |
| 0.044 | T | 0.072 | 0.293 | T | 0.382 | 0.542 |
| 0.639 | T | 0.248 | 0.618 | T | 0.208 | 0.371 |
| 0.135 | T | 0.139 | 0.457 | T | 0.318 | 0.492 |
| . | . | . | . | . | . | . |
| 0.698 | T | 0.378 | 0.736 | T | 0.081 | 0.133 |
| 0.676 | T | 0.225 | 0.589 | T | 0.015 | 0.003 |
| 0.675 | T | 0.24 | 0.608 | T | 0.009 | 0.002 |
| 0.68 | T | 0.237 | 0.604 | T | 0.01 | 0.002 |
| 0.658 | T | 0.241 | 0.61 | T | 0.037 | 0.021 |
| 0.68 | T | 0.237 | 0.604 | T | 0.01 | 0.002 |
| 0.501 | T | 0.14 | 0.46 | T | 0.082 | 0.134 |
| 0.117 | T | 0.081 | 0.319 | T | 0.443 | 0.582 |
| 0.785 | T | 0.499 | 0.811 | T | 0.219 | 0.385 |
| 0.644 | T | 0.258 | 0.629 | T | 0.069 | 0.098 |
| 0.767 | T | 0.389 | 0.744 | T | 0.075 | 0.115 |
| 0.233 | T | 0.103 | 0.38 | T | 0.013 | 0.003 |
| 0.836 | D | 0.413 | 0.761 | T | 0.859 | 0.851 |
| . | . | . | . | . | . | . |
| 0.611 | T | 0.243 | 0.612 | T | 0.703 | 0.725 |
| 0.545 | T | 0.167 | 0.506 | T | 0.007 | 0.002 |

| | | | | | | |
|---|---|---|---|---|---|---|
| . | . | . | . | . | . | . |
| 0.68 | T | 0.237 | 0.604 | T | 0.01 | 0.002 |
| 0.767 | T | 0.389 | 0.744 | T | 0.075 | 0.115 |
| 0.545 | T | 0.167 | 0.506 | T | 0.007 | 0.002 |
| 0.021 | T | 0.023 | 0.099 | T | 0.039 | 0.023 |
| 0.545 | T | 0.167 | 0.506 | T | 0.007 | 0.002 |
| 0.051 | T | 0.026 | 0.111 | T | 0.11 | 0.206 |
| 0.837 | D | 0.562 | 0.841 | D | 0.324 | 0.497 |
| . | . | . | . | . | . | . |
| 0.846 | D | 0.537 | 0.829 | D | 0.022 | 0.006 |
| 0.644 | T | 0.258 | 0.629 | T | 0.069 | 0.098 |
| 0.405 | T | 0.101 | 0.374 | T | 0.005 | 0.001 |
| . | . | . | . | . | . | . |
| 0.909 | D | 0.75 | 0.915 | D | 0.238 | 0.41 |
| 0.759 | T | 0.424 | 0.768 | T | 0.458 | 0.591 |
| 0.692 | T | 0.284 | 0.656 | T | 0.67 | 0.707 |
| 0.237 | T | 0.111 | 0.398 | T | 0.037 | 0.02 |
| . | . | . | . | . | . | . |
| 0.95 | D | 0.87 | 0.957 | D | 0.257 | 0.431 |
| 0.837 | D | 0.524 | 0.823 | D | 0.315 | 0.489 |
| 0.237 | T | 0.111 | 0.398 | T | 0.037 | 0.02 |
| 0.828 | D | 0.463 | 0.792 | T | 0.015 | 0.003 |
| . | . | . | . | . | . | . |
| 0.025 | T | 0.058 | 0.244 | T | 0.788 | 0.784 |
| . | . | . | . | . | . | . |
| 0.838 | D | 0.491 | 0.807 | T | 0.209 | 0.373 |
| 0.592 | T | 0.188 | 0.539 | T | 0.187 | 0.343 |
| . | . | . | . | . | . | . |
| . | . | . | . | . | . | . |
| 0.762 | T | 0.427 | 0.77 | T | 0.386 | 0.545 |
| 0.871 | D | 0.605 | 0.86 | D | 0.902 | 0.895 |
| 0.767 | T | 0.39 | 0.744 | T | 0.649 | 0.695 |
| 0.938 | D | 0.785 | 0.927 | D | 0.824 | 0.815 |
| 0.001 | T | 0.046 | 0.196 | T | 0.67 | 0.707 |
| . | . | . | . | . | . | . |
| 0.794 | T | 0.623 | 0.867 | D | 0.018 | 0.004 |
| 0.767 | T | 0.39 | 0.744 | T | 0.649 | 0.695 |
| 0.881 | D | 0.453 | 0.786 | T | 0.872 | 0.865 |
| 0.012 | T | 0.082 | 0.324 | T | 0.118 | 0.223 |
| 0.059 | T | 0.06 | 0.251 | T | 0.007 | 0.002 |
| 0.9 | D | 0.697 | 0.896 | D | 0.789 | 0.785 |
| 0.975 | D | 0.922 | 0.974 | D | 0.776 | 0.775 |
| 0.613 | T | 0.225 | 0.589 | T | 0.738 | 0.748 |

| | | | | | | |
|---|---|---|---|---|---|---|
| . | . | . | . | . | . | . |
| 0.838 | D | 0.491 | 0.807 | T | 0.209 | 0.373 |
| 0.652 | T | 0.296 | 0.668 | T | 0.216 | 0.381 |
| 0.854 | D | 0.64 | 0.874 | D | 0.323 | 0.496 |
| 0.638 | T | 0.23 | 0.595 | T | 0.173 | 0.322 |
| 0.595 | T | 0.181 | 0.528 | T | 0.561 | 0.648 |
| 0.821 | T | 0.497 | 0.81 | T | 0.769 | 0.769 |
| . | . | . | . | . | . | . |
| 0.231 | T | 0.081 | 0.32 | T | 0.068 | 0.095 |
| 0.981 | D | 0.989 | 0.997 | D | 0.7 | 0.724 |
| 0.841 | D | 0.65 | 0.878 | D | 0.039 | 0.024 |
| 0.254 | T | 0.117 | 0.412 | T | 0.528 | 0.63 |
| 0.961 | D | 0.884 | 0.961 | D | 0.921 | 0.914 |
| 0.301 | T | 0.087 | 0.338 | T | 0.038 | 0.022 |
| 0.924 | D | 0.782 | 0.926 | D | 0.785 | 0.782 |
| 0.614 | T | 0.21 | 0.57 | T | 0.103 | 0.189 |
| 0.539 | T | 0.225 | 0.59 | T | 0.01 | 0.003 |
| 0.778 | T | 0.391 | 0.745 | T | 0.064 | 0.083 |
| 0.16 | T | 0.025 | 0.108 | T | 0.006 | 0.001 |
| 0.539 | T | 0.225 | 0.59 | T | 0.01 | 0.003 |
| 0.632 | T | 0.244 | 0.613 | T | 0.071 | 0.104 |
| 0.688 | T | 0.358 | 0.72 | T | 0.658 | 0.7 |
| 0.828 | D | 0.463 | 0.792 | T | 0.015 | 0.003 |
| 0.539 | T | 0.225 | 0.59 | T | 0.01 | 0.003 |
| 0.395 | T | 0.119 | 0.417 | T | 0.213 | 0.378 |
| . | . | . | . | . | . | . |
| 0.778 | T | 0.391 | 0.745 | T | 0.064 | 0.083 |
| 0.708 | T | 0.33 | 0.698 | T | 0.024 | 0.006 |
| 0.237 | T | 0.111 | 0.398 | T | 0.037 | 0.02 |
| 0.802 | T | 0.415 | 0.762 | T | 0.027 | 0.009 |
| 0.611 | T | 0.249 | 0.619 | T | 0.445 | 0.583 |
| 0.664 | T | 0.271 | 0.643 | T | 0.852 | 0.843 |
| 0.821 | T | 0.497 | 0.81 | T | 0.769 | 0.769 |
| 0.539 | T | 0.225 | 0.59 | T | 0.01 | 0.003 |
| 0.708 | T | 0.33 | 0.698 | T | 0.024 | 0.006 |
| 0.012 | T | 0.082 | 0.324 | T | 0.118 | 0.223 |
| 0.521 | T | 0.193 | 0.546 | T | 0.615 | 0.677 |
| . | . | . | . | . | . | . |
| 0.772 | T | 0.522 | 0.821 | D | 0.642 | 0.691 |
| 0.927 | D | 0.728 | 0.907 | D | 0.983 | 0.984 |
| 0.722 | T | 0.344 | 0.71 | T | 0.036 | 0.018 |
| 0.854 | D | 0.64 | 0.874 | D | 0.323 | 0.496 |
| 0.539 | T | 0.225 | 0.59 | T | 0.01 | 0.003 |

| | | | | | | |
|---|---|---|---|---|---|---|
| 0.003 | T | 0.073 | 0.297 | T | 0.008 | 0.002 |
| 0.708 | T | 0.33 | 0.698 | T | 0.024 | 0.006 |
| 0.006 | T | 0.054 | 0.23 | T | 0.635 | 0.687 |

| MetaRNN_pr | M.CAP_score | M.CAP_ranks | M.CAP_pred | REVEL_score | REVEL_ranks( | MutPred_sco |
|---|---|---|---|---|---|---|
| . | . | . | . | . | . | . |
| . | . | . | . | . | . | . |
| T | 0.019 | 0.407 | T | 0.143 | 0.387 | . |
| D | 0.017 | 0.383 | T | 0.469 | 0.766 | 0.414 |
| T | 0.109 | 0.786 | D | 0.321 | 0.647 | . |
| T | 0.06 | 0.678 | D | 0.291 | 0.614 | 0.136 |
| T | 0.207 | 0.871 | D | 0.693 | 0.89 | . |
| D | 0.051 | 0.646 | D | 0.318 | 0.644 | 0.633 |
| T | 0.23 | 0.882 | D | 0.676 | 0.882 | . |
| T | 0.011 | 0.274 | T | 0.198 | 0.485 | . |
| T | 0.023 | 0.46 | T | 0.233 | 0.539 | . |
| T | 0.019 | 0.407 | T | 0.143 | 0.387 | . |
| T | 0.081 | 0.736 | D | 0.317 | 0.643 | 0.588 |
| T | 0.017 | 0.381 | T | 0.496 | 0.783 | . |
| . | . | . | . | . | . | . |
| T | . | . | . | 0.379 | 0.7 | 0.197 |
| T | 0.46 | 0.945 | D | 0.32 | 0.646 | 0.314 |
| D | 0.082 | 0.739 | D | 0.343 | 0.668 | 0.518 |
| T | 0.023 | 0.46 | T | 0.233 | 0.539 | . |
| T | 0.061 | 0.683 | D | 0.279 | 0.6 | 0.456 |
| D | 0.102 | 0.776 | D | 0.685 | 0.886 | . |
| . | . | . | . | . | . | . |
| D | 0.037 | 0.576 | D | 0.51 | 0.792 | 0.515 |
| D | 0.047 | 0.626 | D | 0.205 | 0.497 | . |
| . | . | . | . | . | . | . |
| D | 0.163 | 0.842 | D | 0.221 | 0.521 | . |
| D | 0.924 | 0.994 | D | 0.826 | 0.945 | . |
| T | 0.012 | 0.303 | T | 0.106 | 0.306 | . |
| T | 0.764 | 0.981 | D | 0.295 | 0.619 | 0.291 |
| D | 0.249 | 0.891 | D | 0.465 | 0.764 | . |
| T | 0.023 | 0.46 | T | 0.233 | 0.539 | . |
| T | 0.419 | 0.938 | D | 0.574 | 0.829 | . |
| T | 0.026 | 0.487 | D | 0.257 | 0.572 | 0.439 |
| T | 0.011 | 0.274 | T | 0.198 | 0.485 | . |
| T | 0.015 | 0.348 | T | 0.085 | 0.252 | 0.386 |
| T | 0.009 | 0.235 | T | 0.198 | 0.485 | . |
| T | 0.023 | 0.458 | T | 0.164 | 0.427 | 0.448 |
| D | 0.031 | 0.529 | D | 0.263 | 0.58 | 0.56 |
| T | 0.198 | 0.865 | D | 0.559 | 0.821 | . |
| D | 0.34 | 0.92 | D | 0.597 | 0.842 | 0.415 |
| T | 0.028 | 0.507 | D | 0.114 | 0.325 | . |
| T | 0.027 | 0.501 | D | 0.273 | 0.593 | . |

| | | | | | | |
|---|---|---|---|---|---|---|
| T | 0.019 | 0.407 | T | 0.143 | 0.387 | . |
| T | 0.069 | 0.706 | D | 0.116 | 0.329 | 0.277 |
| T | 0.027 | 0.499 | D | 0.137 | 0.374 | . |
| D | 0.186 | 0.858 | D | 0.64 | 0.865 | 0.553 |
| D | 0.102 | 0.776 | D | 0.685 | 0.886 | . |
| . | . | . | . | . | . | . |
| . | . | . | . | . | . | . |
| T | 0.01 | 0.268 | T | 0.181 | 0.457 | . |
| D | 0.068 | 0.702 | D | 0.702 | 0.894 | . |
| T | 0.019 | 0.415 | T | 0.139 | 0.379 | . |
| T | 0.061 | 0.682 | D | 0.403 | 0.719 | 0.671 |
| T | 0.016 | 0.375 | T | 0.27 | 0.589 | . |
| T | 0.04 | 0.593 | D | 0.1 | 0.291 | . |
| . | . | . | . | . | . | . |
| T | 0.036 | 0.565 | D | 0.061 | 0.18 | . |
| D | 0.008 | 0.209 | T | 0.279 | 0.6 | 0.375 |
| D | 0.306 | 0.91 | D | 0.932 | 0.984 | 0.453 |
| T | 0.003 | 0.066 | T | 0.125 | 0.349 | . |
| T | 0.022 | 0.447 | T | 0.191 | 0.474 | 0.185 |
| T | 0.033 | 0.548 | D | 0.307 | 0.632 | . |
| T | 0.112 | 0.789 | D | 0.351 | 0.676 | . |
| . | . | . | . | . | . | . |
| T | 0.019 | 0.407 | T | 0.143 | 0.387 | . |
| T | 0.087 | 0.749 | D | 0.523 | 0.8 | . |
| T | 0.023 | 0.464 | T | 0.242 | 0.552 | . |
| T | 0.023 | 0.46 | T | 0.233 | 0.539 | . |
| T | 0.096 | 0.765 | D | 0.283 | 0.605 | 0.197 |
| D | 0.017 | 0.39 | T | 0.087 | 0.257 | 0.51 |
| T | 0.031 | 0.532 | D | 0.358 | 0.682 | . |
| . | . | . | . | . | . | . |
| T | 0.166 | 0.844 | D | 0.602 | 0.845 | 0.352 |
| D | 0.012 | 0.293 | T | 0.211 | 0.506 | . |
| D | 0.107 | 0.783 | D | 0.548 | 0.815 | 0.375 |
| T | 0.23 | 0.882 | D | 0.676 | 0.882 | . |
| T | 0.019 | 0.407 | T | 0.143 | 0.387 | . |
| T | 0.019 | 0.407 | T | 0.143 | 0.387 | . |
| T | 0.038 | 0.578 | D | 0.409 | 0.724 | 0.196 |
| T | 0.042 | 0.604 | D | 0.145 | 0.391 | . |
| T | 0.02 | 0.421 | T | 0.127 | 0.354 | 0.612 |
| D | 0.071 | 0.712 | D | 0.651 | 0.87 | 0.402 |
| T | 0.022 | 0.453 | T | 0.357 | 0.681 | . |
| T | . | . | . | 0.379 | 0.7 | 0.197 |
| T | 0.085 | 0.743 | D | 0.417 | 0.73 | . |

| | | | | | | | | |
|---|---|---|---|---|---|---|---|---|
| T | | 0.029 | 0.519 | D | | 0.137 | 0.374 | . | |
| . | . | . | . | . | . | . | . | |
| T | | 0.047 | 0.625 | D | | 0.275 | 0.595 | . | |
| D | | 0.081 | 0.736 | D | | 0.443 | 0.749 | | 0.514 |
| D | | 0.068 | 0.702 | D | | 0.702 | 0.894 | . | |
| D | | 0.045 | 0.616 | D | | 0.346 | 0.671 | | 0.229 |
| T | | 0.026 | 0.489 | D | | 0.089 | 0.263 | | 0.406 |
| T | | 0.019 | 0.407 | T | | 0.143 | 0.387 | . | |
| D | | 0.042 | 0.603 | D | | 0.401 | 0.718 | . | |
| D | | 0.665 | 0.972 | D | | 0.9 | 0.973 | | 0.194 |
| . | . | . | . | . | . | . | . | |
| T | | 0.109 | 0.786 | D | | 0.321 | 0.647 | . | |
| D | | 0.102 | 0.776 | D | | 0.673 | 0.881 | | 0.738 |
| . | . | . | . | . | . | . | . | |
| . | . | . | . | . | . | . | . | |
| T | | 0.23 | 0.882 | D | | 0.676 | 0.882 | . | |
| . | . | . | . | . | . | . | . | |
| D | | 0.071 | 0.712 | D | | 0.411 | 0.726 | | 0.102 |
| T | | 0.058 | 0.671 | D | | 0.37 | 0.693 | . | |
| . | . | . | . | . | . | . | . | |
| D | | 0.07 | 0.709 | D | | 0.557 | 0.82 | | 0.361 |
| T | | 0.016 | 0.363 | T | | 0.148 | 0.396 | | 0.172 |
| . | . | . | . | . | . | . | . | |
| T | | 0.019 | 0.407 | T | | 0.143 | 0.387 | . | |
| T | | 0.023 | 0.46 | T | | 0.233 | 0.539 | . | |
| T | | 0.015 | 0.356 | T | | 0.277 | 0.598 | . | |
| T | | 0.011 | 0.274 | T | | 0.198 | 0.485 | . | |
| . | . | . | . | . | . | . | . | |
| T | | 0.233 | 0.883 | D | | 0.078 | 0.232 | . | |
| T | | 0.152 | 0.834 | D | | 0.631 | 0.86 | | 0.329 |
| T | | 0.034 | 0.556 | D | | 0.358 | 0.682 | . | |
| T | | 0.009 | 0.241 | T | | 0.239 | 0.548 | . | |
| T | | 0.019 | 0.416 | T | | 0.325 | 0.651 | . | |
| T | | 0.073 | 0.717 | D | | 0.189 | 0.471 | | 0.322 |
| T | | 0.033 | 0.548 | D | | 0.307 | 0.632 | . | |
| T | | 0.078 | 0.729 | D | | 0.374 | 0.696 | | 0.473 |
| D | | 0.07 | 0.708 | D | | 0.349 | 0.674 | | 0.394 |
| D | | 0.03 | 0.527 | D | | 0.413 | 0.727 | | 0.748 |
| D | | 0.155 | 0.836 | D | | 0.508 | 0.791 | | 0.188 |
| T | | 0.05 | 0.641 | D | | 0.735 | 0.909 | . | |
| . | . | . | . | . | . | . | . | |
| . | . | . | . | . | . | . | . | |
| T | | 0.034 | 0.556 | D | | 0.358 | 0.682 | . | |

| | | | | | | |
|---|---|---|---|---|---|---|
| . | . | . | . | . | . | . |
| T | 0.019 | 0.407 | T | 0.143 | 0.387 | . |
| T | 0.033 | 0.548 | D | 0.307 | 0.632 | . |
| D | 0.017 | 0.384 | T | 0.171 | 0.439 | 0.487 |
| T | 0.049 | 0.638 | D | 0.135 | 0.37 | . |
| D | 0.138 | 0.82 | D | 0.496 | 0.783 | 0.285 |
| T | 0.183 | 0.856 | D | 0.72 | 0.902 | . |
| T | 0.173 | 0.85 | D | 0.38 | 0.701 | . |
| D | 0.166 | 0.845 | D | 0.637 | 0.863 | 0.487 |
| . | . | . | . | . | . | . |
| . | . | . | . | . | . | . |
| D | 0.028 | 0.505 | D | 0.276 | 0.596 | 0.282 |
| T | 0.036 | 0.568 | D | 0.363 | 0.686 | 0.092 |
| . | . | . | . | . | . | . |
| T | 0.02 | 0.423 | T | 0.103 | 0.299 | . |
| T | 0.03 | 0.524 | D | 0.177 | 0.45 | 0.122 |
| D | 0.405 | 0.935 | D | 0.616 | 0.852 | 0.205 |
| . | . | . | . | . | . | . |
| T | 0.011 | 0.274 | T | 0.198 | 0.485 | . |
| T | 0.008 | 0.213 | T | 0.064 | 0.19 | 0.237 |
| D | 0.283 | 0.903 | D | 0.847 | 0.953 | 0.729 |
| T | 0.183 | 0.856 | D | 0.72 | 0.902 | . |
| . | . | . | . | . | . | . |
| T | 0.016 | 0.367 | T | 0.239 | 0.548 | 0.244 |
| D | 0.069 | 0.705 | D | 0.261 | 0.577 | 0.526 |
| T | 0.032 | 0.538 | D | 0.051 | 0.147 | . |
| T | 0.008 | 0.207 | T | 0.1 | 0.291 | 0.197 |
| T | 0.025 | 0.481 | D | 0.057 | 0.167 | . |
| T | 0.014 | 0.339 | T | 0.201 | 0.49 | . |
| T | 0.016 | 0.371 | T | 0.361 | 0.685 | . |
| T | 0.031 | 0.532 | D | 0.358 | 0.682 | . |
| T | 0.183 | 0.856 | D | 0.72 | 0.902 | . |
| . | . | . | . | . | . | . |
| T | 0.141 | 0.823 | D | 0.352 | 0.677 | . |
| T | 0.028 | 0.505 | D | 0.195 | 0.481 | . |
| T | 0.023 | 0.459 | T | 0.191 | 0.474 | 0.155 |
| T | 0.132 | 0.814 | D | 0.655 | 0.872 | . |
| T | 0.03 | 0.524 | D | 0.363 | 0.686 | . |
| . | . | . | . | . | . | . |
| T | 0.031 | 0.532 | D | 0.358 | 0.682 | . |
| D | 0.426 | 0.939 | D | 0.825 | 0.945 | 0.536 |
| T | 0.022 | 0.447 | T | 0.335 | 0.661 | . |
| D | 0.019 | 0.406 | T | 0.343 | 0.668 | . |

| | | | | | | |
|---|---|---|---|---|---|---|
| T | 0.056 | 0.664 | D | 0.249 | 0.562 | 0.341 |
| D | 0.05 | 0.639 | D | 0.676 | 0.882 | . |
| . | . | . | . | . | . | . |
| . | . | . | . | . | . | . |
| T | 0.008 | 0.207 | T | 0.1 | 0.291 | 0.197 |
| D | 0.028 | 0.507 | D | 0.219 | 0.518 | 0.355 |
| T | 0.137 | 0.819 | D | 0.523 | 0.8 | . |
| D | 0.283 | 0.903 | D | 0.847 | 0.953 | 0.729 |
| T | 0.027 | 0.503 | D | 0.265 | 0.583 | . |
| . | . | . | . | . | . | . |
| T | 0.05 | 0.64 | D | 0.216 | 0.514 | 0.711 |
| T | 0.019 | 0.409 | T | 0.243 | 0.553 | . |
| D | 0.143 | 0.826 | D | 0.467 | 0.765 | 0.623 |
| D | 0.143 | 0.826 | D | 0.467 | 0.765 | 0.623 |
| . | . | . | . | . | . | . |
| . | . | . | . | . | . | . |
| D | 0.157 | 0.838 | D | 0.818 | 0.942 | 0.539 |
| T | 0.052 | 0.648 | D | 0.386 | 0.706 | 0.214 |
| D | 0.034 | 0.556 | D | 0.419 | 0.731 | 0.65 |
| . | . | . | . | . | . | . |
| T | 0.016 | 0.367 | T | 0.239 | 0.548 | 0.244 |
| T | 0.077 | 0.727 | D | 0.415 | 0.729 | . |
| T | 0.017 | 0.381 | T | 0.496 | 0.783 | . |
| T | 0.535 | 0.956 | D | 0.162 | 0.423 | 0.317 |
| T | 0.024 | 0.467 | T | 0.522 | 0.799 | 0.259 |
| T | 0.02 | 0.428 | T | 0.204 | 0.495 | . |
| . | . | . | . | . | . | . |
| T | 0.05 | 0.64 | D | 0.216 | 0.514 | 0.711 |
| T | 0.013 | 0.328 | T | 0.318 | 0.644 | . |
| T | 0.019 | 0.407 | T | 0.143 | 0.387 | . |
| T | 0.031 | 0.532 | D | 0.358 | 0.682 | . |
| T | 0.019 | 0.409 | T | 0.243 | 0.553 | . |
| T | 0.031 | 0.532 | D | 0.358 | 0.682 | . |
| T | 0.033 | 0.548 | D | 0.307 | 0.632 | . |
| T | 0.025 | 0.484 | D | 0.214 | 0.511 | 0.211 |
| T | 0.485 | 0.949 | D | 0.349 | 0.674 | 0.319 |
| T | 0.028 | 0.507 | D | 0.114 | 0.325 | . |
| T | 0.017 | 0.381 | T | 0.496 | 0.783 | . |
| T | 0.027 | 0.499 | D | 0.137 | 0.374 | . |
| D | 0.212 | 0.873 | D | 0.667 | 0.878 | 0.662 |
| . | . | . | . | . | . | . |
| D | 0.037 | 0.57 | D | 0.186 | 0.466 | . |
| T | 0.028 | 0.505 | D | 0.195 | 0.481 | . |

| | | | | | | |
|---|---|---|---|---|---|---|
| . | . | . | . | . | . | . |
| T | 0.031 | 0.532 | D | 0.358 | 0.682 | . |
| T | 0.017 | 0.381 | T | 0.496 | 0.783 | . |
| T | 0.028 | 0.505 | D | 0.195 | 0.481 | . |
| T | 0.008 | 0.207 | T | 0.1 | 0.291 | 0.197 |
| T | 0.028 | 0.505 | D | 0.195 | 0.481 | . |
| T | 0.011 | 0.276 | T | 0.04 | 0.108 | 0.207 |
| T | 0.03 | 0.526 | D | 0.129 | 0.358 | 0.18 |
| . | . | . | . | . | . | . |
| T | 0.132 | 0.814 | D | 0.655 | 0.872 | . |
| T | 0.028 | 0.507 | D | 0.114 | 0.325 | . |
| T | 0.016 | 0.376 | T | 0.067 | 0.199 | . |
| . | . | . | . | . | . | . |
| T | 0.217 | 0.876 | D | 0.316 | 0.642 | 0.261 |
| T | 0.141 | 0.823 | D | 0.352 | 0.677 | . |
| D | 0.078 | 0.73 | D | 0.323 | 0.649 | 0.637 |
| T | 0.032 | 0.539 | D | 0.312 | 0.637 | . |
| . | . | . | . | . | . | . |
| T | 0.214 | 0.874 | D | 0.678 | 0.883 | 0.425 |
| T | 0.048 | 0.63 | D | 0.289 | 0.612 | 0.127 |
| T | 0.032 | 0.539 | D | 0.312 | 0.637 | . |
| T | 0.181 | 0.855 | D | 0.522 | 0.799 | . |
| . | . | . | . | . | . | . |
| D | 0.042 | 0.6 | D | 0.457 | 0.758 | 0.616 |
| . | . | . | . | . | . | . |
| T | 0.067 | 0.699 | D | 0.605 | 0.846 | 0.792 |
| T | 0.016 | 0.374 | T | 0.097 | 0.284 | 0.26 |
| . | . | . | . | . | . | . |
| . | . | . | . | . | . | . |
| T | 0.069 | 0.706 | D | 0.335 | 0.661 | 0.303 |
| D | 0.129 | 0.811 | D | 0.621 | 0.855 | 0.744 |
| D | 0.083 | 0.74 | D | 0.438 | 0.745 | 0.469 |
| D | 0.522 | 0.955 | D | 0.788 | 0.931 | 0.55 |
| D | 0.008 | 0.215 | T | 0.402 | 0.719 | 0.322 |
| . | . | . | . | . | . | . |
| T | 0.087 | 0.748 | D | 0.265 | 0.583 | . |
| D | 0.083 | 0.74 | D | 0.438 | 0.745 | 0.469 |
| D | 0.207 | 0.871 | D | 0.602 | 0.845 | 0.58 |
| T | 0.01 | 0.252 | T | 0.127 | 0.354 | . |
| T | 0.017 | 0.391 | T | 0.037 | 0.097 | . |
| D | 0.316 | 0.913 | D | 0.645 | 0.867 | 0.525 |
| D | 0.395 | 0.933 | D | 0.78 | 0.928 | . |
| D | 0.072 | 0.714 | D | 0.367 | 0.69 | 0.317 |

| | | | | | | |
|---|---|---|---|---|---|---|
| . | . | . | . | . | . | . |
| T | 0.067 | 0.699 | D | 0.605 | 0.846 | 0.792 |
| T | 0.057 | 0.67 | D | 0.494 | 0.782 | . |
| T | 0.1 | 0.772 | D | 0.577 | 0.831 | 0.326 |
| T | 0.048 | 0.634 | D | 0.204 | 0.495 | . |
| D | 0.023 | 0.463 | T | 0.321 | 0.647 | 0.386 |
| D | 0.068 | 0.703 | D | 0.521 | 0.799 | 0.612 |
| . | . | . | . | . | . | . |
| T | 0.025 | 0.48 | D | 0.082 | 0.243 | 0.369 |
| D | 0.731 | 0.978 | D | 0.442 | 0.748 | 0.166 |
| T | 0.036 | 0.568 | D | 0.363 | 0.686 | 0.092 |
| D | 0.018 | 0.398 | T | 0.201 | 0.49 | 0.605 |
| D | 0.373 | 0.928 | D | 0.913 | 0.977 | 0.764 |
| T | 0.057 | 0.669 | D | 0.292 | 0.615 | . |
| D | 0.223 | 0.879 | D | 0.593 | 0.84 | 0.18 |
| T | 0.009 | 0.226 | T | 0.234 | 0.541 | 0.161 |
| T | 0.024 | 0.472 | T | 0.256 | 0.571 | 0.552 |
| T | 0.494 | 0.95 | D | 0.499 | 0.785 | . |
| T | 0.002 | 0.044 | T | 0.083 | 0.246 | . |
| T | 0.024 | 0.472 | T | 0.256 | 0.571 | 0.552 |
| T | 0.017 | 0.392 | T | 0.295 | 0.619 | 0.153 |
| D | 0.025 | 0.478 | T | 0.356 | 0.68 | 0.433 |
| T | 0.181 | 0.855 | D | 0.522 | 0.799 | . |
| T | 0.024 | 0.472 | T | 0.256 | 0.571 | 0.552 |
| T | 0.041 | 0.597 | D | 0.177 | 0.45 | 0.546 |
| . | . | . | . | . | . | . |
| T | 0.494 | 0.95 | D | 0.499 | 0.785 | . |
| T | 0.025 | 0.479 | T | 0.215 | 0.512 | . |
| T | 0.032 | 0.539 | D | 0.312 | 0.637 | . |
| T | 0.095 | 0.764 | D | 0.23 | 0.535 | . |
| T | 0.21 | 0.872 | D | 0.287 | 0.61 | 0.485 |
| D | 0.032 | 0.539 | D | 0.541 | 0.811 | 0.523 |
| D | 0.068 | 0.703 | D | 0.521 | 0.799 | 0.612 |
| T | 0.024 | 0.472 | T | 0.256 | 0.571 | 0.552 |
| T | 0.025 | 0.479 | T | 0.215 | 0.512 | . |
| T | 0.01 | 0.252 | T | 0.127 | 0.354 | . |
| D | 0.073 | 0.717 | D | 0.275 | 0.595 | 0.692 |
| . | . | . | . | . | . | . |
| D | 0.084 | 0.742 | D | 0.515 | 0.795 | 0.252 |
| D | 0.229 | 0.882 | D | 0.81 | 0.939 | 0.923 |
| T | 0.048 | 0.631 | D | 0.516 | 0.796 | . |
| T | 0.1 | 0.772 | D | 0.577 | 0.831 | 0.326 |
| T | 0.024 | 0.472 | T | 0.256 | 0.571 | 0.552 |

| | | | | | | | | |
|---|---|---|---|---|---|---|---|---|
| T | . | . | . | | 0.124 | 0.347 | . | |
| T | | 0.025 | 0.479 | T | 0.215 | 0.512 | . | |
| D | | 0.037 | 0.573 | D | 0.261 | 0.577 | | 0.54 |

| MutPred_ran | MVP_score | MVP_ranksco | MPC_score | MPC_ranksco | PrimateAI_sc | PrimateAI_ra |
|---|---|---|---|---|---|---|
| . | . | . | . | . | . | . |
| . | . | . | . | . | . | . |
| . | 0.406 | 0.402 | . | . | 0.532 | 0.433 |
| 0.452 | 0.321 | 0.317 | 1.119 | 0.782 | 0.762 | 0.763 |
| . | 0.788 | 0.786 | 1.409 | 0.854 | 0.686 | 0.652 |
| 0.04 | 0.575 | 0.571 | 0.903 | 0.707 | 0.873 | 0.93 |
| . | 0.968 | 0.968 | 2.347 | 0.968 | 0.792 | 0.808 |
| 0.769 | 0.66 | 0.658 | 0.563 | 0.527 | 0.595 | 0.522 |
| . | 0.922 | 0.921 | 1.464 | 0.864 | 0.8 | 0.82 |
| . | 0.212 | 0.208 | 1.047 | 0.76 | 0.765 | 0.767 |
| . | 0.36 | 0.356 | 0.681 | 0.6 | 0.609 | 0.542 |
| . | 0.406 | 0.402 | . | . | 0.532 | 0.433 |
| 0.716 | 0.657 | 0.654 | 0.423 | 0.428 | 0.89 | 0.952 |
| . | 0.608 | 0.605 | 0.709 | 0.616 | 0.627 | 0.568 |
| . | . | . | . | . | . | . |
| 0.11 | 0.815 | 0.814 | 1.097 | 0.776 | 0.803 | 0.825 |
| 0.289 | 0.529 | 0.525 | 0.468 | 0.462 | 0.844 | 0.887 |
| 0.619 | 0.783 | 0.781 | . | . | 0.898 | 0.962 |
| . | 0.36 | 0.356 | 0.681 | 0.6 | 0.609 | 0.542 |
| 0.521 | 0.644 | 0.641 | 0.363 | 0.379 | 0.76 | 0.759 |
| . | 0.773 | 0.771 | 1.543 | 0.878 | 0.83 | 0.865 |
| . | . | . | . | . | . | . |
| 0.614 | 0.396 | 0.392 | 0.51 | 0.491 | 0.665 | 0.622 |
| . | 0.487 | 0.483 | 0.787 | 0.656 | 0.721 | 0.702 |
| . | . | . | . | . | . | . |
| . | 0.472 | 0.468 | 0.779 | 0.652 | 0.53 | 0.431 |
| . | 0.664 | 0.662 | 1.391 | 0.85 | 0.728 | 0.712 |
| . | 0.346 | 0.342 | 0.932 | 0.719 | 0.529 | 0.429 |
| 0.252 | 0.844 | 0.842 | . | . | 0.745 | 0.737 |
| . | 0.34 | 0.336 | 0.708 | 0.615 | 0.836 | 0.874 |
| . | 0.36 | 0.356 | 0.681 | 0.6 | 0.609 | 0.542 |
| . | 0.978 | 0.978 | . | . | 0.846 | 0.89 |
| 0.493 | 0.151 | 0.148 | 0.518 | 0.497 | 0.726 | 0.709 |
| . | 0.212 | 0.208 | 1.047 | 0.76 | 0.765 | 0.767 |
| 0.406 | 0.392 | 0.388 | 0.074 | 0.083 | 0.527 | 0.426 |
| . | 0.162 | 0.158 | 0.032 | 0.034 | 0.435 | 0.299 |
| 0.508 | 0.614 | 0.611 | 1.236 | 0.815 | 0.761 | 0.761 |
| 0.679 | 0.577 | 0.574 | 1.682 | 0.9 | 0.868 | 0.922 |
| . | 0.546 | 0.543 | 1.226 | 0.812 | 0.721 | 0.701 |
| 0.454 | 0.84 | 0.839 | 1.136 | 0.788 | 0.742 | 0.733 |
| . | 0.231 | 0.227 | 0.709 | 0.616 | 0.491 | 0.375 |
| . | 0.42 | 0.416 | 1.181 | 0.8 | 0.799 | 0.818 |

| | | | | | | |
|---|---|---|---|---|---|---|
| . | 0.406 | 0.402 | . | . | 0.532 | 0.433 |
| 0.23 | 0.675 | 0.673 | 0.094 | 0.106 | 0.208 | 0.007 |
| . | 0.537 | 0.533 | 0.691 | 0.606 | 0.506 | 0.397 |
| 0.67 | 0.958 | 0.957 | 0.402 | 0.411 | 0.703 | 0.676 |
| . | 0.773 | 0.771 | 1.543 | 0.878 | 0.83 | 0.865 |
| . | . | . | . | . | . | . |
| . | . | . | . | . | . | . |
| . | 0.709 | 0.707 | 0.165 | 0.186 | 0.391 | 0.238 |
| . | 0.867 | 0.866 | 1.351 | 0.841 | 0.675 | 0.636 |
| . | 0.585 | 0.582 | 1.96 | 0.934 | 0.792 | 0.807 |
| 0.809 | 0.543 | 0.539 | 1.475 | 0.866 | 0.862 | 0.914 |
| . | 0.739 | 0.737 | . | . | 0.776 | 0.783 |
| . | 0.505 | 0.501 | 0.426 | 0.43 | 0.37 | 0.209 |
| . | . | . | . | . | . | . |
| . | 0.33 | 0.326 | 0.093 | 0.105 | 0.261 | 0.05 |
| 0.388 | 0.462 | 0.458 | 0.525 | 0.502 | 0.805 | 0.827 |
| 0.516 | 0.821 | 0.819 | 0.903 | 0.707 | 0.882 | 0.942 |
| . | 0.182 | 0.178 | 0.112 | 0.126 | 0.484 | 0.366 |
| 0.094 | 0.534 | 0.531 | 0.502 | 0.486 | 0.406 | 0.259 |
| . | 0.768 | 0.766 | 0.34 | 0.359 | 0.543 | 0.448 |
| . | 0.592 | 0.589 | 0.808 | 0.666 | 0.62 | 0.558 |
| . | . | . | . | . | . | . |
| . | 0.406 | 0.402 | . | . | 0.532 | 0.433 |
| . | 0.661 | 0.658 | 0.7 | 0.611 | 0.721 | 0.702 |
| . | 0.581 | 0.577 | 0.974 | 0.735 | 0.593 | 0.519 |
| . | 0.36 | 0.356 | 0.681 | 0.6 | 0.609 | 0.542 |
| 0.11 | 0.67 | 0.667 | 0.193 | 0.216 | 0.707 | 0.681 |
| 0.607 | 0.533 | 0.53 | . | . | 0.721 | 0.701 |
| . | 0.694 | 0.692 | 0.143 | 0.161 | 0.615 | 0.55 |
| . | . | . | . | . | . | . |
| 0.351 | 0.825 | 0.823 | 0.454 | 0.451 | 0.68 | 0.643 |
| . | 0.828 | 0.827 | 0.823 | 0.673 | 0.759 | 0.758 |
| 0.388 | 0.865 | 0.864 | 0.563 | 0.528 | 0.703 | 0.676 |
| . | 0.922 | 0.921 | 1.464 | 0.864 | 0.8 | 0.82 |
| . | 0.406 | 0.402 | . | . | 0.532 | 0.433 |
| . | 0.406 | 0.402 | . | . | 0.532 | 0.433 |
| 0.108 | 0.068 | 0.061 | 0.091 | 0.102 | 0.797 | 0.814 |
| . | 0.239 | 0.236 | 0.748 | 0.636 | 0.775 | 0.782 |
| 0.745 | 0.497 | 0.493 | 0.418 | 0.424 | 0.462 | 0.336 |
| 0.432 | 0.864 | 0.862 | 0.147 | 0.166 | 0.55 | 0.458 |
| . | 0.83 | 0.828 | 0.974 | 0.735 | 0.737 | 0.726 |
| 0.11 | 0.815 | 0.814 | 1.097 | 0.776 | 0.803 | 0.825 |
| . | 0.839 | 0.837 | 1.716 | 0.905 | 0.821 | 0.852 |

| | | | | | | |
|---|---|---|---|---|---|---|
| . | 0.225 | 0.221 | 0.74 | 0.632 | 0.634 | 0.577 |
| . | . | . | . | . | . | . |
| . | 0.043 | 0.032 | 0.746 | 0.635 | 0.714 | 0.692 |
| 0.613 | 0.768 | 0.766 | 1.491 | 0.869 | 0.774 | 0.78 |
| . | 0.867 | 0.866 | 1.351 | 0.841 | 0.675 | 0.636 |
| 0.155 | 0.168 | 0.164 | 1.124 | 0.784 | 0.787 | 0.8 |
| 0.439 | 0.665 | 0.663 | 0.323 | 0.345 | 0.762 | 0.762 |
| . | 0.406 | 0.402 | . | . | 0.532 | 0.433 |
| . | 0.817 | 0.816 | 0.17 | 0.191 | 0.819 | 0.848 |
| 0.106 | 0.941 | 0.941 | 2.035 | 0.942 | 0.757 | 0.755 |
| . | . | . | . | . | . | . |
| . | 0.788 | 0.786 | 1.409 | 0.854 | 0.686 | 0.652 |
| 0.871 | 0.825 | 0.824 | 1.424 | 0.857 | 0.701 | 0.673 |
| . | . | . | . | . | . | . |
| . | . | . | . | . | . | . |
| . | 0.922 | 0.921 | 1.464 | 0.864 | 0.8 | 0.82 |
| . | . | . | . | . | . | . |
| 0.016 | 0.7 | 0.698 | 0.382 | 0.396 | 0.495 | 0.381 |
| . | 0.76 | 0.758 | 1.688 | 0.901 | 0.753 | 0.749 |
| . | . | . | . | . | . | . |
| 0.365 | 0.891 | 0.89 | 0.915 | 0.712 | 0.624 | 0.563 |
| 0.078 | 0.129 | 0.125 | 0.974 | 0.735 | 0.598 | 0.526 |
| . | . | . | . | . | . | . |
| . | 0.406 | 0.402 | . | . | 0.532 | 0.433 |
| . | 0.36 | 0.356 | 0.681 | 0.6 | 0.609 | 0.542 |
| . | 0.344 | 0.34 | 0.095 | 0.107 | 0.544 | 0.45 |
| . | 0.212 | 0.208 | 1.047 | 0.76 | 0.765 | 0.767 |
| . | . | . | . | . | . | . |
| . | 0.315 | 0.311 | 2.546 | 0.978 | 0.881 | 0.941 |
| 0.313 | 0.599 | 0.596 | 0.695 | 0.608 | 0.568 | 0.484 |
| . | 0.227 | 0.222 | 0.917 | 0.713 | 0.588 | 0.512 |
| . | 0.3 | 0.296 | 0.166 | 0.188 | 0.488 | 0.371 |
| . | 0.068 | 0.061 | 0.147 | 0.166 | 0.553 | 0.463 |
| 0.302 | 0.872 | 0.871 | 0.028 | 0.029 | 0.405 | 0.258 |
| . | 0.768 | 0.766 | 0.34 | 0.359 | 0.543 | 0.448 |
| 0.548 | 0.643 | 0.64 | 0.856 | 0.687 | 0.754 | 0.75 |
| 0.419 | 0.751 | 0.748 | 0.903 | 0.707 | 0.707 | 0.681 |
| 0.879 | 0.74 | 0.738 | 0.548 | 0.518 | 0.876 | 0.934 |
| 0.098 | 0.72 | 0.718 | 0.856 | 0.688 | 0.692 | 0.659 |
| . | 0.916 | 0.915 | 1.545 | 0.878 | 0.759 | 0.758 |
| . | . | . | . | . | . | . |
| . | . | . | . | . | . | . |
| . | 0.227 | 0.222 | 0.917 | 0.713 | 0.588 | 0.512 |

| | | | | | | | |
|---|---|---|---|---|---|---|---|
| . | . | . | . | . | . | . | |
| . | | 0.406 | 0.402 | . | . | 0.532 | 0.433 |
| . | | 0.768 | 0.766 | 0.34 | 0.359 | 0.543 | 0.448 |
| | 0.571 | 0.611 | 0.608 | . | . | 0.785 | 0.796 |
| . | | 0.806 | 0.804 | 0.179 | 0.202 | 0.491 | 0.376 |
| | 0.243 | 0.75 | 0.748 | 0.83 | 0.676 | 0.825 | 0.858 |
| . | | 0.938 | 0.937 | 0.791 | 0.658 | 0.742 | 0.733 |
| . | | 0.604 | 0.6 | 0.938 | 0.721 | 0.879 | 0.938 |
| | 0.571 | 0.238 | 0.234 | 0.539 | 0.512 | 0.874 | 0.931 |
| . | . | . | . | . | . | . | |
| . | . | . | . | . | . | . | |
| | 0.238 | 0.692 | 0.689 | 0.676 | 0.598 | 0.758 | 0.756 |
| | 0.011 | 0.518 | 0.515 | 0.255 | 0.281 | 0.66 | 0.615 |
| . | . | . | . | . | . | . | |
| . | | 0.493 | 0.489 | 0.608 | 0.556 | 0.525 | 0.424 |
| | 0.029 | 0.103 | 0.098 | 0.578 | 0.537 | 0.563 | 0.477 |
| | 0.121 | 0.918 | 0.917 | 0.607 | 0.555 | 0.806 | 0.829 |
| . | . | . | . | . | . | . | |
| . | | 0.212 | 0.208 | 1.047 | 0.76 | 0.765 | 0.767 |
| | 0.168 | 0.344 | 0.34 | 0.11 | 0.124 | 0.283 | 0.079 |
| | 0.863 | 0.79 | 0.788 | 1.518 | 0.874 | 0.799 | 0.818 |
| . | | 0.938 | 0.937 | 0.791 | 0.658 | 0.742 | 0.733 |
| . | . | . | . | . | . | . | |
| | 0.178 | 0.228 | 0.224 | 0.095 | 0.107 | 0.814 | 0.841 |
| | 0.631 | 0.512 | 0.509 | 0.645 | 0.58 | 0.589 | 0.513 |
| . | | 0.29 | 0.286 | 0.461 | 0.456 | 0.44 | 0.305 |
| | 0.11 | 0.165 | 0.162 | 0.024 | 0.024 | 0.296 | 0.098 |
| . | | 0.54 | 0.537 | 0.37 | 0.386 | 0.374 | 0.215 |
| . | | 0.479 | 0.475 | 0.402 | 0.412 | 0.328 | 0.147 |
| . | | 0.068 | 0.061 | 0.093 | 0.105 | 0.907 | 0.971 |
| . | | 0.694 | 0.692 | 0.143 | 0.161 | 0.615 | 0.55 |
| . | | 0.938 | 0.937 | 0.791 | 0.658 | 0.742 | 0.733 |
| . | . | . | . | . | . | . | |
| . | | 0.696 | 0.693 | 1.138 | 0.788 | 0.745 | 0.737 |
| . | | 0.147 | 0.143 | 0.748 | 0.636 | 0.728 | 0.712 |
| | 0.059 | 0.222 | 0.218 | 0.395 | 0.407 | 0.73 | 0.715 |
| . | | 0.343 | 0.339 | 0.55 | 0.519 | 0.773 | 0.779 |
| . | | 0.391 | 0.387 | 0.182 | 0.205 | 0.762 | 0.762 |
| . | . | . | . | . | . | . | |
| . | | 0.694 | 0.692 | 0.143 | 0.161 | 0.615 | 0.55 |
| | 0.646 | 0.841 | 0.84 | . | . | 0.862 | 0.913 |
| . | | 0.628 | 0.625 | 0.154 | 0.174 | 0.492 | 0.377 |
| . | | 0.702 | 0.699 | 1.837 | 0.921 | 0.829 | 0.864 |

| | | | | | | |
|---|---|---|---|---|---|---|
| 0.333 | 0.748 | 0.745 | 0.469 | 0.462 | 0.473 | 0.351 |
| . | 0.721 | 0.719 | 0.926 | 0.716 | 0.679 | 0.641 |
| . | . | . | . | . | . | . |
| . | . | . | . | . | . | . |
| 0.11 | 0.165 | 0.162 | 0.024 | 0.024 | 0.296 | 0.098 |
| 0.356 | 0.103 | 0.098 | . | . | . | . |
| . | 0.295 | 0.291 | 0.401 | 0.411 | 0.824 | 0.856 |
| 0.863 | 0.79 | 0.788 | 1.518 | 0.874 | 0.799 | 0.818 |
| . | 0.243 | 0.239 | 0.142 | 0.16 | 0.527 | 0.427 |
| . | . | . | . | . | . | . |
| 0.847 | 0.514 | 0.511 | 1.044 | 0.759 | 0.844 | 0.887 |
| . | 0.2 | 0.196 | 0.095 | 0.107 | 0.656 | 0.609 |
| 0.758 | 0.824 | 0.822 | 1.695 | 0.902 | 0.889 | 0.951 |
| 0.758 | 0.824 | 0.822 | 1.695 | 0.902 | 0.889 | 0.951 |
| . | . | . | . | . | . | . |
| . | . | . | . | . | . | . |
| 0.65 | 0.857 | 0.855 | 1.943 | 0.933 | 0.822 | 0.854 |
| 0.133 | 0.499 | 0.495 | 0.007 | 0.007 | 0.735 | 0.722 |
| 0.787 | 0.718 | 0.715 | 1.969 | 0.935 | 0.838 | 0.877 |
| . | . | . | . | . | . | . |
| 0.178 | 0.228 | 0.224 | 0.095 | 0.107 | 0.814 | 0.841 |
| . | 0.372 | 0.368 | 0.51 | 0.491 | 0.784 | 0.796 |
| . | 0.608 | 0.605 | 0.709 | 0.616 | 0.627 | 0.568 |
| 0.294 | 0.708 | 0.706 | 2.487 | 0.976 | 0.915 | 0.979 |
| 0.202 | 0.273 | 0.269 | 0.108 | 0.122 | 0.842 | 0.883 |
| . | 0.187 | 0.183 | 0.436 | 0.437 | 0.888 | 0.95 |
| . | . | . | . | . | . | . |
| 0.847 | 0.514 | 0.511 | 1.044 | 0.759 | 0.844 | 0.887 |
| . | 0.299 | 0.295 | 0.085 | 0.096 | 0.77 | 0.774 |
| . | 0.406 | 0.402 | . | . | 0.532 | 0.433 |
| . | 0.694 | 0.692 | 0.143 | 0.161 | 0.615 | 0.55 |
| . | 0.2 | 0.196 | 0.095 | 0.107 | 0.656 | 0.609 |
| . | 0.694 | 0.692 | 0.143 | 0.161 | 0.615 | 0.55 |
| . | 0.768 | 0.766 | 0.34 | 0.359 | 0.543 | 0.448 |
| 0.129 | 0.162 | 0.158 | 1.033 | 0.755 | 0.496 | 0.383 |
| 0.297 | 0.658 | 0.655 | 0.459 | 0.455 | 0.84 | 0.881 |
| . | 0.231 | 0.227 | 0.709 | 0.616 | 0.491 | 0.375 |
| . | 0.608 | 0.605 | 0.709 | 0.616 | 0.627 | 0.568 |
| . | 0.537 | 0.533 | 0.691 | 0.606 | 0.506 | 0.397 |
| 0.8 | 0.85 | 0.848 | . | . | 0.877 | 0.935 |
| . | . | . | . | . | . | . |
| . | 0.405 | 0.401 | 1.337 | 0.838 | 0.805 | 0.828 |
| . | 0.147 | 0.143 | 0.748 | 0.636 | 0.728 | 0.712 |

| | | | | | | | |
|---|---|---|---|---|---|---|---|
| . | . | . | . | . | . | . | |
| . | | 0.694 | 0.692 | 0.143 | 0.161 | 0.615 | 0.55 |
| . | | 0.608 | 0.605 | 0.709 | 0.616 | 0.627 | 0.568 |
| . | | 0.147 | 0.143 | 0.748 | 0.636 | 0.728 | 0.712 |
| | 0.11 | 0.165 | 0.162 | 0.024 | 0.024 | 0.296 | 0.098 |
| . | | 0.147 | 0.143 | 0.748 | 0.636 | 0.728 | 0.712 |
| | 0.123 | 0.068 | 0.061 | 0.296 | 0.32 | 0.406 | 0.259 |
| | 0.087 | 0.243 | 0.239 | 0.486 | 0.475 | 0.739 | 0.728 |
| . | . | . | . | . | . | . | |
| . | | 0.343 | 0.339 | 0.55 | 0.519 | 0.773 | 0.779 |
| . | | 0.231 | 0.227 | 0.709 | 0.616 | 0.491 | 0.375 |
| . | | 0.379 | 0.375 | 0.269 | 0.294 | 0.305 | 0.112 |
| . | . | . | . | . | . | . | |
| | 0.205 | 0.781 | 0.779 | 0.425 | 0.429 | 0.335 | 0.156 |
| . | | 0.696 | 0.693 | 1.138 | 0.788 | 0.745 | 0.737 |
| | 0.774 | 0.421 | 0.417 | 1.373 | 0.846 | 0.719 | 0.699 |
| . | | 0.117 | 0.112 | 0.862 | 0.69 | 0.535 | 0.438 |
| . | . | . | . | . | . | . | |
| | 0.47 | 0.604 | 0.601 | 0.704 | 0.613 | 0.653 | 0.604 |
| | 0.032 | 0.431 | 0.428 | 0.348 | 0.367 | 0.711 | 0.688 |
| . | | 0.117 | 0.112 | 0.862 | 0.69 | 0.535 | 0.438 |
| . | | 0.279 | 0.275 | 0.086 | 0.097 | 0.648 | 0.597 |
| . | . | . | . | . | . | . | |
| | 0.75 | 0.452 | 0.448 | 0.827 | 0.674 | 0.896 | 0.96 |
| . | . | . | . | . | . | . | |
| | 0.914 | 0.864 | 0.862 | 0.721 | 0.622 | 0.511 | 0.403 |
| | 0.203 | 0.291 | 0.288 | 0.39 | 0.402 | 0.402 | 0.254 |
| . | . | . | . | . | . | . | |
| . | . | . | . | . | . | . | |
| | 0.272 | 0.739 | 0.737 | 0.868 | 0.693 | 0.581 | 0.503 |
| | 0.876 | 0.943 | 0.942 | 0.461 | 0.456 | 0.86 | 0.911 |
| | 0.542 | 0.841 | 0.839 | 0.917 | 0.713 | 0.76 | 0.76 |
| | 0.666 | 0.277 | 0.273 | 0.102 | 0.116 | 0.718 | 0.697 |
| | 0.302 | 0.768 | 0.766 | 0.489 | 0.477 | 0.889 | 0.951 |
| . | . | . | . | . | . | . | |
| . | | 0.766 | 0.764 | 0.279 | 0.304 | 0.315 | 0.127 |
| | 0.542 | 0.841 | 0.839 | 0.917 | 0.713 | 0.76 | 0.76 |
| | 0.706 | 0.706 | 0.704 | 1.635 | 0.893 | 0.768 | 0.771 |
| . | | 0.37 | 0.366 | 0.709 | 0.616 | 0.386 | 0.23 |
| . | | 0.348 | 0.344 | 0.145 | 0.164 | 0.358 | 0.19 |
| | 0.63 | 0.39 | 0.386 | 0.559 | 0.524 | 0.859 | 0.91 |
| . | | 0.687 | 0.684 | 0.75 | 0.637 | 0.855 | 0.904 |
| | 0.294 | 0.567 | 0.564 | 1.146 | 0.791 | 0.899 | 0.963 |

| | | | | | | |
|---|---|---|---|---|---|---|
| . | . | . | . | . | . | . |
| 0.914 | 0.864 | 0.862 | 0.721 | 0.622 | 0.511 | 0.403 |
| . | 0.797 | 0.796 | 1.857 | 0.923 | 0.744 | 0.736 |
| 0.309 | 0.857 | 0.856 | 1.506 | 0.871 | 0.82 | 0.85 |
| . | 0.472 | 0.468 | 0.588 | 0.544 | 0.634 | 0.577 |
| 0.406 | 0.139 | 0.135 | 0.488 | 0.476 | 0.722 | 0.703 |
| 0.745 | 0.741 | 0.739 | 1.426 | 0.857 | 0.785 | 0.796 |
| . | . | . | . | . | . | . |
| 0.378 | 0.339 | 0.336 | 0.316 | 0.339 | 0.477 | 0.357 |
| 0.071 | 0.762 | 0.76 | . | . | . | . |
| 0.011 | 0.518 | 0.515 | 0.255 | 0.281 | 0.66 | 0.615 |
| 0.737 | 0.395 | 0.391 | 0.271 | 0.296 | 0.425 | 0.285 |
| 0.892 | 0.932 | 0.931 | 3.067 | 0.993 | 0.933 | 0.991 |
| . | 0.384 | 0.38 | 0.918 | 0.713 | 0.833 | 0.87 |
| 0.087 | 0.765 | 0.763 | 1.003 | 0.745 | 0.576 | 0.496 |
| 0.065 | 0.147 | 0.143 | 0.158 | 0.179 | 0.315 | 0.127 |
| 0.668 | 0.686 | 0.684 | 0.115 | 0.13 | 0.601 | 0.53 |
| . | 0.345 | 0.341 | 1.001 | 0.744 | 0.888 | 0.949 |
| . | 0.222 | 0.218 | 0.361 | 0.378 | 0.576 | 0.495 |
| 0.668 | 0.686 | 0.684 | 0.115 | 0.13 | 0.601 | 0.53 |
| 0.056 | 0.155 | 0.151 | 0.082 | 0.093 | 0.572 | 0.49 |
| 0.483 | 0.839 | 0.837 | 0.876 | 0.696 | 0.702 | 0.675 |
| . | 0.279 | 0.275 | 0.086 | 0.097 | 0.648 | 0.597 |
| 0.668 | 0.686 | 0.684 | 0.115 | 0.13 | 0.601 | 0.53 |
| 0.66 | 0.234 | 0.23 | 0.363 | 0.379 | 0.773 | 0.779 |
| . | . | . | . | . | . | . |
| . | 0.345 | 0.341 | 1.001 | 0.744 | 0.888 | 0.949 |
| . | 0.625 | 0.622 | 0.344 | 0.363 | 0.706 | 0.681 |
| . | 0.117 | 0.112 | 0.862 | 0.69 | 0.535 | 0.438 |
| . | 0.269 | 0.265 | 0.129 | 0.146 | 0.792 | 0.807 |
| 0.568 | 0.747 | 0.745 | 0.477 | 0.468 | 0.841 | 0.882 |
| 0.627 | 0.681 | 0.678 | 0.501 | 0.485 | 0.923 | 0.985 |
| 0.745 | 0.741 | 0.739 | 1.426 | 0.857 | 0.785 | 0.796 |
| 0.668 | 0.686 | 0.684 | 0.115 | 0.13 | 0.601 | 0.53 |
| . | 0.625 | 0.622 | 0.344 | 0.363 | 0.706 | 0.681 |
| . | 0.37 | 0.366 | 0.709 | 0.616 | 0.386 | 0.23 |
| 0.829 | 0.665 | 0.662 | 1.647 | 0.895 | 0.679 | 0.642 |
| . | . | . | . | . | . | . |
| 0.191 | 0.954 | 0.953 | 0.507 | 0.489 | 0.492 | 0.377 |
| 0.987 | 0.824 | 0.823 | 1.689 | 0.901 | 0.864 | 0.916 |
| . | 0.873 | 0.872 | 1.483 | 0.867 | 0.801 | 0.822 |
| 0.309 | 0.857 | 0.856 | 1.506 | 0.871 | 0.82 | 0.85 |
| 0.668 | 0.686 | 0.684 | 0.115 | 0.13 | 0.601 | 0.53 |

| | | | | | | | |
|---|---|---|---|---|---|---|---|
| . | | 0.465 | 0.461 | 1.761 | 0.911 | 0.572 | 0.489 |
| . | | 0.625 | 0.622 | 0.344 | 0.363 | 0.706 | 0.681 |
| | 0.651 | 0.565 | 0.562 | 1.355 | 0.842 | 0.674 | 0.634 |

| PrimateAI_pr | DEOGEN2_sc | DEOGEN2_ra | DEOGEN2_pr | BayesDel_ad | BayesDel_ad | BayesDel_ad |
|---|---|---|---|---|---|---|
| . | . | . | . | . | . | . |
| . | . | . | . | . | . | . |
| T | 0.45 | 0.791 | T | -0.441 | 0.013 | T |
| T | 0.678 | 0.905 | D | 0.013 | 0.534 | T |
| T | 0.049 | 0.283 | T | -0.238 | 0.155 | T |
| D | 0.135 | 0.467 | T | 0.043 | 0.574 | T |
| T | 0.355 | 0.722 | T | 0.302 | 0.831 | D |
| T | 0.374 | 0.776 | T | -0.138 | 0.302 | T |
| T | 0.246 | 0.914 | T | 0.133 | 0.677 | D |
| T | 0.267 | 0.639 | T | -0.287 | 0.1 | T |
| T | 0.075 | 0.798 | T | -0.142 | 0.296 | T |
| T | 0.45 | 0.791 | T | -0.441 | 0.013 | T |
| D | 0.568 | 0.857 | D | 0.03 | 0.557 | T |
| T | 0.365 | 0.731 | T | -0.037 | 0.464 | T |
| . | . | . | . | . | . | . |
| D | 0.75 | 0.932 | D | -0.259 | 0.13 | T |
| D | 0.031 | 0.221 | T | -0.069 | 0.415 | T |
| D | 0.201 | 0.558 | T | 0.226 | 0.763 | D |
| T | 0.075 | 0.798 | T | -0.142 | 0.296 | T |
| T | 0.5 | 0.821 | D | -0.162 | 0.264 | T |
| D | 0.521 | 0.833 | D | 0.057 | 0.592 | T |
| . | . | . | . | . | . | . |
| T | 0.371 | 0.735 | T | -0.127 | 0.32 | T |
| T | 0.089 | 0.837 | T | 0.049 | 0.582 | T |
| . | . | . | . | 0.622 | 0.988 | D |
| T | 0.095 | 0.844 | T | 0.108 | 0.651 | D |
| T | 0.957 | 0.994 | D | -0.052 | 0.441 | T |
| T | 0.07 | 0.338 | T | -0.23 | 0.166 | T |
| T | 0.104 | 0.412 | T | 0.022 | 0.546 | T |
| D | 0.329 | 0.699 | T | -0.126 | 0.321 | T |
| T | 0.075 | 0.798 | T | -0.142 | 0.296 | T |
| D | 0.559 | 0.852 | D | -0.193 | 0.218 | T |
| T | 0.152 | 0.493 | T | 0.054 | 0.589 | T |
| T | 0.267 | 0.639 | T | -0.287 | 0.1 | T |
| T | 0.251 | 0.621 | T | -0.355 | 0.044 | T |
| T | 0.192 | 0.547 | T | -0.108 | 0.35 | T |
| T | 0.364 | 0.73 | T | -0.139 | 0.301 | T |
| D | 0.643 | 0.891 | D | 0.144 | 0.687 | D |
| T | 0.392 | 0.752 | T | -0.115 | 0.339 | T |
| T | 0.37 | 0.735 | T | 0.212 | 0.75 | D |
| T | 0.171 | 0.869 | T | -0.186 | 0.228 | T |
| T | 0.106 | 0.417 | T | -0.204 | 0.203 | T |

| | | | | | | |
|---|---|---|---|---|---|---|
| T | 0.45 | 0.791 | T | -0.441 | 0.013 | T |
| T | 0.185 | 0.538 | T | -0.097 | 0.369 | T |
| T | 0.738 | 0.927 | D | -0.392 | 0.026 | T |
| T | 0.275 | 0.647 | T | 0.284 | 0.816 | D |
| D | 0.521 | 0.833 | D | 0.057 | 0.592 | T |
| . | . | . | . | 0.625 | 0.994 | D |
| . | . | . | . | . | . | . |
| T | 0.139 | 0.705 | T | -0.171 | 0.251 | T |
| T | 0.828 | 0.959 | D | 0.154 | 0.696 | D |
| T | 0.197 | 0.553 | T | -0.147 | 0.288 | T |
| D | 0.149 | 0.487 | T | -0.134 | 0.308 | T |
| T | 0.409 | 0.764 | T | -0.225 | 0.173 | T |
| T | 0.307 | 0.679 | T | -0.303 | 0.083 | T |
| . | . | . | . | . | . | . |
| T | 0.408 | 0.763 | T | -0.453 | 0.011 | T |
| D | 0.648 | 0.893 | D | -0.174 | 0.246 | T |
| D | 0.838 | 0.962 | D | 0.346 | 0.862 | D |
| T | 0.214 | 0.575 | T | -0.306 | 0.081 | T |
| T | 0.345 | 0.714 | T | 0.089 | 0.631 | D |
| T | 0.285 | 0.658 | T | -0.184 | 0.232 | T |
| T | 0.353 | 0.721 | T | -0.282 | 0.105 | T |
| . | . | . | . | . | . | . |
| T | 0.45 | 0.791 | T | -0.441 | 0.013 | T |
| T | 0.388 | 0.749 | T | -0.339 | 0.054 | T |
| T | 0.438 | 0.784 | T | 0.038 | 0.568 | T |
| T | 0.075 | 0.798 | T | -0.142 | 0.296 | T |
| T | 0.093 | 0.39 | T | 0.035 | 0.564 | T |
| T | 0.15 | 0.49 | T | -0.285 | 0.102 | T |
| T | 0.253 | 0.743 | T | -0.35 | 0.047 | T |
| . | . | . | . | . | . | . |
| T | 0.485 | 0.812 | T | 0.269 | 0.803 | D |
| T | 0.762 | 0.936 | D | -0.078 | 0.4 | T |
| T | 0.734 | 0.926 | D | 0.187 | 0.727 | D |
| T | 0.246 | 0.914 | T | 0.133 | 0.677 | D |
| T | 0.45 | 0.791 | T | -0.441 | 0.013 | T |
| T | 0.45 | 0.791 | T | -0.441 | 0.013 | T |
| T | 0.347 | 0.716 | T | 0.122 | 0.665 | D |
| T | 0.03 | 0.532 | T | -0.103 | 0.359 | T |
| T | 0.317 | 0.689 | T | -0.172 | 0.249 | T |
| T | 0.646 | 0.892 | D | 0.334 | 0.854 | D |
| T | 0.173 | 0.522 | T | -0.024 | 0.483 | T |
| D | 0.75 | 0.932 | D | -0.259 | 0.13 | T |
| D | 0.78 | 0.942 | D | -0.06 | 0.429 | T |

| | | | | | | |
|---|---|---|---|---|---|---|
| T | 0.029 | 0.393 | T | -0.088 | 0.384 | T |
| . | . | . | . | . | . | . |
| T | 0.014 | 0.33 | T | -0.349 | 0.048 | T |
| T | 0.689 | 0.91 | D | 0.201 | 0.74 | D |
| T | 0.828 | 0.959 | D | 0.154 | 0.696 | D |
| T | 0.232 | 0.598 | T | 0.097 | 0.639 | D |
| T | 0.219 | 0.581 | T | -0.048 | 0.448 | T |
| T | 0.45 | 0.791 | T | -0.441 | 0.013 | T |
| D | 0.377 | 0.826 | T | 0.112 | 0.655 | D |
| T | 0.943 | 0.991 | D | 0.383 | 0.889 | D |
| . | . | . | . | . | . | . |
| T | 0.049 | 0.283 | T | -0.238 | 0.155 | T |
| T | 0.82 | 0.956 | D | 0.094 | 0.637 | D |
| . | . | . | . | . | . | . |
| . | . | . | . | . | . | . |
| T | 0.246 | 0.914 | T | 0.133 | 0.677 | D |
| . | . | . | . | . | . | . |
| T | 0.156 | 0.498 | T | 0.193 | 0.733 | D |
| T | 0.306 | 0.678 | T | -0.071 | 0.411 | T |
| . | . | . | . | . | . | . |
| T | 0.846 | 0.965 | D | 0.334 | 0.854 | D |
| T | 0.027 | 0.198 | T | 0.076 | 0.616 | D |
| . | . | . | . | . | . | . |
| T | 0.45 | 0.791 | T | -0.441 | 0.013 | T |
| T | 0.075 | 0.798 | T | -0.142 | 0.296 | T |
| T | 0.109 | 0.423 | T | -0.06 | 0.428 | T |
| T | 0.267 | 0.639 | T | -0.287 | 0.1 | T |
| . | . | . | . | . | . | . |
| D | 0.326 | 0.696 | T | -0.373 | 0.034 | T |
| T | 0.714 | 0.919 | D | 0.256 | 0.791 | D |
| T | 0.394 | 0.753 | T | -0.187 | 0.227 | T |
| T | 0.184 | 0.536 | T | -0.28 | 0.107 | T |
| T | 0.349 | 0.717 | T | -0.321 | 0.069 | T |
| T | 0.685 | 0.908 | D | -0.181 | 0.236 | T |
| T | 0.285 | 0.658 | T | -0.184 | 0.232 | T |
| T | 0.382 | 0.744 | T | 0.067 | 0.605 | T |
| T | 0.199 | 0.556 | T | 0.025 | 0.551 | T |
| D | 0.494 | 0.817 | T | -0.063 | 0.424 | T |
| T | 0.628 | 0.884 | D | 0.072 | 0.611 | D |
| T | 0.757 | 0.934 | D | 0.144 | 0.687 | D |
| . | . | . | . | . | . | . |
| . | . | . | . | . | . | . |
| T | 0.394 | 0.753 | T | -0.187 | 0.227 | T |

| | | | | | | |
|---|---|---|---|---|---|---|
| . | . | . | . | . | . | . |
| T | 0.45 | 0.791 | T | -0.441 | 0.013 | T |
| T | 0.285 | 0.658 | T | -0.184 | 0.232 | T |
| T | 0.418 | 0.771 | T | -0.262 | 0.127 | T |
| T | 0.41 | 0.765 | T | -0.06 | 0.428 | T |
| D | 0.823 | 0.957 | D | 0.264 | 0.799 | D |
| T | 0.487 | 0.813 | T | 0.088 | 0.629 | D |
| D | 0.481 | 0.809 | T | -0.005 | 0.51 | T |
| D | 0.383 | 0.745 | T | 0.298 | 0.828 | D |
| . | . | . | . | 0.364 | 0.876 | D |
| . | . | . | . | 0.625 | 0.994 | D |
| T | 0.232 | 0.908 | T | 0.227 | 0.764 | D |
| T | 0.079 | 0.36 | T | -0.248 | 0.144 | T |
| . | . | . | . | . | . | . |
| T | 0.341 | 0.71 | T | -0.351 | 0.047 | T |
| T | 0.034 | 0.231 | T | -0.022 | 0.486 | T |
| D | 0.527 | 0.836 | D | 0.363 | 0.875 | D |
| . | . | . | . | . | . | . |
| T | 0.267 | 0.639 | T | -0.287 | 0.1 | T |
| T | 0.185 | 0.537 | T | -0.241 | 0.153 | T |
| T | 0.88 | 0.975 | D | 0.187 | 0.727 | D |
| T | 0.487 | 0.813 | T | 0.088 | 0.629 | D |
| . | . | . | . | . | . | . |
| D | 0.153 | 0.494 | T | -0.067 | 0.418 | T |
| T | 0.449 | 0.791 | T | -0.041 | 0.459 | T |
| T | 0.143 | 0.479 | T | -0.203 | 0.203 | T |
| T | 0.182 | 0.534 | T | -0.396 | 0.024 | T |
| T | 0.2 | 0.558 | T | -0.364 | 0.039 | T |
| T | 0.205 | 0.564 | T | -0.358 | 0.043 | T |
| D | 0.378 | 0.741 | T | -0.106 | 0.355 | T |
| T | 0.253 | 0.743 | T | -0.35 | 0.047 | T |
| T | 0.487 | 0.813 | T | 0.088 | 0.629 | D |
| . | . | . | . | . | . | . |
| T | 0.578 | 0.861 | D | 0.009 | 0.529 | T |
| T | 0.33 | 0.7 | T | -0.375 | 0.033 | T |
| T | 0.198 | 0.876 | T | 0.081 | 0.622 | D |
| T | 0.394 | 0.753 | T | 0.038 | 0.568 | T |
| T | 0.279 | 0.652 | T | -0.029 | 0.476 | T |
| . | . | . | . | . | . | . |
| T | 0.253 | 0.743 | T | -0.35 | 0.047 | T |
| D | 0.429 | 0.778 | T | 0.324 | 0.847 | D |
| T | 0.254 | 0.624 | T | -0.131 | 0.313 | T |
| D | 0.195 | 0.551 | T | 0.419 | 0.91 | D |

| | | | | | | | | |
|---|---|---|---|---|---|---|---|---|
| T | | 0.431 | 0.78 | T | | -0.122 | 0.328 | T |
| T | | 0.83 | 0.959 | D | | 0.195 | 0.734 | D |
| . | . | . | | . | . | . | | . |
| . | . | . | | . | . | . | | . |
| T | | 0.182 | 0.534 | T | | -0.396 | 0.024 | T |
| . | . | . | | . | | -0.074 | 0.407 | T |
| D | | 0.368 | 0.733 | T | | -0.009 | 0.504 | T |
| T | | 0.88 | 0.975 | D | | 0.187 | 0.727 | D |
| T | | 0.118 | 0.438 | T | | -0.269 | 0.119 | T |
| . | . | . | | . | . | . | | . |
| D | | 0.2 | 0.557 | T | | -0.26 | 0.129 | T |
| T | | 0.118 | 0.438 | T | | -0.219 | 0.182 | T |
| D | | 0.827 | 0.958 | D | | -0.144 | 0.293 | T |
| D | | 0.827 | 0.958 | D | | -0.144 | 0.293 | T |
| . | . | . | | . | . | . | | . |
| . | . | . | | . | . | . | | . |
| D | | 0.635 | 0.887 | D | | 0.4 | 0.899 | D |
| T | | 0.118 | 0.439 | T | | 0.049 | 0.583 | T |
| D | | 0.709 | 0.917 | D | | 0.186 | 0.726 | D |
| . | . | . | | . | . | . | | . |
| D | | 0.153 | 0.494 | T | | -0.067 | 0.418 | T |
| T | | 0.196 | 0.553 | T | | 0.071 | 0.61 | D |
| T | | 0.365 | 0.731 | T | | -0.037 | 0.464 | T |
| D | | 0.544 | 0.844 | D | | -0.085 | 0.388 | T |
| D | | 0.373 | 0.737 | T | | 0.219 | 0.757 | D |
| D | | 0.129 | 0.456 | T | | -0.272 | 0.115 | T |
| . | . | . | | . | . | . | | . |
| D | | 0.2 | 0.557 | T | | -0.26 | 0.129 | T |
| T | | 0.146 | 0.483 | T | | -0.301 | 0.086 | T |
| T | | 0.45 | 0.791 | T | | -0.441 | 0.013 | T |
| T | | 0.253 | 0.743 | T | | -0.35 | 0.047 | T |
| T | | 0.118 | 0.438 | T | | -0.219 | 0.182 | T |
| T | | 0.253 | 0.743 | T | | -0.35 | 0.047 | T |
| T | | 0.285 | 0.658 | T | | -0.184 | 0.232 | T |
| T | | 0.023 | 0.178 | T | | 0.071 | 0.61 | D |
| D | | 0.021 | 0.162 | T | | -0.094 | 0.374 | T |
| T | | 0.171 | 0.869 | T | | -0.186 | 0.228 | T |
| T | | 0.365 | 0.731 | T | | -0.037 | 0.464 | T |
| T | | 0.738 | 0.927 | D | | -0.392 | 0.026 | T |
| D | | 0.61 | 0.876 | D | | 0.353 | 0.867 | D |
| . | . | . | | . | . | . | | . |
| D | | 0.412 | 0.935 | T | | -0.259 | 0.13 | T |
| T | | 0.33 | 0.7 | T | | -0.375 | 0.033 | T |

| | | | | | | |
|---|---|---|---|---|---|---|
| . | . | . | . | . | . | . |
| T | 0.253 | 0.743 | T | -0.35 | 0.047 | T |
| T | 0.365 | 0.731 | T | -0.037 | 0.464 | T |
| T | 0.33 | 0.7 | T | -0.375 | 0.033 | T |
| T | 0.182 | 0.534 | T | -0.396 | 0.024 | T |
| T | 0.33 | 0.7 | T | -0.375 | 0.033 | T |
| T | 0.029 | 0.21 | T | -0.162 | 0.265 | T |
| T | 0.127 | 0.454 | T | 0.048 | 0.581 | T |
| . | . | . | . | . | . | . |
| T | 0.394 | 0.753 | T | 0.038 | 0.568 | T |
| T | 0.171 | 0.869 | T | -0.186 | 0.228 | T |
| T | 0.223 | 0.587 | T | -0.431 | 0.015 | T |
| . | . | . | . | . | . | . |
| T | 0.225 | 0.59 | T | 0.103 | 0.647 | D |
| T | 0.578 | 0.861 | D | 0.009 | 0.529 | T |
| T | 0.347 | 0.716 | T | 0.097 | 0.64 | D |
| T | 0.189 | 0.543 | T | -0.342 | 0.053 | T |
| . | . | . | . | . | . | . |
| T | 0.185 | 0.954 | T | -0.083 | 0.391 | T |
| T | 0.097 | 0.399 | T | -0.216 | 0.185 | T |
| T | 0.189 | 0.543 | T | -0.342 | 0.053 | T |
| T | 0.365 | 0.731 | T | -0.204 | 0.202 | T |
| . | . | . | . | . | . | . |
| D | 0.252 | 0.622 | T | 0.124 | 0.667 | D |
| . | . | . | . | . | . | . |
| T | 0.743 | 0.929 | D | -0.037 | 0.464 | T |
| T | 0.196 | 0.552 | T | 0.008 | 0.528 | T |
| . | . | . | . | 0.625 | 0.994 | D |
| . | . | . | . | . | . | . |
| T | 0.275 | 0.648 | T | -0.037 | 0.463 | T |
| D | 0.632 | 0.886 | D | 0.248 | 0.784 | D |
| T | 0.349 | 0.717 | T | 0.042 | 0.573 | T |
| T | 0.38 | 0.743 | T | 0.156 | 0.698 | D |
| D | 0.508 | 0.825 | D | 0.057 | 0.592 | T |
| . | . | . | . | . | . | . |
| T | 0.347 | 0.715 | T | -0.356 | 0.043 | T |
| T | 0.349 | 0.717 | T | 0.042 | 0.573 | T |
| T | 0.546 | 0.845 | D | 0.176 | 0.717 | D |
| T | 0.033 | 0.491 | T | -0.3 | 0.087 | T |
| T | 0.221 | 0.585 | T | -0.498 | 0.006 | T |
| D | 0.375 | 0.739 | T | -0.091 | 0.379 | T |
| D | 0.599 | 0.871 | D | 0.115 | 0.659 | D |
| D | 0.669 | 0.902 | D | -0.153 | 0.279 | T |

| | | | | | | |
|---|---|---|---|---|---|---|
| . | . | . | . | . | . | . |
| T | 0.743 | 0.929 | D | -0.037 | 0.464 | T |
| T | 0.021 | 0.166 | T | -0.04 | 0.459 | T |
| D | 0.604 | 0.874 | D | 0.024 | 0.55 | T |
| T | 0.406 | 0.762 | T | -0.193 | 0.219 | T |
| T | 0.375 | 0.739 | T | -0.132 | 0.312 | T |
| T | 0.95 | 0.993 | D | -0.009 | 0.503 | T |
| . | . | . | . | . | . | . |
| T | 0.19 | 0.545 | T | -0.509 | 0.005 | T |
| . | . | . | . | 0.21 | 0.748 | D |
| T | 0.079 | 0.36 | T | -0.248 | 0.144 | T |
| T | 0.62 | 0.881 | D | 0.049 | 0.582 | T |
| D | 0.722 | 0.922 | D | 0.488 | 0.939 | D |
| D | 0.651 | 0.894 | D | -0.104 | 0.358 | T |
| T | 0.739 | 0.928 | D | 0.124 | 0.668 | D |
| T | 0.345 | 0.713 | T | -0.291 | 0.095 | T |
| T | 0.24 | 0.724 | T | -0.443 | 0.013 | T |
| D | 0.185 | 0.538 | T | -0.151 | 0.281 | T |
| T | 0.157 | 0.499 | T | -0.498 | 0.006 | T |
| T | 0.24 | 0.724 | T | -0.443 | 0.013 | T |
| T | 0.346 | 0.715 | T | -0.221 | 0.179 | T |
| T | 0.202 | 0.56 | T | -0.036 | 0.465 | T |
| T | 0.365 | 0.731 | T | -0.204 | 0.202 | T |
| T | 0.24 | 0.724 | T | -0.443 | 0.013 | T |
| T | 0.133 | 0.463 | T | -0.262 | 0.126 | T |
| . | . | . | . | . | . | . |
| D | 0.185 | 0.538 | T | -0.151 | 0.281 | T |
| T | 0.572 | 0.859 | D | -0.265 | 0.123 | T |
| T | 0.189 | 0.543 | T | -0.342 | 0.053 | T |
| T | 0.125 | 0.45 | T | -0.411 | 0.02 | T |
| D | 0.203 | 0.562 | T | 0.085 | 0.626 | D |
| D | 0.399 | 0.757 | T | 0.265 | 0.8 | D |
| T | 0.95 | 0.993 | D | -0.009 | 0.503 | T |
| T | 0.24 | 0.724 | T | -0.443 | 0.013 | T |
| T | 0.572 | 0.859 | D | -0.265 | 0.123 | T |
| T | 0.033 | 0.491 | T | -0.3 | 0.087 | T |
| T | 0.323 | 0.694 | T | 0.025 | 0.551 | T |
| . | . | . | . | . | . | . |
| T | 0.522 | 0.833 | D | 0.263 | 0.798 | D |
| D | 0.356 | 0.723 | T | 0.373 | 0.883 | D |
| T | 0.708 | 0.917 | D | -0.027 | 0.479 | T |
| D | 0.604 | 0.874 | D | 0.024 | 0.55 | T |
| T | 0.24 | 0.724 | T | -0.443 | 0.013 | T |

| | | | | | | |
|---|---|---|---|---|---|---|
| T | 0.169 | 0.516 | T | -0.561 | 0.003 | T |
| T | 0.572 | 0.859 | D | -0.265 | 0.123 | T |
| T | 0.306 | 0.678 | T | 0.08 | 0.62 | D |

| BayesDel_no | BayesDel_no | BayesDel_no | ClinPred_scor | ClinPred_rank | ClinPred_pre | LIST.S2_score |
|---|---|---|---|---|---|---|
| . | . | . | . | . | . | . |
| . | . | . | . | . | . | . |
| -0.405 | 0.329 | T | 0.038 | 0.033 | T | 0.904 |
| -0.155 | 0.588 | T | 0.995 | 0.862 | D | 0.895 |
| -0.121 | 0.617 | T | 0.044 | 0.043 | T | 0.952 |
| -0.176 | 0.569 | T | 0.757 | 0.437 | D | 0.937 |
| 0.388 | 0.917 | D | 0.739 | 0.427 | D | 0.988 |
| -0.328 | 0.417 | T | 0.9 | 0.553 | D | 0.924 |
| 0.418 | 0.927 | D | 0.091 | 0.113 | T | 0.954 |
| -0.18 | 0.565 | T | 0.022 | 0.009 | T | 0.927 |
| 0.02 | 0.716 | D | 0.109 | 0.134 | T | 0.941 |
| -0.405 | 0.329 | T | 0.038 | 0.033 | T | 0.904 |
| -0.195 | 0.551 | T | 0.709 | 0.411 | D | 0.914 |
| 0.092 | 0.763 | D | 0.13 | 0.154 | T | 0.971 |
| . | . | . | . | . | . | . |
| -0.147 | 0.595 | T | 0.027 | 0.015 | T | 0.929 |
| -0.337 | 0.407 | T | 0.492 | 0.323 | T | 0.924 |
| 0.086 | 0.76 | D | 0.991 | 0.816 | D | 0.975 |
| 0.02 | 0.716 | D | 0.109 | 0.134 | T | 0.941 |
| -0.223 | 0.525 | T | 0.332 | 0.263 | T | 0.939 |
| 0.118 | 0.781 | D | 0.445 | 0.306 | T | 0.966 |
| . | . | . | . | . | . | . |
| -0.092 | 0.64 | T | 0.646 | 0.383 | D | 0.595 |
| 0.026 | 0.72 | D | 0.457 | 0.311 | T | 0.981 |
| 0.656 | 0.987 | D | . | . | . | . |
| 0.154 | 0.804 | D | 0.989 | 0.796 | D | 0.973 |
| -0.126 | 0.613 | T | 0.866 | 0.516 | D | 0.952 |
| -0.375 | 0.363 | T | 0.102 | 0.126 | T | 0.829 |
| -0.206 | 0.54 | T | 0.779 | 0.45 | D | 0.577 |
| -0.103 | 0.632 | T | 0.323 | 0.259 | T | 0.97 |
| 0.02 | 0.716 | D | 0.109 | 0.134 | T | 0.941 |
| -0.17 | 0.574 | T | 0.189 | 0.198 | T | 0.723 |
| -0.16 | 0.583 | T | 0.382 | 0.282 | T | 0.888 |
| -0.18 | 0.565 | T | 0.022 | 0.009 | T | 0.927 |
| -0.285 | 0.463 | T | 0.193 | 0.2 | T | 0.919 |
| -0.394 | 0.341 | T | 0.96 | 0.658 | D | 0.824 |
| -0.437 | 0.292 | T | 0.954 | 0.642 | D | 0.923 |
| -0.032 | 0.683 | D | 0.961 | 0.659 | D | 0.859 |
| 0.048 | 0.735 | D | 0.091 | 0.114 | T | 0.982 |
| 0.242 | 0.853 | D | 0.81 | 0.47 | D | 0.91 |
| -0.147 | 0.595 | T | 0.135 | 0.159 | T | 0.975 |
| -0.361 | 0.379 | T | 0.842 | 0.495 | D | 0.924 |

| | | | | | | |
|---|---|---|---|---|---|---|
| -0.405 | 0.329 | T | 0.038 | 0.033 | T | 0.904 |
| -0.377 | 0.36 | T | 0.395 | 0.287 | T | 0.693 |
| -0.36 | 0.38 | T | 0.054 | 0.061 | T | 0.949 |
| 0.17 | 0.814 | D | 0.51 | 0.33 | D | 0.911 |
| 0.118 | 0.781 | D | 0.445 | 0.306 | T | 0.966 |
| 0.66 | 0.994 | D | . | . | . | . |
| . | . | . | . | . | . | . |
| -0.289 | 0.459 | T | 0.251 | 0.229 | T | 0.939 |
| 0.307 | 0.885 | D | 0.419 | 0.297 | T | 0.948 |
| -0.244 | 0.504 | T | 0.265 | 0.235 | T | 0.924 |
| -0.194 | 0.552 | T | 0.445 | 0.306 | T | 0.975 |
| -0.325 | 0.42 | T | 0.211 | 0.21 | T | 0.99 |
| -0.419 | 0.312 | T | 0.29 | 0.246 | T | 0.743 |
| . | . | . | . | . | . | . |
| -0.431 | 0.298 | T | 0.044 | 0.045 | T | 0.792 |
| -0.252 | 0.496 | T | 0.632 | 0.377 | D | 0.964 |
| 0.259 | 0.86 | D | 0.996 | 0.885 | D | 0.925 |
| -0.31 | 0.436 | T | 0.169 | 0.185 | T | 0.832 |
| -0.11 | 0.626 | T | 0.974 | 0.702 | D | 0.731 |
| -0.041 | 0.676 | D | 0.034 | 0.027 | T | 0.854 |
| -0.218 | 0.529 | T | 0.151 | 0.172 | T | 0.862 |
| . | . | . | . | . | . | . |
| -0.405 | 0.329 | T | 0.038 | 0.033 | T | 0.904 |
| -0.279 | 0.469 | T | 0.067 | 0.082 | T | 0.972 |
| -0.183 | 0.562 | T | 0.937 | 0.607 | D | 0.89 |
| 0.02 | 0.716 | D | 0.109 | 0.134 | T | 0.941 |
| -0.08 | 0.649 | T | 0.301 | 0.251 | T | 0.876 |
| -0.319 | 0.427 | T | 0.117 | 0.141 | T | 0.894 |
| -0.278 | 0.47 | T | 0.035 | 0.027 | T | 0.953 |
| . | . | . | . | . | . | . |
| 0.148 | 0.801 | D | 0.989 | 0.792 | D | 0.948 |
| -0.168 | 0.576 | T | 0.87 | 0.52 | D | 0.852 |
| 0.031 | 0.723 | D | 0.933 | 0.599 | D | 0.892 |
| 0.418 | 0.927 | D | 0.091 | 0.113 | T | 0.954 |
| -0.405 | 0.329 | T | 0.038 | 0.033 | T | 0.904 |
| -0.405 | 0.329 | T | 0.038 | 0.033 | T | 0.904 |
| -0.063 | 0.661 | T | 0.738 | 0.426 | D | 0.926 |
| -0.067 | 0.658 | T | 0.258 | 0.232 | T | 0.842 |
| -0.485 | 0.239 | T | 0.812 | 0.472 | D | 0.885 |
| 0.242 | 0.853 | D | 0.985 | 0.763 | D | 0.753 |
| 0.026 | 0.72 | D | 0.252 | 0.229 | T | 0.794 |
| -0.147 | 0.595 | T | 0.027 | 0.015 | T | 0.929 |
| -0.069 | 0.657 | T | 0.631 | 0.377 | D | 0.909 |

| | | | | | | |
|---|---|---|---|---|---|---|
| -0.128 | 0.612 | T | 0.445 | 0.306 | T | 0.95 |
| . | . | . | . | . | . | . |
| -0.273 | 0.475 | T | 0.034 | 0.027 | T | 0.865 |
| 0.051 | 0.736 | D | 0.992 | 0.827 | D | 0.879 |
| 0.307 | 0.885 | D | 0.419 | 0.297 | T | 0.948 |
| -0.083 | 0.647 | T | 0.99 | 0.802 | D | 0.871 |
| -0.306 | 0.441 | T | 0.58 | 0.356 | D | 0.879 |
| -0.405 | 0.329 | T | 0.038 | 0.033 | T | 0.904 |
| 0.092 | 0.764 | D | 0.851 | 0.502 | D | 0.984 |
| 0.312 | 0.887 | D | 0.999 | 0.951 | D | 0.935 |
| . | . | . | . | . | . | . |
| -0.121 | 0.617 | T | 0.044 | 0.043 | T | 0.952 |
| 0.095 | 0.766 | D | 0.979 | 0.726 | D | 0.994 |
| . | . | . | . | . | . | . |
| . | . | . | . | . | . | . |
| 0.418 | 0.927 | D | 0.091 | 0.113 | T | 0.954 |
| . | . | . | . | . | . | . |
| 0.04 | 0.729 | D | 0.527 | 0.336 | D | 0.813 |
| -0.103 | 0.631 | T | 0.331 | 0.263 | T | 0.941 |
| . | . | . | . | . | . | . |
| 0.241 | 0.852 | D | 0.988 | 0.787 | D | 0.936 |
| -0.128 | 0.611 | T | 0.962 | 0.661 | D | 0.853 |
| . | . | . | . | . | . | . |
| -0.405 | 0.329 | T | 0.038 | 0.033 | T | 0.904 |
| 0.02 | 0.716 | D | 0.109 | 0.134 | T | 0.941 |
| -0.06 | 0.663 | T | 0.154 | 0.174 | T | 0.948 |
| -0.18 | 0.565 | T | 0.022 | 0.009 | T | 0.927 |
| . | . | . | . | . | . | . |
| -0.444 | 0.284 | T | 0.265 | 0.235 | T | 0.761 |
| 0.13 | 0.789 | D | 0.982 | 0.743 | D | 0.967 |
| -0.196 | 0.55 | T | 0.185 | 0.196 | T | 0.969 |
| -0.181 | 0.564 | T | 0.079 | 0.098 | T | 0.835 |
| -0.279 | 0.469 | T | 0.04 | 0.037 | T | 0.794 |
| -0.181 | 0.564 | T | 0.132 | 0.155 | T | 0.898 |
| -0.041 | 0.676 | D | 0.034 | 0.027 | T | 0.854 |
| -0.142 | 0.6 | T | 0.774 | 0.447 | D | 0.921 |
| -0.026 | 0.686 | D | 0.572 | 0.353 | D | 0.908 |
| -0.049 | 0.671 | D | 0.528 | 0.337 | D | 0.944 |
| 0.046 | 0.733 | D | 0.757 | 0.437 | D | 0.815 |
| 0.323 | 0.892 | D | 0.24 | 0.224 | T | 0.925 |
| . | . | . | . | . | . | . |
| . | . | . | . | . | . | . |
| -0.196 | 0.55 | T | 0.185 | 0.196 | T | 0.969 |

| | | | | | | |
|---|---|---|---|---|---|---|
| . | . | . | . | . | . | . |
| -0.405 | 0.329 | T | 0.038 | 0.033 | T | 0.904 |
| -0.041 | 0.676 | D | 0.034 | 0.027 | T | 0.854 |
| -0.377 | 0.361 | T | 0.567 | 0.351 | D | 0.987 |
| -0.217 | 0.53 | T | 0.499 | 0.326 | T | 0.971 |
| 0.142 | 0.797 | D | 0.999 | 0.969 | D | 0.98 |
| 0.347 | 0.902 | D | 0.316 | 0.256 | T | 0.981 |
| 0.049 | 0.735 | D | 0.628 | 0.375 | D | 0.889 |
| 0.191 | 0.826 | D | 0.993 | 0.833 | D | 0.986 |
| 0.285 | 0.874 | D | . | . | . | . |
| 0.66 | 0.994 | D | . | . | . | . |
| 0.185 | 0.823 | D | 0.996 | 0.889 | D | 0.964 |
| -0.147 | 0.595 | T | 0.089 | 0.112 | T | 0.834 |
| . | . | . | . | . | . | . |
| -0.388 | 0.348 | T | 0.035 | 0.028 | T | 0.936 |
| -0.269 | 0.479 | T | 0.965 | 0.672 | D | 0.789 |
| 0.283 | 0.873 | D | 0.796 | 0.461 | D | 0.941 |
| . | . | . | . | . | . | . |
| -0.18 | 0.565 | T | 0.022 | 0.009 | T | 0.927 |
| -0.386 | 0.35 | T | 0.113 | 0.137 | T | 0.854 |
| 0.3 | 0.882 | D | 0.486 | 0.321 | T | 0.972 |
| 0.347 | 0.902 | D | 0.316 | 0.256 | T | 0.981 |
| . | . | . | . | . | . | . |
| -0.136 | 0.604 | T | 0.113 | 0.138 | T | 0.946 |
| -0.296 | 0.451 | T | 0.916 | 0.573 | D | 0.798 |
| -0.072 | 0.655 | T | 0.241 | 0.225 | T | 0.839 |
| -0.425 | 0.305 | T | 0.161 | 0.18 | T | 0.704 |
| -0.455 | 0.271 | T | 0.13 | 0.154 | T | 0.683 |
| -0.317 | 0.429 | T | 0.139 | 0.162 | T | 0.824 |
| -0.164 | 0.58 | T | 0.226 | 0.218 | T | 0.964 |
| -0.278 | 0.47 | T | 0.035 | 0.027 | T | 0.953 |
| 0.347 | 0.902 | D | 0.316 | 0.256 | T | 0.981 |
| . | . | . | . | . | . | . |
| -0.027 | 0.685 | D | 0.784 | 0.453 | D | 0.974 |
| -0.311 | 0.435 | T | 0.034 | 0.026 | T | 0.899 |
| -0.121 | 0.617 | T | 0.938 | 0.607 | D | 0.98 |
| 0.165 | 0.811 | D | 0.139 | 0.162 | T | 0.973 |
| 0.075 | 0.753 | D | 0.919 | 0.578 | D | 0.871 |
| . | . | . | . | . | . | . |
| -0.278 | 0.47 | T | 0.035 | 0.027 | T | 0.953 |
| 0.227 | 0.845 | D | 0.992 | 0.828 | D | 0.968 |
| 0.014 | 0.713 | D | 0.112 | 0.136 | T | 0.75 |
| 0.364 | 0.909 | D | 0.812 | 0.472 | D | 0.792 |

| | | | | | | |
|---|---|---|---|---|---|---|
| -0.187 | 0.558 | T | 0.446 | 0.306 | T | 0.874 |
| 0.199 | 0.831 | D | 0.876 | 0.526 | D | 0.972 |
| . | . | . | . | . | . | . |
| . | . | . | . | . | . | . |
| -0.425 | 0.305 | T | 0.161 | 0.18 | T | 0.704 |
| -0.328 | 0.417 | T | 0.763 | 0.44 | D | 0.761 |
| 0.108 | 0.774 | D | 0.303 | 0.251 | T | 0.868 |
| 0.3 | 0.882 | D | 0.486 | 0.321 | T | 0.972 |
| -0.399 | 0.336 | T | 0.462 | 0.312 | T | 0.918 |
| . | . | . | . | . | . | . |
| -0.176 | 0.569 | T | 0.148 | 0.17 | T | 0.891 |
| -0.15 | 0.592 | T | 0.026 | 0.014 | T | 0.799 |
| -0.269 | 0.48 | T | 0.994 | 0.853 | D | 0.969 |
| -0.269 | 0.48 | T | 0.994 | 0.853 | D | 0.969 |
| . | . | . | . | . | . | . |
| . | . | . | . | . | . | . |
| 0.336 | 0.898 | D | 1 | 0.99 | D | 0.97 |
| -0.167 | 0.577 | T | 0.651 | 0.385 | D | 0.846 |
| 0.029 | 0.722 | D | 0.893 | 0.544 | D | 0.953 |
| . | . | . | . | . | . | . |
| -0.136 | 0.604 | T | 0.113 | 0.138 | T | 0.946 |
| 0.236 | 0.85 | D | 0.254 | 0.231 | T | 0.875 |
| 0.092 | 0.763 | D | 0.13 | 0.154 | T | 0.971 |
| -0.36 | 0.38 | T | 0.982 | 0.743 | D | 0.951 |
| 0.252 | 0.857 | D | 0.377 | 0.281 | T | 0.859 |
| -0.337 | 0.407 | T | 0.119 | 0.143 | T | 0.942 |
| . | . | . | . | . | . | . |
| -0.176 | 0.569 | T | 0.148 | 0.17 | T | 0.891 |
| -0.208 | 0.538 | T | 0.072 | 0.089 | T | 0.867 |
| -0.405 | 0.329 | T | 0.038 | 0.033 | T | 0.904 |
| -0.278 | 0.47 | T | 0.035 | 0.027 | T | 0.953 |
| -0.15 | 0.592 | T | 0.026 | 0.014 | T | 0.799 |
| -0.278 | 0.47 | T | 0.035 | 0.027 | T | 0.953 |
| -0.041 | 0.676 | D | 0.034 | 0.027 | T | 0.854 |
| -0.136 | 0.605 | T | 0.952 | 0.638 | D | 0.765 |
| -0.191 | 0.555 | T | 0.451 | 0.308 | T | 0.898 |
| -0.147 | 0.595 | T | 0.135 | 0.159 | T | 0.975 |
| 0.092 | 0.763 | D | 0.13 | 0.154 | T | 0.971 |
| -0.36 | 0.38 | T | 0.054 | 0.061 | T | 0.949 |
| 0.27 | 0.866 | D | 0.989 | 0.792 | D | 0.981 |
| . | . | . | . | . | . | . |
| -0.304 | 0.443 | T | 0.813 | 0.473 | D | 0.976 |
| -0.311 | 0.435 | T | 0.034 | 0.026 | T | 0.899 |

| | | | | | | |
|---|---|---|---|---|---|---|
| . | . | . | . | . | . | . |
| -0.278 | 0.47 | T | 0.035 | 0.027 | T | 0.953 |
| 0.092 | 0.763 | D | 0.13 | 0.154 | T | 0.971 |
| -0.311 | 0.435 | T | 0.034 | 0.026 | T | 0.899 |
| -0.425 | 0.305 | T | 0.161 | 0.18 | T | 0.704 |
| -0.311 | 0.435 | T | 0.034 | 0.026 | T | 0.899 |
| -0.47 | 0.255 | T | 0.836 | 0.49 | D | 0.924 |
| -0.168 | 0.576 | T | 0.765 | 0.442 | D | 0.88 |
| . | . | . | . | . | . | . |
| 0.165 | 0.811 | D | 0.139 | 0.162 | T | 0.973 |
| -0.147 | 0.595 | T | 0.135 | 0.159 | T | 0.975 |
| -0.395 | 0.34 | T | 0.048 | 0.051 | T | 0.728 |
| . | . | . | . | . | . | . |
| -0.089 | 0.642 | T | 0.835 | 0.489 | D | 0.771 |
| -0.027 | 0.685 | D | 0.784 | 0.453 | D | 0.974 |
| -0.098 | 0.636 | T | 0.978 | 0.723 | D | 0.933 |
| -0.271 | 0.477 | T | 0.283 | 0.243 | T | 0.863 |
| . | . | . | . | . | . | . |
| 0.033 | 0.725 | D | 0.294 | 0.248 | T | 0.968 |
| -0.252 | 0.496 | T | 0.315 | 0.256 | T | 0.956 |
| -0.271 | 0.477 | T | 0.283 | 0.243 | T | 0.863 |
| -0.065 | 0.659 | T | 0.212 | 0.211 | T | 0.761 |
| . | . | . | . | . | . | . |
| 0.037 | 0.727 | D | 0.981 | 0.736 | D | 0.989 |
| . | . | . | . | . | . | . |
| 0.168 | 0.813 | D | 0.219 | 0.214 | T | 0.971 |
| -0.226 | 0.522 | T | 0.886 | 0.536 | D | 0.865 |
| 0.66 | 0.994 | D | . | . | . | . |
| . | . | . | . | . | . | . |
| -0.066 | 0.659 | T | 0.466 | 0.314 | T | 0.933 |
| 0.32 | 0.891 | D | 0.707 | 0.41 | D | 0.944 |
| 0.048 | 0.734 | D | 0.732 | 0.423 | D | 0.94 |
| 0.212 | 0.838 | D | 0.958 | 0.65 | D | 0.938 |
| 0.013 | 0.712 | D | 0.829 | 0.485 | D | 0.974 |
| . | . | . | . | . | . | . |
| -0.366 | 0.373 | T | 0.145 | 0.167 | T | 0.803 |
| 0.048 | 0.734 | D | 0.732 | 0.423 | D | 0.94 |
| 0.015 | 0.713 | D | 1 | 0.977 | D | 0.89 |
| -0.31 | 0.436 | T | 0.312 | 0.255 | T | 0.966 |
| -0.489 | 0.235 | T | 0.09 | 0.112 | T | 0.822 |
| -0.049 | 0.671 | D | 0.501 | 0.327 | D | 0.978 |
| 0.284 | 0.874 | D | 0.204 | 0.207 | T | 0.965 |
| -0.165 | 0.579 | T | 0.814 | 0.473 | D | 0.957 |

| | | | | | | |
|---|---|---|---|---|---|---|
| . | . | . | . | . | . | . |
| 0.168 | 0.813 | D | 0.219 | 0.214 | T | 0.971 |
| -0.003 | 0.701 | D | 0.238 | 0.223 | T | 0.937 |
| 0.16 | 0.808 | D | 0.382 | 0.283 | T | 0.974 |
| -0.184 | 0.561 | T | 0.316 | 0.256 | T | 0.946 |
| -0.258 | 0.49 | T | 0.962 | 0.662 | D | 0.956 |
| 0.041 | 0.73 | D | 0.994 | 0.845 | D | 0.989 |
| . | . | . | . | . | . | . |
| -0.562 | 0.162 | T | 0.037 | 0.032 | T | 0.892 |
| 0.063 | 0.745 | D | 0.61 | 0.368 | D | 0.595 |
| -0.147 | 0.595 | T | 0.089 | 0.112 | T | 0.834 |
| -0.167 | 0.577 | T | 0.879 | 0.529 | D | 0.746 |
| 0.463 | 0.939 | D | 0.996 | 0.877 | D | 0.985 |
| 0.065 | 0.746 | D | 0.11 | 0.135 | T | 0.895 |
| 0.166 | 0.812 | D | 0.961 | 0.658 | D | 0.843 |
| -0.293 | 0.454 | T | 0.211 | 0.21 | T | 0.763 |
| -0.404 | 0.329 | T | 0.209 | 0.209 | T | 0.982 |
| 0.004 | 0.706 | D | 0.066 | 0.081 | T | 0.959 |
| -0.487 | 0.237 | T | 0.101 | 0.125 | T | 0.757 |
| -0.404 | 0.329 | T | 0.209 | 0.209 | T | 0.982 |
| -0.177 | 0.568 | T | 0.14 | 0.163 | T | 0.792 |
| 0.041 | 0.73 | D | 0.332 | 0.263 | T | 0.952 |
| -0.065 | 0.659 | T | 0.212 | 0.211 | T | 0.761 |
| -0.404 | 0.329 | T | 0.209 | 0.209 | T | 0.982 |
| -0.366 | 0.373 | T | 0.209 | 0.209 | T | 0.908 |
| . | . | . | . | . | . | . |
| 0.004 | 0.706 | D | 0.066 | 0.081 | T | 0.959 |
| -0.153 | 0.589 | T | 0.095 | 0.118 | T | 0.909 |
| -0.271 | 0.477 | T | 0.283 | 0.243 | T | 0.863 |
| -0.364 | 0.376 | T | 0.12 | 0.144 | T | 0.89 |
| -0.116 | 0.622 | T | 0.728 | 0.421 | D | 0.897 |
| 0.158 | 0.807 | D | 0.994 | 0.843 | D | 0.979 |
| 0.041 | 0.73 | D | 0.994 | 0.845 | D | 0.989 |
| -0.404 | 0.329 | T | 0.209 | 0.209 | T | 0.982 |
| -0.153 | 0.589 | T | 0.095 | 0.118 | T | 0.909 |
| -0.31 | 0.436 | T | 0.312 | 0.255 | T | 0.966 |
| -0.202 | 0.545 | T | 0.872 | 0.522 | D | 0.898 |
| . | . | . | . | . | . | . |
| 0.14 | 0.795 | D | 0.96 | 0.657 | D | 0.906 |
| 0.298 | 0.881 | D | 0.999 | 0.954 | D | 0.956 |
| 0.192 | 0.827 | D | 0.152 | 0.172 | T | 0.971 |
| 0.16 | 0.808 | D | 0.382 | 0.283 | T | 0.974 |
| -0.404 | 0.329 | T | 0.209 | 0.209 | T | 0.982 |

| | | | | | | |
|---|---|---|---|---|---|---|
| -0.58 | 0.146 | T | 0.149 | 0.17 | T | 0.884 |
| -0.153 | 0.589 | T | 0.095 | 0.118 | T | 0.909 |
| -0.123 | 0.615 | T | 0.903 | 0.556 | D | 0.932 |

| LIST.S2_ranks | LIST.S2_pred | Aloft_pred | Aloft_Confide | CADD_raw | CADD_raw_ra | CADD_phred |
|---|---|---|---|---|---|---|
| . | . | . | . | . | . | NA |
| . | . | . | . | . | . | NA |
| 0.93 | D | .;.;.;.;.;.; | .;.;.;.;.;.; | 3.608 | 0.669 | 25.1 |
| 0.635 | D | .;.;.;.;.; | .;.;.;.;.; | 3.387 | 0.612 | 24.4 |
| 0.817 | D | .;.;.; | .;.;.; | 3.703 | 0.694 | 25.4 |
| 0.765 | D | .;.;.;.; | .;.;.;.; | 3.649 | 0.68 | 25.2 |
| 0.963 | D | .;.;.; | .;.;.; | 3.381 | 0.611 | 24.3 |
| 0.721 | D | .;.;.;.;.;.;.;.; | .;.;.;.;.;.;.;.; | 3.948 | 0.769 | 26.6 |
| 0.826 | D | .;.;.;.;.;.;.; | .;.;.;.;.;.;.; | 4.408 | 0.902 | 31 |
| 0.732 | D | .;.;.;.;.; | .;.;.;.;.; | 2.699 | 0.456 | 22.8 |
| 0.795 | D | .;.;.;.;.;.; | .;.;.;.;.;.; | 4.106 | 0.823 | 27.8 |
| 0.93 | D | .;.;.;.;.;.; | .;.;.;.;.;.; | 3.608 | 0.669 | 25.1 |
| 0.729 | D | .;.;.;.; | .;.;.;.; | 3.684 | 0.689 | 25.3 |
| 0.903 | D | .;.; | .;.; | 4.418 | 0.904 | 31 |
| . | . | . | . | . | . | NA |
| 0.738 | D | .;.; | .;.; | 3.141 | 0.552 | 23.7 |
| 0.733 | D | .;.;.; | .;.;.; | 3.134 | 0.551 | 23.7 |
| 0.911 | D | .; | .; | 3.994 | 0.784 | 26.9 |
| 0.795 | D | .;.;.;.;.;.; | .;.;.;.;.;.; | 4.106 | 0.823 | 27.8 |
| 0.806 | D | .;.;.;.; | .;.;.;.; | 3.088 | 0.54 | 23.6 |
| 0.894 | D | .;.;.;.;.; | .;.;.;.;.; | 4.048 | 0.803 | 27.3 |
| . | . | . | . | . | . | NA |
| 0.22 | T | .;.; | .;.; | 3.048 | 0.531 | 23.5 |
| 0.935 | D | .;.;.;.;.;.;.; | .;.;.;.;.;.;.; | 3.618 | 0.672 | 25.1 |
| . | . | Recessive;Re | High;High;.;.; | 7.169 | 0.973 | 37 |
| 0.92 | D | .;.;.;.;.;.; | .;.;.;.;.;.; | 4.039 | 0.799 | 27.3 |
| 0.817 | D | .;.;.;.; | .;.;.;.; | 3.483 | 0.636 | 24.7 |
| 0.493 | T | .;.; | .;.; | 2.123 | 0.331 | 20.3 |
| 0.207 | T | .; | .; | 3.177 | 0.56 | 23.8 |
| 0.893 | D | .; | .; | 3.817 | 0.727 | 25.9 |
| 0.795 | D | .;.;.;.;.;.; | .;.;.;.;.;.; | 4.106 | 0.823 | 27.8 |
| 0.337 | T | .;.; | .;.; | 3.629 | 0.675 | 25.1 |
| 0.788 | D | .;.;.; | .;.;.; | 3.257 | 0.58 | 24 |
| 0.732 | D | .;.;.;.;.; | .;.;.;.;.; | 2.699 | 0.456 | 22.8 |
| 0.709 | D | .;.;.; | .;.;.; | 3.558 | 0.656 | 24.9 |
| 0.485 | T | .;.;.; | .;.;.; | 3.266 | 0.582 | 24 |
| 0.718 | D | .;.;.;.;.; | .;.;.;.;.; | 3.997 | 0.785 | 27 |
| 0.562 | D | .;.; | .;.; | 4.211 | 0.858 | 28.8 |
| 0.939 | D | .;.; | .;.; | 3.569 | 0.659 | 24.9 |
| 0.682 | D | .;.;.;.;.; | .;.;.;.;.; | 5.968 | 0.957 | 35 |
| 0.913 | D | .;.;.;.;.;.;.;.; | .;.;.;.;.;.;.;.; | 3.466 | 0.632 | 24.6 |
| 0.722 | D | .;.;.; | .;.;.; | 3.396 | 0.614 | 24.4 |

| | | | | | | |
|---|---|---|---|---|---|---|
| 0.93 | D | .;.;.;.;.;.; | .;.;.;.;.;.; | 3.608 | 0.669 | 25.1 |
| 0.302 | T | .;.;.; | .;.;.; | 2.249 | 0.358 | 21.2 |
| 0.804 | D | .;.; | .;.; | 4.102 | 0.821 | 27.8 |
| 0.684 | D | .;.; | .;.; | 2.992 | 0.519 | 23.4 |
| 0.894 | D | .;.;.;.;.; | .;.;.;.;.; | 4.048 | 0.803 | 27.3 |
| . | . | Recessive;.; | High;.; | 8.632 | 0.995 | 44 |
| . | . | . | . | . | . | NA |
| 0.793 | D | .;.;.;.;.;.;.;.; | .;.;.;.;.;.;.;.; | 3.146 | 0.553 | 23.7 |
| 0.802 | D | .;.;.;.;.;.;.; | .;.;.;.;.;.;.; | 4.385 | 0.899 | 31 |
| 0.77 | D | .;.;.;.; | .;.;.;.; | 3.378 | 0.61 | 24.3 |
| 0.91 | D | .;.; | .;.; | 3.369 | 0.608 | 24.3 |
| 0.969 | D | .; | .; | 3.022 | 0.525 | 23.5 |
| 0.363 | T | .;.;.; | .;.;.; | 3.401 | 0.616 | 24.4 |
| . | . | . | . | . | . | NA |
| 0.433 | T | .;.;.; | .;.;.; | 2.207 | 0.349 | 20.9 |
| 0.889 | D | .;.; | .;.; | 4.013 | 0.791 | 27.1 |
| 0.724 | D | .;.; | .;.; | 3.91 | 0.756 | 26.4 |
| 0.5 | T | .;.;.; | .;.;.; | 2.938 | 0.507 | 23.3 |
| 0.346 | T | .;.; | .;.; | 3.795 | 0.721 | 25.8 |
| 0.54 | D | .; | .; | 2.684 | 0.452 | 22.8 |
| 0.564 | D | .;.; | .;.; | 2.985 | 0.517 | 23.4 |
| . | . | . | . | . | . | NA |
| 0.93 | D | .;.;.;.;.;.; | .;.;.;.;.;.; | 3.608 | 0.669 | 25.1 |
| 0.897 | D | .; | .; | 3.054 | 0.532 | 23.5 |
| 0.624 | D | .;.; | .;.; | 2.942 | 0.508 | 23.3 |
| 0.795 | D | .;.;.;.;.;.; | .;.;.;.;.;.; | 4.106 | 0.823 | 27.8 |
| 0.588 | D | .; | .; | 3.685 | 0.689 | 25.3 |
| 0.632 | D | .; | .; | 3.505 | 0.642 | 24.7 |
| 0.95 | D | .;.;.;.;.;.;.;.; | .;.;.;.;.;.;.;.; | 4.152 | 0.839 | 28.2 |
| . | . | . | . | . | . | NA |
| 0.809 | D | .;.;.;.;.; | .;.;.;.;.; | 3.921 | 0.76 | 26.5 |
| 0.537 | D | .;.;.;.; | .;.;.;.; | 3.682 | 0.689 | 25.3 |
| 0.627 | D | .;.; | .;.; | 3.329 | 0.597 | 24.2 |
| 0.826 | D | .;.;.;.;.;.;.; | .;.;.;.;.;.;.; | 4.408 | 0.902 | 31 |
| 0.93 | D | .;.;.;.;.;.; | .;.;.;.;.;.; | 3.608 | 0.669 | 25.1 |
| 0.93 | D | .;.;.;.;.;.; | .;.;.;.;.;.; | 3.608 | 0.669 | 25.1 |
| 0.751 | D | .;.;.; | .;.;.; | 3.85 | 0.737 | 26.1 |
| 0.582 | T | .;.;.;.;.;.;.;.;.; | .;.;.;.;.;.;.;.;.; | 3.983 | 0.78 | 26.9 |
| 0.609 | D | .;.; | .;.; | 2.797 | 0.477 | 23 |
| 0.379 | T | .;.; | .;.; | 3.944 | 0.767 | 26.6 |
| 0.436 | T | .;.; | .;.; | 3.465 | 0.632 | 24.6 |
| 0.738 | D | .;.; | .;.; | 3.141 | 0.552 | 23.7 |
| 0.677 | D | .;.;.;.; | .;.;.;.; | 4.076 | 0.812 | 27.6 |

| | | | | | | |
|---|---|---|---|---|---|---|
| 0.809 | D | .;.;.;.;.;.; | .;.;.;.;.;.; | 3.698 | 0.693 | 25.4 |
| . | . | . | . | . | . | NA |
| 0.564 | D | .;.;.;.;.; | .;.;.;.;.; | 4.043 | 0.801 | 27.3 |
| 0.597 | D | .;.;.;.;.;.; | .;.;.;.;.;.; | 4.269 | 0.875 | 29.3 |
| 0.802 | D | .;.;.;.;.;.;.; | .;.;.;.;.;.;.; | 4.385 | 0.899 | 31 |
| 0.576 | D | .; | .; | 3.907 | 0.755 | 26.4 |
| 0.908 | D | .;.;.;.; | .;.;.;.; | 2.722 | 0.461 | 22.9 |
| 0.93 | D | .;.;.;.;.;.; | .;.;.;.;.;.; | 3.608 | 0.669 | 25.1 |
| 0.966 | D | .;.;.;.;.;.;.;.;.;.;. | .;.;.;.;.;.;.;.;.;.;. | 3.985 | 0.781 | 26.9 |
| 0.756 | D | .;.; | .;.; | 4.069 | 0.81 | 27.5 |
| . | . | . | . | . | . | NA |
| 0.817 | D | .;.;.; | .;.;.; | 3.703 | 0.694 | 25.4 |
| 0.987 | D | .;.;.;.;.; | .;.;.;.;.; | 4.345 | 0.892 | 29.9 |
| . | . | . | . | . | . | NA |
| . | . | . | . | . | . | NA |
| 0.826 | D | .;.;.;.;.;.;.; | .;.;.;.;.;.;.; | 4.408 | 0.902 | 31 |
| . | . | . | . | . | . | NA |
| 0.466 | T | .; | .; | 3.231 | 0.573 | 23.9 |
| 0.781 | D | .;.;.; | .;.;.; | 3.355 | 0.604 | 24.3 |
| . | . | . | . | . | . | NA |
| 0.76 | D | .;.; | .;.; | 4.318 | 0.886 | 29.7 |
| 0.539 | D | .; | .; | 3.68 | 0.688 | 25.3 |
| . | . | . | . | . | . | NA |
| 0.93 | D | .;.;.;.;.;.; | .;.;.;.;.;.; | 3.608 | 0.669 | 25.1 |
| 0.795 | D | .;.;.;.;.;.; | .;.;.;.;.;.; | 4.106 | 0.823 | 27.8 |
| 0.799 | D | .;.;.; | .;.;.; | 3.073 | 0.536 | 23.6 |
| 0.732 | D | .;.;.;.;.; | .;.;.;.;.; | 2.699 | 0.456 | 22.8 |
| . | . | . | . | . | . | NA |
| 0.386 | T | .;.; | .;.; | 2.996 | 0.52 | 23.4 |
| 0.905 | D | .;.;.; | .;.;.; | 3.56 | 0.656 | 24.9 |
| 0.886 | D | .; | .; | 4.254 | 0.871 | 29.2 |
| 0.505 | T | .;.;.; | .;.;.; | 3.593 | 0.665 | 25 |
| 0.435 | T | .;.; | .;.; | 3.126 | 0.549 | 23.7 |
| 0.643 | D | .;.;.;.;.;.;.;.;.;.;. | .;.;.;.;.;.;.;.;.;.;. | 2.133 | 0.333 | 20.4 |
| 0.54 | D | .; | .; | 2.684 | 0.452 | 22.8 |
| 0.714 | D | .;.; | .;.; | 3.477 | 0.635 | 24.6 |
| 0.673 | D | .;.; | .;.; | 4.145 | 0.837 | 28.2 |
| 0.788 | D | .;.; | .;.; | 3.825 | 0.729 | 26 |
| 0.469 | T | .; | .; | 3.653 | 0.681 | 25.2 |
| 0.777 | D | .;.;.;.;.;.;.; | .;.;.;.;.;.;.; | 4.102 | 0.822 | 27.8 |
| . | . | . | . | . | . | NA |
| . | . | . | . | . | . | NA |
| 0.886 | D | .; | .; | 4.254 | 0.871 | 29.2 |

| | | | | | | |
|---|---|---|---|---|---|---|
| . | . | . | . | . | . | NA |
| 0.93 | D | .;.;.;.;.;.; | .;.;.;.;.;.; | 3.608 | 0.669 | 25.1 |
| 0.54 | D | .; | .; | 2.684 | 0.452 | 22.8 |
| 0.955 | D | .; | .; | 2.392 | 0.389 | 22.1 |
| 0.894 | D | .;.;.; | .;.;.; | 2.555 | 0.425 | 22.5 |
| 0.93 | D | .;.; | .;.; | 4.179 | 0.848 | 28.5 |
| 0.934 | D | .;.; | .;.; | 3.791 | 0.719 | 25.8 |
| 0.621 | D | .;.; | .;.; | 4.866 | 0.929 | 33 |
| 0.952 | D | .;.;.; | .;.;.; | 3.879 | 0.746 | 26.2 |
| . | . | .;.;.;.;.;.;.;.; | .;.;.;.;.;.;.;.; | 5.738 | 0.952 | 34 |
| . | . | Dominant;.;.; | High;.;.;.;.;.;.;. | 7.994 | 0.987 | 40 |
| 0.867 | D | .;.;.;.;.;.;.;.; | .;.;.;.;.;.;.;.; | 3.88 | 0.746 | 26.2 |
| 0.504 | T | .;.;.; | .;.;.; | 2.218 | 0.351 | 21 |
| . | . | . | . | . | . | NA |
| 0.762 | D | .;.;.; | .;.;.; | 2.597 | 0.434 | 22.6 |
| 0.429 | T | .; | .; | 2.897 | 0.498 | 23.2 |
| 0.778 | D | .;.;.;.;.;.;.;.;.;.;. | .;.;.;.;.;.;.;.;.;.;. | 4.024 | 0.794 | 27.1 |
| . | . | . | . | . | . | NA |
| 0.732 | D | .;.;.;.;.; | .;.;.;.;.; | 2.699 | 0.456 | 22.8 |
| 0.54 | D | .;.; | .;.; | 2.528 | 0.418 | 22.5 |
| 0.899 | D | .;.;.; | .;.;.; | 3.852 | 0.738 | 26.1 |
| 0.934 | D | .;.; | .;.; | 3.791 | 0.719 | 25.8 |
| . | . | . | . | . | . | NA |
| 0.793 | D | .;.;.; | .;.;.; | 3.368 | 0.607 | 24.3 |
| 0.442 | T | .;.;.;.; | .;.;.;.; | 3.733 | 0.703 | 25.5 |
| 0.512 | T | .; | .; | 3.111 | 0.545 | 23.7 |
| 0.314 | T | .;.;.; | .;.;.; | 2.375 | 0.385 | 22 |
| 0.304 | T | .;.;.; | .;.;.; | 2.549 | 0.423 | 22.5 |
| 0.485 | T | .;.; | .;.; | 3.295 | 0.589 | 24.1 |
| 0.866 | D | .;.;.; | .;.;.; | 4.019 | 0.792 | 27.1 |
| 0.95 | D | .;.;.;.;.;.;.;.; | .;.;.;.;.;.;.;.; | 4.152 | 0.839 | 28.2 |
| 0.934 | D | .;.; | .;.; | 3.791 | 0.719 | 25.8 |
| . | . | . | . | . | . | NA |
| 0.908 | D | .;.;.;.;.; | .;.;.;.;.; | 4.185 | 0.85 | 28.5 |
| 0.648 | D | .; | .; | 3.238 | 0.575 | 24 |
| 0.93 | D | .;.;.;.;.;.;.; | .;.;.;.;.;.;.; | 4.014 | 0.791 | 27.1 |
| 0.904 | D | .;.; | .;.; | 3.816 | 0.727 | 25.9 |
| 0.576 | D | .;.;.; | .;.;.; | 3.954 | 0.77 | 26.7 |
| . | . | . | . | . | . | NA |
| 0.95 | D | .;.;.;.;.;.;.;.; | .;.;.;.;.;.;.;.; | 4.152 | 0.839 | 28.2 |
| 0.884 | D | .; | .; | 4.323 | 0.887 | 29.8 |
| 0.372 | T | .;.;.; | .;.;.; | 3.611 | 0.67 | 25.1 |
| 0.438 | T | .;.;.;.;.; | .;.;.;.;.; | 4.392 | 0.9 | 31 |

| | | | | | | |
|---|---|---|---|---|---|---|
| 0.661 | D | .;.;.; | .;.;.; | 3.548 | 0.653 | 24.9 |
| 0.897 | D | .;.; | .;.; | 4.274 | 0.876 | 29.3 |
| . | . | . | . | . | . | NA |
| . | . | . | . | . | . | NA |
| 0.314 | T | .;.;.; | .;.;.; | 2.375 | 0.385 | 22 |
| 0.386 | T | .; | .; | 4.108 | 0.824 | 27.8 |
| 0.57 | D | .;.;.; | .;.;.; | 3.631 | 0.675 | 25.1 |
| 0.899 | D | .;.;.; | .;.;.; | 3.852 | 0.738 | 26.1 |
| 0.704 | D | .;.;.; | .;.;.; | 3.915 | 0.758 | 26.4 |
| . | . | . | . | . | . | NA |
| 0.626 | D | .;.;.;.;.;.;.;.; | .;.;.;.;.;.;.;.; | 4.46 | 0.909 | 32 |
| 0.444 | T | .;.;.; | .;.;.; | 2.953 | 0.51 | 23.3 |
| 0.888 | D | .;.;.;.; | .;.;.;.; | 3.651 | 0.68 | 25.2 |
| 0.888 | D | .;.;.;.; | .;.;.;.; | 3.651 | 0.68 | 25.2 |
| . | . | . | . | . | . | NA |
| . | . | . | . | . | . | NA |
| 0.891 | D | .;.; | .;.; | 4.54 | 0.916 | 32 |
| 0.525 | T | .;.; | .;.; | 2.321 | 0.373 | 21.7 |
| 0.822 | D | .;.; | .;.; | 4.419 | 0.904 | 31 |
| . | . | . | . | . | . | NA |
| 0.793 | D | .;.;.; | .;.;.; | 3.368 | 0.607 | 24.3 |
| 0.586 | D | .;.;.; | .;.;.; | 3.682 | 0.689 | 25.3 |
| 0.903 | D | .;.; | .;.; | 4.418 | 0.904 | 31 |
| 0.826 | D | .;.;.; | .;.;.; | 3.841 | 0.734 | 26 |
| 0.55 | D | .;.;.; | .;.;.; | 2.834 | 0.485 | 23.1 |
| 0.782 | D | .;.;.; | .;.;.; | 2.899 | 0.499 | 23.2 |
| . | . | . | . | . | . | NA |
| 0.626 | D | .;.;.;.;.;.;.;.; | .;.;.;.;.;.;.;.; | 4.46 | 0.909 | 32 |
| 0.568 | D | .;.;.; | .;.;.; | 2.828 | 0.483 | 23.1 |
| 0.93 | D | .;.;.;.;.;.; | .;.;.;.;.;.; | 3.608 | 0.669 | 25.1 |
| 0.95 | D | .;.;.;.;.;.;.;.; | .;.;.;.;.;.;.;.; | 4.152 | 0.839 | 28.2 |
| 0.444 | T | .;.;.; | .;.;.; | 2.953 | 0.51 | 23.3 |
| 0.95 | D | .;.;.;.;.;.;.;.; | .;.;.;.;.;.;.;.; | 4.152 | 0.839 | 28.2 |
| 0.54 | D | .; | .; | 2.684 | 0.452 | 22.8 |
| 0.393 | T | .; | .; | 3.904 | 0.754 | 26.4 |
| 0.659 | D | .;.;.; | .;.;.; | 3.327 | 0.597 | 24.2 |
| 0.913 | D | .;.;.;.;.;.;.;.; | .;.;.;.;.;.;.;.; | 3.466 | 0.632 | 24.6 |
| 0.903 | D | .;.; | .;.; | 4.418 | 0.904 | 31 |
| 0.804 | D | .;.; | .;.; | 4.102 | 0.821 | 27.8 |
| 0.934 | D | .; | .; | 3.032 | 0.527 | 23.5 |
| . | . | . | . | . | . | NA |
| 0.968 | D | .;.;.;.;.;.;.; | .;.;.;.;.;.;.; | 3.19 | 0.564 | 23.8 |
| 0.648 | D | .; | .; | 3.238 | 0.575 | 24 |

| | | | | | | | |
|---|---|---|---|---|---|---|---|
| . | | . | . | . | . | . | NA |
| | 0.95 | D | .;.;.;.;.;.;.;.; | .;.;.;.;.;.;.;.; | 4.152 | 0.839 | 28.2 |
| | 0.903 | D | .;.; | .;.; | 4.418 | 0.904 | 31 |
| | 0.648 | D | .; | .; | 3.238 | 0.575 | 24 |
| | 0.314 | T | .;.;.; | .;.;.; | 2.375 | 0.385 | 22 |
| | 0.648 | D | .; | .; | 3.238 | 0.575 | 24 |
| | 0.722 | D | .; | .; | 2.679 | 0.451 | 22.8 |
| | 0.596 | D | .;.;.; | .;.;.; | 3.403 | 0.616 | 24.4 |
| . | | . | . | . | . | . | NA |
| | 0.904 | D | .;.; | .;.; | 3.816 | 0.727 | 25.9 |
| | 0.913 | D | .;.;.;.;.;.;.;.; | .;.;.;.;.;.;.;.; | 3.466 | 0.632 | 24.6 |
| | 0.343 | T | .;.; | .;.; | 2.725 | 0.461 | 22.9 |
| . | | . | . | . | . | . | NA |
| | 0.401 | T | .;.;.; | .;.;.; | 2.885 | 0.496 | 23.2 |
| | 0.908 | D | .;.;.;.;.; | .;.;.;.;.; | 4.185 | 0.85 | 28.5 |
| | 0.769 | D | .;.; | .;.; | 3.276 | 0.584 | 24.1 |
| | 0.559 | D | .; | .; | 3.552 | 0.654 | 24.9 |
| . | | . | . | . | . | . | NA |
| | 0.885 | D | .;.;.;.;.;.; | .;.;.;.;.;.; | 4.704 | 0.924 | 32 |
| | 0.832 | D | .;.; | .;.; | 3.514 | 0.645 | 24.8 |
| | 0.559 | D | .; | .; | 3.552 | 0.654 | 24.9 |
| | 0.386 | T | .;.;.; | .;.;.; | 3.898 | 0.752 | 26.3 |
| . | | . | . | . | . | . | NA |
| | 0.962 | D | .;.; | .;.; | 3.861 | 0.741 | 26.1 |
| . | | . | . | . | . | . | NA |
| | 0.923 | D | .;.;.;.;.; | .;.;.;.;.; | 4.169 | 0.845 | 28.4 |
| | 0.563 | D | .;.; | .;.; | 3.593 | 0.665 | 25 |
| . | | . | Dominant;.;.; | High;.;.;.;.;.;.;. | 8.746 | 0.995 | 45 |
| . | | . | . | . | . | . | NA |
| | 0.749 | D | .; | .; | 2.714 | 0.459 | 22.9 |
| | 0.97 | D | .;.;.;.; | .;.;.;.; | 3.404 | 0.616 | 24.4 |
| | 0.773 | D | .;.; | .;.; | 4.358 | 0.894 | 30 |
| | 0.766 | D | .;.;.; | .;.;.; | 4.427 | 0.905 | 31 |
| | 0.906 | D | .;.; | .;.; | 4.175 | 0.846 | 28.4 |
| . | | . | . | . | . | . | NA |
| | 0.45 | T | .;.;.; | .;.;.; | 2.961 | 0.512 | 23.3 |
| | 0.773 | D | .;.; | .;.; | 4.358 | 0.894 | 30 |
| | 0.622 | D | .;.;.;.;.; | .;.;.;.;.; | 4.268 | 0.874 | 29.3 |
| | 0.875 | D | .;.;.;.;.;.; | .;.;.;.;.;.; | 3.571 | 0.659 | 24.9 |
| | 0.481 | T | .; | .; | 2.595 | 0.433 | 22.6 |
| | 0.921 | D | .;.; | .;.; | 3.71 | 0.696 | 25.4 |
| | 0.872 | D | .; | .; | 3.928 | 0.762 | 26.5 |
| | 0.839 | D | .;.;.; | .;.;.; | 3.833 | 0.732 | 26 |

| | | | | | | |
|---|---|---|---|---|---|---|
| . | . | . | . | . | . | NA |
| 0.923 | D | .;.;.;.;.; | .;.;.;.;.; | 4.169 | 0.845 | 28.4 |
| 0.769 | D | .;.;.; | .;.;.; | 3.348 | 0.602 | 24.3 |
| 0.908 | D | .;.;.;.;.; | .;.;.;.;.; | 4.267 | 0.874 | 29.3 |
| 0.825 | D | .;.; | .;.; | 4.054 | 0.805 | 27.4 |
| 0.835 | D | .;.;.; | .;.;.; | 3.926 | 0.761 | 26.5 |
| 0.963 | D | .;.;.;.;.; | .;.;.;.;.; | 3.957 | 0.772 | 26.7 |
| . | . | . | . | . | . | NA |
| 0.634 | D | .;.;.;.; | .;.;.;.; | 2.178 | 0.342 | 20.7 |
| 0.221 | T | .;.; | .;.; | 2.194 | 0.346 | 20.8 |
| 0.504 | T | .;.;.; | .;.;.; | 2.218 | 0.351 | 21 |
| 0.366 | T | .;.; | .;.; | 3.839 | 0.734 | 26 |
| 0.952 | D | .;.;.; | .;.;.; | 4.238 | 0.866 | 29 |
| 0.635 | D | .;.; | .;.; | 3.439 | 0.625 | 24.5 |
| 0.52 | T | .; | .; | 3.803 | 0.723 | 25.8 |
| 0.391 | T | .;.;.; | .;.;.; | 3.179 | 0.561 | 23.8 |
| 0.958 | D | .;.;.;.;.;.;.;.;.;.;. | .;.;.;.;.;.;.;.;.;.;. | 4.007 | 0.789 | 27 |
| 0.845 | D | .; | .; | 3.63 | 0.675 | 25.1 |
| 0.38 | T | .;.; | .;.; | 2.805 | 0.478 | 23 |
| 0.958 | D | .;.;.;.;.;.;.;.;.;.;. | .;.;.;.;.;.;.;.;.;.;. | 4.007 | 0.789 | 27 |
| 0.436 | T | .;.;.; | .;.;.; | 2.599 | 0.434 | 22.6 |
| 0.817 | D | .;.; | .;.; | 3.581 | 0.662 | 25 |
| 0.386 | T | .;.;.; | .;.;.; | 3.898 | 0.752 | 26.3 |
| 0.958 | D | .;.;.;.;.;.;.;.;.;.;. | .;.;.;.;.;.;.;.;.;.;. | 4.007 | 0.789 | 27 |
| 0.687 | D | .;.;.;.; | .;.;.;.; | 2.465 | 0.405 | 22.3 |
| . | . | . | . | . | . | NA |
| 0.845 | D | .; | .; | 3.63 | 0.675 | 25.1 |
| 0.699 | D | .;.;.;.; | .;.;.;.; | 3.893 | 0.751 | 26.3 |
| 0.559 | D | .; | .; | 3.552 | 0.654 | 24.9 |
| 0.623 | D | .;.; | .;.; | 3.282 | 0.586 | 24.1 |
| 0.707 | D | .;.;.;.; | .;.;.;.; | 3.97 | 0.776 | 26.8 |
| 0.929 | D | .;.; | .;.; | 4.533 | 0.915 | 32 |
| 0.963 | D | .;.;.;.;.; | .;.;.;.;.; | 3.957 | 0.772 | 26.7 |
| 0.958 | D | .;.;.;.;.;.;.;.;.;.;. | .;.;.;.;.;.;.;.;.;.;. | 4.007 | 0.789 | 27 |
| 0.699 | D | .;.;.;.; | .;.;.;.; | 3.893 | 0.751 | 26.3 |
| 0.875 | D | .;.;.;.;.;.; | .;.;.;.;.;.; | 3.571 | 0.659 | 24.9 |
| 0.642 | D | .;.; | .;.; | 3.515 | 0.645 | 24.8 |
| . | . | . | . | . | . | NA |
| 0.67 | D | .; | .; | 3.214 | 0.569 | 23.9 |
| 0.832 | D | .;.;.; | .;.;.; | 3.809 | 0.725 | 25.9 |
| 0.903 | D | .;.;.;.;.; | .;.;.;.;.; | 3.875 | 0.745 | 26.2 |
| 0.908 | D | .;.;.;.;.; | .;.;.;.;.; | 4.267 | 0.874 | 29.3 |
| 0.958 | D | .;.;.;.;.;.;.;.;.;.;. | .;.;.;.;.;.;.;.;.;.;. | 4.007 | 0.789 | 27 |

| | | | | | | |
|---|---|---|---|---|---|---|
| 0.728 | D | .;.; | .;.; | 3.151 | 0.554 | 23.7 |
| 0.699 | D | .;.;.;.; | .;.;.;.; | 3.893 | 0.751 | 26.3 |
| 0.747 | D | .;.;.;.;.; | .;.;.;.;.; | 4.603 | 0.92 | 32 |

| DANN_score | DANN_ranksc | fathmm.MKL_ | fathmm.MKL_ | fathmm.MKL_ | fathmm.XF_c | fathmm.XF_c |
|---|---|---|---|---|---|---|
| . | . | . | . | . | . | . |
| . | . | . | . | . | . | . |
| 0.999 | 0.935 | 0.833 | 0.425 | D | 0.76 | 0.698 |
| 0.987 | 0.447 | 0.972 | 0.732 | D | 0.969 | 0.992 |
| 0.999 | 0.933 | 0.658 | 0.329 | D | 0.327 | 0.428 |
| 0.993 | 0.591 | 0.913 | 0.533 | D | 0.668 | 0.636 |
| 0.998 | 0.862 | 0.63 | 0.32 | D | 0.747 | 0.689 |
| 0.999 | 0.985 | 0.986 | 0.846 | D | 0.92 | 0.891 |
| 0.999 | 0.993 | 0.976 | 0.758 | D | 0.904 | 0.854 |
| 0.987 | 0.446 | 0.981 | 0.797 | D | 0.869 | 0.79 |
| 0.996 | 0.718 | 0.991 | 0.911 | D | 0.785 | 0.716 |
| 0.999 | 0.935 | 0.833 | 0.425 | D | 0.76 | 0.698 |
| 0.996 | 0.74 | 0.902 | 0.512 | D | 0.431 | 0.493 |
| 0.996 | 0.729 | 0.996 | 0.978 | D | 0.852 | 0.769 |
| . | . | . | . | . | . | . |
| 0.988 | 0.459 | 0.857 | 0.448 | D | 0.526 | 0.548 |
| 0.955 | 0.271 | 0.909 | 0.524 | D | 0.557 | 0.567 |
| 0.997 | 0.837 | 0.989 | 0.884 | D | 0.962 | 0.984 |
| 0.996 | 0.718 | 0.991 | 0.911 | D | 0.785 | 0.716 |
| 0.995 | 0.71 | 0.925 | 0.56 | D | 0.52 | 0.545 |
| 0.999 | 0.985 | 0.985 | 0.84 | D | 0.927 | 0.908 |
| . | . | . | . | . | . | . |
| 0.989 | 0.484 | 0.993 | 0.938 | D | 0.675 | 0.64 |
| 0.998 | 0.919 | 0.974 | 0.745 | D | 0.946 | 0.955 |
| 0.992 | 0.547 | 0.989 | 0.89 | D | 0.346 | 0.441 |
| 0.999 | 0.999 | 0.988 | 0.873 | D | 0.759 | 0.697 |
| 0.998 | 0.89 | 0.981 | 0.797 | D | 0.939 | 0.938 |
| 0.904 | 0.197 | 0.609 | 0.313 | D | 0.494 | 0.53 |
| 0.993 | 0.587 | 0.975 | 0.753 | D | 0.661 | 0.631 |
| 0.999 | 0.986 | 0.992 | 0.925 | D | 0.88 | 0.806 |
| 0.996 | 0.718 | 0.991 | 0.911 | D | 0.785 | 0.716 |
| 0.997 | 0.837 | 0.904 | 0.516 | D | 0.379 | 0.462 |
| 0.999 | 0.975 | 0.975 | 0.752 | D | 0.787 | 0.717 |
| 0.987 | 0.446 | 0.981 | 0.797 | D | 0.869 | 0.79 |
| 0.999 | 0.999 | 0.79 | 0.39 | D | 0.217 | 0.342 |
| 0.997 | 0.824 | 0.993 | 0.942 | D | 0.47 | 0.516 |
| 0.998 | 0.922 | 0.99 | 0.9 | D | 0.732 | 0.679 |
| 0.99 | 0.501 | 0.806 | 0.402 | D | 0.732 | 0.679 |
| 0.999 | 0.992 | 0.961 | 0.675 | D | 0.637 | 0.616 |
| 0.998 | 0.905 | 0.928 | 0.565 | D | 0.861 | 0.779 |
| 0.999 | 0.999 | 0.91 | 0.527 | D | 0.828 | 0.747 |
| 0.999 | 0.998 | 0.95 | 0.63 | D | 0.512 | 0.54 |

| | | | | | | |
|---|---|---|---|---|---|---|
| 0.999 | 0.935 | 0.833 | 0.425 | D | 0.76 | 0.698 |
| 0.982 | 0.395 | 0.742 | 0.363 | D | 0.56 | 0.569 |
| 0.999 | 0.999 | 0.974 | 0.748 | D | 0.868 | 0.787 |
| 0.986 | 0.439 | 0.971 | 0.729 | D | 0.676 | 0.641 |
| 0.999 | 0.985 | 0.985 | 0.84 | D | 0.927 | 0.908 |
| 0.997 | 0.837 | 0.998 | 0.997 | D | 0.33 | 0.43 |
| . | . | . | . | . | . | . |
| 0.99 | 0.491 | 0.848 | 0.438 | D | 0.581 | 0.581 |
| 0.992 | 0.559 | 0.961 | 0.677 | D | 0.969 | 0.992 |
| 0.999 | 0.984 | 0.965 | 0.693 | D | 0.626 | 0.609 |
| 0.999 | 0.957 | 0.962 | 0.68 | D | 0.207 | 0.333 |
| 0.999 | 0.937 | 0.992 | 0.93 | D | 0.915 | 0.88 |
| 0.997 | 0.8 | 0.856 | 0.448 | D | 0.131 | 0.25 |
| . | . | . | . | . | . | . |
| 0.946 | 0.252 | 0.37 | 0.257 | N | 0.094 | 0.19 |
| 0.999 | 0.989 | 0.997 | 0.985 | D | 0.871 | 0.792 |
| 0.994 | 0.624 | 0.995 | 0.966 | D | 0.951 | 0.966 |
| 0.994 | 0.643 | 0.954 | 0.645 | D | 0.371 | 0.457 |
| 0.994 | 0.641 | 0.72 | 0.352 | D | 0.295 | 0.405 |
| 0.998 | 0.87 | 0.938 | 0.593 | D | 0.452 | 0.506 |
| 0.997 | 0.796 | 0.974 | 0.743 | D | 0.924 | 0.901 |
| . | . | . | . | . | . | . |
| 0.999 | 0.935 | 0.833 | 0.425 | D | 0.76 | 0.698 |
| 0.998 | 0.927 | 0.983 | 0.814 | D | 0.859 | 0.776 |
| 0.998 | 0.928 | 0.826 | 0.418 | D | 0.497 | 0.532 |
| 0.996 | 0.718 | 0.991 | 0.911 | D | 0.785 | 0.716 |
| 0.998 | 0.917 | 0.993 | 0.941 | D | 0.631 | 0.612 |
| 0.999 | 0.992 | 0.99 | 0.899 | D | 0.95 | 0.963 |
| 0.998 | 0.848 | 0.953 | 0.642 | D | 0.947 | 0.958 |
| . | . | . | . | . | . | . |
| 0.999 | 0.96 | 0.914 | 0.535 | D | 0.6 | 0.593 |
| 0.999 | 0.936 | 0.817 | 0.41 | D | 0.716 | 0.668 |
| 0.999 | 0.936 | 0.949 | 0.629 | D | 0.586 | 0.584 |
| 0.999 | 0.993 | 0.976 | 0.758 | D | 0.904 | 0.854 |
| 0.999 | 0.935 | 0.833 | 0.425 | D | 0.76 | 0.698 |
| 0.999 | 0.935 | 0.833 | 0.425 | D | 0.76 | 0.698 |
| 0.917 | 0.21 | 0.961 | 0.678 | D | 0.892 | 0.828 |
| 0.997 | 0.798 | 0.998 | 0.992 | D | 0.929 | 0.913 |
| 0.998 | 0.888 | 0.925 | 0.559 | D | 0.374 | 0.459 |
| 0.998 | 0.875 | 0.97 | 0.719 | D | 0.803 | 0.728 |
| 0.999 | 0.962 | 0.953 | 0.643 | D | 0.966 | 0.988 |
| 0.988 | 0.459 | 0.857 | 0.448 | D | 0.526 | 0.548 |
| 0.999 | 0.983 | 0.941 | 0.601 | D | 0.867 | 0.786 |

| | | | | | | |
|---|---|---|---|---|---|---|
| 1 | 1 | 0.964 | 0.688 | D | 0.824 | 0.744 |
| . | . | . | . | . | . | . |
| 0.996 | 0.737 | 0.551 | 0.298 | D | 0.236 | 0.358 |
| 0.998 | 0.905 | 0.949 | 0.626 | D | 0.931 | 0.919 |
| 0.992 | 0.559 | 0.961 | 0.677 | D | 0.969 | 0.992 |
| 0.997 | 0.799 | 0.99 | 0.896 | D | 0.9 | 0.844 |
| 0.998 | 0.874 | 0.934 | 0.582 | D | 0.538 | 0.555 |
| 0.999 | 0.935 | 0.833 | 0.425 | D | 0.76 | 0.698 |
| 0.998 | 0.91 | 0.992 | 0.923 | D | 0.943 | 0.947 |
| 0.999 | 0.962 | 0.95 | 0.632 | D | 0.927 | 0.91 |
| . | . | . | . | . | . | . |
| 0.999 | 0.933 | 0.658 | 0.329 | D | 0.327 | 0.428 |
| 0.999 | 0.962 | 0.982 | 0.805 | D | 0.968 | 0.991 |
| . | . | . | . | . | . | . |
| . | . | . | . | . | . | . |
| 0.999 | 0.993 | 0.976 | 0.758 | D | 0.904 | 0.854 |
| . | . | . | . | . | . | . |
| 0.965 | 0.299 | 0.987 | 0.852 | D | 0.751 | 0.692 |
| 0.981 | 0.386 | 0.95 | 0.632 | D | 0.371 | 0.457 |
| . | . | . | . | . | . | . |
| 0.996 | 0.72 | 0.998 | 0.995 | D | 0.962 | 0.984 |
| 0.999 | 0.934 | 0.989 | 0.877 | D | 0.605 | 0.596 |
| . | . | . | . | . | . | . |
| 0.999 | 0.935 | 0.833 | 0.425 | D | 0.76 | 0.698 |
| 0.996 | 0.718 | 0.991 | 0.911 | D | 0.785 | 0.716 |
| 0.96 | 0.284 | 0.935 | 0.584 | D | 0.752 | 0.692 |
| 0.987 | 0.446 | 0.981 | 0.797 | D | 0.869 | 0.79 |
| . | . | . | . | . | . | . |
| 0.974 | 0.34 | 0.457 | 0.276 | N | 0.084 | 0.17 |
| 0.995 | 0.679 | 0.854 | 0.446 | D | 0.237 | 0.359 |
| 0.999 | 0.998 | 0.969 | 0.715 | D | 0.771 | 0.706 |
| 0.999 | 0.963 | 0.988 | 0.869 | D | 0.869 | 0.789 |
| 0.912 | 0.205 | 0.966 | 0.701 | D | 0.655 | 0.627 |
| 0.997 | 0.785 | 0.974 | 0.747 | D | 0.269 | 0.385 |
| 0.998 | 0.87 | 0.938 | 0.593 | D | 0.452 | 0.506 |
| 0.999 | 0.974 | 0.969 | 0.718 | D | 0.688 | 0.649 |
| 0.999 | 0.998 | 0.928 | 0.565 | D | 0.698 | 0.656 |
| 0.999 | 0.973 | 0.998 | 0.996 | D | 0.908 | 0.862 |
| 0.998 | 0.85 | 0.984 | 0.825 | D | 0.866 | 0.785 |
| 0.985 | 0.42 | 0.98 | 0.79 | D | 0.966 | 0.989 |
| . | . | . | . | . | . | . |
| . | . | . | . | . | . | . |
| 0.999 | 0.998 | 0.969 | 0.715 | D | 0.771 | 0.706 |

| | | | | | | |
|---|---|---|---|---|---|---|
| . | . | . | . | . | . | . |
| 0.999 | 0.935 | 0.833 | 0.425 | D | 0.76 | 0.698 |
| 0.998 | 0.87 | 0.938 | 0.593 | D | 0.452 | 0.506 |
| 0.999 | 0.935 | 0.998 | 0.994 | D | 0.962 | 0.984 |
| 0.983 | 0.398 | 0.925 | 0.558 | D | 0.495 | 0.531 |
| 0.999 | 0.95 | 0.903 | 0.514 | D | 0.932 | 0.923 |
| 0.998 | 0.889 | 0.971 | 0.729 | D | 0.976 | 0.997 |
| 0.999 | 0.993 | 0.992 | 0.933 | D | 0.846 | 0.763 |
| 0.999 | 0.945 | 0.965 | 0.693 | D | 0.916 | 0.882 |
| 0.994 | 0.63 | 0.977 | 0.767 | D | . | . |
| 0.998 | 0.872 | 0.918 | 0.543 | D | 0.153 | 0.278 |
| 0.999 | 0.993 | 0.946 | 0.617 | D | 0.824 | 0.744 |
| 0.995 | 0.686 | 0.977 | 0.766 | D | 0.707 | 0.662 |
| . | . | . | . | . | . | . |
| 0.985 | 0.428 | 0.779 | 0.383 | D | 0.513 | 0.541 |
| 0.99 | 0.512 | 0.875 | 0.471 | D | 0.276 | 0.391 |
| 0.995 | 0.671 | 0.997 | 0.991 | D | 0.948 | 0.959 |
| . | . | . | . | . | . | . |
| 0.987 | 0.446 | 0.981 | 0.797 | D | 0.869 | 0.79 |
| 0.997 | 0.836 | 0.958 | 0.66 | D | 0.492 | 0.529 |
| 0.999 | 0.979 | 0.951 | 0.635 | D | 0.479 | 0.521 |
| 0.998 | 0.889 | 0.971 | 0.729 | D | 0.976 | 0.997 |
| . | . | . | . | . | . | . |
| 0.83 | 0.145 | 0.908 | 0.522 | D | 0.512 | 0.541 |
| 0.996 | 0.771 | 0.991 | 0.907 | D | 0.872 | 0.793 |
| 0.996 | 0.749 | 0.982 | 0.807 | D | 0.642 | 0.619 |
| 0.994 | 0.614 | 0.841 | 0.432 | D | 0.415 | 0.484 |
| 0.994 | 0.649 | 0.954 | 0.647 | D | 0.309 | 0.415 |
| 0.99 | 0.496 | 0.978 | 0.769 | D | 0.584 | 0.583 |
| 0.977 | 0.355 | 0.975 | 0.751 | D | 0.887 | 0.819 |
| 0.998 | 0.848 | 0.953 | 0.642 | D | 0.947 | 0.958 |
| 0.998 | 0.889 | 0.971 | 0.729 | D | 0.976 | 0.997 |
| . | . | . | . | . | . | . |
| 0.999 | 0.966 | 0.999 | 0.998 | D | 0.921 | 0.895 |
| 0.997 | 0.774 | 0.958 | 0.66 | D | 0.538 | 0.556 |
| 0.981 | 0.382 | 0.988 | 0.865 | D | 0.959 | 0.979 |
| 0.993 | 0.6 | 0.98 | 0.787 | D | 0.916 | 0.881 |
| 0.999 | 0.979 | 0.976 | 0.756 | D | 0.836 | 0.754 |
| . | . | . | . | . | . | . |
| 0.998 | 0.848 | 0.953 | 0.642 | D | 0.947 | 0.958 |
| 0.998 | 0.888 | 0.994 | 0.949 | D | 0.965 | 0.987 |
| 0.989 | 0.488 | 0.923 | 0.553 | D | 0.281 | 0.394 |
| 0.998 | 0.883 | 0.993 | 0.939 | D | 0.917 | 0.883 |

| | | | | | | |
|---|---|---|---|---|---|---|
| 0.999 | 0.938 | 0.863 | 0.456 | D | 0.415 | 0.484 |
| 0.998 | 0.856 | 0.983 | 0.815 | D | 0.985 | 1 |
| . | . | . | . | . | . | . |
| . | . | . | . | . | . | . |
| 0.994 | 0.614 | 0.841 | 0.432 | D | 0.415 | 0.484 |
| 0.999 | 0.997 | 0.971 | 0.724 | D | 0.823 | 0.743 |
| 0.975 | 0.341 | 0.998 | 0.993 | D | 0.898 | 0.841 |
| 0.999 | 0.979 | 0.951 | 0.635 | D | 0.479 | 0.521 |
| 0.999 | 0.932 | 0.97 | 0.724 | D | 0.871 | 0.792 |
| . | . | . | . | . | . | . |
| 0.999 | 0.98 | 0.989 | 0.888 | D | 0.816 | 0.738 |
| 0.974 | 0.339 | 0.905 | 0.518 | D | . | . |
| 0.997 | 0.785 | 0.916 | 0.539 | D | 0.816 | 0.738 |
| 0.997 | 0.785 | 0.916 | 0.539 | D | 0.816 | 0.738 |
| . | . | . | . | . | . | . |
| . | . | . | . | . | . | . |
| 0.999 | 0.999 | 0.981 | 0.795 | D | 0.916 | 0.882 |
| 0.957 | 0.275 | 0.995 | 0.965 | D | 0.959 | 0.979 |
| 1 | 1 | 0.983 | 0.816 | D | 0.89 | 0.825 |
| . | . | . | . | . | . | . |
| 0.83 | 0.145 | 0.908 | 0.522 | D | 0.512 | 0.541 |
| 0.982 | 0.394 | 0.965 | 0.693 | D | 0.917 | 0.884 |
| 0.996 | 0.729 | 0.996 | 0.978 | D | 0.852 | 0.769 |
| 0.998 | 0.878 | 0.99 | 0.897 | D | 0.877 | 0.801 |
| 0.895 | 0.189 | 0.971 | 0.725 | D | 0.899 | 0.842 |
| 0.988 | 0.461 | 0.959 | 0.667 | D | 0.849 | 0.766 |
| . | . | . | . | . | . | . |
| 0.999 | 0.98 | 0.989 | 0.888 | D | 0.816 | 0.738 |
| 0.922 | 0.216 | 0.863 | 0.456 | D | . | . |
| 0.999 | 0.935 | 0.833 | 0.425 | D | 0.76 | 0.698 |
| 0.998 | 0.848 | 0.953 | 0.642 | D | 0.947 | 0.958 |
| 0.974 | 0.339 | 0.905 | 0.518 | D | . | . |
| 0.998 | 0.848 | 0.953 | 0.642 | D | 0.947 | 0.958 |
| 0.998 | 0.87 | 0.938 | 0.593 | D | 0.452 | 0.506 |
| 0.998 | 0.895 | 0.987 | 0.863 | D | 0.94 | 0.942 |
| 0.973 | 0.331 | 0.951 | 0.635 | D | 0.738 | 0.683 |
| 0.999 | 0.999 | 0.91 | 0.527 | D | 0.828 | 0.747 |
| 0.996 | 0.729 | 0.996 | 0.978 | D | 0.852 | 0.769 |
| 0.999 | 0.999 | 0.974 | 0.748 | D | 0.868 | 0.787 |
| 0.999 | 0.985 | 0.991 | 0.918 | D | 0.957 | 0.976 |
| . | . | . | . | . | . | . |
| 0.998 | 0.918 | 0.954 | 0.647 | D | 0.602 | 0.594 |
| 0.997 | 0.774 | 0.958 | 0.66 | D | 0.538 | 0.556 |

| | | | | | | |
|---|---|---|---|---|---|---|
| . | . | . | . | . | . | . |
| 0.998 | 0.848 | 0.953 | 0.642 | D | 0.947 | 0.958 |
| 0.996 | 0.729 | 0.996 | 0.978 | D | 0.852 | 0.769 |
| 0.997 | 0.774 | 0.958 | 0.66 | D | 0.538 | 0.556 |
| 0.994 | 0.614 | 0.841 | 0.432 | D | 0.415 | 0.484 |
| 0.997 | 0.774 | 0.958 | 0.66 | D | 0.538 | 0.556 |
| 0.994 | 0.636 | 0.744 | 0.364 | D | 0.087 | 0.176 |
| 0.989 | 0.474 | 0.961 | 0.674 | D | 0.714 | 0.666 |
| . | . | . | . | . | . | . |
| 0.993 | 0.6 | 0.98 | 0.787 | D | 0.916 | 0.881 |
| 0.999 | 0.999 | 0.91 | 0.527 | D | 0.828 | 0.747 |
| 0.992 | 0.559 | 0.675 | 0.334 | D | 0.243 | 0.365 |
| . | . | . | . | . | . | . |
| 0.994 | 0.623 | 0.975 | 0.749 | D | 0.337 | 0.435 |
| 0.999 | 0.966 | 0.999 | 0.998 | D | 0.921 | 0.895 |
| 0.997 | 0.806 | 0.983 | 0.813 | D | 0.666 | 0.635 |
| 0.999 | 0.984 | 0.973 | 0.739 | D | 0.676 | 0.641 |
| . | . | . | . | . | . | . |
| 0.998 | 0.856 | 0.974 | 0.746 | D | 0.908 | 0.862 |
| 0.99 | 0.499 | 0.982 | 0.802 | D | 0.844 | 0.761 |
| 0.999 | 0.984 | 0.973 | 0.739 | D | 0.676 | 0.641 |
| 0.947 | 0.253 | 0.97 | 0.719 | D | 0.822 | 0.742 |
| . | . | . | . | . | . | . |
| 0.999 | 0.99 | 0.854 | 0.446 | D | 0.528 | 0.55 |
| . | . | . | . | . | . | . |
| 0.999 | 0.999 | 0.954 | 0.644 | D | 0.918 | 0.887 |
| 0.999 | 0.967 | 0.943 | 0.607 | D | 0.536 | 0.555 |
| 0.998 | 0.888 | 0.985 | 0.831 | D | 0.215 | 0.34 |
| . | . | . | . | . | . | . |
| 0.999 | 0.947 | 0.868 | 0.462 | D | 0.763 | 0.7 |
| 0.997 | 0.825 | 0.983 | 0.813 | D | 0.951 | 0.965 |
| 0.999 | 0.941 | 0.968 | 0.709 | D | 0.801 | 0.727 |
| 0.972 | 0.327 | 0.964 | 0.692 | D | . | . |
| 0.999 | 0.995 | 0.999 | 0.999 | D | 0.91 | 0.866 |
| . | . | . | . | . | . | . |
| 0.997 | 0.835 | 0.827 | 0.418 | D | 0.181 | 0.309 |
| 0.999 | 0.941 | 0.968 | 0.709 | D | 0.801 | 0.727 |
| 0.999 | 0.972 | 0.973 | 0.738 | D | 0.939 | 0.939 |
| 1 | 1 | 0.952 | 0.638 | D | 0.643 | 0.62 |
| 0.988 | 0.472 | 0.79 | 0.39 | D | . | . |
| 0.988 | 0.465 | 0.984 | 0.824 | D | 0.912 | 0.871 |
| 0.999 | 0.968 | 0.971 | 0.724 | D | 0.653 | 0.626 |
| 0.999 | 0.952 | 0.994 | 0.959 | D | 0.903 | 0.852 |

| | | | | | | |
|---|---|---|---|---|---|---|
| . | . | . | . | . | . | . |
| 0.999 | 0.999 | 0.954 | 0.644 | D | 0.918 | 0.887 |
| 0.996 | 0.751 | 0.985 | 0.836 | D | 0.688 | 0.649 |
| 0.996 | 0.763 | 0.965 | 0.695 | D | 0.917 | 0.883 |
| 0.999 | 0.964 | 0.959 | 0.668 | D | 0.928 | 0.912 |
| 0.992 | 0.544 | 0.969 | 0.715 | D | 0.898 | 0.841 |
| 0.998 | 0.928 | 0.84 | 0.431 | D | 0.808 | 0.732 |
| . | . | . | . | . | . | . |
| 0.984 | 0.407 | 0.828 | 0.42 | D | 0.34 | 0.437 |
| 0.995 | 0.678 | 0.968 | 0.713 | D | 0.752 | 0.692 |
| 0.995 | 0.686 | 0.977 | 0.766 | D | 0.707 | 0.662 |
| 0.985 | 0.419 | 0.958 | 0.661 | D | 0.701 | 0.658 |
| 0.996 | 0.74 | 0.992 | 0.923 | D | 0.941 | 0.944 |
| 0.997 | 0.808 | 0.991 | 0.918 | D | 0.925 | 0.904 |
| 0.982 | 0.394 | 0.975 | 0.749 | D | 0.857 | 0.774 |
| 0.939 | 0.239 | 0.955 | 0.649 | D | 0.908 | 0.864 |
| 0.99 | 0.496 | 0.994 | 0.952 | D | 0.952 | 0.966 |
| 0.999 | 0.986 | 0.946 | 0.616 | D | 0.249 | 0.369 |
| 0.996 | 0.74 | 0.891 | 0.493 | D | 0.649 | 0.624 |
| 0.99 | 0.496 | 0.994 | 0.952 | D | 0.952 | 0.966 |
| 0.948 | 0.256 | 0.856 | 0.447 | D | 0.404 | 0.478 |
| 1 | 1 | 0.967 | 0.702 | D | 0.772 | 0.706 |
| 0.947 | 0.253 | 0.97 | 0.719 | D | 0.822 | 0.742 |
| 0.99 | 0.496 | 0.994 | 0.952 | D | 0.952 | 0.966 |
| 0.993 | 0.586 | 0.858 | 0.45 | D | 0.211 | 0.336 |
| . | . | . | . | . | . | . |
| 0.999 | 0.986 | 0.946 | 0.616 | D | 0.249 | 0.369 |
| 0.999 | 0.996 | 0.949 | 0.627 | D | 0.512 | 0.54 |
| 0.999 | 0.984 | 0.973 | 0.739 | D | 0.676 | 0.641 |
| 0.945 | 0.249 | 0.992 | 0.925 | D | 0.722 | 0.672 |
| 0.991 | 0.525 | 0.893 | 0.497 | D | 0.571 | 0.575 |
| 0.997 | 0.839 | 0.983 | 0.817 | D | 0.914 | 0.876 |
| 0.998 | 0.928 | 0.84 | 0.431 | D | 0.808 | 0.732 |
| 0.99 | 0.496 | 0.994 | 0.952 | D | 0.952 | 0.966 |
| 0.999 | 0.996 | 0.949 | 0.627 | D | 0.512 | 0.54 |
| 1 | 1 | 0.952 | 0.638 | D | 0.643 | 0.62 |
| 0.999 | 0.964 | 0.947 | 0.619 | D | 0.94 | 0.94 |
| . | . | . | . | . | . | . |
| 0.996 | 0.764 | 0.963 | 0.685 | D | 0.781 | 0.713 |
| 0.994 | 0.625 | 0.99 | 0.9 | D | 0.938 | 0.938 |
| 0.999 | 0.984 | 0.962 | 0.68 | D | 0.945 | 0.953 |
| 0.996 | 0.763 | 0.965 | 0.695 | D | 0.917 | 0.883 |
| 0.99 | 0.496 | 0.994 | 0.952 | D | 0.952 | 0.966 |

| | | | | | | |
|---|---|---|---|---|---|---|
| 0.999 | 0.945 | 0.992 | 0.925 | D | 0.856 | 0.773 |
| 0.999 | 0.996 | 0.949 | 0.627 | D | 0.512 | 0.54 |
| 0.998 | 0.902 | 0.994 | 0.955 | D | 0.92 | 0.891 |

| fathmm.XF_c | Eigen.raw_co | Eigen.raw_co | Eigen.PC.raw | Eigen.PC.raw | GenoCanyon | GenoCanyon |
|---|---|---|---|---|---|---|
| . | . | . | . | . | . | . |
| . | . | . | . | . | . | . |
| D | 0.642 | 0.758 | 0.668 | 0.8 | 1 | 0.983 |
| D | 0.939 | 0.935 | 0.928 | 0.967 | 1 | 0.748 |
| N | 0.256 | 0.539 | 0.232 | 0.516 | 1 | 0.748 |
| D | 0.139 | 0.483 | 0.157 | 0.475 | 1 | 0.425 |
| D | 0.218 | 0.521 | 0.114 | 0.453 | 0.76 | 0.235 |
| D | 0.581 | 0.72 | 0.495 | 0.676 | 1 | 0.748 |
| D | 0.795 | 0.858 | 0.761 | 0.87 | 1 | 0.748 |
| D | 0.159 | 0.492 | 0.313 | 0.563 | 1 | 0.748 |
| D | 0.512 | 0.678 | 0.552 | 0.716 | 1 | 0.748 |
| D | 0.642 | 0.758 | 0.668 | 0.8 | 1 | 0.983 |
| N | 0.253 | 0.538 | 0.218 | 0.509 | 1 | 0.418 |
| D | 0.965 | 0.946 | 0.936 | 0.97 | 1 | 0.748 |
| . | . | . | . | . | . | . |
| D | 0.02 | 0.428 | 0.123 | 0.457 | 1 | 0.748 |
| D | -0.198 | 0.332 | -0.055 | 0.373 | 0.003 | 0.097 |
| D | 0.632 | 0.752 | 0.628 | 0.77 | 1 | 0.748 |
| D | 0.512 | 0.678 | 0.552 | 0.716 | 1 | 0.748 |
| D | -0.188 | 0.336 | -0.244 | 0.3 | 1 | 0.748 |
| D | 0.736 | 0.82 | 0.757 | 0.867 | 1 | 0.748 |
| . | . | . | . | . | . | . |
| D | 0.457 | 0.646 | 0.417 | 0.626 | 1 | 0.748 |
| D | 0.583 | 0.721 | 0.657 | 0.791 | 1 | 0.983 |
| N | 1.023 | 0.964 | 0.915 | 0.962 | 1 | 0.748 |
| D | 0.612 | 0.74 | 0.583 | 0.737 | 1 | 0.748 |
| D | 0.722 | 0.811 | 0.644 | 0.782 | 1 | 0.748 |
| N | -0.569 | 0.199 | -0.622 | 0.189 | 1 | 0.748 |
| D | -0.06 | 0.392 | 0.118 | 0.455 | 1 | 0.748 |
| D | 0.786 | 0.852 | 0.783 | 0.886 | 1 | 0.748 |
| D | 0.512 | 0.678 | 0.552 | 0.716 | 1 | 0.748 |
| N | 0.353 | 0.589 | 0.259 | 0.532 | 1 | 0.748 |
| D | 0.675 | 0.78 | 0.662 | 0.795 | 1 | 0.748 |
| D | 0.159 | 0.492 | 0.313 | 0.563 | 1 | 0.748 |
| N | 0.411 | 0.62 | 0.452 | 0.649 | 1 | 0.748 |
| N | 0.22 | 0.522 | 0.346 | 0.582 | 1 | 0.748 |
| D | 0.58 | 0.719 | 0.552 | 0.716 | 1 | 0.748 |
| D | 0.7 | 0.797 | 0.717 | 0.837 | 1 | 0.748 |
| D | 0.406 | 0.617 | 0.341 | 0.58 | 0.991 | 0.323 |
| D | 0.614 | 0.74 | 0.687 | 0.814 | 1 | 0.748 |
| D | 0.441 | 0.637 | 0.451 | 0.648 | 1 | 0.748 |
| D | 0.613 | 0.74 | 0.63 | 0.771 | 1 | 0.748 |

| | | | | | | |
|---|---|---|---|---|---|---|
| D | 0.642 | 0.758 | 0.668 | 0.8 | 1 | 0.983 |
| D | -0.33 | 0.28 | -0.27 | 0.291 | 1 | 0.748 |
| D | 0.689 | 0.789 | 0.722 | 0.84 | 1 | 0.748 |
| D | 0.082 | 0.456 | 0.175 | 0.485 | 1 | 0.748 |
| D | 0.736 | 0.82 | 0.757 | 0.867 | 1 | 0.748 |
| N | 1.132 | 0.987 | 0.992 | 0.985 | 1 | 0.748 |
| . | . | . | . | . | . | . |
| D | -0.212 | 0.327 | -0.124 | 0.344 | 1 | 0.748 |
| D | 0.854 | 0.894 | 0.811 | 0.906 | 1 | 0.748 |
| D | 0.532 | 0.69 | 0.578 | 0.733 | 1 | 0.748 |
| N | 0.371 | 0.598 | 0.387 | 0.608 | 1 | 0.467 |
| D | 0.112 | 0.47 | 0.253 | 0.528 | 1 | 0.983 |
| N | 0.27 | 0.546 | 0.163 | 0.478 | 0.999 | 0.383 |
| . | . | . | . | . | . | . |
| N | -0.06 | 0.392 | -0.093 | 0.357 | 1 | 0.501 |
| D | 0.829 | 0.879 | 0.857 | 0.934 | 1 | 0.748 |
| D | 0.846 | 0.889 | 0.81 | 0.905 | 1 | 0.983 |
| N | 0.387 | 0.607 | 0.453 | 0.649 | 1 | 0.748 |
| N | 0.399 | 0.614 | 0.396 | 0.613 | 1 | 0.518 |
| N | -0.081 | 0.382 | -0.072 | 0.366 | 1 | 0.748 |
| D | 0.567 | 0.711 | 0.506 | 0.684 | 1 | 0.748 |
| . | . | . | . | . | . | . |
| D | 0.642 | 0.758 | 0.668 | 0.8 | 1 | 0.983 |
| D | 0.294 | 0.558 | 0.378 | 0.602 | 1 | 0.748 |
| N | 0.381 | 0.604 | 0.427 | 0.633 | 1 | 0.422 |
| D | 0.512 | 0.678 | 0.552 | 0.716 | 1 | 0.748 |
| D | 0.532 | 0.69 | 0.582 | 0.736 | 1 | 0.748 |
| D | 0.163 | 0.494 | 0.314 | 0.563 | 1 | 0.748 |
| D | 0.78 | 0.848 | 0.743 | 0.857 | 1 | 0.748 |
| . | . | . | . | . | . | . |
| D | 0.424 | 0.627 | 0.451 | 0.648 | 0.995 | 0.338 |
| D | -0.033 | 0.404 | -0.099 | 0.354 | 1 | 0.748 |
| D | 0.189 | 0.507 | 0.274 | 0.54 | 1 | 0.748 |
| D | 0.795 | 0.858 | 0.761 | 0.87 | 1 | 0.748 |
| D | 0.642 | 0.758 | 0.668 | 0.8 | 1 | 0.983 |
| D | 0.642 | 0.758 | 0.668 | 0.8 | 1 | 0.983 |
| D | 0.584 | 0.722 | 0.638 | 0.777 | 1 | 0.748 |
| D | 0.644 | 0.76 | 0.671 | 0.802 | 1 | 0.748 |
| N | 0.398 | 0.613 | 0.42 | 0.628 | 1 | 0.748 |
| D | 0.532 | 0.69 | 0.56 | 0.721 | 1 | 0.748 |
| D | 0.431 | 0.631 | 0.49 | 0.673 | 1 | 0.748 |
| D | 0.02 | 0.428 | 0.123 | 0.457 | 1 | 0.748 |
| D | 0.402 | 0.616 | 0.291 | 0.55 | 1 | 0.748 |

| | | | | | | | |
|---|---|---|---|---|---|---|---|
| D | | 0.682 | 0.784 | 0.675 | 0.805 | 1 | 0.748 |
| . | . | . | . | . | . | . | |
| N | | 0.008 | 0.422 | 0.033 | 0.413 | 1 | 0.518 |
| D | | 0.796 | 0.859 | 0.767 | 0.874 | 1 | 0.983 |
| D | | 0.854 | 0.894 | 0.811 | 0.906 | 1 | 0.748 |
| D | | 0.838 | 0.884 | 0.816 | 0.909 | 1 | 0.748 |
| D | | 0.386 | 0.607 | 0.46 | 0.654 | 1 | 0.748 |
| D | | 0.642 | 0.758 | 0.668 | 0.8 | 1 | 0.983 |
| D | | 0.854 | 0.893 | 0.831 | 0.919 | 1 | 0.748 |
| D | | 0.792 | 0.856 | 0.696 | 0.82 | 1 | 0.748 |
| . | . | . | . | . | . | . | |
| N | | 0.256 | 0.539 | 0.232 | 0.516 | 1 | 0.748 |
| D | | 0.965 | 0.946 | 0.9 | 0.956 | 1 | 0.983 |
| . | . | . | . | . | . | . | |
| . | . | . | . | . | . | . | |
| D | | 0.795 | 0.858 | 0.761 | 0.87 | 1 | 0.748 |
| . | . | . | . | . | . | . | |
| D | | 0.405 | 0.617 | 0.465 | 0.657 | 1 | 0.748 |
| N | | 0.203 | 0.514 | 0.312 | 0.562 | 1 | 0.459 |
| . | . | . | . | . | . | . | |
| D | | 0.817 | 0.872 | 0.846 | 0.928 | 1 | 0.748 |
| D | | 0.679 | 0.782 | 0.65 | 0.786 | 1 | 0.748 |
| . | . | . | . | . | . | . | |
| D | | 0.642 | 0.758 | 0.668 | 0.8 | 1 | 0.983 |
| D | | 0.512 | 0.678 | 0.552 | 0.716 | 1 | 0.748 |
| D | | 0.187 | 0.506 | 0.383 | 0.605 | 1 | 0.748 |
| D | | 0.159 | 0.492 | 0.313 | 0.563 | 1 | 0.748 |
| . | . | . | . | . | . | . | |
| N | | -0.251 | 0.311 | -0.273 | 0.29 | 1 | 0.748 |
| N | | 0.604 | 0.734 | 0.528 | 0.699 | 0.653 | 0.222 |
| D | | 0.631 | 0.751 | 0.63 | 0.771 | 1 | 0.748 |
| D | | 0.754 | 0.832 | 0.774 | 0.879 | 1 | 0.748 |
| D | | 0.403 | 0.616 | 0.48 | 0.666 | 1 | 0.748 |
| N | | -0.506 | 0.219 | -0.416 | 0.245 | 1 | 0.748 |
| N | | -0.081 | 0.382 | -0.072 | 0.366 | 1 | 0.748 |
| D | | 0.593 | 0.727 | 0.638 | 0.777 | 1 | 0.748 |
| D | | 0.468 | 0.652 | 0.519 | 0.693 | 1 | 0.748 |
| D | | 0.802 | 0.862 | 0.8 | 0.898 | 1 | 0.983 |
| D | | 0.631 | 0.752 | 0.648 | 0.784 | 1 | 0.983 |
| D | | 0.806 | 0.865 | 0.773 | 0.879 | 1 | 0.748 |
| . | . | . | . | . | . | . | |
| . | . | . | . | . | . | . | |
| D | | 0.631 | 0.751 | 0.63 | 0.771 | 1 | 0.748 |

| | | | | | | |
|---|---|---|---|---|---|---|
| . | . | . | . | . | . | . |
| D | 0.642 | 0.758 | 0.668 | 0.8 | 1 | 0.983 |
| N | -0.081 | 0.382 | -0.072 | 0.366 | 1 | 0.748 |
| D | 0.142 | 0.484 | 0.249 | 0.526 | 1 | 0.983 |
| N | -0.141 | 0.356 | -0.108 | 0.351 | 1 | 0.748 |
| D | 0.463 | 0.649 | 0.416 | 0.626 | 0.999 | 0.38 |
| D | 1.011 | 0.961 | 0.972 | 0.98 | 1 | 0.748 |
| D | 0.448 | 0.641 | 0.401 | 0.616 | 1 | 0.748 |
| D | 0.835 | 0.882 | 0.805 | 0.901 | 1 | 0.748 |
| . | 1.091 | 0.98 | 0.938 | 0.97 | 1 | 0.748 |
| N | 1.106 | 0.982 | 0.97 | 0.979 | 0.998 | 0.367 |
| D | 0.614 | 0.74 | 0.578 | 0.734 | 1 | 0.748 |
| D | 0.073 | 0.452 | 0.24 | 0.521 | 1 | 0.748 |
| . | . | . | . | . | . | . |
| D | 0.023 | 0.429 | 0.135 | 0.464 | 1 | 0.518 |
| N | 0.255 | 0.539 | 0.192 | 0.494 | 0.14 | 0.172 |
| D | 0.674 | 0.779 | 0.715 | 0.835 | 1 | 0.748 |
| . | . | . | . | . | . | . |
| D | 0.159 | 0.492 | 0.313 | 0.563 | 1 | 0.748 |
| N | -0.076 | 0.385 | -0.03 | 0.384 | 1 | 0.748 |
| N | 0.717 | 0.808 | 0.672 | 0.803 | 1 | 0.748 |
| D | 1.011 | 0.961 | 0.972 | 0.98 | 1 | 0.748 |
| . | . | . | . | . | . | . |
| D | 0.484 | 0.662 | 0.57 | 0.728 | 1 | 0.748 |
| D | 0.476 | 0.657 | 0.419 | 0.627 | 1 | 0.748 |
| D | -0.088 | 0.379 | -0.004 | 0.395 | 1 | 0.983 |
| N | 0.055 | 0.444 | 0.147 | 0.47 | 1 | 0.748 |
| N | 0.244 | 0.533 | 0.223 | 0.511 | 1 | 0.446 |
| D | 0.516 | 0.68 | 0.461 | 0.654 | 1 | 0.748 |
| D | 0.902 | 0.918 | 0.904 | 0.958 | 1 | 0.748 |
| D | 0.78 | 0.848 | 0.743 | 0.857 | 1 | 0.748 |
| D | 1.011 | 0.961 | 0.972 | 0.98 | 1 | 0.748 |
| . | . | . | . | . | . | . |
| D | 0.837 | 0.884 | 0.826 | 0.916 | 1 | 0.748 |
| D | 0.316 | 0.57 | 0.416 | 0.626 | 0.998 | 0.362 |
| D | 0.758 | 0.834 | 0.744 | 0.857 | 1 | 0.748 |
| D | 0.761 | 0.836 | 0.756 | 0.866 | 1 | 0.748 |
| D | 0.844 | 0.888 | 0.855 | 0.933 | 1 | 0.748 |
| . | . | . | . | . | . | . |
| D | 0.78 | 0.848 | 0.743 | 0.857 | 1 | 0.748 |
| D | 0.835 | 0.882 | 0.769 | 0.876 | 1 | 0.748 |
| N | 0.546 | 0.698 | 0.56 | 0.721 | 1 | 0.748 |
| D | 0.724 | 0.812 | 0.73 | 0.847 | 1 | 0.748 |

| | | | | | | |
|---|---|---|---|---|---|---|
| N | 0.212 | 0.518 | 0.186 | 0.491 | 1 | 0.748 |
| D | 0.752 | 0.831 | 0.755 | 0.866 | 1 | 0.748 |
| . | . | . | . | . | . | . |
| . | . | . | . | . | . | . |
| N | 0.055 | 0.444 | 0.147 | 0.47 | 1 | 0.748 |
| D | 0.75 | 0.829 | 0.769 | 0.876 | 1 | 0.748 |
| D | 0.764 | 0.838 | 0.759 | 0.868 | 1 | 0.748 |
| N | 0.717 | 0.808 | 0.672 | 0.803 | 1 | 0.748 |
| D | 0.73 | 0.816 | 0.796 | 0.895 | 1 | 0.983 |
| . | . | . | . | . | . | . |
| D | 0.562 | 0.708 | 0.541 | 0.708 | 1 | 0.748 |
| . | 0.195 | 0.51 | 0.33 | 0.573 | 1 | 0.748 |
| D | 0.45 | 0.642 | 0.243 | 0.522 | 1 | 0.748 |
| D | 0.45 | 0.642 | 0.243 | 0.522 | 1 | 0.748 |
| . | . | . | . | . | . | . |
| . | . | . | . | . | . | . |
| D | 0.852 | 0.892 | 0.81 | 0.905 | 1 | 0.748 |
| D | 0.265 | 0.544 | 0.408 | 0.621 | 1 | 0.748 |
| D | 0.883 | 0.909 | 0.883 | 0.948 | 1 | 0.983 |
| . | . | . | . | . | . | . |
| D | 0.484 | 0.662 | 0.57 | 0.728 | 1 | 0.748 |
| D | 0.692 | 0.791 | 0.712 | 0.833 | 1 | 0.748 |
| D | 0.965 | 0.946 | 0.936 | 0.97 | 1 | 0.748 |
| D | 0.426 | 0.629 | 0.396 | 0.613 | 1 | 0.748 |
| D | 0.224 | 0.524 | 0.387 | 0.608 | 1 | 0.748 |
| D | 0.72 | 0.81 | 0.764 | 0.872 | 1 | 0.748 |
| . | . | . | . | . | . | . |
| D | 0.562 | 0.708 | 0.541 | 0.708 | 1 | 0.748 |
| . | 0.206 | 0.515 | 0.298 | 0.554 | 1 | 0.748 |
| D | 0.642 | 0.758 | 0.668 | 0.8 | 1 | 0.983 |
| D | 0.78 | 0.848 | 0.743 | 0.857 | 1 | 0.748 |
| . | 0.195 | 0.51 | 0.33 | 0.573 | 1 | 0.748 |
| D | 0.78 | 0.848 | 0.743 | 0.857 | 1 | 0.748 |
| N | -0.081 | 0.382 | -0.072 | 0.366 | 1 | 0.748 |
| D | 0.569 | 0.712 | 0.542 | 0.709 | 1 | 0.748 |
| D | -0.095 | 0.376 | 0.088 | 0.439 | 0.44 | 0.204 |
| D | 0.441 | 0.637 | 0.451 | 0.648 | 1 | 0.748 |
| D | 0.965 | 0.946 | 0.936 | 0.97 | 1 | 0.748 |
| D | 0.689 | 0.789 | 0.722 | 0.84 | 1 | 0.748 |
| D | 0.549 | 0.7 | 0.469 | 0.659 | 1 | 0.983 |
| . | . | . | . | . | . | . |
| D | 0.401 | 0.615 | 0.316 | 0.565 | 0.996 | 0.342 |
| D | 0.316 | 0.57 | 0.416 | 0.626 | 0.998 | 0.362 |

| | | | | | | |
|---|---|---|---|---|---|---|
| . | . | . | . | . | . | . |
| D | 0.78 | 0.848 | 0.743 | 0.857 | 1 | 0.748 |
| D | 0.965 | 0.946 | 0.936 | 0.97 | 1 | 0.748 |
| D | 0.316 | 0.57 | 0.416 | 0.626 | 0.998 | 0.362 |
| N | 0.055 | 0.444 | 0.147 | 0.47 | 1 | 0.748 |
| D | 0.316 | 0.57 | 0.416 | 0.626 | 0.998 | 0.362 |
| N | 0.012 | 0.424 | 0.157 | 0.475 | 0.614 | 0.218 |
| D | 0.715 | 0.806 | 0.734 | 0.85 | 1 | 0.748 |
| . | . | . | . | . | . | . |
| D | 0.761 | 0.836 | 0.756 | 0.866 | 1 | 0.748 |
| D | 0.441 | 0.637 | 0.451 | 0.648 | 1 | 0.748 |
| N | 0.145 | 0.486 | 0.071 | 0.431 | 1 | 0.748 |
| . | . | . | . | . | . | . |
| N | 0.326 | 0.575 | 0.216 | 0.507 | 1 | 0.748 |
| D | 0.837 | 0.884 | 0.826 | 0.916 | 1 | 0.748 |
| D | 0.482 | 0.66 | 0.552 | 0.715 | 0.901 | 0.261 |
| D | 0.629 | 0.75 | 0.551 | 0.715 | 1 | 0.748 |
| . | . | . | . | . | . | . |
| D | 0.693 | 0.791 | 0.72 | 0.839 | 1 | 0.748 |
| D | 0.683 | 0.785 | 0.653 | 0.789 | 1 | 0.748 |
| D | 0.629 | 0.75 | 0.551 | 0.715 | 1 | 0.748 |
| D | 0.484 | 0.662 | 0.499 | 0.68 | 1 | 0.983 |
| . | . | . | . | . | . | . |
| D | 0.413 | 0.621 | 0.373 | 0.599 | 0.997 | 0.35 |
| . | . | . | . | . | . | . |
| D | 0.725 | 0.812 | 0.66 | 0.793 | 1 | 0.748 |
| D | 0.498 | 0.67 | 0.556 | 0.718 | 1 | 0.748 |
| N | 1.219 | 0.997 | 1.091 | 0.997 | 1 | 0.748 |
| . | . | . | . | . | . | . |
| D | -0.026 | 0.407 | 0.145 | 0.469 | 1 | 0.748 |
| D | 0.731 | 0.816 | 0.653 | 0.788 | 1 | 0.748 |
| D | 0.623 | 0.746 | 0.651 | 0.787 | 1 | 0.748 |
| . | 0.754 | 0.832 | 0.71 | 0.831 | 1 | 0.983 |
| D | 0.732 | 0.817 | 0.773 | 0.879 | 1 | 0.983 |
| . | . | . | . | . | . | . |
| N | 0.127 | 0.477 | 0.005 | 0.399 | 0.999 | 0.388 |
| D | 0.623 | 0.746 | 0.651 | 0.787 | 1 | 0.748 |
| D | 1.059 | 0.973 | 0.931 | 0.968 | 1 | 0.748 |
| D | 0.676 | 0.781 | 0.659 | 0.793 | 1 | 0.748 |
| . | -0.399 | 0.255 | -0.346 | 0.266 | 1 | 0.449 |
| D | 0.763 | 0.837 | 0.733 | 0.849 | 1 | 0.748 |
| D | 0.804 | 0.863 | 0.812 | 0.906 | 1 | 0.748 |
| D | 0.635 | 0.754 | 0.616 | 0.761 | 1 | 0.748 |

| | | | | | | |
|---|---|---|---|---|---|---|
| . | . | . | . | . | . | . |
| D | 0.725 | 0.812 | 0.66 | 0.793 | 1 | 0.748 |
| D | 0.321 | 0.572 | 0.373 | 0.599 | 1 | 0.748 |
| D | 0.734 | 0.819 | 0.791 | 0.892 | 1 | 0.748 |
| D | 0.544 | 0.697 | 0.577 | 0.733 | 1 | 0.748 |
| D | 0.514 | 0.679 | 0.572 | 0.73 | 1 | 0.748 |
| D | 0.527 | 0.687 | 0.375 | 0.6 | 1 | 0.748 |
| . | . | . | . | . | . | . |
| N | -0.428 | 0.245 | -0.35 | 0.265 | 0.999 | 0.378 |
| D | 0.7 | 0.796 | 0.723 | 0.842 | 1 | 0.748 |
| D | 0.073 | 0.452 | 0.24 | 0.521 | 1 | 0.748 |
| D | 0.464 | 0.65 | 0.526 | 0.698 | 1 | 0.748 |
| D | 0.888 | 0.912 | 0.86 | 0.936 | 1 | 0.748 |
| D | 0.204 | 0.514 | 0.354 | 0.588 | 1 | 0.748 |
| D | 0.564 | 0.709 | 0.588 | 0.741 | 1 | 0.983 |
| D | 0.086 | 0.458 | 0.266 | 0.536 | 1 | 0.748 |
| D | 0.574 | 0.715 | 0.59 | 0.742 | 1 | 0.748 |
| N | 0.482 | 0.66 | 0.412 | 0.623 | 0.933 | 0.271 |
| D | 0.128 | 0.478 | 0.312 | 0.562 | 1 | 0.748 |
| D | 0.574 | 0.715 | 0.59 | 0.742 | 1 | 0.748 |
| N | -0.176 | 0.341 | 0.022 | 0.407 | 1 | 0.748 |
| D | 0.486 | 0.663 | 0.495 | 0.677 | 1 | 0.748 |
| D | 0.484 | 0.662 | 0.499 | 0.68 | 1 | 0.983 |
| D | 0.574 | 0.715 | 0.59 | 0.742 | 1 | 0.748 |
| N | -0.011 | 0.413 | -0.079 | 0.362 | 1 | 0.518 |
| . | . | . | . | . | . | . |
| N | 0.482 | 0.66 | 0.412 | 0.623 | 0.933 | 0.271 |
| D | 0.786 | 0.853 | 0.781 | 0.884 | 1 | 0.748 |
| D | 0.629 | 0.75 | 0.551 | 0.715 | 1 | 0.748 |
| D | 0.439 | 0.636 | 0.477 | 0.665 | 1 | 0.748 |
| D | 0.218 | 0.521 | 0.133 | 0.463 | 1 | 0.429 |
| D | 0.845 | 0.888 | 0.833 | 0.92 | 1 | 0.748 |
| D | 0.527 | 0.687 | 0.375 | 0.6 | 1 | 0.748 |
| D | 0.574 | 0.715 | 0.59 | 0.742 | 1 | 0.748 |
| D | 0.786 | 0.853 | 0.781 | 0.884 | 1 | 0.748 |
| D | 0.676 | 0.781 | 0.659 | 0.793 | 1 | 0.748 |
| D | 0.003 | 0.42 | 0.119 | 0.455 | 1 | 0.748 |
| . | . | . | . | . | . | . |
| D | -0.038 | 0.401 | 0.155 | 0.474 | 1 | 0.748 |
| D | 1.049 | 0.971 | 0.982 | 0.982 | 1 | 0.748 |
| D | 0.63 | 0.751 | 0.615 | 0.76 | 1 | 0.748 |
| D | 0.734 | 0.819 | 0.791 | 0.892 | 1 | 0.748 |
| D | 0.574 | 0.715 | 0.59 | 0.742 | 1 | 0.748 |

| | | | | | | |
|---|---|---|---|---|---|---|
| D | 0.541 | 0.695 | 0.648 | 0.784 | 1 | 0.748 |
| D | 0.786 | 0.853 | 0.781 | 0.884 | 1 | 0.748 |
| D | 0.657 | 0.768 | 0.63 | 0.771 | 1 | 0.748 |

| integrated_fi | integrated_fi | integrated_cc | LINSIGHT | LINSIGHT_rar | GERP.._NR | GERP.._RS |
|---|---|---|---|---|---|---|
| . | . | . | . | . | . | . |
| . | . | . | . | . | . | . |
| 0.549 | 0.229 | 0 | . | . | 5.84 | 5.84 |
| 0.632 | 0.41 | 0 | . | . | 6.11 | 6.11 |
| 0.707 | 0.731 | 0 | . | . | 4.66 | 4.66 |
| 0.707 | 0.731 | 0 | . | . | 5.61 | 3.72 |
| 0.707 | 0.731 | 0 | . | . | 5.12 | 3 |
| 0.732 | 0.924 | 0 | . | . | 5.74 | 4.86 |
| 0.707 | 0.731 | 0 | . | . | 5.6 | 5.6 |
| 0.672 | 0.526 | 0 | . | . | 4.99 | 4.99 |
| 0.722 | 0.854 | 0 | . | . | 5.71 | 5.71 |
| 0.549 | 0.229 | 0 | . | . | 5.84 | 5.84 |
| 0.707 | 0.731 | 0 | . | . | 5.42 | 4.29 |
| 0.707 | 0.731 | 0 | . | . | 5.46 | 5.46 |
| . | . | . | . | . | . | . |
| 0.652 | 0.481 | 0 | . | . | 4.39 | 4.39 |
| 0.707 | 0.731 | 0 | . | . | 4.41 | 3.33 |
| 0.675 | 0.551 | 0 | . | . | 5.12 | 5.12 |
| 0.722 | 0.854 | 0 | . | . | 5.71 | 5.71 |
| 0.696 | 0.57 | 0 | . | . | 5.11 | 4.19 |
| 0.615 | 0.376 | 0 | . | . | 5.73 | 5.73 |
| . | . | . | . | . | . | . |
| 0.581 | 0.331 | 0 | . | . | 5.46 | 4.51 |
| 0.707 | 0.731 | 0 | . | . | 5.87 | 5.87 |
| 0.549 | 0.229 | 0 | . | . | 5.74 | 5.74 |
| 0.719 | 0.831 | 0 | . | . | 5.02 | 5.02 |
| 0.652 | 0.481 | 0 | . | . | 4.64 | 4.64 |
| 0.628 | 0.405 | 0 | . | . | 3.91 | 1.7 |
| 0.765 | 0.991 | 0 | . | . | 4.44 | 4.44 |
| 0.707 | 0.731 | 0 | . | . | 5.48 | 5.48 |
| 0.722 | 0.854 | 0 | . | . | 5.71 | 5.71 |
| 0.623 | 0.398 | 0 | . | . | 4.85 | 1.96 |
| 0.563 | 0.315 | 0 | . | . | 5.08 | 5.08 |
| 0.672 | 0.526 | 0 | . | . | 4.99 | 4.99 |
| 0.737 | 0.974 | 0 | . | . | 5.87 | 4.99 |
| 0.706 | 0.612 | 0 | . | . | 5.87 | 5.87 |
| 0.707 | 0.731 | 0 | . | . | 5.7 | 4.79 |
| 0.437 | 0.071 | 0 | . | . | 5.74 | 5.74 |
| 0.732 | 0.924 | 0 | . | . | 5.27 | 3.36 |
| 0.628 | 0.405 | 0 | . | . | 5.72 | 5.72 |
| 0.707 | 0.731 | 0 | . | . | 5.9 | 4.97 |
| 0.731 | 0.879 | 0 | . | . | 5.96 | 5.96 |

| | | | | | | |
|---|---|---|---|---|---|---|
| 0.549 | 0.229 | 0 | . | . | 5.84 | 5.84 |
| 0.672 | 0.526 | 0 | . | . | 5.4 | 5.4 |
| 0.707 | 0.731 | 0 | . | . | 5.72 | 5.72 |
| 0.707 | 0.731 | 0 | . | . | 5.36 | 5.36 |
| 0.615 | 0.376 | 0 | . | . | 5.73 | 5.73 |
| 0.581 | 0.331 | 0 | . | . | 5.46 | 5.46 |
| . | . | . | . | . | . | . |
| 0.732 | 0.924 | 0 | . | . | 5.38 | 3.47 |
| 0.685 | 0.555 | 0 | . | . | 5.07 | 5.07 |
| 0.672 | 0.526 | 0 | . | . | 5.96 | 5.96 |
| 0.722 | 0.854 | 0 | . | . | 5.64 | 4.76 |
| 0.722 | 0.854 | 0 | . | . | 5.4 | 5.4 |
| 0.672 | 0.526 | 0 | . | . | 5.26 | 2.82 |
| . | . | . | . | . | . | . |
| 0.554 | 0.252 | 0 | . | . | 5.73 | 4.86 |
| 0.653 | 0.485 | 0 | . | . | 5.79 | 5.79 |
| 0.707 | 0.731 | 0 | . | . | 5.26 | 5.26 |
| 0.706 | 0.612 | 0 | . | . | 5.45 | 4.56 |
| 0.635 | 0.418 | 0 | . | . | 6.04 | 4.82 |
| 0.707 | 0.731 | 0 | . | . | 5.98 | 5.09 |
| 0.732 | 0.924 | 0 | . | . | 5.05 | 5.05 |
| . | . | . | . | . | . | . |
| 0.549 | 0.229 | 0 | . | . | 5.84 | 5.84 |
| 0.707 | 0.731 | 0 | . | . | 4.6 | 4.6 |
| 0.707 | 0.731 | 0 | . | . | 5.09 | 5.09 |
| 0.722 | 0.854 | 0 | . | . | 5.71 | 5.71 |
| 0.707 | 0.731 | 0 | . | . | 5.73 | 5.73 |
| 0.722 | 0.854 | 0 | . | . | 5.94 | 4.91 |
| 0.732 | 0.924 | 0 | . | . | 5.38 | 5.38 |
| . | . | . | . | . | . | . |
| 0.707 | 0.731 | 0 | . | . | 6.03 | 4.09 |
| 0.722 | 0.854 | 0 | . | . | 4.38 | 3.49 |
| 0.707 | 0.731 | 0 | . | . | 5.9 | 4.73 |
| 0.707 | 0.731 | 0 | . | . | 5.6 | 5.6 |
| 0.549 | 0.229 | 0 | . | . | 5.84 | 5.84 |
| 0.549 | 0.229 | 0 | . | . | 5.84 | 5.84 |
| 0.707 | 0.731 | 0 | . | . | 5.64 | 5.64 |
| 0.707 | 0.731 | 0 | . | . | 5.51 | 5.51 |
| 0.707 | 0.731 | 0 | . | . | 5.79 | 3.98 |
| 0.706 | 0.612 | 0 | . | . | 5.9 | 5.9 |
| 0.707 | 0.731 | 0 | . | . | 5.18 | 5.18 |
| 0.652 | 0.481 | 0 | . | . | 4.39 | 4.39 |
| 0.628 | 0.405 | 0 | . | . | 4.57 | 3.68 |

| | | | | | | | |
|---|---|---|---|---|---|---|---|
| | 0.719 | 0.831 | 0 . | . | | 5.94 | 5.94 |
| . | . | . | . | . | . | . | |
| | 0.598 | 0.345 | 0 . | . | | 5.27 | 2.88 |
| | 0.61 | 0.356 | 0 . | . | | 5.2 | 5.2 |
| | 0.685 | 0.555 | 0 . | . | | 5.07 | 5.07 |
| | 0.615 | 0.376 | 0 . | . | | 5.08 | 5.08 |
| | 0.707 | 0.731 | 0 . | . | | 5.61 | 5.61 |
| | 0.549 | 0.229 | 0 . | . | | 5.84 | 5.84 |
| | 0.672 | 0.526 | 0 . | . | | 5.55 | 5.55 |
| | 0.726 | 0.872 | 0 . | . | | 4.26 | 4.26 |
| . | . | . | . | . | . | . | |
| | 0.707 | 0.731 | 0 . | . | | 4.66 | 4.66 |
| | 0.677 | 0.553 | 0 . | . | | 4.94 | 4.94 |
| . | . | . | . | . | . | . | |
| . | . | . | . | . | . | . | |
| | 0.707 | 0.731 | 0 . | . | | 5.6 | 5.6 |
| . | . | . | . | . | . | . | |
| | 0.707 | 0.731 | 0 . | . | | 5.59 | 5.59 |
| | 0.732 | 0.924 | 0 . | . | | 5.29 | 5.29 |
| . | . | . | . | . | . | . | |
| | 0.707 | 0.731 | 0 . | . | | 5.89 | 5.89 |
| | 0.706 | 0.612 | 0 . | . | | 5.08 | 5.08 |
| . | . | . | . | . | . | . | |
| | 0.549 | 0.229 | 0 . | . | | 5.84 | 5.84 |
| | 0.722 | 0.854 | 0 . | . | | 5.71 | 5.71 |
| | 0.615 | 0.376 | 0 . | . | | 5.84 | 5.84 |
| | 0.672 | 0.526 | 0 . | . | | 4.99 | 4.99 |
| . | . | . | . | . | . | . | |
| | 0.658 | 0.49 | 0 . | . | | 4.05 | 3.18 |
| | 0.732 | 0.924 | 0 . | . | | 5.19 | 4.31 |
| | 0.646 | 0.454 | 0 . | . | | 5.52 | 5.52 |
| | 0.707 | 0.731 | 0 . | . | | 6.06 | 6.06 |
| | 0.563 | 0.315 | 0 . | . | | 4.94 | 4.94 |
| | 0.722 | 0.854 | 0 . | . | | 5.57 | 2.67 |
| | 0.707 | 0.731 | 0 . | . | | 5.98 | 5.09 |
| | 0.732 | 0.924 | 0 . | . | | 5.82 | 5.82 |
| | 0.745 | 0.986 | 0 . | . | | 5.4 | 5.4 |
| | 0.707 | 0.731 | 0 . | . | | 5.41 | 5.41 |
| | 0.707 | 0.731 | 0 . | . | | 5.37 | 5.37 |
| | 0.677 | 0.553 | 0 . | . | | 5.31 | 5.31 |
| . | . | . | . | . | . | . | |
| . | . | . | . | . | . | . | |
| | 0.646 | 0.454 | 0 . | . | | 5.52 | 5.52 |

| | | | | | | | |
|---|---|---|---|---|---|---|---|
| . | . | . | . | . | . | . | |
| | 0.549 | 0.229 | 0 . | . | | 5.84 | 5.84 |
| | 0.707 | 0.731 | 0 . | . | | 5.98 | 5.09 |
| | 0.722 | 0.854 | 0 . | . | | 5.39 | 5.39 |
| | 0.615 | 0.376 | 0 . | . | | 5.76 | 4.89 |
| | 0.732 | 0.924 | 0 . | . | | 5.05 | 2.9 |
| | 0.632 | 0.41 | 0 . | . | | 6.08 | 6.08 |
| | 0.628 | 0.405 | 0 . | . | | 4.17 | 4.17 |
| | 0.563 | 0.315 | 0 . | . | | 5.22 | 5.22 |
| | 0.156 | 0.033 | 0 | 0.977 | 0.808 | 5.43 | 5.43 |
| | 0.732 | 0.924 | 0 . | . | | 5.85 | 5.85 |
| | 0.707 | 0.731 | 0 . | . | | 5.91 | 4.97 |
| | 0.651 | 0.469 | 0 . | . | | 5.51 | 5.51 |
| . | . | . | . | . | . | . | |
| | 0.73 | 0.876 | 0 . | . | | 5.93 | 5.93 |
| | 0.706 | 0.612 | 0 . | . | | 5.18 | 2.23 |
| | 0.707 | 0.731 | 0 . | . | | 5.83 | 5.83 |
| . | . | . | . | . | . | . | |
| | 0.672 | 0.526 | 0 . | . | | 4.99 | 4.99 |
| | 0.635 | 0.418 | 0 . | . | | 5.93 | 5.06 |
| | 0.732 | 0.924 | 0 . | . | | 4.94 | 4.94 |
| | 0.632 | 0.41 | 0 . | . | | 6.08 | 6.08 |
| . | . | . | . | . | . | . | |
| | 0.563 | 0.315 | 0 . | . | | 5.99 | 5.99 |
| | 0.628 | 0.405 | 0 . | . | | 5.47 | 3.38 |
| | 0.025 | 0.001 | 3 . | . | | 5.23 | 4.36 |
| | 0.706 | 0.612 | 0 . | . | | 5.87 | 3.99 |
| | 0.672 | 0.526 | 0 . | . | | 5.87 | 4.69 |
| | 0.672 | 0.526 | 0 . | . | | 6.08 | 6.08 |
| | 0.563 | 0.315 | 0 . | . | | 5.77 | 5.77 |
| | 0.732 | 0.924 | 0 . | . | | 5.38 | 5.38 |
| | 0.632 | 0.41 | 0 . | . | | 6.08 | 6.08 |
| . | . | . | . | . | . | . | |
| | 0.672 | 0.526 | 0 . | . | | 5.44 | 5.44 |
| | 0.732 | 0.924 | 0 . | . | | 5.1 | 5.1 |
| | 0.707 | 0.731 | 0 . | . | | 5.32 | 5.32 |
| | 0.563 | 0.315 | 0 . | . | | 5.52 | 5.52 |
| | 0.707 | 0.731 | 0 . | . | | 5.96 | 5.96 |
| . | . | . | . | . | . | . | |
| | 0.732 | 0.924 | 0 . | . | | 5.38 | 5.38 |
| | 0.731 | 0.879 | 0 . | . | | 5.15 | 5.15 |
| | 0.706 | 0.612 | 0 . | . | | 5.74 | 5.74 |
| | 0.672 | 0.526 | 0 . | . | | 5.74 | 5.74 |

| | | | | | | |
|---|---|---|---|---|---|---|
| 0.615 | 0.376 | 0 | . | . | 5.76 | 5.76 |
| 0.732 | 0.924 | 0 | . | . | 5.49 | 5.49 |
| . | . | . | . | . | . | . |
| . | . | . | . | . | . | . |
| 0.706 | 0.612 | 0 | . | . | 5.87 | 3.99 |
| 0.516 | 0.209 | 0 | . | . | 5.83 | 5.83 |
| 0.651 | 0.469 | 0 | . | . | 5.4 | 5.4 |
| 0.732 | 0.924 | 0 | . | . | 4.94 | 4.94 |
| 0.707 | 0.731 | 0 | . | . | 5.99 | 5.99 |
| . | . | . | . | . | . | . |
| 0.442 | 0.08 | 0 | . | . | 5.38 | 4.48 |
| 0.677 | 0.553 | 0 | . | . | 5.55 | 5.55 |
| 0.628 | 0.405 | 0 | . | . | 4.57 | 0.795 |
| 0.628 | 0.405 | 0 | . | . | 4.57 | 0.795 |
| . | . | . | . | . | . | . |
| . | . | . | . | . | . | . |
| 0.732 | 0.924 | 0 | . | . | 5.27 | 5.27 |
| 0.707 | 0.731 | 0 | . | . | 5.34 | 5.34 |
| 0.554 | 0.289 | 0 | . | . | 5.75 | 5.75 |
| . | . | . | . | . | . | . |
| 0.563 | 0.315 | 0 | . | . | 5.99 | 5.99 |
| 0.563 | 0.315 | 0 | . | . | 5.78 | 5.78 |
| 0.707 | 0.731 | 0 | . | . | 5.46 | 5.46 |
| 0.517 | 0.214 | 0 | . | . | 3.62 | 3.62 |
| 0.707 | 0.731 | 0 | . | . | 5.64 | 5.64 |
| 0.615 | 0.376 | 0 | . | . | 5.78 | 5.78 |
| . | . | . | . | . | . | . |
| 0.442 | 0.08 | 0 | . | . | 5.38 | 4.48 |
| 0.722 | 0.854 | 0 | . | . | 4.94 | 4.94 |
| 0.549 | 0.229 | 0 | . | . | 5.84 | 5.84 |
| 0.732 | 0.924 | 0 | . | . | 5.38 | 5.38 |
| 0.677 | 0.553 | 0 | . | . | 5.55 | 5.55 |
| 0.732 | 0.924 | 0 | . | . | 5.38 | 5.38 |
| 0.707 | 0.731 | 0 | . | . | 5.98 | 5.09 |
| 0.706 | 0.612 | 0 | . | . | 5.08 | 5.08 |
| 0.707 | 0.731 | 0 | . | . | 4.41 | 4.41 |
| 0.707 | 0.731 | 0 | . | . | 5.9 | 4.97 |
| 0.707 | 0.731 | 0 | . | . | 5.46 | 5.46 |
| 0.707 | 0.731 | 0 | . | . | 5.72 | 5.72 |
| 0.675 | 0.551 | 0 | . | . | 5.45 | 5.45 |
| . | . | . | . | . | . | . |
| 0.707 | 0.731 | 0 | . | . | 5.87 | 1.71 |
| 0.732 | 0.924 | 0 | . | . | 5.1 | 5.1 |

| | | | | | | | |
|---|---|---|---|---|---|---|---|
| . | . | . | . | . | . | . | |
| | 0.732 | 0.924 | 0 | . | . | 5.38 | 5.38 |
| | 0.707 | 0.731 | 0 | . | . | 5.46 | 5.46 |
| | 0.732 | 0.924 | 0 | . | . | 5.1 | 5.1 |
| | 0.706 | 0.612 | 0 | . | . | 5.87 | 3.99 |
| | 0.732 | 0.924 | 0 | . | . | 5.1 | 5.1 |
| | 0.706 | 0.612 | 0 | . | . | 5.08 | 5.08 |
| | 0.707 | 0.731 | 0 | . | . | 5.75 | 5.75 |
| . | . | . | . | . | . | . | |
| | 0.563 | 0.315 | 0 | . | . | 5.52 | 5.52 |
| | 0.707 | 0.731 | 0 | . | . | 5.9 | 4.97 |
| | 0.635 | 0.418 | 0 | . | . | 6.04 | 5.16 |
| . | . | . | . | . | . | . | |
| | 0.672 | 0.526 | 0 | . | . | 5.4 | 5.4 |
| | 0.672 | 0.526 | 0 | . | . | 5.44 | 5.44 |
| | 0.707 | 0.731 | 0 | . | . | 5.52 | 5.52 |
| | 0.706 | 0.612 | 0 | . | . | 5.18 | 5.18 |
| . | . | . | . | . | . | . | |
| | 0.707 | 0.731 | 0 | . | . | 5.86 | 5.86 |
| | 0.563 | 0.315 | 0 | . | . | 4.51 | 4.51 |
| | 0.706 | 0.612 | 0 | . | . | 5.18 | 5.18 |
| | 0.437 | 0.071 | 0 | . | . | 4.39 | 4.39 |
| . | . | . | . | . | . | . | |
| | 0.707 | 0.731 | 0 | . | . | 5.63 | 3.79 |
| . | . | . | . | . | . | . | |
| | 0.677 | 0.553 | 0 | . | . | 5.2 | 4.31 |
| | 0.672 | 0.526 | 0 | . | . | 5.87 | 5.87 |
| | 0.732 | 0.924 | 0 | . | . | 5.95 | 5.95 |
| . | . | . | . | . | . | . | |
| | 0.707 | 0.731 | 0 | . | . | 5.01 | 5.01 |
| | 0.696 | 0.57 | 0 | . | . | 5.56 | 5.56 |
| | 0.745 | 0.986 | 0 | . | . | 5.66 | 5.66 |
| | 0.443 | 0.088 | 1 | . | . | 4.24 | 4.24 |
| | 0.707 | 0.731 | 0 | . | . | 5.41 | 5.41 |
| . | . | . | . | . | . | . | |
| | 0.672 | 0.526 | 0 | . | . | 5.3 | 4.4 |
| | 0.745 | 0.986 | 0 | . | . | 5.66 | 5.66 |
| | 0.543 | 0.224 | 0 | . | . | 4.71 | 4.71 |
| | 0.707 | 0.731 | 0 | . | . | 5.13 | 5.13 |
| | 0.732 | 0.924 | 0 | . | . | 4.75 | 2.94 |
| | 0.563 | 0.315 | 0 | . | . | 4.92 | 4.92 |
| | 0.646 | 0.454 | 0 | . | . | 6.04 | 6.04 |
| | 0.707 | 0.731 | 0 | . | . | 5.2 | 5.2 |

| | | | | | | |
|---|---|---|---|---|---|---|
| . | . | . | . | . | . | . |
| 0.677 | 0.553 | 0 | . | . | 5.2 | 4.31 |
| 0.707 | 0.731 | 0 | . | . | 4.47 | 4.47 |
| 0.707 | 0.731 | 0 | . | . | 5.9 | 5.9 |
| 0.707 | 0.731 | 0 | . | . | 5.38 | 5.38 |
| 0.563 | 0.315 | 0 | . | . | 5.31 | 5.31 |
| 0.628 | 0.405 | 0 | . | . | 5.4 | 0.724 |
| . | . | . | . | . | . | . |
| 0.672 | 0.526 | 0 | . | . | 5.56 | 4.65 |
| 0.487 | 0.14 | 0 | . | . | 5.79 | 5.79 |
| 0.651 | 0.469 | 0 | . | . | 5.51 | 5.51 |
| 0.706 | 0.612 | 0 | . | . | 5.9 | 5.9 |
| 0.706 | 0.612 | 0 | . | . | 5.06 | 5.06 |
| 0.722 | 0.854 | 0 | . | . | 5.35 | 5.35 |
| 0.707 | 0.731 | 0 | . | . | 5.37 | 5.37 |
| 0.563 | 0.315 | 0 | . | . | 5.45 | 5.45 |
| 0.706 | 0.612 | 0 | . | . | 5.46 | 5.46 |
| 0.646 | 0.454 | 0 | . | . | 4.39 | 4.39 |
| 0.722 | 0.854 | 0 | . | . | 5.77 | 4.76 |
| 0.706 | 0.612 | 0 | . | . | 5.46 | 5.46 |
| 0.437 | 0.071 | 0 | . | . | 4.39 | 4.39 |
| 0.745 | 0.986 | 0 | . | . | 5.66 | 5.66 |
| 0.437 | 0.071 | 0 | . | . | 4.39 | 4.39 |
| 0.706 | 0.612 | 0 | . | . | 5.46 | 5.46 |
| 0.672 | 0.526 | 0 | . | . | 4.96 | 4.96 |
| . | . | . | . | . | . | . |
| 0.646 | 0.454 | 0 | . | . | 4.39 | 4.39 |
| 0.707 | 0.731 | 0 | . | . | 5.61 | 5.61 |
| 0.706 | 0.612 | 0 | . | . | 5.18 | 5.18 |
| 0.563 | 0.315 | 0 | . | . | 5.11 | 5.11 |
| 0.707 | 0.731 | 0 | . | . | 5.26 | 4.13 |
| 0.651 | 0.469 | 0 | . | . | 5.52 | 5.52 |
| 0.628 | 0.405 | 0 | . | . | 5.4 | 0.724 |
| 0.706 | 0.612 | 0 | . | . | 5.46 | 5.46 |
| 0.707 | 0.731 | 0 | . | . | 5.61 | 5.61 |
| 0.707 | 0.731 | 0 | . | . | 5.13 | 5.13 |
| 0.447 | 0.091 | 0 | . | . | 4.53 | 3.64 |
| . | . | . | . | . | . | . |
| 0.707 | 0.731 | 0 | . | . | 5.51 | 5.51 |
| 0.615 | 0.376 | 0 | . | . | 5.86 | 5.86 |
| 0.677 | 0.553 | 0 | . | . | 5.3 | 4.41 |
| 0.707 | 0.731 | 0 | . | . | 5.9 | 5.9 |
| 0.706 | 0.612 | 0 | . | . | 5.46 | 5.46 |

| | | | | | |
|---|---|---|---|---|---|
| 0.628 | 0.405 | 0 . | . | 6.03 | 6.03 |
| 0.707 | 0.731 | 0 . | . | 5.61 | 5.61 |
| 0.707 | 0.731 | 0 . | . | 5.7 | 5.7 |

| GERP.._RS_ra | phyloP100wa | phyloP100wa | phyloP30way | phyloP30way | phastCons10( | phastCons10( |
|---|---|---|---|---|---|---|
| . | . | . | . | . | . | . |
| . | . | . | . | . | . | . |
| 0.934 | 5.502 | 0.667 | 1.138 | 0.647 | 1 | 0.716 |
| 0.991 | 7.713 | 0.837 | 1.026 | 0.459 | 1 | 0.716 |
| 0.579 | 2.516 | 0.452 | 1.083 | 0.538 | 0.933 | 0.324 |
| 0.419 | 1.6 | 0.364 | 0.218 | 0.243 | 1 | 0.716 |
| 0.338 | 0.425 | 0.211 | 1.026 | 0.459 | 0.034 | 0.207 |
| 0.626 | 6.727 | 0.746 | 1.176 | 0.789 | 1 | 0.716 |
| 0.85 | 6.036 | 0.706 | 1.026 | 0.459 | 1 | 0.716 |
| 0.659 | 6.477 | 0.735 | 1.067 | 0.529 | 1 | 0.716 |
| 0.89 | 7.547 | 0.811 | 1.312 | 0.947 | 1 | 0.716 |
| 0.934 | 5.502 | 0.667 | 1.138 | 0.647 | 1 | 0.716 |
| 0.504 | 2.781 | 0.474 | 1.138 | 0.647 | 1 | 0.716 |
| 0.8 | 10.003 | 0.997 | 1.155 | 0.708 | 1 | 0.716 |
| . | . | . | . | . | . | . |
| 0.522 | 5.008 | 0.637 | 1.303 | 0.88 | 1 | 0.716 |
| 0.372 | 4.509 | 0.602 | 1.298 | 0.875 | 1 | 0.716 |
| 0.695 | 7.972 | 0.876 | 1.138 | 0.647 | 1 | 0.716 |
| 0.89 | 7.547 | 0.811 | 1.312 | 0.947 | 1 | 0.716 |
| 0.486 | 6.503 | 0.736 | 1.097 | 0.546 | 1 | 0.716 |
| 0.897 | 7.568 | 0.815 | 1.026 | 0.459 | 1 | 0.716 |
| . | . | . | . | . | . | . |
| 0.546 | 3.856 | 0.557 | 1.022 | 0.399 | 0.964 | 0.339 |
| 0.943 | 9.246 | 0.946 | 1.312 | 0.947 | 1 | 0.716 |
| 0.901 | 9.325 | 0.96 | 1.312 | 0.947 | 1 | 0.716 |
| 0.667 | 3.704 | 0.545 | 1.023 | 0.404 | 0.999 | 0.427 |
| 0.574 | 7.807 | 0.846 | 1.138 | 0.647 | 1 | 0.716 |
| 0.234 | 4.639 | 0.611 | 0.932 | 0.336 | 0.77 | 0.294 |
| 0.532 | 4.777 | 0.621 | 1.101 | 0.549 | 1 | 0.716 |
| 0.807 | 7.884 | 0.858 | 1.026 | 0.459 | 1 | 0.716 |
| 0.89 | 7.547 | 0.811 | 1.312 | 0.947 | 1 | 0.716 |
| 0.252 | 0.252 | 0.18 | 0.085 | 0.158 | 0.899 | 0.314 |
| 0.684 | 6.131 | 0.714 | 0.129 | 0.186 | 1 | 0.716 |
| 0.659 | 6.477 | 0.735 | 1.067 | 0.529 | 1 | 0.716 |
| 0.659 | 3.404 | 0.524 | 0.126 | 0.18 | 1 | 0.716 |
| 0.943 | 5.111 | 0.645 | 1.176 | 0.789 | 1 | 0.716 |
| 0.609 | 4.734 | 0.617 | 1.026 | 0.459 | 1 | 0.716 |
| 0.901 | 7.021 | 0.762 | 1.12 | 0.565 | 1 | 0.716 |
| 0.376 | 5.619 | 0.674 | 1.176 | 0.789 | 1 | 0.716 |
| 0.894 | 6.645 | 0.742 | 1.176 | 0.789 | 1 | 0.716 |
| 0.654 | 5.809 | 0.688 | 1.176 | 0.789 | 1 | 0.716 |
| 0.967 | 3.419 | 0.525 | 1.026 | 0.459 | 1 | 0.716 |

| | | | | | | |
|---|---|---|---|---|---|---|
| 0.934 | 5.502 | 0.667 | 1.138 | 0.647 | 1 | 0.716 |
| 0.78 | 4.21 | 0.583 | 1.312 | 0.947 | 0.917 | 0.319 |
| 0.894 | 5.972 | 0.701 | 1.176 | 0.789 | 1 | 0.716 |
| 0.766 | 5.458 | 0.664 | 1.123 | 0.569 | 1 | 0.716 |
| 0.897 | 7.568 | 0.815 | 1.026 | 0.459 | 1 | 0.716 |
| 0.8 | 7.847 | 0.852 | 1.019 | 0.393 | 1 | 0.716 |
| . | . | . | . | . | . | . |
| 0.388 | 3.144 | 0.503 | 1.176 | 0.789 | 1 | 0.716 |
| 0.681 | 7.906 | 0.868 | 1.176 | 0.789 | 1 | 0.716 |
| 0.967 | 6.17 | 0.718 | 1.026 | 0.459 | 1 | 0.716 |
| 0.602 | 2.127 | 0.417 | 0.125 | 0.178 | 1 | 0.716 |
| 0.78 | 7.852 | 0.852 | 1.016 | 0.388 | 1 | 0.716 |
| 0.321 | 2.301 | 0.433 | 1.312 | 0.947 | 0.912 | 0.317 |
| . | . | . | . | . | . | . |
| 0.626 | 0.689 | 0.251 | 1.026 | 0.459 | 0.002 | 0.153 |
| 0.918 | 9.864 | 0.984 | 1.176 | 0.789 | 1 | 0.716 |
| 0.735 | 7.902 | 0.861 | 1.026 | 0.459 | 1 | 0.716 |
| 0.556 | 1.727 | 0.377 | 1.026 | 0.459 | 1 | 0.716 |
| 0.616 | 1.066 | 0.302 | 1.138 | 0.647 | 0.804 | 0.298 |
| 0.686 | 3.213 | 0.509 | 1.176 | 0.789 | 1 | 0.716 |
| 0.676 | 5.545 | 0.669 | 1.026 | 0.459 | 1 | 0.716 |
| . | . | . | . | . | . | . |
| 0.934 | 5.502 | 0.667 | 1.138 | 0.647 | 1 | 0.716 |
| 0.565 | 7.923 | 0.87 | 1.134 | 0.584 | 1 | 0.716 |
| 0.686 | 6.224 | 0.722 | 1.312 | 0.947 | 1 | 0.716 |
| 0.89 | 7.547 | 0.811 | 1.312 | 0.947 | 1 | 0.716 |
| 0.897 | 7.259 | 0.778 | 1.138 | 0.647 | 1 | 0.716 |
| 0.639 | 7.86 | 0.854 | 1.026 | 0.459 | 1 | 0.716 |
| 0.773 | 8.964 | 0.93 | 1.176 | 0.789 | 1 | 0.716 |
| . | . | . | . | . | . | . |
| 0.47 | 2.338 | 0.436 | 1.026 | 0.459 | 1 | 0.716 |
| 0.391 | 2.73 | 0.47 | 1.026 | 0.459 | 0.929 | 0.322 |
| 0.595 | 4.958 | 0.634 | 1.138 | 0.647 | 1 | 0.716 |
| 0.85 | 6.036 | 0.706 | 1.026 | 0.459 | 1 | 0.716 |
| 0.934 | 5.502 | 0.667 | 1.138 | 0.647 | 1 | 0.716 |
| 0.934 | 5.502 | 0.667 | 1.138 | 0.647 | 1 | 0.716 |
| 0.865 | 8.017 | 0.887 | 1.138 | 0.647 | 1 | 0.716 |
| 0.818 | 7.263 | 0.778 | 1.176 | 0.789 | 1 | 0.716 |
| 0.454 | 3.871 | 0.558 | 1.026 | 0.459 | 1 | 0.716 |
| 0.95 | 4.478 | 0.6 | 1.312 | 0.947 | 1 | 0.716 |
| 0.711 | 7.568 | 0.815 | 1.312 | 0.947 | 1 | 0.716 |
| 0.522 | 5.008 | 0.637 | 1.303 | 0.88 | 1 | 0.716 |
| 0.414 | 2.742 | 0.471 | 1.026 | 0.459 | 0.986 | 0.362 |

| | | | | | | |
|---|---|---|---|---|---|---|
| 0.962 | 6.829 | 0.751 | 1.176 | 0.789 | 1 | 0.716 |
| . | . | . | . | . | . | . |
| 0.326 | 3.198 | 0.507 | 0.281 | 0.271 | 1 | 0.716 |
| 0.717 | 6.09 | 0.71 | 1.026 | 0.459 | 1 | 0.716 |
| 0.681 | 7.906 | 0.868 | 1.176 | 0.789 | 1 | 0.716 |
| 0.684 | 9.153 | 0.938 | 1.176 | 0.789 | 1 | 0.716 |
| 0.853 | 2.169 | 0.421 | 1.026 | 0.459 | 1 | 0.716 |
| 0.934 | 5.502 | 0.667 | 1.138 | 0.647 | 1 | 0.716 |
| 0.833 | 8.477 | 0.903 | 1.176 | 0.789 | 1 | 0.716 |
| 0.498 | 7.39 | 0.791 | 0.962 | 0.354 | 1 | 0.716 |
| . | . | . | . | . | . | . |
| 0.579 | 2.516 | 0.452 | 1.083 | 0.538 | 0.933 | 0.324 |
| 0.646 | 9.979 | 0.992 | 1.176 | 0.789 | 1 | 0.716 |
| . | . | . | . | . | . | . |
| . | . | . | . | . | . | . |
| 0.85 | 6.036 | 0.706 | 1.026 | 0.459 | 1 | 0.716 |
| . | . | . | . | . | . | . |
| 0.847 | 4.915 | 0.631 | 1.312 | 0.947 | 1 | 0.716 |
| 0.744 | 2.39 | 0.441 | 1.026 | 0.459 | 0.979 | 0.352 |
| . | . | . | . | . | . | . |
| 0.948 | 7.813 | 0.847 | 1.167 | 0.722 | 1 | 0.716 |
| 0.684 | 5.623 | 0.674 | 1.138 | 0.647 | 1 | 0.716 |
| . | . | . | . | . | . | . |
| 0.934 | 5.502 | 0.667 | 1.138 | 0.647 | 1 | 0.716 |
| 0.89 | 7.547 | 0.811 | 1.312 | 0.947 | 1 | 0.716 |
| 0.934 | 6.634 | 0.741 | 1.176 | 0.789 | 1 | 0.716 |
| 0.659 | 6.477 | 0.735 | 1.067 | 0.529 | 1 | 0.716 |
| . | . | . | . | . | . | . |
| 0.356 | 1.734 | 0.378 | 0.944 | 0.341 | 0.009 | 0.182 |
| 0.507 | 0.525 | 0.227 | 1.026 | 0.459 | 0.992 | 0.376 |
| 0.822 | 9.302 | 0.953 | 1.176 | 0.789 | 1 | 0.716 |
| 0.983 | 7.583 | 0.818 | 1.134 | 0.584 | 1 | 0.716 |
| 0.646 | 5.522 | 0.668 | 1.176 | 0.789 | 1 | 0.716 |
| 0.308 | 2.446 | 0.446 | 1.176 | 0.789 | 0.997 | 0.402 |
| 0.686 | 3.213 | 0.509 | 1.176 | 0.789 | 1 | 0.716 |
| 0.927 | 3.824 | 0.554 | 1.172 | 0.73 | 1 | 0.716 |
| 0.78 | 4.653 | 0.612 | 1.176 | 0.789 | 1 | 0.716 |
| 0.783 | 7.726 | 0.838 | 1.026 | 0.459 | 1 | 0.716 |
| 0.769 | 5.587 | 0.672 | 1.026 | 0.459 | 1 | 0.716 |
| 0.751 | 7.952 | 0.873 | 1.138 | 0.647 | 1 | 0.716 |
| . | . | . | . | . | . | . |
| . | . | . | . | . | . | . |
| 0.822 | 9.302 | 0.953 | 1.176 | 0.789 | 1 | 0.716 |

| | | | | | | |
|---|---|---|---|---|---|---|
| . | . | . | . | . | . | . |
| 0.934 | 5.502 | 0.667 | 1.138 | 0.647 | 1 | 0.716 |
| 0.686 | 3.213 | 0.509 | 1.176 | 0.789 | 1 | 0.716 |
| 0.776 | 7.849 | 0.852 | 1.016 | 0.388 | 1 | 0.716 |
| 0.634 | 3.473 | 0.529 | 1.026 | 0.459 | 0.999 | 0.427 |
| 0.328 | 2.25 | 0.428 | 1.026 | 0.459 | 0.992 | 0.376 |
| 0.99 | 7.905 | 0.865 | 1.026 | 0.459 | 1 | 0.716 |
| 0.483 | 5.161 | 0.648 | 1.155 | 0.708 | 1 | 0.716 |
| 0.723 | 7.854 | 0.853 | 1.026 | 0.459 | 1 | 0.716 |
| 0.79 | 7.206 | 0.774 | 1.138 | 0.647 | 1 | 0.716 |
| 0.937 | 3.408 | 0.524 | 1.026 | 0.459 | 1 | 0.716 |
| 0.654 | 3.71 | 0.546 | 1.026 | 0.459 | 1 | 0.716 |
| 0.818 | 6.252 | 0.724 | 1.138 | 0.647 | 1 | 0.716 |
| . | . | . | . | . | . | . |
| 0.959 | 4.323 | 0.59 | 1.176 | 0.789 | 0.98 | 0.353 |
| 0.272 | 1.438 | 0.346 | -0.242 | 0.084 | 1 | 0.716 |
| 0.931 | 9.22 | 0.943 | 1.312 | 0.947 | 1 | 0.716 |
| . | . | . | . | . | . | . |
| 0.659 | 6.477 | 0.735 | 1.067 | 0.529 | 1 | 0.716 |
| 0.678 | 3.78 | 0.551 | 1.176 | 0.789 | 0.999 | 0.427 |
| 0.646 | 4.534 | 0.603 | 1.176 | 0.789 | 1 | 0.716 |
| 0.99 | 7.905 | 0.865 | 1.026 | 0.459 | 1 | 0.716 |
| . | . | . | . | . | . | . |
| 0.973 | 3.217 | 0.509 | 1.176 | 0.789 | 1 | 0.716 |
| 0.378 | 4.965 | 0.634 | 1.002 | 0.37 | 1 | 0.716 |
| 0.516 | 4.773 | 0.62 | 1.026 | 0.459 | 0.998 | 0.413 |
| 0.455 | 1.75 | 0.38 | 1.176 | 0.789 | 1 | 0.716 |
| 0.585 | 2.007 | 0.405 | 1.312 | 0.947 | 0.922 | 0.32 |
| 0.99 | 4.515 | 0.602 | 1.176 | 0.789 | 1 | 0.716 |
| 0.911 | 10.003 | 0.997 | 1.176 | 0.789 | 1 | 0.716 |
| 0.773 | 8.964 | 0.93 | 1.176 | 0.789 | 1 | 0.716 |
| 0.99 | 7.905 | 0.865 | 1.026 | 0.459 | 1 | 0.716 |
| . | . | . | . | . | . | . |
| 0.793 | 9.996 | 0.993 | 1.176 | 0.789 | 1 | 0.716 |
| 0.689 | 5.885 | 0.695 | 1.176 | 0.789 | 1 | 0.716 |
| 0.754 | 7.899 | 0.86 | 1.026 | 0.459 | 1 | 0.716 |
| 0.822 | 7.969 | 0.876 | 1.134 | 0.584 | 1 | 0.716 |
| 0.967 | 7.634 | 0.824 | 1.176 | 0.789 | 1 | 0.716 |
| . | . | . | . | . | . | . |
| 0.773 | 8.964 | 0.93 | 1.176 | 0.789 | 1 | 0.716 |
| 0.703 | 9.265 | 0.948 | 1.312 | 0.947 | 1 | 0.716 |
| 0.901 | 3.362 | 0.52 | 1.312 | 0.947 | 0.98 | 0.353 |
| 0.901 | 7.235 | 0.776 | 1.138 | 0.647 | 1 | 0.716 |

| | | | | | | |
|---|---|---|---|---|---|---|
| 0.907 | 4.11 | 0.576 | 1.026 | 0.459 | 0.999 | 0.427 |
| 0.81 | 9.325 | 0.96 | 1.312 | 0.947 | 1 | 0.716 |
| . | . | . | . | . | . | . |
| . | . | . | . | . | . | . |
| 0.455 | 1.75 | 0.38 | 1.176 | 0.789 | 1 | 0.716 |
| 0.931 | 7.272 | 0.779 | 1.176 | 0.789 | 1 | 0.716 |
| 0.78 | 9.325 | 0.96 | 1.312 | 0.947 | 1 | 0.716 |
| 0.646 | 4.534 | 0.603 | 1.176 | 0.789 | 1 | 0.716 |
| 0.973 | 9.602 | 0.976 | 1.176 | 0.789 | 1 | 0.716 |
| . | . | . | . | . | . | . |
| 0.54 | 3.972 | 0.565 | 1.176 | 0.789 | 1 | 0.716 |
| 0.833 | 4.205 | 0.583 | 1.176 | 0.789 | 1 | 0.716 |
| 0.178 | 0.951 | 0.287 | -0.138 | 0.114 | 1 | 0.716 |
| 0.178 | 0.951 | 0.287 | -0.138 | 0.114 | 1 | 0.716 |
| . | . | . | . | . | . | . |
| . | . | . | . | . | . | . |
| 0.738 | 7.987 | 0.878 | 1.176 | 0.789 | 1 | 0.716 |
| 0.76 | 7.229 | 0.776 | 1.312 | 0.947 | 1 | 0.716 |
| 0.904 | 9.602 | 0.976 | 1.176 | 0.789 | 1 | 0.716 |
| . | . | . | . | . | . | . |
| 0.973 | 3.217 | 0.509 | 1.176 | 0.789 | 1 | 0.716 |
| 0.914 | 6.137 | 0.715 | 1.026 | 0.459 | 1 | 0.716 |
| 0.8 | 10.003 | 0.997 | 1.155 | 0.708 | 1 | 0.716 |
| 0.406 | 7.23 | 0.776 | 0.941 | 0.339 | 1 | 0.716 |
| 0.865 | 9.325 | 0.96 | 1.312 | 0.947 | 1 | 0.716 |
| 0.914 | 7.905 | 0.865 | 1.026 | 0.459 | 1 | 0.716 |
| . | . | . | . | . | . | . |
| 0.54 | 3.972 | 0.565 | 1.176 | 0.789 | 1 | 0.716 |
| 0.646 | 5.066 | 0.642 | 1.312 | 0.947 | 1 | 0.716 |
| 0.934 | 5.502 | 0.667 | 1.138 | 0.647 | 1 | 0.716 |
| 0.773 | 8.964 | 0.93 | 1.176 | 0.789 | 1 | 0.716 |
| 0.833 | 4.205 | 0.583 | 1.176 | 0.789 | 1 | 0.716 |
| 0.773 | 8.964 | 0.93 | 1.176 | 0.789 | 1 | 0.716 |
| 0.686 | 3.213 | 0.509 | 1.176 | 0.789 | 1 | 0.716 |
| 0.684 | 7.325 | 0.784 | 1.138 | 0.647 | 1 | 0.716 |
| 0.526 | 6.21 | 0.721 | 1.298 | 0.875 | 1 | 0.716 |
| 0.654 | 5.809 | 0.688 | 1.176 | 0.789 | 1 | 0.716 |
| 0.8 | 10.003 | 0.997 | 1.155 | 0.708 | 1 | 0.716 |
| 0.894 | 5.972 | 0.701 | 1.176 | 0.789 | 1 | 0.716 |
| 0.797 | 9.942 | 0.989 | 1.176 | 0.789 | 1 | 0.716 |
| . | . | . | . | . | . | . |
| 0.234 | 0.779 | 0.264 | 1.026 | 0.459 | 0.999 | 0.427 |
| 0.689 | 5.885 | 0.695 | 1.176 | 0.789 | 1 | 0.716 |

| | | | | | | |
|---|---|---|---|---|---|---|
| . | . | . | . | . | . | . |
| 0.773 | 8.964 | 0.93 | 1.176 | 0.789 | 1 | 0.716 |
| 0.8 | 10.003 | 0.997 | 1.155 | 0.708 | 1 | 0.716 |
| 0.689 | 5.885 | 0.695 | 1.176 | 0.789 | 1 | 0.716 |
| 0.455 | 1.75 | 0.38 | 1.176 | 0.789 | 1 | 0.716 |
| 0.689 | 5.885 | 0.695 | 1.176 | 0.789 | 1 | 0.716 |
| 0.684 | 0.528 | 0.227 | 1.026 | 0.459 | 0.999 | 0.427 |
| 0.904 | 5.871 | 0.694 | 1.164 | 0.718 | 1 | 0.716 |
| . | . | . | . | . | . | . |
| 0.822 | 7.969 | 0.876 | 1.134 | 0.584 | 1 | 0.716 |
| 0.654 | 5.809 | 0.688 | 1.176 | 0.789 | 1 | 0.716 |
| 0.706 | 1.727 | 0.377 | 1.026 | 0.459 | 0.327 | 0.256 |
| . | . | . | . | . | . | . |
| 0.78 | 2.705 | 0.468 | 1.312 | 0.947 | 0.982 | 0.355 |
| 0.793 | 9.996 | 0.993 | 1.176 | 0.789 | 1 | 0.716 |
| 0.822 | 4.093 | 0.574 | 1.01 | 0.38 | 0.996 | 0.394 |
| 0.711 | 7.224 | 0.775 | 1.026 | 0.459 | 1 | 0.716 |
| . | . | . | . | . | . | . |
| 0.939 | 6.787 | 0.749 | 1.312 | 0.947 | 1 | 0.716 |
| 0.546 | 7.858 | 0.853 | 1.026 | 0.459 | 1 | 0.716 |
| 0.711 | 7.224 | 0.775 | 1.026 | 0.459 | 1 | 0.716 |
| 0.522 | 6.706 | 0.745 | 1.172 | 0.73 | 1 | 0.716 |
| . | . | . | . | . | . | . |
| 0.428 | 2.129 | 0.417 | 1.026 | 0.459 | 1 | 0.716 |
| . | . | . | . | . | . | . |
| 0.507 | 6.164 | 0.717 | 1.026 | 0.459 | 1 | 0.716 |
| 0.943 | 8.069 | 0.893 | 1.312 | 0.947 | 1 | 0.716 |
| 0.964 | 7.063 | 0.764 | 1.026 | 0.459 | 1 | 0.716 |
| . | . | . | . | . | . | . |
| 0.665 | 4.734 | 0.617 | 1.176 | 0.789 | 1 | 0.716 |
| 0.837 | 7.79 | 0.844 | 1.026 | 0.459 | 1 | 0.716 |
| 0.873 | 6.717 | 0.745 | 1.176 | 0.789 | 0.998 | 0.413 |
| 0.495 | 9.864 | 0.984 | 1.176 | 0.789 | 1 | 0.716 |
| 0.783 | 9.775 | 0.982 | 1.176 | 0.789 | 1 | 0.716 |
| . | . | . | . | . | . | . |
| 0.524 | 1.955 | 0.4 | 1.176 | 0.789 | 0.994 | 0.383 |
| 0.873 | 6.717 | 0.745 | 1.176 | 0.789 | 0.998 | 0.413 |
| 0.59 | 9.566 | 0.973 | 1.142 | 0.701 | 1 | 0.716 |
| 0.697 | 7.889 | 0.858 | 1.006 | 0.373 | 1 | 0.716 |
| 0.332 | 2.139 | 0.418 | 0.952 | 0.349 | 0.96 | 0.336 |
| 0.641 | 7.518 | 0.807 | 1.026 | 0.459 | 1 | 0.716 |
| 0.98 | 6.254 | 0.724 | 1.176 | 0.789 | 1 | 0.716 |
| 0.717 | 9.317 | 0.954 | 1.176 | 0.789 | 1 | 0.716 |

| | | | | | | |
|---|---|---|---|---|---|---|
| . | . | . | . | . | . | . |
| 0.507 | 6.164 | 0.717 | 1.026 | 0.459 | 1 | 0.716 |
| 0.538 | 4.851 | 0.626 | 1.083 | 0.538 | 1 | 0.716 |
| 0.95 | 9.409 | 0.967 | 1.176 | 0.789 | 1 | 0.716 |
| 0.773 | 2.601 | 0.459 | 1.026 | 0.459 | 1 | 0.716 |
| 0.751 | 7.622 | 0.823 | 1.138 | 0.647 | 1 | 0.716 |
| 0.174 | 1.223 | 0.321 | 1.176 | 0.789 | 1 | 0.716 |
| . | . | . | . | . | . | . |
| 0.576 | 5.475 | 0.665 | 1.172 | 0.73 | 0.84 | 0.303 |
| 0.918 | 5.528 | 0.668 | 1.026 | 0.459 | 1 | 0.716 |
| 0.818 | 6.252 | 0.724 | 1.138 | 0.647 | 1 | 0.716 |
| 0.95 | 3.698 | 0.545 | 1.138 | 0.647 | 1 | 0.716 |
| 0.678 | 10.003 | 0.997 | 1.176 | 0.789 | 1 | 0.716 |
| 0.763 | 6.736 | 0.746 | 0.221 | 0.244 | 1 | 0.716 |
| 0.769 | 6.576 | 0.739 | 1.176 | 0.789 | 1 | 0.716 |
| 0.797 | 8.01 | 0.882 | 1.176 | 0.789 | 1 | 0.716 |
| 0.8 | 7.39 | 0.791 | 1.312 | 0.947 | 1 | 0.716 |
| 0.522 | 2.071 | 0.411 | 1.019 | 0.393 | 1 | 0.716 |
| 0.602 | 4.684 | 0.614 | 1.018 | 0.391 | 1 | 0.716 |
| 0.8 | 7.39 | 0.791 | 1.312 | 0.947 | 1 | 0.716 |
| 0.522 | 4.691 | 0.614 | 1.172 | 0.73 | 1 | 0.716 |
| 0.873 | 6.717 | 0.745 | 1.176 | 0.789 | 0.999 | 0.427 |
| 0.522 | 6.706 | 0.745 | 1.172 | 0.73 | 1 | 0.716 |
| 0.8 | 7.39 | 0.791 | 1.312 | 0.947 | 1 | 0.716 |
| 0.652 | 1.961 | 0.401 | 0.31 | 0.277 | 0.968 | 0.341 |
| . | . | . | . | . | . | . |
| 0.522 | 2.071 | 0.411 | 1.019 | 0.393 | 1 | 0.716 |
| 0.853 | 5.938 | 0.699 | 1.026 | 0.459 | 1 | 0.716 |
| 0.711 | 7.224 | 0.775 | 1.026 | 0.459 | 1 | 0.716 |
| 0.692 | 3.638 | 0.541 | 1.026 | 0.459 | 0.995 | 0.388 |
| 0.477 | 4.463 | 0.599 | 1.122 | 0.568 | 1 | 0.716 |
| 0.822 | 8.006 | 0.881 | 1.172 | 0.73 | 1 | 0.716 |
| 0.174 | 1.223 | 0.321 | 1.176 | 0.789 | 1 | 0.716 |
| 0.8 | 7.39 | 0.791 | 1.312 | 0.947 | 1 | 0.716 |
| 0.853 | 5.938 | 0.699 | 1.026 | 0.459 | 1 | 0.716 |
| 0.697 | 7.889 | 0.858 | 1.006 | 0.373 | 1 | 0.716 |
| 0.409 | 5.443 | 0.663 | 1.172 | 0.73 | 1 | 0.716 |
| . | . | . | . | . | . | . |
| 0.818 | 3.052 | 0.496 | 1.176 | 0.789 | 1 | 0.716 |
| 0.939 | 7.905 | 0.865 | 1.026 | 0.459 | 1 | 0.716 |
| 0.526 | 7.888 | 0.858 | 1.026 | 0.459 | 1 | 0.716 |
| 0.95 | 9.409 | 0.967 | 1.176 | 0.789 | 1 | 0.716 |
| 0.8 | 7.39 | 0.791 | 1.312 | 0.947 | 1 | 0.716 |

| | | | | | | |
|---|---|---|---|---|---|---|
| 0.978 | 7.679 | 0.834 | 1.176 | 0.789 | 1 | 0.716 |
| 0.853 | 5.938 | 0.699 | 1.026 | 0.459 | 1 | 0.716 |
| 0.887 | 7.66 | 0.828 | 1.138 | 0.647 | 1 | 0.716 |

| phastCons30\ | phastCons30\ | SiPhy_29way_ | SiPhy_29way_ | Interpro_dom | GTEx_V8_ger | GTEx_V8_tiss |
|---|---|---|---|---|---|---|
| . | . | . | . | . | . | . |
| . | . | . | . | . | . | . |
| 1 | 0.863 | 16.205 | 0.819 | .;.;.;.;.;. | . | . |
| 1 | 0.863 | 20.734 | 0.997 | .;.;.;.;. | . | . |
| 0.95 | 0.418 | 17.341 | 0.872 | .;.;. | . | . |
| 1 | 0.863 | 13.462 | 0.607 | .;.;.;. | . | . |
| 1 | 0.863 | 14.298 | 0.659 | .;.;. | . | . |
| 0.417 | 0.259 | 14.533 | 0.675 | .;.;.;.;.;.;.;. | . | . |
| 0.997 | 0.62 | 19.21 | 0.937 | .;.;AAA+ ATPa | . | . |
| 0.998 | 0.659 | 16.148 | 0.814 | Ribosomal pr | . | . |
| 0.997 | 0.62 | 15.996 | 0.8 | .;.;.;.;.;. | . | . |
| 1 | 0.863 | 16.205 | 0.819 | .;.;.;.;.;. | . | . |
| 1 | 0.863 | 7.758 | 0.281 | .;.;.;. | . | . |
| 1 | 0.863 | 19.512 | 0.951 | Lysine-specifi | . | . |
| . | . | . | . | . | . | . |
| 1 | 0.863 | 10.468 | 0.438 | Sox developn | . | . |
| 1 | 0.863 | 7.564 | 0.27 | .;.;. | . | . |
| 1 | 0.863 | 14.117 | 0.647 | . | . | . |
| 0.997 | 0.62 | 15.996 | 0.8 | .;.;.;.;.;. | . | . |
| 0.709 | 0.311 | 13.616 | 0.616 | .;.;.;. | . | . |
| 1 | 0.863 | 18.89 | 0.924 | Sterile alpha | . | . |
| . | . | . | . | . | . | . |
| 0.752 | 0.322 | 13.864 | 0.631 | .;. | . | . |
| 1 | 0.863 | 16.238 | 0.822 | .;.;.;.;.;.;. | . | . |
| 1 | 0.863 | 16.343 | 0.829 | .;.;.;. | . | . |
| 0.832 | 0.346 | 13.658 | 0.618 | .;.;.;.;.;. | . | . |
| 0.999 | 0.704 | 14.347 | 0.662 | .;.;.;. | . | . |
| 0.344 | 0.248 | 8.579 | 0.328 | Ribosomal pr | . | . |
| 0.996 | 0.595 | 15.832 | 0.785 | . | . | . |
| 0.97 | 0.45 | 19.356 | 0.944 | Zinc finger, Pl | . | . |
| 0.997 | 0.62 | 15.996 | 0.8 | .;.;.;.;.;. | . | . |
| 0.999 | 0.704 | 9.114 | 0.359 | Distal-less-lik | . | . |
| 0.967 | 0.444 | 18.487 | 0.908 | .;.;. | . | . |
| 0.998 | 0.659 | 16.148 | 0.814 | Ribosomal pr | . | . |
| 0.964 | 0.439 | 12.411 | 0.548 | Nuclear recep | . | . |
| 0.878 | 0.365 | 20.583 | 0.994 | Nuclear recep | . | . |
| 0.991 | 0.532 | 13.751 | 0.624 | .;.;.;.;. | . | . |
| 1 | 0.863 | 15.037 | 0.714 | .;. | . | . |
| 0.914 | 0.386 | 14.102 | 0.646 | .;. | . | . |
| 1 | 0.863 | 18.867 | 0.923 | .;.;.;.;. | . | . |
| 0.944 | 0.412 | 16.961 | 0.861 | .;.;.;.;.;.;.;. | . | . |
| 0.998 | 0.659 | 20.422 | 0.991 | Ephrin recept | . | . |

| | | | | | | |
|---|---|---|---|---|---|---|
| 1 | 0.863 | 16.205 | 0.819 | .;.;.;.;.;. | . | . |
| 0.455 | 0.266 | 8.429 | 0.319 | .;.;. | . | . |
| 1 | 0.863 | 13.454 | 0.606 | .;. | . | . |
| 0.935 | 0.403 | 15.365 | 0.742 | .;. | . | . |
| 1 | 0.863 | 18.89 | 0.924 | Sterile alpha | . | . |
| 0.927 | 0.396 | 19.305 | 0.941 | .;. | . | . |
| . | . | . | . | . | . | . |
| 0.01 | 0.119 | 6.691 | 0.224 | .;.;.;.;.;.;.;. | . | . |
| 1 | 0.863 | 17.623 | 0.88 | TGF-beta, pro | . | . |
| 0.729 | 0.316 | 19.397 | 0.946 | .;.;.;. | . | . |
| 0.977 | 0.468 | 14.713 | 0.689 | JmjC domain | . | . |
| 0.598 | 0.289 | 18.146 | 0.896 | Transcription | . | . |
| 0.763 | 0.325 | 5.223 | 0.147 | .;.;. | . | . |
| . | . | . | . | . | . | . |
| 0.449 | 0.265 | 11.887 | 0.519 | .;.;. | . | . |
| 1 | 0.863 | 20.026 | 0.975 | .;. | . | . |
| 0.998 | 0.659 | 19.221 | 0.938 | .;. | . | . |
| 0.998 | 0.659 | 14.452 | 0.67 | Nuclear recep | . | . |
| 0.999 | 0.704 | 6.719 | 0.225 | .;. | . | . |
| 0.529 | 0.277 | 17.082 | 0.865 | . | . | . |
| 0.256 | 0.233 | 18.762 | 0.918 | .;. | . | . |
| . | . | . | . | . | . | . |
| 1 | 0.863 | 16.205 | 0.819 | .;.;.;.;.;. | . | . |
| 1 | 0.863 | 14 | 0.639 | . | . | . |
| 0.965 | 0.44 | 15.152 | 0.724 | .;. | . | . |
| 0.997 | 0.62 | 15.996 | 0.8 | .;.;.;.;.;. | . | . |
| 1 | 0.863 | 16.033 | 0.804 | . | . | . |
| 0.986 | 0.501 | 14.192 | 0.652 | Transcription | . | . |
| 0.084 | 0.188 | 17.306 | 0.871 | .;.;.;.;.;.;.;. | . | . |
| . | . | . | . | . | . | . |
| 1 | 0.863 | 12.67 | 0.562 | .;.;.;.;. | . | . |
| 0.817 | 0.34 | 11.808 | 0.514 | Histidine kina | . | . |
| 0.995 | 0.577 | 12.592 | 0.558 | .;. | . | . |
| 0.997 | 0.62 | 19.21 | 0.937 | .;.;AAA+ ATPa | . | . |
| 1 | 0.863 | 16.205 | 0.819 | .;.;.;.;.;. | . | . |
| 1 | 0.863 | 16.205 | 0.819 | .;.;.;.;.;. | . | . |
| 1 | 0.863 | 15.84 | 0.786 | .;.;. | . | . |
| 0.996 | 0.595 | 19.451 | 0.949 | .;.;.;.;.;.;.;.;. | . | . |
| 0.995 | 0.577 | 8.707 | 0.335 | Rho GTPase-a | . | . |
| 0.987 | 0.506 | 15.998 | 0.801 | .;. | . | . |
| 1 | 0.863 | 15.342 | 0.74 | .;. | . | . |
| 1 | 0.863 | 10.468 | 0.438 | Sox developm | . | . |
| 0.574 | 0.285 | 14.143 | 0.649 | Histidine kina | . | . |

| | | | | | | | |
|---|---|---|---|---|---|---|---|
| | 0.701 | 0.31 | 18.165 | 0.897 | .;.;.;.;.;. | . | . |
| . | . | . | . | | . | . | . |
| | 0.996 | 0.595 | 9.562 | 0.385 | .;.;.;.;. | . | . |
| | 1 | 0.863 | 17.908 | 0.888 | TGF-beta, prc | . | . |
| | 1 | 0.863 | 17.623 | 0.88 | TGF-beta, prc | . | . |
| | 0.975 | 0.462 | 18.649 | 0.914 | . | . | . |
| | 0.999 | 0.704 | 18.99 | 0.928 | .;.;.;. | . | . |
| | 1 | 0.863 | 16.205 | 0.819 | .;.;.;.;.;. | . | . |
| | 0.976 | 0.465 | 17.684 | 0.882 | .;.;.;.;.;.;.;.;.;.;. | . | . |
| | 1 | 0.863 | 16.665 | 0.851 | .;. | . | . |
| . | . | . | . | | . | . | . |
| | 0.95 | 0.418 | 17.341 | 0.872 | .;.;. | . | . |
| | 0.976 | 0.465 | 17.332 | 0.872 | TGF-beta, prc | . | . |
| . | . | . | . | | . | . | . |
| . | . | . | . | | . | . | . |
| | 0.997 | 0.62 | 19.21 | 0.937 | .;.;AAA+ ATPa | . | . |
| . | . | . | . | | . | . | . |
| | 0.995 | 0.577 | 15.748 | 0.777 | . | . | . |
| | 1 | 0.863 | 13.371 | 0.602 | .;.;. | . | . |
| . | . | . | . | | . | . | . |
| | 1 | 0.863 | 20.257 | 0.984 | .;. | . | . |
| | 1 | 0.863 | 15.001 | 0.711 | . | . | . |
| . | . | . | . | | . | . | . |
| | 1 | 0.863 | 16.205 | 0.819 | .;.;.;.;.;. | . | . |
| | 0.997 | 0.62 | 15.996 | 0.8 | .;.;.;.;.;. | . | . |
| | 0.994 | 0.563 | 20.139 | 0.981 | .;.;. | . | . |
| | 0.998 | 0.659 | 16.148 | 0.814 | Ribosomal pr | . | . |
| . | . | . | . | | . | . | . |
| | 0.997 | 0.62 | 11.913 | 0.52 | .;. | . | . |
| | 1 | 0.863 | 9.608 | 0.388 | SLC12A trans | . | . |
| | 0.567 | 0.284 | 19.796 | 0.965 | . | . | . |
| | 1 | 0.863 | 16.613 | 0.847 | Nuclear recep | . | . |
| | 1 | 0.863 | 11.981 | 0.524 | .;. | . | . |
| | 0.897 | 0.375 | 5.524 | 0.162 | .;.;.;.;.;.;.;.;.;.;.;. | . | . |
| | 0.529 | 0.277 | 17.082 | 0.865 | . | . | . |
| | 0.988 | 0.511 | 20.099 | 0.979 | .;. | . | . |
| | 0.962 | 0.435 | 18.532 | 0.909 | .;. | . | . |
| | 0.98 | 0.477 | 19.378 | 0.945 | SH3 domain\| | . | . |
| | 1 | 0.863 | 13.652 | 0.618 | . | . | . |
| | 1 | 0.863 | 14.603 | 0.68 | TGF-beta, prc | . | . |
| . | . | . | . | | . | . | . |
| . | . | . | . | | . | . | . |
| | 0.567 | 0.284 | 19.796 | 0.965 | . | . | . |

| | | | | | | |
|---|---|---|---|---|---|---|
| . | . | . | . | . | . | . |
| 1 | 0.863 | 16.205 | 0.819 | .;.;.;.;.;. | . | . |
| 0.529 | 0.277 | 17.082 | 0.865 | . | . | . |
| 0.098 | 0.193 | 18.131 | 0.895 | Transcription | . | . |
| 0.024 | 0.146 | 14.766 | 0.693 | Rho GTPase-a | . | . |
| 0.987 | 0.506 | 12.593 | 0.558 | .;. | . | . |
| 1 | 0.863 | 20.663 | 0.997 | .;. | . | . |
| 1 | 0.863 | 12.794 | 0.569 | .;. | . | . |
| 1 | 0.863 | 18.787 | 0.919 | Matrin/U1-C- | . | . |
| 0.992 | 0.541 | 15.485 | 0.753 | .;.;.;.;.;.;.;. | . | . |
| 0.994 | 0.563 | 12.752 | 0.567 | p53, tetrame | . | . |
| 1 | 0.863 | 14.183 | 0.651 | .;.;.;.;.;.;.;. | . | . |
| 0.991 | 0.532 | 15.617 | 0.765 | .;.;. | . | . |
| . | . | . | . | . | . | . |
| 0.822 | 0.342 | 10.22 | 0.424 | .;.;. | . | . |
| 0.982 | 0.484 | 10.991 | 0.468 | . | . | . |
| 1 | 0.863 | 15.385 | 0.744 | .;.;.;.;.;.;.;.;.;.;. | . | . |
| . | . | . | . | . | . | . |
| 0.998 | 0.659 | 16.148 | 0.814 | Ribosomal pr | . | . |
| 0.075 | 0.184 | 14.662 | 0.685 | .;. | . | . |
| 1 | 0.863 | 18.365 | 0.903 | Amino acid p | . | . |
| 1 | 0.863 | 20.663 | 0.997 | .;. | . | . |
| . | . | . | . | . | . | . |
| 1 | 0.863 | 15.222 | 0.73 | Zinc finger C2 | . | . |
| 0.988 | 0.511 | 11.724 | 0.51 | .;.;.;. | . | . |
| 0.64 | 0.297 | 14.064 | 0.643 | . | . | . |
| 0.98 | 0.477 | 15.293 | 0.736 | Nuclear recep | . | . |
| 0.982 | 0.484 | 10.332 | 0.43 | .;.;. | . | . |
| 0.886 | 0.369 | 18.46 | 0.907 | .;. | . | . |
| 0.999 | 0.704 | 19.983 | 0.973 | Zinc finger C2 | . | . |
| 0.084 | 0.188 | 17.306 | 0.871 | .;.;.;.;.;.;.;. | . | . |
| 1 | 0.863 | 20.663 | 0.997 | .;. | . | . |
| . | . | . | . | . | . | . |
| 0.966 | 0.442 | 18.249 | 0.899 | .;.;.;.;. | . | . |
| 1 | 0.863 | 18.872 | 0.923 | Zinc finger, Pl | . | . |
| 1 | 0.863 | 18.603 | 0.912 | .;.;.;.;.;.;. | . | . |
| 1 | 0.863 | 15.677 | 0.771 | .;. | . | . |
| 1 | 0.863 | 20.395 | 0.99 | Nuclear recep | . | . |
| . | . | . | . | . | . | . |
| 0.084 | 0.188 | 17.306 | 0.871 | .;.;.;.;.;.;.;. | . | . |
| 0.889 | 0.37 | 14.14 | 0.648 | Transcription | . | . |
| 0.959 | 0.431 | 10.98 | 0.467 | Nuclear recep | . | . |
| 1 | 0.863 | 13.986 | 0.639 | .;.;.;.;. | . | . |

| | | | | | | |
|---|---|---|---|---|---|---|
| 0.016 | 0.134 | 19.956 | 0.972 | Rho GTPase-a | . | . |
| 0.996 | 0.595 | 15.874 | 0.789 | .;. | . | . |
| . | . | . | . | . | . | . |
| . | . | . | . | . | . | . |
| 0.98 | 0.477 | 15.293 | 0.736 | Nuclear recep | . | . |
| 0.944 | 0.412 | 19.733 | 0.962 | . | . | . |
| 1 | 0.863 | 15.435 | 0.749 | .;.;. | . | . |
| 1 | 0.863 | 18.365 | 0.903 | Amino acid pe | . | . |
| 0.975 | 0.462 | 20.475 | 0.992 | Nuclear recep | . | . |
| . | . | . | . | . | . | . |
| 0.979 | 0.474 | 11.107 | 0.474 | .;.;.;.;.;.;.;. | . | . |
| 0.984 | 0.492 | 13.758 | 0.624 | .;.;. | . | . |
| 0.998 | 0.659 | 8.273 | 0.31 | Histidine kina | . | . |
| 0.998 | 0.659 | 8.273 | 0.31 | Histidine kina | . | . |
| . | . | . | . | . | . | . |
| . | . | . | . | . | . | . |
| 0.991 | 0.532 | 19.252 | 0.939 | .;. | . | . |
| 1 | 0.863 | 15.482 | 0.753 | .;. | . | . |
| 0.994 | 0.563 | 19.942 | 0.972 | .;. | . | . |
| . | . | . | . | . | . | . |
| 1 | 0.863 | 15.222 | 0.73 | Zinc finger C2 | . | . |
| 0.506 | 0.274 | 20.367 | 0.989 | .;.;. | . | . |
| 1 | 0.863 | 19.512 | 0.951 | Lysine-specifi | . | . |
| 0.996 | 0.595 | 15.648 | 0.768 | .;.;. | . | . |
| 1 | 0.863 | 15.84 | 0.786 | .;.;. | . | . |
| 0.996 | 0.595 | 20.367 | 0.989 | Zinc finger C2 | . | . |
| . | . | . | . | . | . | . |
| 0.979 | 0.474 | 11.107 | 0.474 | .;.;.;.;.;.;.;. | . | . |
| 0.998 | 0.659 | 10.229 | 0.424 | .;.;. | . | . |
| 1 | 0.863 | 16.205 | 0.819 | .;.;.;.;.;. | . | . |
| 0.084 | 0.188 | 17.306 | 0.871 | .;.;.;.;.;.;.;. | . | . |
| 0.984 | 0.492 | 13.758 | 0.624 | .;.;. | . | . |
| 0.084 | 0.188 | 17.306 | 0.871 | .;.;.;.;.;.;.;. | . | . |
| 0.529 | 0.277 | 17.082 | 0.865 | . | . | . |
| 0.838 | 0.348 | 15.001 | 0.711 | . | . | . |
| 1 | 0.863 | 11.269 | 0.484 | .;.;. | . | . |
| 0.944 | 0.412 | 16.961 | 0.861 | .;.;.;.;.;.;.;. | . | . |
| 1 | 0.863 | 19.512 | 0.951 | Lysine-specifi | . | . |
| 1 | 0.863 | 13.454 | 0.606 | .;. | . | . |
| 0.017 | 0.135 | 18.248 | 0.899 | . | . | . |
| . | . | . | . | . | . | . |
| 1 | 0.863 | 9.964 | 0.409 | .;.;AAA+ ATPa | . | . |
| 1 | 0.863 | 18.872 | 0.923 | Zinc finger, PI | . | . |

| | | | | | | |
|---|---|---|---|---|---|---|
| . | . | . | . | . | . | . |
| 0.084 | 0.188 | 17.306 | 0.871 | .;.;.;.;.;.;.;. | . | . |
| 1 | 0.863 | 19.512 | 0.951 | Lysine-specifi | . | . |
| 1 | 0.863 | 18.872 | 0.923 | Zinc finger, PI | . | . |
| 0.98 | 0.477 | 15.293 | 0.736 | Nuclear recep | . | . |
| 1 | 0.863 | 18.872 | 0.923 | Zinc finger, PI | . | . |
| 1 | 0.863 | 8.605 | 0.329 | . | . | . |
| 0.998 | 0.659 | 19.945 | 0.972 | Zinc finger C2 | . | . |
| . | . | . | . | . | . | . |
| 1 | 0.863 | 15.677 | 0.771 | .;. | . | . |
| 0.944 | 0.412 | 16.961 | 0.861 | .;.;.;.;.;.;.;. | . | . |
| 0.15 | 0.209 | 13.601 | 0.615 | .;. | . | . |
| . | . | . | . | . | . | . |
| 0.015 | 0.131 | 13.29 | 0.597 | .;.;. | . | . |
| 0.966 | 0.442 | 18.249 | 0.899 | .;.;.;.;. | . | . |
| 0.996 | 0.595 | 13.077 | 0.585 | ARID DNA-bir | . | . |
| 0.072 | 0.182 | 18.869 | 0.923 | . | . | . |
| . | . | . | . | . | . | . |
| 1 | 0.863 | 14.482 | 0.672 | .;.;Calponin h | . | . |
| 1 | 0.863 | 17.587 | 0.879 | .;. | . | . |
| 0.072 | 0.182 | 18.869 | 0.923 | . | . | . |
| 1 | 0.863 | 16.59 | 0.846 | .;.;. | . | . |
| . | . | . | . | . | . | . |
| 0.998 | 0.659 | 13.692 | 0.62 | .;. | . | . |
| . | . | . | . | . | . | . |
| 1 | 0.863 | 13.233 | 0.594 | TGF-beta, prc | . | . |
| 0.991 | 0.532 | 14.838 | 0.698 | .;. | . | . |
| 1 | 0.863 | 17.549 | 0.878 | .;.;.;.;.;.;.;.;.;.;.;. | . | . |
| . | . | . | . | . | . | . |
| 1 | 0.863 | 18.3 | 0.901 | . | . | . |
| 0.896 | 0.374 | 18.098 | 0.894 | .;.;.;. | . | . |
| 0.995 | 0.577 | 20.125 | 0.98 | .;. | . | . |
| 1 | 0.863 | 16.983 | 0.862 | .;.;. | . | . |
| 0.999 | 0.704 | 19.378 | 0.945 | SH3 domain;S | . | . |
| . | . | . | . | . | . | . |
| 0.164 | 0.213 | 10.787 | 0.456 | .;.;. | . | . |
| 0.995 | 0.577 | 20.125 | 0.98 | .;. | . | . |
| 0.999 | 0.704 | 18.092 | 0.894 | .;.;.;.;. | . | . |
| 0.827 | 0.344 | 12.011 | 0.526 | .;.;.;.;.;. | . | . |
| 0.999 | 0.704 | 8.634 | 0.331 | . | . | . |
| 1 | 0.863 | 18.476 | 0.907 | .;. | . | . |
| 0.985 | 0.496 | 20.579 | 0.994 | Tubby, C-tern | . | . |
| 0.077 | 0.185 | 17.907 | 0.888 | Zinc finger, G | . | . |

| | | | | | | |
|---|---|---|---|---|---|---|
| . | . | . | . | . | . | . |
| 1 | 0.863 | 13.233 | 0.594 | TGF-beta, prc | . | . |
| 0.999 | 0.704 | 16.078 | 0.808 | .;.;. | . | . |
| 1 | 0.863 | 19.267 | 0.94 | Sterile alpha | . | . |
| 0.999 | 0.704 | 19.128 | 0.934 | .;. | . | . |
| 1 | 0.863 | 15.275 | 0.734 | Zinc finger C2 | . | . |
| 1 | 0.863 | 9.557 | 0.385 | Transforming | . | . |
| . | . | . | . | . | . | . |
| 0.088 | 0.19 | 13.1 | 0.586 | .;.;.;. | . | . |
| 1 | 0.863 | 19.025 | 0.929 | .;. | . | . |
| 0.991 | 0.532 | 15.617 | 0.765 | .;.;. | . | . |
| 1 | 0.863 | 15.998 | 0.801 | .;. | . | . |
| 1 | 0.863 | 19.036 | 0.93 | .;.;SNF2-relat | . | . |
| 0.998 | 0.659 | 19.056 | 0.93 | .;. | . | . |
| 1 | 0.863 | 18.7 | 0.916 | . | . | . |
| 0.972 | 0.455 | 14.842 | 0.699 | .;.;. | . | . |
| 1 | 0.863 | 14.094 | 0.645 | .;.;.;.;.;.;.;.;.;.;. | . | . |
| 0.892 | 0.372 | 10.944 | 0.465 | . | . | . |
| 0.998 | 0.659 | 11.775 | 0.512 | .;. | . | . |
| 1 | 0.863 | 14.094 | 0.645 | .;.;.;.;.;.;.;.;.;.;. | . | . |
| 1 | 0.863 | 9.88 | 0.404 | .;.;. | . | . |
| 0.99 | 0.524 | 20.125 | 0.98 | .;. | . | . |
| 1 | 0.863 | 16.59 | 0.846 | .;.;. | . | . |
| 1 | 0.863 | 14.094 | 0.645 | .;.;.;.;.;.;.;.;.;.;. | . | . |
| 0.078 | 0.185 | 14.945 | 0.707 | .;.;.;. | . | . |
| . | . | . | . | . | . | . |
| 0.892 | 0.372 | 10.944 | 0.465 | . | . | . |
| 1 | 0.863 | 18.99 | 0.928 | .;.;.;. | . | . |
| 0.072 | 0.182 | 18.869 | 0.923 | . | . | . |
| 0.942 | 0.41 | 18.551 | 0.91 | Zinc finger C2 | . | . |
| 1 | 0.863 | 9.908 | 0.406 | .;.;.;. | . | . |
| 0.999 | 0.704 | 19.474 | 0.95 | Zinc finger C2 | . | . |
| 1 | 0.863 | 9.557 | 0.385 | Transforming | . | . |
| 1 | 0.863 | 14.094 | 0.645 | .;.;.;.;.;.;.;.;.;.;. | . | . |
| 1 | 0.863 | 18.99 | 0.928 | .;.;.;. | . | . |
| 0.827 | 0.344 | 12.011 | 0.526 | .;.;.;.;.;. | . | . |
| 1 | 0.863 | 10.498 | 0.44 | Insulin-like gr | . | . |
| . | . | . | . | . | . | . |
| 1 | 0.863 | 10.897 | 0.462 | . | . | . |
| 0.994 | 0.563 | 20.178 | 0.982 | Ephrin recept | . | . |
| 1 | 0.863 | 12.996 | 0.58 | TGF-beta, prc | . | . |
| 1 | 0.863 | 19.267 | 0.94 | Sterile alpha | . | . |
| 1 | 0.863 | 14.094 | 0.645 | .;.;.;.;.;.;.;.;.;.;. | . | . |

| | | | | | | |
|---|---|---|---|---|---|---|
| 0.962 | 0.435 | 20.557 | 0.993 | .;. | . | . |
| 1 | 0.863 | 18.99 | 0.928 | .;.;.;. | . | . |
| 0.684 | 0.306 | 15.138 | 0.723 | .;.;.;.;. | . | . |

| dbscSNV_AD | dbscSNV_RF_ | AF | AF_popmax | AF_male | AF_female | AF_raw |
|---|---|---|---|---|---|---|
| . | . | . | 0 | . | . | . |
| . | . | . | 0 | . | . | . |
| . | . | 0.0023 | 0.004 | 0.0023 | 0.0024 | 0.0023 |
| . | . | 1.20E-05 | 2.66E-05 | 7.39E-06 | 1.74E-05 | 1.19E-05 |
| . | . | 0.0017 | 0.0022 | 0.0016 | 0.0018 | 0.0017 |
| . | . | 3.99E-06 | 5.44E-05 | 0 | 8.69E-06 | 3.98E-06 |
| . | . | 3.58E-05 | 7.03E-05 | 4.41E-05 | 2.60E-05 | 3.58E-05 |
| . | . | 1.59E-05 | 3.52E-05 | 2.21E-05 | 8.66E-06 | 1.59E-05 |
| . | . | 0.0021 | 0.004 | 0.0021 | 0.0021 | 0.0021 |
| . | . | 0.0011 | 0.002 | 0.001 | 0.0012 | 0.0011 |
| . | . | 0.0011 | 0.002 | 0.0011 | 0.001 | 0.0011 |
| . | . | 0.0023 | 0.004 | 0.0023 | 0.0024 | 0.0023 |
| . | . | . | 0 | . | . | . |
| . | . | 0.001 | 0.0013 | 0.0011 | 0.001 | 0.001 |
| . | . | . | 0 | . | . | . |
| . | . | 0.0006 | 0.0032 | 0.0007 | 0.0004 | 0.0005 |
| . | . | . | 0 | . | . | . |
| 0.0919 | 0.394 | 0 | 0 | 0 | 0 | 3.98E-06 |
| . | . | 0.0011 | 0.002 | 0.0011 | 0.001 | 0.0011 |
| . | . | 6.35E-05 | 2.00E-04 | 6.07E-05 | 6.66E-05 | 7.04E-05 |
| . | . | 3.58E-05 | 2.00E-04 | 4.42E-05 | 2.60E-05 | 3.58E-05 |
| 0.0003 | 0.094 | . | 0 | . | . | . |
| . | . | 7.24E-05 | 5.00E-04 | 8.91E-05 | 5.27E-05 | 7.16E-05 |
| . | . | 4.57E-05 | 8.11E-05 | 2.30E-05 | 7.23E-05 | 4.37E-05 |
| . | . | . | 0 | . | . | . |
| . | . | 1.60E-05 | 1.00E-04 | 2.22E-05 | 8.68E-06 | 1.59E-05 |
| . | . | 3.55E-05 | 8.17E-05 | 4.04E-05 | 2.96E-05 | 3.60E-05 |
| . | . | 4.84E-05 | 8.13E-05 | 5.21E-05 | 4.40E-05 | 4.81E-05 |
| . | . | . | 0 | . | . | . |
| . | . | 0.0002 | 4.00E-04 | 0.0002 | 0.0003 | 0.0002 |
| . | . | 0.0011 | 0.002 | 0.0011 | 0.001 | 0.0011 |
| . | . | 0.0008 | 5.00E-04 | 0.0008 | 0.0008 | 0.0008 |
| . | . | 8.40E-05 | 0 | 7.39E-05 | 9.60E-05 | 8.35E-05 |
| . | . | 0.0011 | 0.002 | 0.001 | 0.0012 | 0.0011 |
| . | . | 0.0004 | 0.0032 | 0.0006 | 0.0001 | 0.0004 |
| . | . | 4.00E-06 | 8.84E-06 | 7.40E-06 | 0 | 3.98E-06 |
| . | . | 3.99E-06 | 8.86E-06 | 0 | 8.69E-06 | 3.98E-06 |
| . | . | . | 0 | . | . | . |
| . | . | 0.0014 | 0.0012 | 0.0013 | 0.0015 | 0.0014 |
| 0.9999 | 0.964 | 1.20E-05 | 5.79E-05 | 7.37E-06 | 1.73E-05 | 1.19E-05 |
| . | . | 0.0003 | 5.00E-04 | 0.0003 | 0.0003 | 0.0003 |
| . | . | 1.19E-05 | 5.44E-05 | 1.47E-05 | 8.66E-06 | 1.19E-05 |

| | | | | | | |
|---|---|---|---|---|---|---|
| . | . | 0.0023 | 0.004 | 0.0023 | 0.0024 | 0.0023 |
| . | . | . | 0 | . | . | . |
| . | . | 0.0011 | 0.0017 | 0.0011 | 0.0012 | 0.0012 |
| . | . | 3.98E-06 | 8.80E-06 | 7.36E-06 | 0 | 3.98E-06 |
| . | . | 3.58E-05 | 2.00E-04 | 4.42E-05 | 2.60E-05 | 3.58E-05 |
| . | . | . | 0 | . | . | . |
| . | . | . | 0 | . | . | . |
| . | . | 3.69E-05 | 8.24E-05 | 3.79E-05 | 3.58E-05 | 3.98E-05 |
| . | . | 0.0002 | 4.00E-04 | 0.0002 | 0.0002 | 0.0002 |
| . | . | 7.56E-05 | 1.00E-04 | 7.36E-05 | 7.79E-05 | 7.95E-05 |
| . | . | 1.60E-05 | 1.00E-04 | 2.22E-05 | 8.76E-06 | 1.60E-05 |
| . | . | 7.56E-05 | 1.00E-04 | 5.89E-05 | 9.52E-05 | 7.56E-05 |
| . | . | 3.59E-05 | 1.00E-04 | 2.22E-05 | 5.21E-05 | 3.58E-05 |
| . | . | . | 0 | . | . | . |
| . | . | 0.0004 | 8.00E-04 | 0.0004 | 0.0004 | 0.0004 |
| . | . | 2.78E-05 | 1.00E-04 | 2.94E-05 | 2.60E-05 | 2.78E-05 |
| . | . | 3.98E-06 | 8.81E-06 | 7.37E-06 | 0 | 7.95E-06 |
| . | . | 0.0004 | 7.00E-04 | 0.0005 | 0.0004 | 0.0004 |
| . | . | . | 0 | . | . | . |
| . | . | 0.0014 | 0.0021 | 0.0016 | 0.0012 | 0.0014 |
| . | . | 0.0003 | 6.00E-04 | 0.0003 | 0.0003 | 0.0003 |
| . | . | 0.0004 | 0.0011 | 0.0003 | 0.0004 | 0.0004 |
| . | . | 0.0023 | 0.004 | 0.0023 | 0.0024 | 0.0023 |
| . | . | 0.001 | 0.0017 | 0.001 | 0.001 | 0.001 |
| . | . | . | 0 | . | . | . |
| . | . | 0.0011 | 0.002 | 0.0011 | 0.001 | 0.0011 |
| . | . | 2.01E-05 | 4.43E-05 | 2.96E-05 | 8.78E-06 | 2.00E-05 |
| . | . | 9.55E-05 | 5.00E-04 | 5.89E-05 | 0.0001 | 9.54E-05 |
| . | . | 0.002 | 0.0033 | 0.002 | 0.0019 | 0.002 |
| 0.0002 | 0.006 | 3.21E-05 | 7.09E-05 | 2.96E-05 | 3.52E-05 | 3.20E-05 |
| . | . | 3.99E-06 | 3.27E-05 | 7.38E-06 | 0 | 7.95E-06 |
| . | . | 1.61E-05 | 6.18E-05 | 2.24E-05 | 8.77E-06 | 1.99E-05 |
| . | . | . | 0 | . | . | . |
| . | . | 0.0021 | 0.004 | 0.0021 | 0.0021 | 0.0021 |
| . | . | 0.0023 | 0.004 | 0.0023 | 0.0024 | 0.0023 |
| . | . | 0.0023 | 0.004 | 0.0023 | 0.0024 | 0.0023 |
| . | . | . | 0 | . | . | . |
| . | . | 0.0002 | 4.00E-04 | 0.0002 | 0.0001 | 0.0002 |
| . | . | 3.98E-06 | 8.80E-06 | 7.36E-06 | 0 | 3.98E-06 |
| . | . | . | 0 | . | . | . |
| . | . | 0.0001 | 3.00E-04 | 0.0001 | 0.0002 | 0.0001 |
| . | . | 0.0006 | 0.0032 | 0.0007 | 0.0004 | 0.0005 |
| . | . | 9.15E-05 | 2.00E-04 | 9.57E-05 | 8.65E-05 | 9.15E-05 |

| | | | | | | |
|---|---|---|---|---|---|---|
| . | . | 7.56E-05 | 1.00E-04 | 8.84E-05 | 6.07E-05 | 7.56E-05 |
| . | . | 0 | 0 | 0 | 0 | 0.0013 |
| . | . | 0.0013 | 0.0021 | 0.0014 | 0.0012 | 0.0013 |
| . | . | . | 0 | . | . | . |
| . | . | 0.0002 | 4.00E-04 | 0.0002 | 0.0002 | 0.0002 |
| . | . | 7.97E-06 | 1.76E-05 | 1.47E-05 | 0 | 7.95E-06 |
| . | . | . | 0 | . | . | . |
| . | . | 0.0023 | 0.004 | 0.0023 | 0.0024 | 0.0023 |
| . | . | 1.99E-05 | 5.44E-05 | 2.21E-05 | 1.74E-05 | 2.39E-05 |
| . | . | . | 0 | . | . | . |
| . | . | 1.68E-05 | 3.63E-05 | 0 | 3.70E-05 | 1.59E-05 |
| . | . | 0.0017 | 0.0022 | 0.0016 | 0.0018 | 0.0017 |
| . | . | 2.78E-05 | 8.67E-05 | 2.21E-05 | 3.46E-05 | 3.18E-05 |
| . | . | . | 0 | . | . | . |
| . | . | . | 0 | . | . | . |
| . | . | 0.0021 | 0.004 | 0.0021 | 0.0021 | 0.0021 |
| . | . | 3.62E-05 | 6.19E-05 | 7.43E-06 | 7.01E-05 | 0.0007 |
| . | . | . | 0 | . | . | . |
| . | . | 7.56E-05 | 1.00E-04 | 6.62E-05 | 8.65E-05 | 7.56E-05 |
| . | . | 7.76E-05 | 9.81E-05 | 9.00E-05 | 6.28E-05 | 7.56E-05 |
| . | . | . | 0 | . | . | . |
| . | . | . | 0 | . | . | . |
| . | . | . | 0 | . | . | . |
| . | . | 0.0023 | 0.004 | 0.0023 | 0.0024 | 0.0023 |
| . | . | 0.0011 | 0.002 | 0.0011 | 0.001 | 0.0011 |
| . | . | 7.95E-05 | 2.00E-04 | 7.36E-05 | 8.65E-05 | 7.95E-05 |
| . | . | 0.0011 | 0.002 | 0.001 | 0.0012 | 0.0011 |
| . | . | . | 0 | . | . | . |
| . | . | 0.0002 | 5.00E-04 | 0.0002 | 0.0001 | 0.0001 |
| . | . | . | 0 | . | . | . |
| . | . | 0.0002 | 4.00E-04 | 0.0002 | 0.0002 | 0.0002 |
| . | . | 0.0004 | 0.0023 | 0.0004 | 0.0005 | 0.0004 |
| . | . | 0.0006 | 0.0012 | 0.0006 | 0.0006 | 0.0006 |
| . | . | 7.36E-05 | 4.00E-04 | 0.0001 | 3.59E-05 | 7.58E-05 |
| . | . | 0.0014 | 0.0021 | 0.0016 | 0.0012 | 0.0014 |
| . | . | . | 0 | . | . | . |
| . | . | 1.59E-05 | 5.78E-05 | 1.47E-05 | 1.73E-05 | 1.59E-05 |
| . | . | 1.61E-05 | 2.00E-04 | 0 | 3.51E-05 | 1.59E-05 |
| . | . | 2.39E-05 | 5.30E-05 | 2.95E-05 | 1.74E-05 | 2.39E-05 |
| . | . | 0.0003 | 7.00E-04 | 0.0003 | 0.0003 | 0.0003 |
| . | . | 0.0006 | 0.0021 | 0.0007 | 0.0004 | 0.0006 |
| . | . | 1.59E-05 | 6.53E-05 | 2.94E-05 | 0 | 1.59E-05 |
| . | . | 0.0002 | 4.00E-04 | 0.0002 | 0.0002 | 0.0002 |

| | | | | | | |
|---|---|---|---|---|---|---|
| . | . | . | 0 | . | . | . |
| . | . | 0.0023 | 0.004 | 0.0023 | 0.0024 | 0.0023 |
| . | . | 0.0014 | 0.0021 | 0.0016 | 0.0012 | 0.0014 |
| . | . | 4.37E-05 | 1.00E-04 | 3.68E-05 | 5.19E-05 | 4.37E-05 |
| . | . | 1.59E-05 | 3.52E-05 | 7.36E-06 | 2.60E-05 | 1.59E-05 |
| . | . | . | 0 | . | . | . |
| . | . | 0.0008 | 0.0021 | 0.0007 | 0.001 | 0.0008 |
| . | . | 0.0001 | 2.00E-04 | 0.0002 | 0.0001 | 0.0001 |
| . | . | . | 0 | . | . | . |
| 0.9909 | 0.614 | . | 0 | . | . | . |
| . | . | . | 0 | . | . | . |
| . | . | 3.98E-06 | 3.27E-05 | 7.36E-06 | 0 | 3.98E-06 |
| . | . | 0.0001 | 0.0016 | 0.0001 | 0.0001 | 0.0001 |
| . | . | 7.95E-06 | 1.76E-05 | 7.36E-06 | 8.65E-06 | 7.95E-06 |
| . | . | 0.0001 | 6.00E-04 | 7.36E-05 | 0.0001 | 0.0001 |
| . | . | . | 0 | . | . | . |
| . | . | . | 0 | . | . | . |
| . | . | 1.21E-05 | 2.67E-05 | 2.25E-05 | 0 | 1.19E-05 |
| . | . | 0.0011 | 0.002 | 0.001 | 0.0012 | 0.0011 |
| . | . | 5.20E-05 | 7.97E-05 | 4.43E-05 | 6.11E-05 | 5.17E-05 |
| . | . | 2.39E-05 | 2.00E-04 | 7.37E-06 | 4.34E-05 | 2.78E-05 |
| . | . | 0.0008 | 0.0021 | 0.0007 | 0.001 | 0.0008 |
| . | . | . | 0 | . | . | . |
| . | . | 4.77E-05 | 7.91E-05 | 6.62E-05 | 2.60E-05 | 4.77E-05 |
| . | . | . | 0 | . | . | . |
| . | . | 9.16E-05 | 0.0012 | 6.63E-05 | 0.0001 | 9.15E-05 |
| . | . | 0.0001 | 8.00E-04 | 0.0001 | 0.0001 | 0.0001 |
| . | . | 3.19E-05 | 5.00E-04 | 1.48E-05 | 5.21E-05 | 3.18E-05 |
| . | . | 8.78E-05 | 0.0014 | 5.91E-05 | 0.0001 | 8.75E-05 |
| . | . | 2.39E-05 | 1.00E-04 | 2.21E-05 | 2.60E-05 | 2.78E-05 |
| . | . | 0.002 | 0.0033 | 0.002 | 0.0019 | 0.002 |
| . | . | 0.0008 | 0.0021 | 0.0007 | 0.001 | 0.0008 |
| . | . | 4.01E-06 | 2.90E-05 | 7.39E-06 | 0 | 4.00E-06 |
| . | . | 4.78E-05 | 9.69E-05 | 5.89E-05 | 3.47E-05 | 4.77E-05 |
| . | . | 0.0007 | 0.0043 | 0.0006 | 0.0008 | 0.0007 |
| . | . | . | 0 | . | . | . |
| . | . | 0.0006 | 5.00E-04 | 0.0006 | 0.0007 | 0.0006 |
| . | . | 0.0001 | 6.00E-04 | 0.0001 | 0.0001 | 0.0001 |
| . | . | . | 0 | . | . | . |
| . | . | 0.002 | 0.0033 | 0.002 | 0.0019 | 0.002 |
| . | . | . | 0 | . | . | . |
| . | . | 0.0009 | 0.0013 | 0.0008 | 0.001 | 0.0009 |
| . | . | . | 0 | . | . | . |

| | | | | | | |
|---|---|---|---|---|---|---|
| . | . | 1.99E-05 | 1.00E-04 | 1.47E-05 | 2.60E-05 | 1.99E-05 |
| . | . | 1.99E-05 | 4.40E-05 | 7.36E-06 | 3.46E-05 | 1.99E-05 |
| . | . | 1.22E-05 | 5.81E-05 | 1.50E-05 | 8.89E-06 | 1.19E-05 |
| . | . | 8.05E-06 | 8.87E-06 | 1.48E-05 | 0 | 8.39E-05 |
| . | . | 0.0001 | 8.00E-04 | 0.0001 | 0.0001 | 0.0001 |
| . | . | 6.49E-06 | 1.68E-05 | 1.22E-05 | 0 | 6.23E-06 |
| . | . | 0.0002 | 4.00E-04 | 0.0002 | 0.0002 | 0.0002 |
| . | . | 2.39E-05 | 2.00E-04 | 7.37E-06 | 4.34E-05 | 2.78E-05 |
| . | . | 7.18E-05 | 1.00E-04 | 9.60E-05 | 4.34E-05 | 7.16E-05 |
| . | . | 0.0004 | 0.0011 | 0.0004 | 0.0004 | 0.0004 |
| . | . | 0.0001 | 0.0015 | 0.0002 | 0.0001 | 0.0001 |
| . | . | 0.0004 | 0.001 | 0.0003 | 0.0004 | 0.0004 |
| . | . | 1.19E-05 | 8.67E-05 | 1.47E-05 | 8.65E-06 | 1.19E-05 |
| . | . | 1.19E-05 | 8.67E-05 | 1.47E-05 | 8.65E-06 | 1.19E-05 |
| . | . | . | 0 | . | . | . |
| . | . | 4.08E-06 | 3.31E-05 | 7.55E-06 | 0 | 3.98E-06 |
| . | . | . | 0 | . | . | . |
| . | . | . | 0 | . | . | . |
| . | . | 3.98E-06 | 8.81E-06 | 0 | 8.66E-06 | 3.98E-06 |
| . | . | 3.99E-06 | 2.89E-05 | 7.38E-06 | 0 | 3.98E-06 |
| . | . | 4.77E-05 | 7.91E-05 | 6.62E-05 | 2.60E-05 | 4.77E-05 |
| . | . | 5.97E-05 | 8.00E-04 | 4.42E-05 | 7.80E-05 | 5.96E-05 |
| . | . | 0.001 | 0.0013 | 0.0011 | 0.001 | 0.001 |
| . | . | 6.30E-06 | 3.75E-05 | 1.14E-05 | 0 | 2.46E-05 |
| . | . | 7.95E-06 | 5.78E-05 | 7.36E-06 | 8.66E-06 | 7.95E-06 |
| . | . | 7.16E-05 | 2.00E-04 | 8.83E-05 | 5.19E-05 | 7.16E-05 |
| . | . | . | 0 | . | . | . |
| . | . | 0.0001 | 0.0015 | 0.0002 | 0.0001 | 0.0001 |
| . | . | 0.0001 | 0.0019 | 9.63E-05 | 0.0002 | 0.0001 |
| . | . | 0.0023 | 0.004 | 0.0023 | 0.0024 | 0.0023 |
| . | . | 0.002 | 0.0033 | 0.002 | 0.0019 | 0.002 |
| . | . | 0.0004 | 0.001 | 0.0003 | 0.0004 | 0.0004 |
| . | . | 0.002 | 0.0033 | 0.002 | 0.0019 | 0.002 |
| . | . | 0.0014 | 0.0021 | 0.0016 | 0.0012 | 0.0014 |
| . | . | . | 0 | . | . | . |
| . | . | 7.96E-06 | 6.15E-05 | 1.47E-05 | 0 | 7.95E-06 |
| . | . | 0.0003 | 5.00E-04 | 0.0003 | 0.0003 | 0.0003 |
| . | . | 0.001 | 0.0013 | 0.0011 | 0.001 | 0.001 |
| . | . | 0.0011 | 0.0017 | 0.0011 | 0.0012 | 0.0012 |
| . | . | 7.96E-06 | 8.81E-06 | 0 | 1.73E-05 | 1.59E-05 |
| . | . | 0.0004 | 0.0011 | 0.0004 | 0.0004 | 0.0004 |
| . | . | 0.0001 | 3.00E-04 | 9.57E-05 | 0.0001 | 0.0001 |
| . | . | 0.0007 | 0.0043 | 0.0006 | 0.0008 | 0.0007 |

| | | | | | | |
|---|---|---|---|---|---|---|
| . | . | 1.73E-05 | 6.83E-05 | 1.60E-05 | 1.88E-05 | 1.59E-05 |
| . | . | 0.002 | 0.0033 | 0.002 | 0.0019 | 0.002 |
| . | . | 0.001 | 0.0013 | 0.0011 | 0.001 | 0.001 |
| . | . | 0.0007 | 0.0043 | 0.0006 | 0.0008 | 0.0007 |
| . | . | 0.0001 | 8.00E-04 | 0.0001 | 0.0001 | 0.0001 |
| . | . | 0.0007 | 0.0043 | 0.0006 | 0.0008 | 0.0007 |
| . | . | . | 0 | . | . | . |
| . | . | . | 0 | . | . | . |
| . | . | 0.0005 | 0.0042 | 0.0006 | 0.0005 | 0.0002 |
| . | . | 0.0006 | 5.00E-04 | 0.0006 | 0.0007 | 0.0006 |
| . | . | 0.0003 | 5.00E-04 | 0.0003 | 0.0003 | 0.0003 |
| . | . | 0.0002 | 0.0033 | 0.0002 | 0.0003 | 0.0002 |
| . | . | 0.0004 | 0.0011 | 0.0004 | 0.0004 | 0.0004 |
| . | . | . | 0 | . | . | . |
| . | . | 4.78E-05 | 9.69E-05 | 5.89E-05 | 3.47E-05 | 4.77E-05 |
| . | . | . | 0 | . | . | . |
| . | . | 0.0002 | 0.002 | 0.0001 | 0.0002 | 0.0002 |
| . | . | 0.0026 | 0.0034 | 0.0026 | 0.0025 | 0.0011 |
| . | . | 5.97E-05 | 8.00E-04 | 3.68E-05 | 8.66E-05 | 5.96E-05 |
| . | . | 1.59E-05 | 2.00E-04 | 1.47E-05 | 1.73E-05 | 1.59E-05 |
| . | . | 0.0002 | 0.002 | 0.0001 | 0.0002 | 0.0002 |
| . | . | 0.0002 | 0.0031 | 0.0002 | 0.0002 | 0.0002 |
| . | . | 0.0002 | 0.0028 | 0.0002 | 0.0002 | 0.0003 |
| . | . | 7.99E-06 | 3.27E-05 | 7.39E-06 | 8.70E-06 | 7.95E-06 |
| . | . | 1.21E-05 | 2.00E-04 | 7.42E-06 | 1.75E-05 | 1.19E-05 |
| . | . | 0.0002 | 0.0022 | 0.0002 | 0.0001 | 0.0002 |
| . | . | 4.01E-06 | 5.47E-05 | 0 | 8.74E-06 | 3.98E-06 |
| . | . | . | 0 | . | . | . |
| . | . | 2.62E-05 | 7.52E-05 | 3.22E-05 | 1.91E-05 | 2.39E-05 |
| . | . | 7.97E-06 | 1.00E-04 | 0 | 1.74E-05 | 7.95E-06 |
| . | . | 1.87E-05 | 1.00E-04 | 8.58E-06 | 3.09E-05 | 1.59E-05 |
| . | . | 1.19E-05 | 2.00E-04 | 7.37E-06 | 1.73E-05 | 1.19E-05 |
| . | . | 8.09E-06 | 1.00E-04 | 7.46E-06 | 8.84E-06 | 7.99E-06 |
| . | . | 4.04E-06 | 5.45E-05 | 0 | 8.85E-06 | 3.98E-06 |
| . | . | 9.21E-05 | 0.0011 | 8.89E-05 | 9.58E-05 | 9.15E-05 |
| . | . | 7.98E-05 | 0.001 | 9.60E-05 | 6.08E-05 | 7.95E-05 |
| . | . | 1.19E-05 | 2.00E-04 | 7.37E-06 | 1.73E-05 | 1.19E-05 |
| . | . | . | 0 | . | . | . |
| . | . | 0.0001 | 6.00E-04 | 8.05E-05 | 0.0001 | 0.0001 |
| . | . | 0.0003 | 0.0042 | 0.0003 | 0.0003 | 0.0003 |
| . | . | 2.78E-05 | 4.00E-04 | 2.94E-05 | 2.60E-05 | 2.78E-05 |
| . | . | 0.0003 | 6.00E-04 | 0.0003 | 0.0003 | 0.0003 |
| . | . | 1.59E-05 | 2.00E-04 | 7.36E-06 | 2.60E-05 | 1.59E-05 |

| | | | | | | |
|---|---|---|---|---|---|---|
| . | . | 0.0002 | 0.0028 | 0.0002 | 0.0002 | 0.0003 |
| . | . | 0.0002 | 0.0022 | 0.0002 | 0.0001 | 0.0002 |
| . | . | 5.18E-05 | 2.00E-04 | 5.90E-05 | 4.34E-05 | 5.17E-05 |
| . | . | 4.38E-05 | 6.00E-04 | 3.69E-05 | 5.20E-05 | 4.37E-05 |
| . | . | 8.75E-05 | 4.00E-04 | 7.36E-05 | 0.0001 | 8.75E-05 |
| . | . | 3.98E-06 | 5.44E-05 | 7.36E-06 | 0 | 3.98E-06 |
| . | . | 1.99E-05 | 3.00E-04 | 7.36E-06 | 3.47E-05 | 1.99E-05 |
| . | . | 1.21E-05 | 2.00E-04 | 7.42E-06 | 1.75E-05 | 1.19E-05 |
| . | . | 9.61E-05 | 9.00E-04 | 7.22E-05 | 0.0001 | 7.62E-05 |
| . | . | . | 0 | . | . | . |
| . | . | 0.0001 | 0.0016 | 0.0001 | 0.0001 | 0.0001 |
| . | . | . | 0 | . | . | . |
| . | . | . | 0 | . | . | . |
| . | . | 0.0001 | 0.0015 | 0.0001 | 0.0001 | 0.0001 |
| . | . | 1.20E-05 | 2.00E-04 | 7.37E-06 | 1.74E-05 | 1.19E-05 |
| . | . | 5.57E-05 | 6.00E-04 | 3.68E-05 | 7.79E-05 | 5.57E-05 |
| . | . | 0.0003 | 0.004 | 0.0003 | 0.0003 | 0.0003 |
| . | . | 0.0005 | 0.0022 | 0.0004 | 0.0006 | 0.0005 |
| . | . | 0.0003 | 0.004 | 0.0002 | 0.0004 | 0.0003 |
| . | . | 0.0003 | 0.004 | 0.0003 | 0.0003 | 0.0003 |
| . | . | 9.53E-05 | 7.00E-04 | 0.0001 | 7.30E-05 | 9.19E-05 |
| . | . | 3.19E-05 | 4.00E-04 | 3.68E-05 | 2.60E-05 | 3.18E-05 |
| . | . | 0.0002 | 0.0031 | 0.0002 | 0.0002 | 0.0002 |
| . | . | 0.0003 | 0.004 | 0.0003 | 0.0003 | 0.0003 |
| . | . | 1.78E-05 | 1.00E-04 | 3.57E-05 | 0 | 4.24E-05 |
| . | . | 0.0002 | 0.0024 | 0.0002 | 0.0001 | 0.0002 |
| . | . | 0.0005 | 0.0022 | 0.0004 | 0.0006 | 0.0005 |
| . | . | 0.0005 | 0.0034 | 0.0005 | 0.0005 | 0.0005 |
| . | . | 0.0002 | 0.002 | 0.0001 | 0.0002 | 0.0002 |
| . | . | 0.0001 | 0.0016 | 0.0001 | 0.0001 | 0.0001 |
| . | . | . | 0 | . | . | . |
| . | . | 8.00E-06 | 1.78E-05 | 0 | 1.74E-05 | 7.95E-06 |
| . | . | 1.99E-05 | 3.00E-04 | 7.36E-06 | 3.47E-05 | 1.99E-05 |
| . | . | 0.0003 | 0.004 | 0.0003 | 0.0003 | 0.0003 |
| . | . | 0.0005 | 0.0034 | 0.0005 | 0.0005 | 0.0005 |
| . | . | 0.0001 | 6.00E-04 | 8.05E-05 | 0.0001 | 0.0001 |
| . | . | 2.21E-05 | 2.00E-04 | 2.66E-05 | 1.65E-05 | 2.82E-05 |
| . | . | 8.76E-05 | 5.00E-04 | 9.57E-05 | 7.80E-05 | 8.75E-05 |
| . | . | . | 0 | . | . | . |
| . | . | . | 0 | . | . | . |
| . | . | 0.0003 | 0.0033 | 0.0003 | 0.0003 | 0.0003 |
| . | . | 4.38E-05 | 6.00E-04 | 3.69E-05 | 5.20E-05 | 4.37E-05 |
| . | . | 0.0003 | 0.004 | 0.0003 | 0.0003 | 0.0003 |

| | | | | | | |
|---|---|---|---|---|---|---|
| . | . | 0.0002 | 0.0031 | 0.0003 | 0.0002 | 0.0002 |
| . | . | 0.0005 | 0.0034 | 0.0005 | 0.0005 | 0.0005 |
| 0.9001 | 0.48 | . | 0 | . | . | . |

| AF_afr | AF_sas | AF_amr | AF_eas | AF_nfe | AF_fin | AF_asj |
|---|---|---|---|---|---|---|
| . | . | . | . | . | . | . |
| . | . | . | . | . | . | . |
| 0.0009 | 0.0001 | 0.0021 | 0 | 0.004 | 0.0001 | 0.0026 |
| 0 | 0 | 0 | 0 | 2.66E-05 | 0 | 0 |
| 0.0002 | 0.0001 | 0.0001 | 0 | 0.0022 | 0.0068 | 0 |
| 0 | 0 | 0 | 5.44E-05 | 0 | 0 | 0 |
| 0 | 0 | 2.89E-05 | 0 | 7.03E-05 | 0 | 0 |
| 0 | 0 | 0 | 0 | 3.52E-05 | 0 | 0 |
| 0.0005 | 0.0003 | 0.0013 | 0 | 0.004 | 0.0003 | 0 |
| 6.15E-05 | 0.0002 | 8.67E-05 | 0 | 0.002 | 0.0014 | 0.0005 |
| 0.0002 | 3.27E-05 | 0.0004 | 0 | 0.002 | 0.0003 | 9.92E-05 |
| 0.0009 | 0.0001 | 0.0021 | 0 | 0.004 | 0.0001 | 0.0026 |
| . | . | . | . | . | . | . |
| 0.0001 | 0.0002 | 0.0013 | 0 | 0.0013 | 0.0019 | 0.0003 |
| . | . | . | . | . | . | . |
| 0.0002 | 0.0032 | 2.91E-05 | 0 | 0.0003 | 0 | 0 |
| . | . | . | . | . | . | . |
| 0 | 0 | 0 | 0 | 0 | 0 | 0 |
| 0.0002 | 3.27E-05 | 0.0004 | 0 | 0.002 | 0.0003 | 9.92E-05 |
| 0 | 0 | 0 | 0.0001 | 0.0002 | 0 | 0 |
| 0 | 0.0002 | 0 | 0 | 1.76E-05 | 0 | 9.93E-05 |
| . | . | . | . | . | . | . |
| 6.20E-05 | 0.0005 | 0 | 5.44E-05 | 1.80E-05 | 0 | 0 |
| 0 | 0 | 0 | 0 | 8.11E-05 | 0 | 0 |
| . | . | . | . | . | . | . |
| 0.0001 | 3.27E-05 | 0 | 0 | 8.85E-06 | 0 | 0 |
| 0 | 0 | 0 | 0 | 8.17E-05 | 0 | 0 |
| 6.21E-05 | 0 | 2.90E-05 | 5.44E-05 | 8.13E-05 | 0 | 0 |
| . | . | . | . | . | . | . |
| 0 | 0 | 0 | 5.57E-05 | 0.0004 | 4.64E-05 | 0 |
| 0.0002 | 3.27E-05 | 0.0004 | 0 | 0.002 | 0.0003 | 9.92E-05 |
| 6.16E-05 | 0 | 0.0001 | 0 | 0.0005 | 0.006 | 0 |
| 0 | 0 | 0 | 0 | 0 | 0.001 | 0 |
| 6.15E-05 | 0.0002 | 8.67E-05 | 0 | 0.002 | 0.0014 | 0.0005 |
| 0 | 0.0032 | 2.89E-05 | 5.44E-05 | 5.31E-05 | 0 | 0 |
| 0 | 0 | 0 | 0 | 8.84E-06 | 0 | 0 |
| 0 | 0 | 0 | 0 | 8.86E-06 | 0 | 0 |
| . | . | . | . | . | . | . |
| 0.0002 | 3.27E-05 | 2.90E-05 | 5.57E-05 | 0.0012 | 0.0092 | 0 |
| 0 | 0 | 5.79E-05 | 0 | 8.81E-06 | 0 | 0 |
| 0.0004 | 0.0005 | 2.89E-05 | 0 | 0.0005 | 0 | 9.93E-05 |
| 0 | 0 | 0 | 5.44E-05 | 1.76E-05 | 0 | 0 |

| | | | | | | |
|---|---|---|---|---|---|---|
| 0.0009 | 0.0001 | 0.0021 | 0 | 0.004 | 0.0001 | 0.0026 |
| . | . | . | . | . | . | . |
| 0.0001 | 0.0007 | 0.0017 | 0.0001 | 0.001 | 0 | 0.0063 |
| 0 | 0 | 0 | 0 | 8.80E-06 | 0 | 0 |
| 0 | 0.0002 | 0 | 0 | 1.76E-05 | 0 | 9.93E-05 |
| . | . | . | . | . | . | . |
| . | . | . | . | . | . | . |
| 0 | 0 | 0 | 0 | 8.24E-05 | 0 | 0 |
| 0 | 0 | 0 | 0 | 0.0004 | 0 | 0 |
| 6.15E-05 | 9.80E-05 | 0 | 5.44E-05 | 0.0001 | 4.62E-05 | 0 |
| 0.0001 | 3.27E-05 | 2.90E-05 | 0 | 0 | 0 | 0 |
| 0 | 0 | 0 | 0 | 0.0001 | 0.0002 | 9.92E-05 |
| 0 | 3.27E-05 | 0.0001 | 0 | 2.66E-05 | 0 | 0 |
| . | . | . | . | . | . | . |
| 0.0002 | 0 | 2.89E-05 | 0 | 0.0008 | 9.27E-05 | 0 |
| 0.0001 | 0 | 2.89E-05 | 0 | 3.52E-05 | 0 | 0 |
| 0 | 0 | 0 | 0 | 8.81E-06 | 0 | 0 |
| 0 | 0.0007 | 5.79E-05 | 0 | 0.0007 | 4.63E-05 | 0 |
| . | . | . | . | . | . | . |
| 0.0002 | 0.001 | 0.0021 | 0 | 0.0019 | 0.0002 | 0.0012 |
| 0.0002 | 0.0005 | 0 | 0 | 0.0006 | 0 | 0 |
| 0.001 | 0.0002 | 0.0004 | 0.0011 | 0.0003 | 0 | 0 |
| 0.0009 | 0.0001 | 0.0021 | 0 | 0.004 | 0.0001 | 0.0026 |
| 0.0002 | 0.0007 | 0.0003 | 0 | 0.0017 | 0.0008 | 9.93E-05 |
| . | . | . | . | . | . | . |
| 0.0002 | 3.27E-05 | 0.0004 | 0 | 0.002 | 0.0003 | 9.92E-05 |
| 0 | 0 | 0 | 0 | 4.43E-05 | 0 | 0 |
| 0 | 0 | 0.0005 | 0 | 2.64E-05 | 0 | 0 |
| 0.0006 | 0.0001 | 0.0017 | 0.0002 | 0.0033 | 0.0009 | 0.0016 |
| 0 | 0 | 0 | 0 | 7.09E-05 | 0 | 0 |
| 0 | 3.27E-05 | 0 | 0 | 0 | 0 | 0 |
| 6.18E-05 | 0 | 0 | 0 | 2.67E-05 | 0 | 0 |
| . | . | . | . | . | . | . |
| 0.0005 | 0.0003 | 0.0013 | 0 | 0.004 | 0.0003 | 0 |
| 0.0009 | 0.0001 | 0.0021 | 0 | 0.004 | 0.0001 | 0.0026 |
| 0.0009 | 0.0001 | 0.0021 | 0 | 0.004 | 0.0001 | 0.0026 |
| . | . | . | . | . | . | . |
| 0 | 0 | 5.80E-05 | 0 | 0.0004 | 4.77E-05 | 0 |
| 0 | 0 | 0 | 0 | 8.80E-06 | 0 | 0 |
| . | . | . | . | . | . | . |
| 6.16E-05 | 0 | 0 | 0 | 0.0003 | 9.24E-05 | 0 |
| 0.0002 | 0.0032 | 2.91E-05 | 0 | 0.0003 | 0 | 0 |
| 0.0001 | 0 | 0.0002 | 0 | 0.0001 | 0 | 0 |

| | | | | | | |
|---|---|---|---|---|---|---|
| 0 | 3.27E-05 | 5.79E-05 | 0 | 0.0001 | 0 | 0 |
| 0 | 0 | 0 | 0 | 0 | 0 | 0 |
| 0.0004 | 0 | 0.0013 | 0 | 0.0021 | 9.31E-05 | 0.0016 |
| . | . | . | . | . | . | . |
| 0 | 0 | 0 | 0 | 0.0004 | 0 | 0 |
| 0 | 0 | 0 | 0 | 1.76E-05 | 0 | 0 |
| . | . | . | . | . | . | . |
| 0.0009 | 0.0001 | 0.0021 | 0 | 0.004 | 0.0001 | 0.0026 |
| 0 | 0 | 2.90E-05 | 5.44E-05 | 2.64E-05 | 0 | 0 |
| . | . | . | . | . | . | . |
| 0 | 0 | 0 | 0 | 3.63E-05 | 0 | 0 |
| 0.0002 | 0.0001 | 0.0001 | 0 | 0.0022 | 0.0068 | 0 |
| 0 | 6.53E-05 | 8.67E-05 | 0 | 8.79E-06 | 0 | 0 |
| . | . | . | . | . | . | . |
| . | . | . | . | . | . | . |
| 0.0005 | 0.0003 | 0.0013 | 0 | 0.004 | 0.0003 | 0 |
| 6.19E-05 | 0 | 0 | 0 | 0 | 0.0004 | 0 |
| . | . | . | . | . | . | . |
| 0 | 0.0001 | 2.89E-05 | 0 | 0.0001 | 4.62E-05 | 0 |
| 0 | 9.81E-05 | 5.81E-05 | 0 | 8.26E-05 | 0 | 0.0004 |
| . | . | . | . | . | . | . |
| . | . | . | . | . | . | . |
| . | . | . | . | . | . | . |
| 0.0009 | 0.0001 | 0.0021 | 0 | 0.004 | 0.0001 | 0.0026 |
| 0.0002 | 3.27E-05 | 0.0004 | 0 | 0.002 | 0.0003 | 9.92E-05 |
| 0 | 0 | 0 | 0 | 0.0002 | 0 | 0 |
| 6.15E-05 | 0.0002 | 8.67E-05 | 0 | 0.002 | 0.0014 | 0.0005 |
| . | . | . | . | . | . | . |
| 0 | 0 | 0 | 0 | 0.0005 | 0.0002 | 0 |
| . | . | . | . | . | . | . |
| 0 | 6.57E-05 | 0.0001 | 5.55E-05 | 0.0004 | 0.0001 | 0 |
| 6.15E-05 | 0 | 0.0023 | 0 | 0.0003 | 0 | 0 |
| 0.0002 | 9.80E-05 | 2.89E-05 | 0 | 0.0012 | 0.0002 | 9.92E-05 |
| 0 | 0.0004 | 2.90E-05 | 0 | 4.43E-05 | 0 | 0 |
| 0.0002 | 0.001 | 0.0021 | 0 | 0.0019 | 0.0002 | 0.0012 |
| . | . | . | . | . | . | . |
| 0 | 3.27E-05 | 5.78E-05 | 0 | 8.80E-06 | 0 | 0 |
| 0.0002 | 0 | 2.89E-05 | 0 | 0 | 0 | 0 |
| 0 | 0 | 0 | 0 | 5.30E-05 | 0 | 0 |
| 0 | 0 | 0.0007 | 0 | 4.46E-05 | 0 | 0.0043 |
| 6.57E-05 | 0.0021 | 0.0002 | 0.0005 | 0.0005 | 0 | 0 |
| 0 | 6.53E-05 | 0 | 0 | 1.76E-05 | 0 | 0 |
| 0 | 6.57E-05 | 0.0001 | 5.55E-05 | 0.0004 | 0.0001 | 0 |

| | | | | | | |
|---|---|---|---|---|---|---|
| . | . | . | . | . | . | . |
| 0.0009 | 0.0001 | 0.0021 | 0 | 0.004 | 0.0001 | 0.0026 |
| 0.0002 | 0.001 | 0.0021 | 0 | 0.0019 | 0.0002 | 0.0012 |
| 0 | 0.0001 | 0 | 0 | 5.27E-05 | 0 | 9.92E-05 |
| 0 | 0 | 0 | 0 | 3.52E-05 | 0 | 0 |
| . | . | . | . | . | . | . |
| 0.0003 | 0.0002 | 0.0021 | 0 | 0.0009 | 0.0002 | 0 |
| 6.25E-05 | 6.55E-05 | 0 | 0 | 0.0002 | 0.0002 | 9.99E-05 |
| . | . | . | . | . | . | . |
| . | . | . | . | . | . | . |
| . | . | . | . | . | . | . |
| 0 | 3.27E-05 | 0 | 0 | 0 | 0 | 0 |
| 0 | 0 | 0 | 0.0016 | 8.79E-06 | 0 | 0 |
| 0 | 0 | 0 | 0 | 1.76E-05 | 0 | 0 |
| 0 | 0 | 0.0006 | 0 | 5.28E-05 | 0 | 0 |
| . | . | . | . | . | . | . |
| . | . | . | . | . | . | . |
| 0 | 0 | 0 | 0 | 2.67E-05 | 0 | 0 |
| 6.15E-05 | 0.0002 | 8.67E-05 | 0 | 0.002 | 0.0014 | 0.0005 |
| 0 | 6.53E-05 | 0 | 0 | 7.97E-05 | 0 | 0 |
| 0 | 0 | 0.0002 | 0 | 0 | 0 | 0 |
| 0.0003 | 0.0002 | 0.0021 | 0 | 0.0009 | 0.0002 | 0 |
| . | . | . | . | . | . | . |
| 0 | 0 | 5.78E-05 | 0 | 7.91E-05 | 0 | 0 |
| . | . | . | . | . | . | . |
| 0.0012 | 0 | 5.79E-05 | 5.44E-05 | 0 | 0 | 0 |
| 0 | 0 | 0.0008 | 0 | 0 | 0 | 0 |
| 0.0005 | 0 | 0 | 0 | 0 | 0 | 0 |
| 0.0014 | 0 | 0 | 0 | 0 | 0 | 0 |
| 0 | 0 | 0.0001 | 0 | 8.79E-06 | 0 | 0 |
| 0.0006 | 0.0001 | 0.0017 | 0.0002 | 0.0033 | 0.0009 | 0.0016 |
| 0.0003 | 0.0002 | 0.0021 | 0 | 0.0009 | 0.0002 | 0 |
| 0 | 0 | 2.90E-05 | 0 | 0 | 0 | 0 |
| 6.16E-05 | 0 | 0 | 0 | 9.69E-05 | 0 | 0 |
| 0.0043 | 0 | 0.0018 | 0 | 0.0002 | 0 | 9.93E-05 |
| . | . | . | . | . | . | . |
| 6.15E-05 | 9.80E-05 | 0.0005 | 0 | 0.0004 | 0 | 0.0081 |
| 0 | 0 | 0.0006 | 0 | 9.75E-05 | 0 | 9.99E-05 |
| . | . | . | . | . | . | . |
| 0.0006 | 0.0001 | 0.0017 | 0.0002 | 0.0033 | 0.0009 | 0.0016 |
| . | . | . | . | . | . | . |
| 0.0001 | 4.11E-05 | 0.0007 | 0 | 0.0013 | 0 | 0.0035 |
| . | . | . | . | . | . | . |

| | | | | | | |
|---|---|---|---|---|---|---|
| 0 | 0 | 0.0001 | 0 | 8.80E-06 | 0 | 0 |
| 0 | 0 | 0 | 0 | 4.40E-05 | 0 | 0 |
| 0 | 0 | 5.81E-05 | 5.49E-05 | 0 | 0 | 0 |
| 0 | 0 | 0 | 0 | 8.87E-06 | 0 | 9.96E-05 |
| 0 | 0 | 0.0008 | 0 | 0 | 0 | 0 |
| 0 | 0 | 0 | 0 | 1.68E-05 | 0 | 0 |
| 0 | 3.27E-05 | 5.78E-05 | 0 | 0.0004 | 0 | 0.0009 |
| 0 | 0 | 0.0002 | 0 | 0 | 0 | 0 |
| 0 | 6.53E-05 | 5.79E-05 | 0 | 0.0001 | 0 | 0 |
| 0.0009 | 0.0003 | 0.0011 | 5.53E-05 | 0.0002 | 0 | 0.0002 |
| 0 | 0 | 0.0001 | 0.0015 | 0 | 0 | 0 |
| 6.15E-05 | 6.53E-05 | 0.001 | 0 | 0.0005 | 0 | 0 |
| 0 | 0 | 8.67E-05 | 0 | 0 | 0 | 0 |
| 0 | 0 | 8.67E-05 | 0 | 0 | 0 | 0 |
| . | . | . | . | . | . | . |
| 0 | 3.31E-05 | 0 | 0 | 0 | 0 | 0 |
| . | . | . | . | . | . | . |
| . | . | . | . | . | . | . |
| 0 | 0 | 0 | 0 | 8.81E-06 | 0 | 0 |
| 0 | 0 | 2.89E-05 | 0 | 0 | 0 | 0 |
| 0 | 0 | 5.78E-05 | 0 | 7.91E-05 | 0 | 0 |
| 0.0008 | 0 | 0 | 0 | 0 | 0 | 0 |
| 0.0001 | 0.0002 | 0.0013 | 0 | 0.0013 | 0.0019 | 0.0003 |
| 0 | 0 | 3.75E-05 | 0 | 0 | 0 | 0 |
| 0 | 0 | 5.78E-05 | 0 | 0 | 0 | 0 |
| 0.0001 | 0.0002 | 5.78E-05 | 0.0002 | 3.52E-05 | 0 | 0 |
| . | . | . | . | . | . | . |
| 0 | 0 | 0.0001 | 0.0015 | 0 | 0 | 0 |
| 0.0019 | 0 | 5.79E-05 | 0 | 0 | 0 | 0 |
| 0.0009 | 0.0001 | 0.0021 | 0 | 0.004 | 0.0001 | 0.0026 |
| 0.0006 | 0.0001 | 0.0017 | 0.0002 | 0.0033 | 0.0009 | 0.0016 |
| 6.15E-05 | 6.53E-05 | 0.001 | 0 | 0.0005 | 0 | 0 |
| 0.0006 | 0.0001 | 0.0017 | 0.0002 | 0.0033 | 0.0009 | 0.0016 |
| 0.0002 | 0.001 | 0.0021 | 0 | 0.0019 | 0.0002 | 0.0012 |
| . | . | . | . | . | . | . |
| 6.15E-05 | 0 | 2.89E-05 | 0 | 0 | 0 | 0 |
| 0.0004 | 0.0005 | 2.89E-05 | 0 | 0.0005 | 0 | 9.93E-05 |
| 0.0001 | 0.0002 | 0.0013 | 0 | 0.0013 | 0.0019 | 0.0003 |
| 0.0001 | 0.0007 | 0.0017 | 0.0001 | 0.001 | 0 | 0.0063 |
| 0 | 0 | 0 | 0 | 8.81E-06 | 4.64E-05 | 0 |
| 0.0009 | 0.0003 | 0.0011 | 5.53E-05 | 0.0002 | 0 | 0.0002 |
| 0.0003 | 0 | 0.0003 | 0 | 7.03E-05 | 0 | 0 |
| 0.0043 | 0 | 0.0018 | 0 | 0.0002 | 0 | 9.93E-05 |

| | | | | | | |
|---|---|---|---|---|---|---|
| 6.83E-05 | 0 | 0 | 0 | 2.85E-05 | 0 | 0 |
| 0.0006 | 0.0001 | 0.0017 | 0.0002 | 0.0033 | 0.0009 | 0.0016 |
| 0.0001 | 0.0002 | 0.0013 | 0 | 0.0013 | 0.0019 | 0.0003 |
| 0.0043 | 0 | 0.0018 | 0 | 0.0002 | 0 | 9.93E-05 |
| 0 | 0 | 0.0008 | 0 | 0 | 0 | 0 |
| 0.0043 | 0 | 0.0018 | 0 | 0.0002 | 0 | 9.93E-05 |
| . | . | . | . | . | . | . |
| . | . | . | . | . | . | . |
| 0 | 0.0042 | 0 | 0 | 0.0001 | 0 | 0 |
| 6.15E-05 | 9.80E-05 | 0.0005 | 0 | 0.0004 | 0 | 0.0081 |
| 0.0004 | 0.0005 | 2.89E-05 | 0 | 0.0005 | 0 | 9.93E-05 |
| 0.0033 | 3.27E-05 | 0.0002 | 0 | 0 | 0 | 0 |
| 0.0009 | 0.0003 | 0.0011 | 5.53E-05 | 0.0002 | 0 | 0.0002 |
| . | . | . | . | . | . | . |
| 6.16E-05 | 0 | 0 | 0 | 9.69E-05 | 0 | 0 |
| . | . | . | . | . | . | . |
| 0 | 6.53E-05 | 0 | 0.002 | 8.80E-06 | 0 | 0 |
| 0.0013 | 0.0023 | 0.0014 | 0.0034 | 0.0016 | 0 | 0.019 |
| 0 | 0 | 0 | 0.0008 | 0 | 0 | 0 |
| 0.0002 | 0 | 0 | 0 | 8.79E-06 | 0 | 0 |
| 0 | 6.53E-05 | 0 | 0.002 | 8.80E-06 | 0 | 0 |
| 0 | 0 | 0 | 0.0031 | 0 | 0 | 0 |
| 0 | 3.27E-05 | 0 | 0.0028 | 0 | 0 | 0 |
| 0 | 3.27E-05 | 0 | 0 | 8.87E-06 | 0 | 0 |
| 0 | 0 | 0 | 0.0002 | 0 | 0 | 0 |
| 0 | 6.54E-05 | 0 | 0.0022 | 8.83E-06 | 0 | 0 |
| 0 | 0 | 0 | 5.47E-05 | 0 | 0 | 0 |
| . | . | . | . | . | . | . |
| 0 | 7.52E-05 | 0 | 0 | 2.88E-05 | 4.95E-05 | 0 |
| 0 | 0 | 0 | 0.0001 | 0 | 0 | 0 |
| 0 | 0 | 0.0001 | 0 | 0 | 0 | 0 |
| 0 | 0 | 0 | 0.0002 | 0 | 0 | 0 |
| 0 | 0 | 0 | 0.0001 | 0 | 0 | 0 |
| 0 | 0 | 0 | 5.45E-05 | 0 | 0 | 0 |
| 0 | 0 | 5.85E-05 | 0.0011 | 8.82E-06 | 0 | 0 |
| 0 | 3.27E-05 | 0 | 0.001 | 8.85E-06 | 0 | 0 |
| 0 | 0 | 0 | 0.0002 | 0 | 0 | 0 |
| . | . | . | . | . | . | . |
| 6.28E-05 | 0 | 3.25E-05 | 0.0006 | 9.47E-05 | 4.91E-05 | 0 |
| 0 | 3.29E-05 | 0 | 0.0042 | 0 | 0 | 0 |
| 0 | 0 | 0 | 0.0004 | 0 | 0 | 0 |
| 0.0002 | 3.37E-05 | 9.10E-05 | 0.0003 | 0.0006 | 5.00E-05 | 0 |
| 0 | 0 | 0 | 0.0002 | 0 | 0 | 0 |

| | | | | | | |
|---|---|---|---|---|---|---|
| 0 | 3.27E-05 | 0 | 0.0028 | 0 | 0 | 0 |
| 0 | 6.54E-05 | 0 | 0.0022 | 8.83E-06 | 0 | 0 |
| 0 | 0.0002 | 0 | 0.0001 | 3.53E-05 | 0 | 0 |
| 0 | 0 | 0 | 0.0006 | 0 | 0 | 0 |
| 0.0002 | 0 | 0 | 0.0004 | 7.04E-05 | 0 | 0 |
| 0 | 0 | 0 | 5.44E-05 | 0 | 0 | 0 |
| 0 | 0 | 0 | 0.0003 | 0 | 0 | 0 |
| 0 | 0 | 0 | 0.0002 | 0 | 0 | 0 |
| 0.0001 | 0 | 0 | 0.0009 | 4.16E-05 | 0 | 0 |
| . | . | . | . | . | . | . |
| 0 | 0 | 0 | 0.0016 | 8.79E-06 | 0 | 0 |
| . | . | . | . | . | . | . |
| . | . | . | . | . | . | . |
| 0.0015 | 0 | 0.0001 | 5.58E-05 | 1.83E-05 | 0 | 0 |
| 0 | 0 | 0 | 0.0002 | 0 | 0 | 0 |
| 0 | 9.80E-05 | 0 | 0.0006 | 0 | 0 | 0 |
| 0 | 0 | 0 | 0.004 | 0 | 0 | 0 |
| 0.0001 | 0.0007 | 0.0008 | 0.0022 | 0.0001 | 8.50E-05 | 0.0013 |
| 0 | 3.27E-05 | 2.90E-05 | 0.004 | 0 | 0 | 0 |
| 0 | 0 | 0 | 0.004 | 0 | 0 | 0 |
| 6.73E-05 | 3.34E-05 | 6.02E-05 | 0.0007 | 4.61E-05 | 0 | 0 |
| 0 | 0 | 0 | 0.0004 | 0 | 0 | 0 |
| 0 | 0 | 0 | 0.0031 | 0 | 0 | 0 |
| 0 | 0 | 0 | 0.004 | 0 | 0 | 0 |
| 0 | 0 | 0 | 0.0001 | 0 | 0 | 0 |
| 0 | 0 | 0 | 0.0024 | 0 | 0 | 0 |
| 0.0001 | 0.0007 | 0.0008 | 0.0022 | 0.0001 | 8.50E-05 | 0.0013 |
| 0 | 0.0008 | 0 | 0.0034 | 0.0003 | 0.0002 | 0 |
| 0 | 6.53E-05 | 0 | 0.002 | 8.80E-06 | 0 | 0 |
| 0 | 0 | 0 | 0.0016 | 0 | 0 | 0 |
| . | . | . | . | . | . | . |
| 0 | 0 | 0 | 0 | 1.78E-05 | 0 | 0 |
| 0 | 0 | 0 | 0.0003 | 0 | 0 | 0 |
| 0 | 0 | 0 | 0.004 | 0 | 0 | 0 |
| 0 | 0.0008 | 0 | 0.0034 | 0.0003 | 0.0002 | 0 |
| 6.28E-05 | 0 | 3.25E-05 | 0.0006 | 9.47E-05 | 4.91E-05 | 0 |
| 0 | 0 | 4.72E-05 | 0.0002 | 0 | 0 | 0 |
| 6.17E-05 | 0 | 2.89E-05 | 0.0005 | 7.92E-05 | 0 | 0.0002 |
| . | . | . | . | . | . | . |
| . | . | . | . | . | . | . |
| 0 | 0.0003 | 0.0002 | 0.0033 | 8.82E-06 | 0 | 0 |
| 0 | 0 | 0 | 0.0006 | 0 | 0 | 0 |
| 0 | 0 | 0 | 0.004 | 0 | 0 | 0 |

| | | | | | | |
|---|---|---|---|---|---|---|
| 6.15E-05 | 6.53E-05 | 0 | 0.0031 | 8.80E-06 | 0 | 0 |
| 0 | 0.0008 | 0 | 0.0034 | 0.0003 | 0.0002 | 0 |
| . | . | . | . | . | . | . |

| AF_oth | non_topmed_ | non_neuro_A | non_cancer_ | controls_AF_ | gnomad312_ | gnomad312_ |
|---|---|---|---|---|---|---|
| . | . | . | . | . | . | . |
| . | . | . | . | . | 0.0011 | 0.0011 |
| 0.0018 | 0.004 | 0.0042 | 0.004 | 0.0038 | 0.0022 | 0.0022 |
| 0 | 2.71E-05 | 2.23E-05 | 2.94E-05 | . | . | . |
| 0.0029 | 0.0022 | 0.0019 | 0.0023 | 0.0025 | 0.0012 | 0.0012 |
| 0 | 5.44E-05 | 7.46E-05 | 5.66E-05 | . | 6.59E-06 | 6.57E-06 |
| 0 | 7.16E-05 | 7.82E-05 | 6.81E-05 | 2.34E-05 | 2.63E-05 | 2.63E-05 |
| 0 | 3.59E-05 | 4.47E-05 | 3.90E-05 | 7.02E-05 | . | . |
| 0.0018 | 0.004 | 0.0042 | 0.0039 | 0.0038 | 0.0022 | 0.0022 |
| 0.001 | 0.002 | 0.0017 | 0.0018 | 0.0016 | 0.0013 | 0.0013 |
| 0.001 | 0.0021 | 0.0023 | 0.002 | 0.0018 | 0.0011 | 0.0011 |
| 0.0018 | 0.004 | 0.0042 | 0.004 | 0.0038 | 0.0022 | 0.0022 |
| . | . | . | . | . | 6.57E-06 | 6.57E-06 |
| 0.0012 | 0.0013 | 0.0013 | 0.0013 | 0.0014 | 0.0011 | 0.0011 |
| . | . | . | . | . | 1.97E-05 | 1.97E-05 |
| 0.0008 | 0.0032 | 0.0032 | 0.0032 | 0.0032 | 9.21E-05 | 9.20E-05 |
| . | . | . | . | . | . | . |
| 0 | . | . | . | . | . | . |
| 0.001 | 0.0021 | 0.0023 | 0.002 | 0.0018 | 0.0011 | 0.0011 |
| 0 | 0.0002 | 0.0001 | 0.0001 | 0.0003 | 0.0001 | 0.0001 |
| 0 | 0.0002 | 0.0002 | 0.0002 | 0.0001 | 3.29E-05 | 3.28E-05 |
| . | . | . | . | . | 1.97E-05 | 1.97E-05 |
| 0 | 0.0005 | 0.0005 | 0.0005 | 0.0005 | 5.26E-05 | 5.25E-05 |
| 0.0003 | 8.27E-05 | 8.04E-05 | 6.00E-05 | 4.73E-05 | 7.89E-05 | 7.88E-05 |
| . | . | . | . | . | . | . |
| 0 | 8.32E-05 | 0.0001 | 0.0001 | 0.0001 | 3.95E-05 | 3.94E-05 |
| 0 | 8.29E-05 | 0.0001 | 7.74E-05 | 0.0001 | 1.32E-05 | 1.31E-05 |
| 0 | 8.41E-05 | 9.18E-05 | 8.02E-05 | 9.37E-05 | 3.29E-05 | 3.28E-05 |
| . | . | . | . | . | . | . |
| 0.0003 | 0.0004 | 0.0004 | 0.0004 | 0.0002 | 0.0002 | 0.0002 |
| 0.001 | 0.0021 | 0.0023 | 0.002 | 0.0018 | 0.0011 | 0.0011 |
| 0.0002 | 0.0006 | 0.0005 | 0.0006 | 0.0005 | 0.0008 | 0.0008 |
| 0 | . | . | . | . | 9.21E-05 | 9.19E-05 |
| 0.001 | 0.002 | 0.0017 | 0.0018 | 0.0016 | 0.0013 | 0.0013 |
| 0 | 0.0032 | 0.0032 | 0.0031 | 0.0029 | 0.0002 | 0.0002 |
| 0 | 9.00E-06 | 1.12E-05 | 9.79E-06 | . | 6.57E-06 | 6.57E-06 |
| 0 | 9.03E-06 | 1.13E-05 | 9.82E-06 | . | 6.57E-06 | 6.57E-06 |
| . | . | . | . | . | . | . |
| 0.0015 | 0.0012 | 0.0011 | 0.0013 | 0.0015 | 0.001 | 0.001 |
| 0 | 5.81E-05 | 3.28E-05 | 5.85E-05 | 5.85E-05 | . | . |
| 0 | 0.0005 | 0.0005 | 0.0005 | 0.0007 | 0.0004 | 0.0004 |
| 0 | 5.44E-05 | 7.46E-05 | 5.65E-05 | 2.34E-05 | 3.29E-05 | 3.28E-05 |

| | | | | | | |
|---|---|---|---|---|---|---|
| 0.0018 | 0.004 | 0.0042 | 0.004 | 0.0038 | 0.0022 | 0.0022 |
| . | . | . | . | . | 6.57E-06 | 6.57E-06 |
| 0.0039 | 0.0017 | 0.0018 | 0.0017 | 0.002 | 0.0008 | 0.0008 |
| 0 | 8.96E-06 | . | 9.74E-06 | . | . | . |
| 0 | 0.0002 | 0.0002 | 0.0002 | 0.0001 | 3.29E-05 | 3.28E-05 |
| . | . | . | . | . | . | . |
| . | . | . | . | . | 6.57E-06 | 6.57E-06 |
| 0 | 7.46E-05 | 8.06E-05 | 8.08E-05 | 7.26E-05 | 7.98E-05 | 7.90E-05 |
| 0 | 0.0004 | 0.0004 | 0.0004 | 0.0003 | 0.0002 | 0.0002 |
| 0 | 0.0001 | 0.0001 | 0.0001 | 0.0002 | 7.89E-05 | 7.88E-05 |
| 0 | 0.0002 | 0.0001 | 0.0001 | . | 1.32E-05 | 1.31E-05 |
| 0 | 0.0001 | 0.0001 | 0.0001 | 0.0001 | 4.60E-05 | 4.60E-05 |
| 0 | 0.0001 | 0.0001 | 0.0001 | 0.0002 | 1.97E-05 | 1.97E-05 |
| . | . | . | . | . | 3.33E-05 | 3.29E-05 |
| 0.0005 | 0.0007 | 0.0008 | 0.0008 | 0.0006 | 0.0005 | 0.0005 |
| 0 | 2.90E-05 | 0.0001 | 0.0001 | 0.0001 | 3.94E-05 | 3.94E-05 |
| 0 | 8.97E-06 | 1.12E-05 | 9.75E-06 | 2.34E-05 | 6.58E-06 | 1.31E-05 |
| 0 | 0.0007 | 0.0008 | 0.0007 | 0.0008 | 0.0004 | 0.0004 |
| . | . | . | . | . | 6.57E-06 | 6.57E-06 |
| 0.0031 | 0.0021 | 0.0022 | 0.0021 | 0.0029 | 0.001 | 0.001 |
| 0 | 0.0005 | 0.0005 | 0.0005 | 0.0006 | 0.0002 | 0.0002 |
| 0.0007 | 0.0011 | 0.0011 | 0.0011 | 0.0014 | 0.0004 | 0.0004 |
| 0.0018 | 0.004 | 0.0042 | 0.004 | 0.0038 | 0.0022 | 0.0022 |
| 0.0005 | 0.0017 | 0.0017 | 0.0017 | 0.0016 | 0.0009 | 0.0009 |
| . | . | . | . | . | 6.57E-06 | 6.57E-06 |
| 0.001 | 0.0021 | 0.0023 | 0.002 | 0.0018 | 0.0011 | 0.0011 |
| 0 | 4.52E-05 | 4.49E-05 | 4.91E-05 | 2.35E-05 | . | . |
| 0.0007 | 0.0005 | 0.0005 | 0.0005 | 0.0002 | 1.31E-05 | 1.31E-05 |
| 0.0013 | 0.0033 | 0.0033 | 0.0033 | 0.004 | 0.002 | 0.002 |
| 0 | 7.22E-05 | 9.01E-05 | 7.85E-05 | 9.41E-05 | 2.64E-05 | 2.63E-05 |
| 0 | 3.27E-05 | 3.27E-05 | 3.28E-05 | . | . | . |
| 0 | 8.36E-05 | 6.19E-05 | 6.74E-05 | 0.0001 | 3.95E-05 | 3.94E-05 |
| . | . | . | . | . | 1.31E-05 | 1.31E-05 |
| 0.0018 | 0.004 | 0.0042 | 0.0039 | 0.0038 | 0.0022 | 0.0022 |
| 0.0018 | 0.004 | 0.0042 | 0.004 | 0.0038 | 0.0022 | 0.0022 |
| 0.0018 | 0.004 | 0.0042 | 0.004 | 0.0038 | 0.0022 | 0.0022 |
| . | . | . | . | . | . | . |
| 0.0002 | 0.0004 | 0.0004 | 0.0004 | 0.0003 | 0.0001 | 0.0001 |
| 0 | 8.97E-06 | 1.12E-05 | 9.75E-06 | 2.34E-05 | 6.57E-06 | 6.57E-06 |
| . | . | . | . | . | . | . |
| 0 | 0.0003 | 0.0003 | 0.0003 | 0.0003 | 9.20E-05 | 9.19E-05 |
| 0.0008 | 0.0032 | 0.0032 | 0.0032 | 0.0032 | 9.21E-05 | 9.20E-05 |
| 0 | 0.0002 | 0.0002 | 0.0002 | 0.0003 | 0.0001 | 0.0001 |

| | | | | | | |
|---|---|---|---|---|---|---|
| 0.0002 | 0.0001 | 0.0001 | 0.0001 | 0.0001 | 6.57E-05 | 6.57E-05 |
| 0 | . | . | . | . | 0.0002 | 0.0002 |
| 0.0018 | 0.0021 | 0.0022 | 0.0022 | 0.002 | 0.0015 | 0.0015 |
| . | . | . | . | . | . | . |
| 0 | 0.0004 | 0.0004 | 0.0004 | 0.0003 | 0.0002 | 0.0002 |
| 0 | 1.80E-05 | 2.24E-05 | 1.95E-05 | 2.34E-05 | 6.58E-06 | 6.57E-06 |
| . | . | . | . | . | 1.32E-05 | 1.31E-05 |
| 0.0018 | 0.004 | 0.0042 | 0.004 | 0.0038 | 0.0022 | 0.0022 |
| 0 | 5.45E-05 | 1.12E-05 | 5.66E-05 | 0.0001 | 1.31E-05 | 1.31E-05 |
| . | . | . | . | . | 6.57E-06 | 6.57E-06 |
| 0 | 3.70E-05 | 4.62E-05 | 1.99E-05 | 2.41E-05 | 1.32E-05 | 1.31E-05 |
| 0.0029 | 0.0022 | 0.0019 | 0.0023 | 0.0025 | 0.0012 | 0.0012 |
| 0.0002 | 8.71E-05 | 9.83E-05 | 8.76E-05 | 6.37E-05 | 1.97E-05 | 1.97E-05 |
| . | . | . | . | . | 1.32E-05 | 1.31E-05 |
| . | . | . | . | . | 0.0011 | 0.0011 |
| 0.0018 | 0.004 | 0.0042 | 0.0039 | 0.0038 | 0.0022 | 0.0022 |
| 0 | 8.37E-05 | 6.20E-05 | 6.75E-05 | 0.0001 | 0 | 6.57E-06 |
| . | . | . | . | . | . | . |
| 0 | 0.0001 | 0.0001 | 0.0001 | 7.02E-05 | 7.24E-05 | 7.23E-05 |
| 0.0002 | 9.81E-05 | 9.82E-05 | 9.84E-05 | 0.0002 | 5.26E-05 | 5.25E-05 |
| . | . | . | . | . | 6.60E-06 | 6.58E-06 |
| . | . | . | . | . | . | . |
| . | . | . | . | . | 2.66E-05 | 2.63E-05 |
| 0.0018 | 0.004 | 0.0042 | 0.004 | 0.0038 | 0.0022 | 0.0022 |
| 0.001 | 0.0021 | 0.0023 | 0.002 | 0.0018 | 0.0011 | 0.0011 |
| 0 | 0.0002 | 0.0002 | 0.0002 | 0.0003 | 7.23E-05 | 7.22E-05 |
| 0.001 | 0.002 | 0.0017 | 0.0018 | 0.0016 | 0.0013 | 0.0013 |
| . | . | . | . | . | . | . |
| 0 | 0.0005 | 0.0005 | 0.0005 | 0.0007 | 0.0003 | 0.0003 |
| . | . | . | . | . | . | . |
| 0.0002 | 0.0004 | 0.0004 | 0.0004 | 0.0004 | 0.0002 | 0.0002 |
| 0.0003 | 0.0023 | 0.0022 | 0.0023 | 0.0028 | 0.0003 | 0.0004 |
| 0 | 0.0012 | 0.001 | 0.0012 | 0.0013 | 0.0005 | 0.0005 |
| 0.0002 | 0.0004 | 0.0004 | 0.0004 | 0.0004 | 1.97E-05 | 1.97E-05 |
| 0.0031 | 0.0021 | 0.0022 | 0.0021 | 0.0029 | 0.001 | 0.001 |
| . | . | . | . | . | 1.97E-05 | 1.97E-05 |
| 0 | 5.81E-05 | 6.55E-05 | 5.84E-05 | 0.0001 | 6.57E-06 | 6.57E-06 |
| 0 | 0.0002 | 0.0002 | 0.0002 | 0.0003 | 3.29E-05 | 3.28E-05 |
| 0 | 5.40E-05 | 3.36E-05 | 5.86E-05 | 2.34E-05 | 1.97E-05 | 1.97E-05 |
| 0.0018 | 0.0006 | 0.0007 | 0.0007 | 0.0007 | 0.0001 | 0.0001 |
| 0.0008 | 0.0021 | 0.0021 | 0.0021 | 0.0021 | 0.0004 | 0.0004 |
| 0 | 6.53E-05 | 6.54E-05 | 6.55E-05 | 6.37E-05 | 6.57E-06 | 6.57E-06 |
| 0.0002 | 0.0004 | 0.0004 | 0.0004 | 0.0004 | 0.0002 | 0.0002 |

| | | | | | | |
|---|---|---|---|---|---|---|
| . | . | . | . | . | 1.97E-05 | 1.97E-05 |
| 0.0018 | 0.004 | 0.0042 | 0.004 | 0.0038 | 0.0022 | 0.0022 |
| 0.0031 | 0.0021 | 0.0022 | 0.0021 | 0.0029 | 0.001 | 0.001 |
| 0 | 0.0001 | 0.0001 | 0.0001 | 0.0001 | 1.97E-05 | 1.97E-05 |
| 0 | 2.69E-05 | 4.47E-05 | 3.90E-05 | 7.02E-05 | 3.29E-05 | 3.28E-05 |
| . | . | . | . | . | . | . |
| 0.0024 | 0.0021 | 0.0021 | 0.0021 | 0.0018 | 0.0006 | 0.0006 |
| 0.0002 | 0.0002 | 0.0002 | 0.0003 | 0.0003 | 0.0002 | 0.0002 |
| . | . | . | . | . | . | . |
| . | . | . | . | . | . | . |
| . | . | . | . | . | . | . |
| 0 | 3.27E-05 | 3.27E-05 | 3.28E-05 | 6.37E-05 | 1.32E-05 | 1.97E-05 |
| 0 | 0.0016 | 0.0016 | 0.0016 | 0.0017 | 3.29E-05 | 3.28E-05 |
| 0 | 1.79E-05 | 2.23E-05 | 1.95E-05 | 4.68E-05 | 6.57E-06 | 6.57E-06 |
| 0 | 0.0006 | 0.0006 | 0.0006 | 0.0006 | 7.89E-05 | 7.88E-05 |
| . | . | . | . | . | . | . |
| . | . | . | . | . | . | . |
| 0 | 2.73E-05 | 3.38E-05 | 1.98E-05 | 2.35E-05 | 1.97E-05 | 1.97E-05 |
| 0.001 | 0.002 | 0.0017 | 0.0018 | 0.0016 | 0.0013 | 0.0013 |
| 0.0003 | 8.12E-05 | 8.95E-05 | 6.86E-05 | 2.34E-05 | 1.97E-05 | 1.97E-05 |
| 0 | 0.0001 | 0.0002 | 0.0002 | 0.0002 | 6.58E-06 | 6.57E-06 |
| 0.0024 | 0.0021 | 0.0021 | 0.0021 | 0.0018 | 0.0006 | 0.0006 |
| . | . | . | . | . | 0.0001 | 0.0001 |
| 0.0002 | 7.16E-05 | 0.0001 | 7.79E-05 | 0.0001 | 1.97E-05 | 1.97E-05 |
| . | . | . | . | . | . | . |
| 0 | 0.001 | 0.0012 | 0.0013 | 0.0011 | 0.0004 | 0.0004 |
| 0 | 0.0008 | 0.0005 | 0.0008 | 0.0009 | 0.0004 | 0.0004 |
| 0 | 0.0005 | 0.0005 | 0.0005 | 0.0006 | 9.86E-05 | 9.85E-05 |
| 0 | 0.0012 | 0.0014 | 0.0015 | 0.0015 | 0.0003 | 0.0003 |
| 0.0002 | 0.0001 | 0.0001 | 0.0001 | 0.0001 | 1.31E-05 | 1.31E-05 |
| 0.0013 | 0.0033 | 0.0033 | 0.0033 | 0.004 | 0.002 | 0.002 |
| 0.0024 | 0.0021 | 0.0021 | 0.0021 | 0.0018 | 0.0006 | 0.0006 |
| 0 | 2.91E-05 | 3.28E-05 | 2.92E-05 | . | . | . |
| 0 | 9.87E-05 | 0.0001 | 7.80E-05 | 0.0001 | 9.86E-05 | 9.85E-05 |
| 0.002 | 0.0047 | 0.0043 | 0.0045 | 0.0046 | 0.0015 | 0.0015 |
| . | . | . | . | . | 6.57E-06 | 6.57E-06 |
| 0.0018 | 0.0005 | 0.0005 | 0.0005 | 0.0004 | 0.0005 | 0.0005 |
| 0 | 0.0006 | 0.0006 | 0.0006 | 0.0007 | 9.20E-05 | 9.20E-05 |
| . | . | . | . | . | 6.57E-06 | 6.57E-06 |
| 0.0013 | 0.0033 | 0.0033 | 0.0033 | 0.004 | 0.002 | 0.002 |
| . | . | . | . | . | 6.57E-06 | 6.57E-06 |
| 0.001 | 0.0013 | 0.0013 | 0.0013 | 0.0013 | 0.0009 | 0.0009 |
| . | . | . | . | . | 1.31E-05 | 1.31E-05 |

| | | | | | | |
|---|---|---|---|---|---|---|
| 0 | 0.0001 | 0.0001 | 0.0001 | 5.85E-05 | 6.57E-06 | 6.57E-06 |
| 0 | 4.48E-05 | 3.35E-05 | 3.89E-05 | . | 1.97E-05 | 1.97E-05 |
| 0 | 5.83E-05 | 6.59E-05 | 5.86E-05 | 0.0001 | 6.58E-06 | 6.57E-06 |
| 0 | 9.04E-06 | . | 9.82E-06 | . | . | . |
| 0 | 0.0008 | 0.0005 | 0.0008 | 0.0009 | 0.0004 | 0.0004 |
| 0 | 1.72E-05 | 2.04E-05 | 1.77E-05 | 5.18E-05 | 6.57E-06 | 6.57E-06 |
| 0.0003 | 0.0004 | 0.0004 | 0.0004 | 0.0004 | 0.0003 | 0.0003 |
| 0 | 0.0001 | 0.0002 | 0.0002 | 0.0002 | 6.58E-06 | 6.57E-06 |
| 0.0002 | 0.0001 | 0.0001 | 0.0001 | 6.38E-05 | 6.57E-05 | 6.57E-05 |
| 0.0007 | 0.0011 | 0.0011 | 0.0011 | 0.0012 | 0.0004 | 0.0004 |
| 0.0002 | 0.0015 | 0.0019 | 0.0015 | 0.0008 | 3.94E-05 | 3.94E-05 |
| 0.0003 | 0.0009 | 0.0008 | 0.001 | 0.0009 | 0.0006 | 0.0006 |
| 0 | 8.71E-05 | 9.83E-05 | 8.76E-05 | 0.0002 | 6.57E-06 | 6.57E-06 |
| 0 | 8.71E-05 | 9.83E-05 | 8.76E-05 | 0.0002 | 6.57E-06 | 6.57E-06 |
| . | . | . | . | . | 6.57E-06 | 6.57E-06 |
| 0 | 3.31E-05 | 3.31E-05 | 3.32E-05 | . | . | . |
| . | . | . | . | . | . | . |
| . | . | . | . | . | 6.57E-06 | 6.57E-06 |
| 0 | 8.98E-06 | . | 9.76E-06 | . | . | . |
| 0 | 2.90E-05 | 3.28E-05 | 2.92E-05 | . | . | . |
| 0.0002 | 7.16E-05 | 0.0001 | 7.79E-05 | 0.0001 | 1.97E-05 | 1.97E-05 |
| 0.0003 | 0.0007 | 0.0008 | 0.0009 | 0.0011 | 0.0004 | 0.0004 |
| 0.0012 | 0.0013 | 0.0013 | 0.0013 | 0.0014 | 0.0011 | 0.0011 |
| 0 | 3.76E-05 | 4.40E-05 | 3.76E-05 | 7.56E-05 | 1.33E-05 | 1.32E-05 |
| 0 | 5.81E-05 | 6.55E-05 | 5.84E-05 | 0.0001 | 1.31E-05 | 1.31E-05 |
| 0 | 0.0002 | 0.0002 | 0.0002 | 0.0003 | 6.57E-05 | 6.57E-05 |
| . | . | . | . | . | . | . |
| 0.0002 | 0.0015 | 0.0019 | 0.0015 | 0.0008 | 3.94E-05 | 3.94E-05 |
| 0 | 0.002 | 0.0019 | 0.0016 | 0.0018 | 0.0007 | 0.0007 |
| 0.0018 | 0.004 | 0.0042 | 0.004 | 0.0038 | 0.0022 | 0.0022 |
| 0.0013 | 0.0033 | 0.0033 | 0.0033 | 0.004 | 0.002 | 0.002 |
| 0.0003 | 0.0009 | 0.0008 | 0.001 | 0.0009 | 0.0006 | 0.0006 |
| 0.0013 | 0.0033 | 0.0033 | 0.0033 | 0.004 | 0.002 | 0.002 |
| 0.0031 | 0.0021 | 0.0022 | 0.0021 | 0.0029 | 0.001 | 0.001 |
| . | . | . | . | . | 0 | 6.57E-06 |
| 0 | 8.32E-05 | 6.17E-05 | 6.71E-05 | 5.85E-05 | . | . |
| 0 | 0.0005 | 0.0005 | 0.0005 | 0.0007 | 0.0004 | 0.0004 |
| 0.0012 | 0.0013 | 0.0013 | 0.0013 | 0.0014 | 0.0011 | 0.0011 |
| 0.0039 | 0.0017 | 0.0018 | 0.0017 | 0.002 | 0.0008 | 0.0008 |
| 0 | 8.97E-06 | 1.12E-05 | . | . | . | . |
| 0.0007 | 0.0011 | 0.0011 | 0.0011 | 0.0012 | 0.0004 | 0.0004 |
| 0.0003 | 0.0003 | 0.0003 | 0.0003 | 0.0005 | 0.0001 | 0.0001 |
| 0.002 | 0.0047 | 0.0043 | 0.0045 | 0.0046 | 0.0015 | 0.0015 |

| | | | | | | |
|---|---|---|---|---|---|---|
| 0 | 8.84E-05 | 6.85E-05 | 7.40E-05 | 7.52E-05 | 1.32E-05 | 1.31E-05 |
| 0.0013 | 0.0033 | 0.0033 | 0.0033 | 0.004 | 0.002 | 0.002 |
| 0.0012 | 0.0013 | 0.0013 | 0.0013 | 0.0014 | 0.0011 | 0.0011 |
| 0.002 | 0.0047 | 0.0043 | 0.0045 | 0.0046 | 0.0015 | 0.0015 |
| 0 | 0.0008 | 0.0005 | 0.0008 | 0.0009 | 0.0004 | 0.0004 |
| 0.002 | 0.0047 | 0.0043 | 0.0045 | 0.0046 | 0.0015 | 0.0015 |
| . | . | . | . | . | . | . |
| . | . | . | . | . | . | . |
| 0 | 0.0042 | 0.0042 | 0.0042 | 0.0046 | 0.0001 | 0.0001 |
| 0.0018 | 0.0005 | 0.0005 | 0.0005 | 0.0004 | 0.0005 | 0.0005 |
| 0 | 0.0005 | 0.0005 | 0.0005 | 0.0007 | 0.0004 | 0.0004 |
| 0.0002 | 0.0033 | 0.0033 | 0.0032 | 0.0032 | 0.0008 | 0.0008 |
| 0.0007 | 0.0011 | 0.0011 | 0.0011 | 0.0012 | 0.0004 | 0.0004 |
| . | . | . | . | . | . | . |
| 0 | 9.87E-05 | 0.0001 | 7.80E-05 | 0.0001 | 9.86E-05 | 9.85E-05 |
| . | . | . | . | . | . | . |
| 0.0002 | 0.002 | 0.0019 | 0.0019 | 0.0018 | 0.0002 | 0.0002 |
| 0.0027 | 0.0034 | 0.004 | 0.0035 | 0.0042 | 0.001 | 0.001 |
| 0 | 0.0008 | 0.0006 | 0.0008 | 0.001 | 6.57E-06 | 6.57E-06 |
| 0 | 8.32E-05 | 0.0002 | 0.0002 | 0.0003 | 6.57E-05 | 6.57E-05 |
| 0.0002 | 0.002 | 0.0019 | 0.0019 | 0.0018 | 0.0002 | 0.0002 |
| 0.0002 | 0.0031 | 0.003 | 0.0032 | 0.0033 | 0.0001 | 0.0001 |
| 0.0005 | 0.0028 | 0.0024 | 0.0029 | 0.0018 | 0.0002 | 0.0002 |
| 0 | 3.27E-05 | 3.27E-05 | 3.28E-05 | . | . | . |
| 0 | 0.0002 | 0.0001 | 0.0002 | . | 2.67E-05 | 2.65E-05 |
| 0.0002 | 0.0022 | 0.0022 | 0.0023 | 0.0023 | 3.28E-05 | 3.28E-05 |
| 0 | 5.47E-05 | . | 5.68E-05 | 0.0001 | . | . |
| . | . | . | . | . | . | . |
| 0 | 7.52E-05 | 7.52E-05 | 7.54E-05 | 7.76E-05 | 3.95E-05 | 5.26E-05 |
| 0 | 0.0001 | 0.0001 | 0.0001 | 0.0002 | . | . |
| 0 | 0.0001 | 0.0001 | 0.0001 | 6.32E-05 | 6.58E-06 | 6.57E-06 |
| 0 | 0.0002 | 7.45E-05 | 0.0002 | . | 6.57E-06 | 6.57E-06 |
| 0 | 0.0001 | 7.49E-05 | 0.0001 | 0.0001 | 6.58E-06 | 6.57E-06 |
| 0 | 5.45E-05 | . | 5.65E-05 | 0.0001 | . | . |
| 0 | 0.0011 | 0.0008 | 0.0011 | 0.0011 | 3.94E-05 | 3.94E-05 |
| 0 | 0.001 | 0.0008 | 0.0009 | 0.0007 | 9.20E-05 | 9.20E-05 |
| 0 | 0.0002 | 7.45E-05 | 0.0002 | . | 6.57E-06 | 6.57E-06 |
| . | . | . | . | . | . | . |
| 0.0002 | 0.0006 | 0.0006 | 0.0005 | 0.0003 | 0.0001 | 0.0001 |
| 0.0002 | 0.0042 | 0.0034 | 0.0043 | 0.0047 | 0.0002 | 0.0002 |
| 0 | 0.0004 | 0.0003 | 0.0004 | 0.0003 | 6.57E-06 | 6.57E-06 |
| 0.0002 | 0.0006 | 0.0006 | 0.0006 | 0.0005 | 0.0003 | 0.0003 |
| 0 | 0.0002 | 0.0002 | 0.0002 | 0.0001 | 6.57E-06 | 6.57E-06 |

| | | | | | | |
|---|---|---|---|---|---|---|
| 0.0005 | 0.0028 | 0.0024 | 0.0029 | 0.0018 | 0.0002 | 0.0002 |
| 0.0002 | 0.0022 | 0.0022 | 0.0023 | 0.0023 | 3.28E-05 | 3.28E-05 |
| 0 | 0.0002 | 0.0002 | 0.0002 | 0.0001 | 5.92E-05 | 5.91E-05 |
| 0 | 0.0006 | 0.0007 | 0.0005 | 0.0004 | 2.63E-05 | 2.63E-05 |
| 0.0003 | 0.0004 | 0.0006 | 0.0004 | 0.0004 | 7.90E-05 | 7.88E-05 |
| 0 | 5.44E-05 | 7.45E-05 | 5.65E-05 | 0.0001 | . | . |
| 0 | 0.0003 | 0.0002 | 0.0003 | 0.0002 | 3.29E-05 | 3.28E-05 |
| 0 | 0.0002 | 0.0001 | 0.0002 | . | 2.67E-05 | 2.65E-05 |
| 0 | 0.0009 | 0.0011 | 0.001 | 0.001 | 2.64E-05 | 2.63E-05 |
| . | . | . | . | . | . | . |
| 0 | 0.0016 | 0.0016 | 0.0016 | 0.0017 | 3.29E-05 | 3.28E-05 |
| . | . | . | . | . | . | . |
| . | . | . | . | . | . | . |
| 0 | 0.0017 | 0.0015 | 0.0016 | 0.0023 | 0.0005 | 0.0005 |
| 0 | 0.0002 | 0.0001 | 0.0002 | 0.0001 | . | . |
| 0 | 0.0006 | 0.0004 | 0.0006 | 0.0007 | 1.31E-05 | 1.31E-05 |
| 0.0002 | 0.004 | 0.0029 | 0.0042 | 0.0039 | 0.0001 | 0.0001 |
| 0.0011 | 0.0022 | 0.0024 | 0.0022 | 0.0019 | 0.0004 | 0.0004 |
| 0.0002 | 0.004 | 0.0038 | 0.004 | 0.0032 | 0.0001 | 0.0001 |
| 0.0002 | 0.004 | 0.0029 | 0.0042 | 0.0039 | 0.0001 | 0.0001 |
| 0.0002 | 0.0007 | 0.0009 | 0.0007 | 0.0008 | 5.31E-05 | 5.27E-05 |
| 0 | 0.0004 | 0.0006 | 0.0005 | 0.0004 | 1.32E-05 | 1.31E-05 |
| 0.0002 | 0.0031 | 0.003 | 0.0032 | 0.0033 | 0.0001 | 0.0001 |
| 0.0002 | 0.004 | 0.0029 | 0.0042 | 0.0039 | 0.0001 | 0.0001 |
| 0 | 0.0001 | 0.0001 | 0.0001 | 0.0003 | 1.97E-05 | 1.97E-05 |
| 0 | 0.0024 | 0.0017 | 0.0024 | 0.0022 | 5.93E-05 | 5.91E-05 |
| 0.0011 | 0.0022 | 0.0024 | 0.0022 | 0.0019 | 0.0004 | 0.0004 |
| 0 | 0.0034 | 0.003 | 0.0034 | 0.0031 | 0.0004 | 0.0004 |
| 0.0002 | 0.002 | 0.0019 | 0.0019 | 0.0018 | 0.0002 | 0.0002 |
| 0 | 0.0016 | 0.0014 | 0.0015 | 0.0016 | 9.86E-05 | 9.85E-05 |
| . | . | . | . | . | . | . |
| 0 | 1.81E-05 | 2.26E-05 | 9.86E-06 | . | . | . |
| 0 | 0.0003 | 0.0002 | 0.0003 | 0.0002 | 3.29E-05 | 3.28E-05 |
| 0.0002 | 0.004 | 0.0029 | 0.0042 | 0.0039 | 0.0001 | 0.0001 |
| 0 | 0.0034 | 0.003 | 0.0034 | 0.0031 | 0.0004 | 0.0004 |
| 0.0002 | 0.0006 | 0.0006 | 0.0005 | 0.0003 | 0.0001 | 0.0001 |
| 0 | 0.0002 | 0.0002 | 0.0002 | 0.0002 | . | . |
| 0 | 0.0005 | 0.0007 | 0.0005 | 0.0008 | 8.55E-05 | 8.54E-05 |
| . | . | . | . | . | . | . |
| . | . | . | . | . | . | . |
| 0 | 0.0033 | 0.0031 | 0.0034 | 0.0035 | 0.0001 | 0.0001 |
| 0 | 0.0006 | 0.0007 | 0.0005 | 0.0004 | 2.63E-05 | 2.63E-05 |
| 0.0002 | 0.004 | 0.0029 | 0.0042 | 0.0039 | 0.0001 | 0.0001 |

| | | | | | | | |
|---|---|---|---|---|---|---|---|
| | 0 | 0.0031 | 0.0031 | 0.0031 | 0.004 | 0.0002 | 0.0002 |
| | 0 | 0.0034 | 0.003 | 0.0034 | 0.0031 | 0.0004 | 0.0004 |
| . | . | . | . | . | | 6.57E-06 | 6.57E-06 |

| gnomad312_ | gnomad312_ | gnomad312_ | gnomad312_ | gnomad312_ | gnomad312_ | gnomad312_ |
|---|---|---|---|---|---|---|
| . | . | 0 | . | . | . | . |
| 0.0012 | 0.001 | 0.0019 | 0.0017 | 0.0007 | 0 | 0.0003 |
| 0.0022 | 0.0022 | 0.0038 | 0.003 | 0.0007 | 0 | 0.0038 |
| . | . | 0 | . | . | . | . |
| 0.001 | 0.0015 | 0.0015 | 0.0013 | 0.0003 | 0 | 6.54E-05 |
| 0 | 1.35E-05 | 1.47E-05 | 0 | 0 | 0 | 0 |
| 5.14E-05 | 0 | 5.88E-05 | 1.97E-05 | 0 | 0 | 0 |
| . | . | 0 | . | . | . | . |
| 0.0022 | 0.0022 | 0.0041 | 0.0037 | 0.0007 | 0.0011 | 0.0012 |
| 0.0012 | 0.0014 | 0.0024 | 0.0021 | 7.24E-05 | 0 | 0.0002 |
| 0.0013 | 0.0009 | 0.002 | 0.0018 | 0.0003 | 0.0044 | 0.0003 |
| 0.0022 | 0.0022 | 0.0038 | 0.003 | 0.0007 | 0 | 0.0038 |
| 1.29E-05 | 0 | 1.47E-05 | 0 | 0 | 0 | 0 |
| 0.001 | 0.0012 | 0.0026 | 0.002 | 0.0002 | 0 | 0.0026 |
| 2.57E-05 | 1.35E-05 | 4.41E-05 | 1.17E-05 | 0 | 0 | 0 |
| 9.00E-05 | 9.43E-05 | 0.0021 | 0.0011 | 2.42E-05 | 0 | 0 |
| . | . | 0 | . | . | . | . |
| . | . | 0 | . | . | . | . |
| 0.0013 | 0.0009 | 0.002 | 0.0018 | 0.0003 | 0.0044 | 0.0003 |
| 0.0002 | 5.51E-05 | 1.00E-04 | 5.90E-05 | 2.49E-05 | 0.0067 | 0 |
| 5.14E-05 | 1.35E-05 | 5.88E-05 | 1.97E-05 | 0 | 0 | 0 |
| 2.57E-05 | 1.35E-05 | 2.94E-05 | 4.88E-06 | 2.41E-05 | 0 | 0 |
| 6.42E-05 | 4.04E-05 | 2.00E-04 | 0 | 4.83E-05 | 0 | 0 |
| 6.42E-05 | 9.42E-05 | 1.00E-04 | 7.91E-05 | 4.83E-05 | 0 | 0 |
| . | . | 0 | . | . | . | . |
| 3.86E-05 | 4.04E-05 | 1.00E-04 | 6.29E-05 | 0.0001 | 0 | 0 |
| 2.57E-05 | 0 | 2.94E-05 | 4.88E-06 | 0 | 0 | 0 |
| 3.85E-05 | 2.69E-05 | 4.82E-05 | 7.99E-06 | 4.82E-05 | 0 | 0 |
| . | . | 0 | . | . | . | . |
| 0.0001 | 0.0002 | 4.00E-04 | 0.0003 | 0 | 0 | 0 |
| 0.0013 | 0.0009 | 0.002 | 0.0018 | 0.0003 | 0.0044 | 0.0003 |
| 0.0004 | 0.0012 | 5.00E-04 | 0.0003 | 0.0001 | 0 | 6.54E-05 |
| 6.43E-05 | 0.0001 | 0 | . | 0 | 0 | 0 |
| 0.0012 | 0.0014 | 0.0024 | 0.0021 | 7.24E-05 | 0 | 0.0002 |
| 5.14E-05 | 0.0003 | 0.0044 | 0.0029 | 2.41E-05 | 0 | 0 |
| 1.29E-05 | 0 | 1.47E-05 | 0 | 0 | 0 | 0 |
| 1.28E-05 | 0 | 1.47E-05 | 0 | 0 | 0 | 0 |
| . | . | 0 | . | . | . | . |
| 0.0009 | 0.0011 | 0.001 | 0.0008 | 0.0002 | 0 | 0.0003 |
| . | . | 0 | . | . | . | . |
| 0.0004 | 0.0004 | 0.0023 | 0.0013 | 0.0002 | 0 | 6.55E-05 |
| 5.14E-05 | 1.35E-05 | 4.83E-05 | 8.00E-06 | 4.83E-05 | 0 | 0 |

| | | | | | | |
|---|---|---|---|---|---|---|
| 0.0022 | 0.0022 | 0.0038 | 0.003 | 0.0007 | 0 | 0.0038 |
| 0 | 1.35E-05 | 1.47E-05 | 0 | 0 | 0 | 0 |
| 0.0008 | 0.0008 | 0.001 | 0.0004 | 0.0005 | 0 | 0.0007 |
| . | . | 0 | . | . | . | . |
| 5.14E-05 | 1.35E-05 | 5.88E-05 | 1.97E-05 | 0 | 0 | 0 |
| . | . | 0 | . | . | . | . |
| 1.28E-05 | 0 | 1.47E-05 | 0 | 0 | 0 | 0 |
| 6.48E-05 | 9.57E-05 | 1.00E-04 | 7.95E-05 | 4.91E-05 | 0 | 0 |
| 0.0003 | 6.73E-05 | 4.00E-04 | 0.0003 | 0 | 0 | 0 |
| 0.0001 | 5.38E-05 | 1.00E-04 | 6.81E-05 | 4.83E-05 | 0 | 6.55E-05 |
| 0 | 2.69E-05 | 2.00E-04 | 0 | 0 | 0 | 0 |
| 5.14E-05 | 4.03E-05 | 8.82E-05 | 3.76E-05 | 0 | 0 | 0 |
| 1.29E-05 | 2.69E-05 | 6.56E-05 | 0 | 0 | 0 | 6.56E-05 |
| 3.90E-05 | 2.73E-05 | 7.43E-05 | 2.87E-05 | 0 | 0 | 0 |
| 0.0006 | 0.0004 | 9.00E-04 | 0.0008 | 0.0002 | 0 | 0 |
| 6.43E-05 | 1.35E-05 | 4.00E-04 | 6.84E-05 | 7.24E-05 | 0 | 0 |
| 0 | 1.35E-05 | 1.47E-05 | 0 | 0 | 0 | 0 |
| 0.0004 | 0.0004 | 8.00E-04 | 0.0007 | 2.41E-05 | 0 | 6.55E-05 |
| 1.29E-05 | 0 | 1.47E-05 | 0 | 0 | 0 | 0 |
| 0.001 | 0.0009 | 0.0023 | 0.0017 | 0.0003 | 0 | 0.0023 |
| 0.0003 | 0.0002 | 0.001 | 0.0004 | 4.83E-05 | 0 | 0 |
| 0.0004 | 0.0004 | 0.0016 | 0.0008 | 0.0006 | 0 | 0.0002 |
| 0.0022 | 0.0022 | 0.0038 | 0.003 | 0.0007 | 0 | 0.0038 |
| 0.0009 | 0.0008 | 0.0015 | 0.0013 | 0.0002 | 0 | 0.0004 |
| 0 | 1.35E-05 | 0 | . | 0 | 0 | 0 |
| 0.0013 | 0.0009 | 0.002 | 0.0018 | 0.0003 | 0.0044 | 0.0003 |
| . | . | 0 | . | . | . | . |
| 2.57E-05 | 0 | 6.54E-05 | 0 | 2.41E-05 | 0 | 6.54E-05 |
| 0.002 | 0.002 | 0.0033 | 0.0029 | 0.0004 | 0 | 0.0028 |
| 3.86E-05 | 1.35E-05 | 5.88E-05 | 1.97E-05 | 0 | 0 | 0 |
| . | . | 0 | . | . | . | . |
| 5.14E-05 | 2.70E-05 | 7.35E-05 | 2.85E-05 | 2.42E-05 | 0 | 0 |
| 1.29E-05 | 1.35E-05 | 4.82E-05 | 7.99E-06 | 4.82E-05 | 0 | 0 |
| 0.0022 | 0.0022 | 0.0041 | 0.0037 | 0.0007 | 0.0011 | 0.0012 |
| 0.0022 | 0.0022 | 0.0038 | 0.003 | 0.0007 | 0 | 0.0038 |
| 0.0022 | 0.0022 | 0.0038 | 0.003 | 0.0007 | 0 | 0.0038 |
| . | . | 0 | . | . | . | . |
| 0.0002 | 0.0001 | 2.00E-04 | 0.0002 | 4.83E-05 | 0 | 0.0001 |
| 1.29E-05 | 0 | 1.47E-05 | 0 | 0 | 0 | 0 |
| . | . | 0 | . | . | . | . |
| 0.0001 | 8.07E-05 | 2.00E-04 | 9.05E-05 | 2.41E-05 | 0 | 0 |
| 9.00E-05 | 9.43E-05 | 0.0021 | 0.0011 | 2.42E-05 | 0 | 0 |
| 0.0002 | 8.07E-05 | 2.00E-04 | 0.0001 | 9.65E-05 | 0 | 6.55E-05 |

| | | | | | | |
|---|---|---|---|---|---|---|
| 3.85E-05 | 9.41E-05 | 2.00E-04 | 0 | 2.41E-05 | 0 | 0 |
| 0.0003 | 0.0001 | 7.00E-04 | 0.0005 | 0.0007 | 0 | 6.60E-05 |
| 0.0017 | 0.0013 | 0.0024 | 0.0021 | 0.0005 | 0 | 0.0022 |
| . | . | 0 | . | . | . | . |
| 0.0003 | 6.73E-05 | 4.00E-04 | 0.0003 | 0 | 0 | 0 |
| 0 | 1.35E-05 | 6.55E-05 | 0 | 0 | 0 | 6.55E-05 |
| 1.29E-05 | 1.35E-05 | 0 | . | 0 | 0 | 0 |
| 0.0022 | 0.0022 | 0.0038 | 0.003 | 0.0007 | 0 | 0.0038 |
| 2.57E-05 | 0 | 2.94E-05 | 4.88E-06 | 0 | 0 | 0 |
| 0 | 1.35E-05 | 1.47E-05 | 0 | 0 | 0 | 0 |
| 2.57E-05 | 0 | 2.94E-05 | 4.88E-06 | 0 | 0 | 0 |
| 0.001 | 0.0015 | 0.0015 | 0.0013 | 0.0003 | 0 | 6.54E-05 |
| 1.28E-05 | 2.69E-05 | 4.41E-05 | 1.17E-05 | 0 | 0 | 0 |
| 0 | 2.69E-05 | 4.00E-04 | 6.84E-05 | 0 | 0 | 0 |
| 0.0012 | 0.001 | 0.0019 | 0.0017 | 0.0007 | 0 | 0.0003 |
| 0.0022 | 0.0022 | 0.0041 | 0.0037 | 0.0007 | 0.0011 | 0.0012 |
| 0 | 0 | 0 | . | 0 | 0 | 0 |
| . | . | 0 | . | . | . | . |
| 7.72E-05 | 6.73E-05 | 2.00E-04 | 9.05E-05 | 0 | 0 | 0 |
| 7.71E-05 | 2.69E-05 | 1.00E-04 | 2.26E-05 | 0 | 0 | 0.0001 |
| 1.29E-05 | 0 | 1.47E-05 | 0 | 0 | 0 | 0 |
| . | . | 0 | . | . | . | . |
| 0 | 5.45E-05 | 4.00E-04 | 7.28E-05 | 2.42E-05 | 0 | 0 |
| 0.0022 | 0.0022 | 0.0038 | 0.003 | 0.0007 | 0 | 0.0038 |
| 0.0013 | 0.0009 | 0.002 | 0.0018 | 0.0003 | 0.0044 | 0.0003 |
| 6.42E-05 | 8.07E-05 | 2.00E-04 | 9.05E-05 | 0 | 0 | 0 |
| 0.0012 | 0.0014 | 0.0024 | 0.0021 | 7.24E-05 | 0 | 0.0002 |
| . | . | 0 | . | . | . | . |
| 0.0002 | 0.0003 | 6.00E-04 | 0.0004 | 4.83E-05 | 0 | 0 |
| . | . | 0 | . | . | . | . |
| 0.0002 | 0.0001 | 2.00E-04 | 0.0001 | 0.0001 | 0 | 6.54E-05 |
| 0.0003 | 0.0004 | 0.0022 | 0.0016 | 0.0001 | 0 | 0.0022 |
| 0.0006 | 0.0004 | 0.001 | 0.0008 | 0.0001 | 0 | 0.0001 |
| 1.28E-05 | 2.69E-05 | 4.00E-04 | 7.28E-05 | 0 | 0 | 0 |
| 0.001 | 0.0009 | 0.0023 | 0.0017 | 0.0003 | 0 | 0.0023 |
| 2.57E-05 | 1.35E-05 | 4.41E-05 | 1.17E-05 | 0 | 0 | 0 |
| 1.28E-05 | 0 | 2.41E-05 | 0 | 2.41E-05 | 0 | 0 |
| 3.86E-05 | 2.69E-05 | 2.00E-04 | 0 | 9.67E-05 | 0 | 0 |
| 1.29E-05 | 2.69E-05 | 4.41E-05 | 1.17E-05 | 0 | 0 | 0 |
| 0.0002 | 0.0001 | 3.00E-04 | 8.88E-05 | 0 | 0 | 0.0003 |
| 0.0004 | 0.0004 | 0.0023 | 0.0013 | 0.0001 | 0 | 0.0003 |
| 1.29E-05 | 0 | 1.47E-05 | 0 | 0 | 0 | 0 |
| 0.0002 | 0.0001 | 2.00E-04 | 0.0001 | 0.0001 | 0 | 6.54E-05 |

| | | | | | | |
|---|---|---|---|---|---|---|
| 0 | 4.03E-05 | 2.94E-05 | 4.88E-06 | 2.41E-05 | 0 | 0 |
| 0.0022 | 0.0022 | 0.0038 | 0.003 | 0.0007 | 0 | 0.0038 |
| 0.001 | 0.0009 | 0.0023 | 0.0017 | 0.0003 | 0 | 0.0023 |
| 1.28E-05 | 2.69E-05 | 4.41E-05 | 1.17E-05 | 0 | 0 | 0 |
| 3.86E-05 | 2.69E-05 | 7.35E-05 | 2.85E-05 | 0 | 0 | 0 |
| . | . | 0 | . | . | . | . |
| 0.0006 | 0.0006 | 0.0018 | 0.0013 | 0.0002 | 0 | 0.0018 |
| 0.0002 | 0.0003 | 3.00E-04 | 0.0002 | 4.83E-05 | 0.0077 | 0 |
| . | . | 0 | . | . | . | . |
| . | . | 0 | . | . | . | . |
| . | . | 0 | . | . | . | . |
| 2.57E-05 | 0 | 2.41E-05 | 0 | 2.41E-05 | 0 | 0 |
| 1.29E-05 | 5.38E-05 | 0.001 | 0.0004 | 0 | 0 | 0 |
| 0 | 1.35E-05 | 1.47E-05 | 0 | 0 | 0 | 0 |
| 0.0001 | 5.38E-05 | 3.00E-04 | 8.88E-05 | 4.83E-05 | 0 | 0.0003 |
| . | . | 0 | . | . | . | . |
| . | . | 0 | . | . | . | . |
| 2.57E-05 | 1.35E-05 | 4.41E-05 | 1.17E-05 | 0 | 0 | 0 |
| 0.0012 | 0.0014 | 0.0024 | 0.0021 | 7.24E-05 | 0 | 0.0002 |
| 1.29E-05 | 2.69E-05 | 2.94E-05 | 4.88E-06 | 0 | 0 | 0 |
| 1.29E-05 | 0 | 2.41E-05 | 0 | 2.41E-05 | 0 | 0 |
| 0.0006 | 0.0006 | 0.0018 | 0.0013 | 0.0002 | 0 | 0.0018 |
| 0.0002 | 8.19E-05 | 3.00E-04 | 9.06E-05 | 0 | 0 | 0.0003 |
| 3.85E-05 | 0 | 4.41E-05 | 1.17E-05 | 0 | 0 | 0 |
| . | . | 0 | . | . | . | . |
| 0.0005 | 0.0003 | 0.0015 | 0.0012 | 0.0015 | 0 | 0.0001 |
| 0.0002 | 0.0006 | 0.0042 | 0.0034 | 2.42E-05 | 0 | 0.0042 |
| 8.99E-05 | 0.0001 | 3.00E-04 | 0.0002 | 0.0003 | 0 | 6.55E-05 |
| 0.0004 | 0.0003 | 0.0013 | 0.001 | 0.0013 | 0 | 0 |
| 1.29E-05 | 1.35E-05 | 2.94E-05 | 4.88E-06 | 0 | 0 | 0 |
| 0.002 | 0.002 | 0.0033 | 0.0029 | 0.0004 | 0 | 0.0028 |
| 0.0006 | 0.0006 | 0.0018 | 0.0013 | 0.0002 | 0 | 0.0018 |
| . | . | 0 | . | . | . | . |
| 0.0001 | 5.38E-05 | 2.00E-04 | 9.05E-05 | 9.65E-05 | 0 | 0 |
| 0.0015 | 0.0015 | 0.0047 | 0.0041 | 0.0047 | 0 | 0.0007 |
| 1.29E-05 | 0 | 6.55E-05 | 0 | 0 | 0 | 6.55E-05 |
| 0.0005 | 0.0005 | 8.00E-04 | 0.0005 | 4.83E-05 | 0 | 0.0008 |
| 0.0001 | 8.07E-05 | 7.00E-04 | 0.0004 | 0 | 0 | 0.0007 |
| 1.29E-05 | 0 | 6.54E-05 | 0 | 0 | 0 | 6.54E-05 |
| 0.002 | 0.002 | 0.0033 | 0.0029 | 0.0004 | 0 | 0.0028 |
| 0 | 1.35E-05 | 6.55E-05 | 0 | 0 | 0 | 6.55E-05 |
| 0.001 | 0.0007 | 0.0013 | 0.0011 | 0.0002 | 0 | 0.001 |
| 0 | 2.69E-05 | 1.00E-04 | 2.26E-05 | 0 | 0 | 0.0001 |

| | | | | | | |
|---|---|---|---|---|---|---|
| 0 | 1.35E-05 | 0 | . | 0 | 0 | 0 |
| 2.57E-05 | 1.35E-05 | 2.94E-05 | 4.88E-06 | 0 | 0 | 0 |
| 0 | 1.35E-05 | 6.56E-05 | 0 | 0 | 0 | 6.56E-05 |
| . | . | 0 | . | . | . | . |
| 0.0002 | 0.0006 | 0.0042 | 0.0034 | 2.42E-05 | 0 | 0.0042 |
| 1.29E-05 | 0 | 2.41E-05 | 0 | 2.41E-05 | 0 | 0 |
| 0.0004 | 0.0003 | 6.00E-04 | 0.0004 | 2.41E-05 | 0 | 0.0001 |
| 1.29E-05 | 0 | 2.41E-05 | 0 | 2.41E-05 | 0 | 0 |
| 2.57E-05 | 0.0001 | 2.00E-04 | 0 | 4.83E-05 | 0 | 6.55E-05 |
| 0.0005 | 0.0004 | 0.0011 | 0.0008 | 0.0011 | 0 | 0.0006 |
| 3.86E-05 | 4.04E-05 | 0.001 | 0.0004 | 0 | 0 | 6.55E-05 |
| 0.0004 | 0.0007 | 0.0032 | 0.0025 | 9.65E-05 | 0 | 0.0032 |
| 0 | 1.35E-05 | 6.54E-05 | 0 | 0 | 0 | 6.54E-05 |
| 0 | 1.35E-05 | 6.54E-05 | 0 | 0 | 0 | 6.54E-05 |
| 1.29E-05 | 0 | 6.54E-05 | 0 | 0 | 0 | 6.54E-05 |
| . | . | 0 | . | . | . | . |
| . | . | 0 | . | . | . | . |
| 0 | 1.35E-05 | 2.00E-04 | 0 | 0 | 0 | 0 |
| . | . | 0 | . | . | . | . |
| . | . | 0 | . | . | . | . |
| 3.85E-05 | 0 | 4.41E-05 | 1.17E-05 | 0 | 0 | 0 |
| 0.0004 | 0.0003 | 0.0012 | 0.0009 | 0.0012 | 0 | 0.0001 |
| 0.001 | 0.0012 | 0.0026 | 0.002 | 0.0002 | 0 | 0.0026 |
| 1.30E-05 | 1.36E-05 | 6.59E-05 | 0 | 0 | 0 | 6.59E-05 |
| 1.29E-05 | 1.35E-05 | 1.00E-04 | 2.26E-05 | 0 | 0 | 0.0001 |
| 8.99E-05 | 4.04E-05 | 2.00E-04 | 0 | 0.0001 | 0 | 0 |
| . | . | 0 | . | . | . | . |
| 3.86E-05 | 4.04E-05 | 0.001 | 0.0004 | 0 | 0 | 6.55E-05 |
| 0.0007 | 0.0007 | 0.0024 | 0.0021 | 0.0024 | 0 | 0 |
| 0.0022 | 0.0022 | 0.0038 | 0.003 | 0.0007 | 0 | 0.0038 |
| 0.002 | 0.002 | 0.0033 | 0.0029 | 0.0004 | 0 | 0.0028 |
| 0.0004 | 0.0007 | 0.0032 | 0.0025 | 9.65E-05 | 0 | 0.0032 |
| 0.002 | 0.002 | 0.0033 | 0.0029 | 0.0004 | 0 | 0.0028 |
| 0.001 | 0.0009 | 0.0023 | 0.0017 | 0.0003 | 0 | 0.0023 |
| 0 | 0 | 0 | . | 0 | 0 | 0 |
| . | . | 0 | . | . | . | . |
| 0.0004 | 0.0004 | 0.0023 | 0.0013 | 0.0002 | 0 | 6.55E-05 |
| 0.001 | 0.0012 | 0.0026 | 0.002 | 0.0002 | 0 | 0.0026 |
| 0.0008 | 0.0008 | 0.001 | 0.0004 | 0.0005 | 0 | 0.0007 |
| . | . | 0 | . | . | . | . |
| 0.0005 | 0.0004 | 0.0011 | 0.0008 | 0.0011 | 0 | 0.0006 |
| 0.0001 | 0.0002 | 3.00E-04 | 0.0001 | 0.0003 | 0 | 0.0003 |
| 0.0015 | 0.0015 | 0.0047 | 0.0041 | 0.0047 | 0 | 0.0007 |

| | | | | | | |
|---|---|---|---|---|---|---|
| 2.57E-05 | 0 | 4.83E-05 | 8.00E-06 | 4.83E-05 | 0 | 0 |
| 0.002 | 0.002 | 0.0033 | 0.0029 | 0.0004 | 0 | 0.0028 |
| 0.001 | 0.0012 | 0.0026 | 0.002 | 0.0002 | 0 | 0.0026 |
| 0.0015 | 0.0015 | 0.0047 | 0.0041 | 0.0047 | 0 | 0.0007 |
| 0.0002 | 0.0006 | 0.0042 | 0.0034 | 2.42E-05 | 0 | 0.0042 |
| 0.0015 | 0.0015 | 0.0047 | 0.0041 | 0.0047 | 0 | 0.0007 |
| . | . | 0 | . | . | . | . |
| . | . | 0 | . | . | . | . |
| 0.0001 | 0.0002 | 0.0026 | 0.0015 | 0.0001 | 0 | 0 |
| 0.0005 | 0.0005 | 8.00E-04 | 0.0005 | 4.83E-05 | 0 | 0.0008 |
| 0.0004 | 0.0004 | 0.0023 | 0.0013 | 0.0002 | 0 | 6.55E-05 |
| 0.0008 | 0.0008 | 0.0029 | 0.0025 | 0.0029 | 0 | 0.0003 |
| 0.0005 | 0.0004 | 0.0011 | 0.0008 | 0.0011 | 0 | 0.0006 |
| . | . | 0 | . | . | . | . |
| 0.0001 | 5.38E-05 | 2.00E-04 | 9.05E-05 | 9.65E-05 | 0 | 0 |
| . | . | 0 | . | . | . | . |
| 0.0002 | 0.0001 | 0.0033 | 0.0021 | 0 | 0 | 0.0001 |
| 0.0009 | 0.001 | 0.0011 | 0.0009 | 0.0006 | 0 | 0.0006 |
| 1.29E-05 | 0 | 2.00E-04 | 0 | 0 | 0 | 0 |
| 7.71E-05 | 5.38E-05 | 2.00E-04 | 0.0001 | 0.0002 | 0 | 6.54E-05 |
| 0.0002 | 0.0001 | 0.0033 | 0.0021 | 0 | 0 | 0.0001 |
| 0.0001 | 0.0001 | 0.0035 | 0.0023 | 0 | 0 | 0 |
| 0.0001 | 0.0002 | 0.005 | 0.0035 | 0 | 0 | 0 |
| . | . | 0 | . | . | . | . |
| 2.61E-05 | 2.73E-05 | 8.00E-04 | 0.0003 | 0 | 0 | 0 |
| 2.57E-05 | 4.03E-05 | 8.00E-04 | 0.0003 | 0 | 0 | 0 |
| . | . | 0 | . | . | . | . |
| . | . | 0 | . | . | . | . |
| 0 | 8.09E-05 | 0.0012 | 0.0005 | 0 | 0 | 0 |
| . | . | 0 | . | . | . | . |
| 1.29E-05 | 0 | 6.54E-05 | 0 | 0 | 0 | 6.54E-05 |
| 1.29E-05 | 0 | 2.00E-04 | 0 | 0 | 0 | 0 |
| 0 | 1.35E-05 | 2.00E-04 | 0 | 0 | 0 | 0 |
| . | . | 0 | . | . | . | . |
| 6.42E-05 | 1.35E-05 | 6.00E-04 | 0.0002 | 4.82E-05 | 0 | 0 |
| 0.0001 | 6.73E-05 | 0.0023 | 0.0013 | 0 | 0 | 6.55E-05 |
| 1.29E-05 | 0 | 2.00E-04 | 0 | 0 | 0 | 0 |
| . | . | 0 | . | . | . | . |
| 0.0001 | 0.0001 | 8.00E-04 | 0.0003 | 9.66E-05 | 0 | 0.0002 |
| 9.00E-05 | 0.0002 | 0.0042 | 0.0029 | 0 | 0 | 6.55E-05 |
| 1.28E-05 | 0 | 2.00E-04 | 0 | 0 | 0 | 0 |
| 0.0004 | 0.0003 | 5.00E-04 | 0.0004 | 0.0003 | 0.0011 | 0 |
| 0 | 1.35E-05 | 2.00E-04 | 0 | 0 | 0 | 0 |

| | | | | | | |
|---|---|---|---|---|---|---|
| 0.0001 | 0.0002 | 0.005 | 0.0035 | 0 | 0 | 0 |
| 2.57E-05 | 4.03E-05 | 8.00E-04 | 0.0003 | 0 | 0 | 0 |
| 6.43E-05 | 5.38E-05 | 2.00E-04 | 0 | 7.24E-05 | 0 | 0 |
| 1.29E-05 | 4.04E-05 | 6.00E-04 | 0.0002 | 0 | 0 | 6.56E-05 |
| 5.15E-05 | 0.0001 | 0.0014 | 0.0006 | 0.0001 | 0 | 0 |
| . | . | 0 | . | . | . | . |
| 5.14E-05 | 1.35E-05 | 6.54E-05 | 0 | 2.41E-05 | 0 | 6.54E-05 |
| 2.61E-05 | 2.73E-05 | 8.00E-04 | 0.0003 | 0 | 0 | 0 |
| 2.58E-05 | 2.71E-05 | 6.00E-04 | 0.0002 | 0 | 0 | 0 |
| . | . | 0 | . | . | . | . |
| 1.29E-05 | 5.38E-05 | 0.001 | 0.0004 | 0 | 0 | 0 |
| . | . | 0 | . | . | . | . |
| . | . | 0 | . | . | . | . |
| 0.0005 | 0.0006 | 0.0019 | 0.0015 | 0.0019 | 0 | 0 |
| . | . | 0 | . | . | . | . |
| 2.57E-05 | 0 | 4.00E-04 | 6.82E-05 | 0 | 0 | 0 |
| 0.0001 | 0.0001 | 0.0037 | 0.0024 | 0 | 0 | 6.54E-05 |
| 0.0004 | 0.0005 | 0.0031 | 0.0019 | 7.24E-05 | 0 | 0.0012 |
| 0.0002 | 0.0001 | 0.0038 | 0.0025 | 4.83E-05 | 0 | 0 |
| 0.0001 | 0.0001 | 0.0037 | 0.0024 | 0 | 0 | 6.54E-05 |
| 5.19E-05 | 5.45E-05 | 6.00E-04 | 0.0002 | 2.44E-05 | 0 | 0 |
| 1.29E-05 | 1.35E-05 | 2.00E-04 | 0 | 2.41E-05 | 0 | 0 |
| 0.0001 | 0.0001 | 0.0035 | 0.0023 | 0 | 0 | 0 |
| 0.0001 | 0.0001 | 0.0037 | 0.0024 | 0 | 0 | 6.54E-05 |
| 0 | 4.04E-05 | 6.00E-04 | 0.0002 | 0 | 0 | 0 |
| 6.44E-05 | 5.40E-05 | 0.0017 | 0.0009 | 0 | 0 | 0 |
| 0.0004 | 0.0005 | 0.0031 | 0.0019 | 7.24E-05 | 0 | 0.0012 |
| 0.0003 | 0.0005 | 0.0037 | 0.0024 | 0 | 0 | 0 |
| 0.0002 | 0.0001 | 0.0033 | 0.0021 | 0 | 0 | 0.0001 |
| 6.43E-05 | 0.0001 | 0.0029 | 0.0018 | 0 | 0 | 0 |
| . | . | 0 | . | . | . | . |
| . | . | 0 | . | . | . | . |
| 5.14E-05 | 1.35E-05 | 6.54E-05 | 0 | 2.41E-05 | 0 | 6.54E-05 |
| 0.0001 | 0.0001 | 0.0037 | 0.0024 | 0 | 0 | 6.54E-05 |
| 0.0003 | 0.0005 | 0.0037 | 0.0024 | 0 | 0 | 0 |
| 0.0001 | 0.0001 | 8.00E-04 | 0.0003 | 9.66E-05 | 0 | 0.0002 |
| . | . | 0 | . | . | . | . |
| 9.00E-05 | 8.08E-05 | 2.00E-04 | 0 | 0.0001 | 0 | 0 |
| . | . | 0 | . | . | . | . |
| . | . | 0 | . | . | . | . |
| 8.99E-05 | 0.0001 | 0.0025 | 0.0015 | 0 | 0 | 0.0001 |
| 1.29E-05 | 4.04E-05 | 6.00E-04 | 0.0002 | 0 | 0 | 6.56E-05 |
| 0.0001 | 0.0001 | 0.0037 | 0.0024 | 0 | 0 | 6.54E-05 |

| | | | | | | |
|---|---|---|---|---|---|---|
| 8.99E-05 | 0.0003 | 0.0044 | 0.003 | 2.41E-05 | 0 | 0 |
| 0.0003 | 0.0005 | 0.0037 | 0.0024 | 0 | 0 | 0 |
| 0 | 1.35E-05 | 2.00E-04 | 0 | 0 | 0 | 0 |

| gnomad312_ | gnomad312_ | gnomad312_ | gnomad312_ | gnomad312_ | gnomad312_ | gnomad312_ |
|---|---|---|---|---|---|---|
| . | . | . | . | . | . | . |
| 0 | 0.0002 | 0.0002 | 0 | 0.0019 | 0 | 0.0002 |
| 0.0029 | 0 | 0 | 0 | 0.0034 | 0.0038 | 0 |
| . | . | . | . | . | . | . |
| 0 | 0.001 | 0.006 | 0 | 0.0015 | 0.001 | 0.0002 |
| 0 | 0 | 0 | 0 | 1.47E-05 | 0 | 0 |
| 0 | 0 | 0 | 0 | 5.88E-05 | 0 | 0 |
| . | . | . | . | . | . | . |
| 0 | 0 | 0.0004 | 0 | 0.0041 | 0.0014 | 0 |
| 0.0012 | 0 | 0.0016 | 0 | 0.0024 | 0.0014 | 0.0004 |
| 0.0006 | 0 | 0.0007 | 0.0032 | 0.002 | 0.0005 | 0 |
| 0.0029 | 0 | 0 | 0 | 0.0034 | 0.0038 | 0 |
| 0 | 0 | 0 | 0 | 1.47E-05 | 0 | 0 |
| 0.0003 | 0 | 0.0023 | 0 | 0.0013 | 0.0005 | 0 |
| 0 | 0 | 0 | 0 | 4.41E-05 | 0 | 0 |
| 0 | 0 | 0 | 0 | 4.41E-05 | 0 | 0.0021 |
| . | . | . | . | . | . | . |
| . | . | . | . | . | . | . |
| 0.0006 | 0 | 0.0007 | 0.0032 | 0.002 | 0.0005 | 0 |
| 0 | 0 | 0 | 0 | 0.0001 | 0.0005 | 0 |
| 0.0003 | 0 | 0 | 0 | 5.88E-05 | 0 | 0 |
| 0 | 0 | 0 | 0 | 2.94E-05 | 0 | 0 |
| 0 | 0 | 0 | 0 | 7.35E-05 | 0 | 0.0002 |
| 0 | 0 | 0 | 0 | 0.0001 | 0 | 0 |
| . | . | . | . | . | . | . |
| 0 | 0 | 0 | 0 | 0 | 0 | 0 |
| 0 | 0 | 0 | 0 | 2.94E-05 | 0 | 0 |
| 0 | 0 | 0 | 0 | 4.41E-05 | 0 | 0 |
| . | . | . | . | . | . | . |
| 0 | 0 | 9.41E-05 | 0 | 0.0004 | 0 | 0 |
| 0.0006 | 0 | 0.0007 | 0.0032 | 0.002 | 0.0005 | 0 |
| 0 | 0 | 0.0077 | 0 | 0.0005 | 0 | 0 |
| 0 | 0 | 0.0013 | 0 | 0 | 0 | 0 |
| 0.0012 | 0 | 0.0016 | 0 | 0.0024 | 0.0014 | 0.0004 |
| 0 | 0 | 0 | 0 | 1.47E-05 | 0 | 0.0044 |
| 0 | 0 | 0 | 0 | 1.47E-05 | 0 | 0 |
| 0 | 0 | 0 | 0 | 1.47E-05 | 0 | 0 |
| . | . | . | . | . | . | . |
| 0 | 0 | 0.0064 | 0 | 0.001 | 0.0014 | 0.0002 |
| . | . | . | . | . | . | . |
| 0 | 0 | 0 | 0 | 0.0006 | 0.0005 | 0.0023 |
| 0 | 0 | 0 | 0 | 4.41E-05 | 0 | 0 |

| | | | | | | |
|---|---|---|---|---|---|---|
| 0.0029 | 0 | 0 | 0 | 0.0034 | 0.0038 | 0 |
| 0 | 0 | 0 | 0 | 1.47E-05 | 0 | 0 |
| 0.006 | 0 | 0 | 0.0095 | 0.0007 | 0.0048 | 0.001 |
| . | . | . | . | . | . | . |
| 0.0003 | 0 | 0 | 0 | 5.88E-05 | 0 | 0 |
| . | . | . | . | . | . | . |
| 0 | 0 | 0 | 0 | 1.47E-05 | 0 | 0 |
| 0 | 0 | 0 | 0 | 0.0001 | 0 | 0 |
| 0 | 0 | 0 | 0 | 0.0004 | 0 | 0 |
| 0 | 0 | 0 | 0 | 0.0001 | 0 | 0 |
| 0 | 0 | 0 | 0 | 1.47E-05 | 0 | 0.0002 |
| 0 | 0 | 9.41E-05 | 0 | 8.82E-05 | 0 | 0 |
| 0 | 0 | 0 | 0 | 2.94E-05 | 0 | 0 |
| 0 | 0 | 0 | 0 | 7.43E-05 | 0 | 0 |
| 0 | 0 | 0.0005 | 0 | 0.0009 | 0 | 0 |
| 0 | 0.0004 | 0 | 0 | 1.47E-05 | 0 | 0 |
| 0 | 0 | 0 | 0 | 1.47E-05 | 0 | 0 |
| 0 | 0 | 0 | 0 | 0.0008 | 0 | 0.0006 |
| 0 | 0 | 0 | 0 | 1.47E-05 | 0 | 0 |
| 0 | 0 | 9.42E-05 | 0.0158 | 0.0013 | 0 | 0.0004 |
| 0 | 0 | 0 | 0 | 0.0004 | 0 | 0.001 |
| 0.0003 | 0.0016 | 0 | 0 | 0.0003 | 0.0005 | 0.0008 |
| 0.0029 | 0 | 0 | 0 | 0.0034 | 0.0038 | 0 |
| 0 | 0 | 0.0006 | 0 | 0.0015 | 0.001 | 0.0008 |
| 0 | 0 | 0 | 0 | 0 | 0.0005 | 0 |
| 0.0006 | 0 | 0.0007 | 0.0032 | 0.002 | 0.0005 | 0 |
| . | . | . | . | . | . | . |
| 0 | 0 | 0 | 0 | 0 | 0 | 0 |
| 0.002 | 0.0002 | 0.0003 | 0 | 0.0033 | 0.0024 | 0 |
| 0 | 0 | 0 | 0 | 5.88E-05 | 0 | 0 |
| . | . | . | . | . | . | . |
| 0 | 0 | 0 | 0 | 7.35E-05 | 0 | 0 |
| 0 | 0 | 0 | 0 | 0 | 0 | 0 |
| 0 | 0 | 0.0004 | 0 | 0.0041 | 0.0014 | 0 |
| 0.0029 | 0 | 0 | 0 | 0.0034 | 0.0038 | 0 |
| 0.0029 | 0 | 0 | 0 | 0.0034 | 0.0038 | 0 |
| . | . | . | . | . | . | . |
| 0 | 0 | 0 | 0 | 0.0002 | 0 | 0 |
| 0 | 0 | 0 | 0 | 1.47E-05 | 0 | 0 |
| . | . | . | . | . | . | . |
| 0 | 0 | 9.41E-05 | 0 | 0.0002 | 0.0005 | 0 |
| 0 | 0 | 0 | 0 | 4.41E-05 | 0 | 0.0021 |
| 0 | 0 | 0 | 0 | 0.0002 | 0 | 0 |

| | | | | | | |
|---|---|---|---|---|---|---|
| 0 | 0.0002 | 0 | 0 | 0.0001 | 0 | 0 |
| 0 | 0 | 0 | 0 | 5.92E-05 | 0 | 0.0004 |
| 0.002 | 0 | 0 | 0 | 0.0024 | 0.0029 | 0 |
| . | . | . | . | . | . | . |
| 0 | 0 | 0 | 0 | 0.0004 | 0 | 0 |
| 0 | 0 | 0 | 0 | 0 | 0 | 0 |
| 0 | 0 | 0 | 0.0063 | 0 | 0 | 0 |
| 0.0029 | 0 | 0 | 0 | 0.0034 | 0.0038 | 0 |
| 0 | 0 | 0 | 0 | 2.94E-05 | 0 | 0 |
| 0 | 0 | 0 | 0 | 1.47E-05 | 0 | 0 |
| 0 | 0 | 0 | 0 | 2.94E-05 | 0 | 0 |
| 0 | 0.001 | 0.006 | 0 | 0.0015 | 0.001 | 0.0002 |
| 0 | 0 | 0 | 0 | 4.41E-05 | 0 | 0 |
| 0 | 0.0004 | 0 | 0 | 0 | 0 | 0 |
| 0 | 0.0002 | 0.0002 | 0 | 0.0019 | 0 | 0.0002 |
| 0 | 0 | 0.0004 | 0 | 0.0041 | 0.0014 | 0 |
| 0 | 0 | 0 | 0 | 0 | 0 | 0 |
| . | . | . | . | . | . | . |
| 0 | 0 | 0 | 0 | 0.0002 | 0 | 0 |
| 0.0003 | 0 | 0 | 0 | 5.88E-05 | 0.0005 | 0 |
| 0 | 0 | 0 | 0 | 1.47E-05 | 0 | 0 |
| . | . | . | . | . | . | . |
| 0 | 0.0002 | 0 | 0 | 0 | 0 | 0.0004 |
| 0.0029 | 0 | 0 | 0 | 0.0034 | 0.0038 | 0 |
| 0.0006 | 0 | 0.0007 | 0.0032 | 0.002 | 0.0005 | 0 |
| 0 | 0 | 0 | 0 | 0.0002 | 0 | 0 |
| 0.0012 | 0 | 0.0016 | 0 | 0.0024 | 0.0014 | 0.0004 |
| . | . | . | . | . | . | . |
| 0 | 0 | 9.46E-05 | 0 | 0.0006 | 0 | 0 |
| . | . | . | . | . | . | . |
| 0 | 0 | 9.42E-05 | 0 | 0.0002 | 0.0005 | 0 |
| 0 | 0 | 0 | 0 | 0.0002 | 0.0014 | 0 |
| 0 | 0 | 0 | 0 | 0.001 | 0.0005 | 0.0002 |
| 0 | 0 | 0 | 0 | 0 | 0.0005 | 0.0004 |
| 0 | 0 | 9.42E-05 | 0.0158 | 0.0013 | 0 | 0.0004 |
| 0 | 0 | 0 | 0 | 4.41E-05 | 0 | 0 |
| 0 | 0 | 0 | 0 | 0 | 0 | 0 |
| 0 | 0.0002 | 0 | 0 | 0 | 0 | 0 |
| 0 | 0 | 0 | 0 | 4.41E-05 | 0 | 0 |
| 0.0043 | 0 | 0 | 0 | 4.41E-05 | 0 | 0 |
| 0 | 0.0004 | 0 | 0.0096 | 0.0004 | 0.0005 | 0.0023 |
| 0 | 0 | 0 | 0 | 1.47E-05 | 0 | 0 |
| 0 | 0 | 9.42E-05 | 0 | 0.0002 | 0.0005 | 0 |

| | | | | | | |
|---|---|---|---|---|---|---|
| 0 | 0 | 0 | 0 | 2.94E-05 | 0 | 0 |
| 0.0029 | 0 | 0 | 0 | 0.0034 | 0.0038 | 0 |
| 0 | 0 | 9.42E-05 | 0.0158 | 0.0013 | 0 | 0.0004 |
| 0 | 0 | 0 | 0 | 4.41E-05 | 0 | 0 |
| 0 | 0 | 0 | 0 | 7.35E-05 | 0 | 0 |
| . | . | . | . | . | . | . |
| 0 | 0 | 0.0004 | 0 | 0.0007 | 0.0014 | 0.0004 |
| 0 | 0 | 0.0004 | 0 | 0.0003 | 0 | 0 |
| . | . | . | . | . | . | . |
| . | . | . | . | . | . | . |
| . | . | . | . | . | . | . |
| 0 | 0 | 0 | 0 | 1.47E-05 | 0 | 0 |
| 0 | 0.001 | 0 | 0 | 0 | 0 | 0 |
| 0 | 0 | 0 | 0 | 1.47E-05 | 0 | 0 |
| 0 | 0.0002 | 0 | 0 | 5.88E-05 | 0.0005 | 0 |
| . | . | . | . | . | . | . |
| . | . | . | . | . | . | . |
| 0 | 0 | 0 | 0 | 4.41E-05 | 0 | 0 |
| 0.0012 | 0 | 0.0016 | 0 | 0.0024 | 0.0014 | 0.0004 |
| 0 | 0 | 0 | 0 | 2.94E-05 | 0.0005 | 0 |
| 0 | 0 | 0 | 0 | 0 | 0 | 0 |
| 0 | 0 | 0.0004 | 0 | 0.0007 | 0.0014 | 0.0004 |
| 0 | 0 | 0 | 0 | 0.0003 | 0 | 0 |
| 0 | 0 | 0 | 0 | 4.41E-05 | 0 | 0 |
| . | . | . | . | . | . | . |
| 0 | 0 | 0 | 0 | 2.94E-05 | 0 | 0 |
| 0 | 0 | 0 | 0 | 0 | 0 | 0 |
| 0 | 0 | 0 | 0 | 0 | 0 | 0 |
| 0 | 0 | 0 | 0 | 0 | 0 | 0 |
| 0 | 0 | 0 | 0 | 2.94E-05 | 0 | 0 |
| 0.002 | 0.0002 | 0.0003 | 0 | 0.0033 | 0.0024 | 0 |
| 0 | 0 | 0.0004 | 0 | 0.0007 | 0.0014 | 0.0004 |
| . | . | . | . | . | . | . |
| 0 | 0 | 0 | 0 | 0.0002 | 0 | 0 |
| 0 | 0 | 0 | 0 | 0.0003 | 0.0014 | 0 |
| 0 | 0 | 0 | 0 | 0 | 0 | 0 |
| 0.0075 | 0 | 0 | 0.0095 | 0.0005 | 0.0014 | 0 |
| 0 | 0 | 0 | 0 | 4.41E-05 | 0 | 0 |
| 0 | 0 | 0 | 0 | 0 | 0 | 0 |
| 0.002 | 0.0002 | 0.0003 | 0 | 0.0033 | 0.0024 | 0 |
| 0 | 0 | 0 | 0 | 0 | 0 | 0 |
| 0.004 | 0 | 0 | 0 | 0.0013 | 0.0019 | 0.0002 |
| 0 | 0 | 0 | 0 | 0 | 0 | 0 |

| | | | | | | |
|---|---|---|---|---|---|---|
| 0.0003 | 0 | 0 | 0 | 0 | 0 | 0 |
| 0 | 0 | 0 | 0 | 2.94E-05 | 0.0005 | 0 |
| 0 | 0 | 0 | 0 | 0 | 0 | 0 |
| . | . | . | . | . | . | . |
| 0 | 0 | 0 | 0 | 0 | 0 | 0 |
| 0 | 0 | 0 | 0 | 0 | 0 | 0 |
| 0.0014 | 0 | 9.43E-05 | 0 | 0.0006 | 0 | 0 |
| 0 | 0 | 0 | 0 | 0 | 0 | 0 |
| 0 | 0 | 0 | 0 | 7.35E-05 | 0.0005 | 0.0002 |
| 0.0003 | 0 | 9.43E-05 | 0 | 0.0001 | 0.0014 | 0 |
| 0 | 0.001 | 0 | 0 | 0 | 0 | 0 |
| 0 | 0 | 9.41E-05 | 0 | 0.0005 | 0 | 0 |
| 0 | 0 | 0 | 0 | 0 | 0 | 0 |
| 0 | 0 | 0 | 0 | 0 | 0 | 0 |
| 0 | 0 | 0 | 0 | 0 | 0 | 0 |
| . | . | . | . | . | . | . |
| . | . | . | . | . | . | . |
| 0 | 0 | 0 | 0 | 0 | 0 | 0.0002 |
| . | . | . | . | . | . | . |
| . | . | . | . | . | . | . |
| 0 | 0 | 0 | 0 | 4.41E-05 | 0 | 0 |
| 0 | 0 | 0 | 0 | 0 | 0.0014 | 0 |
| 0.0003 | 0 | 0.0023 | 0 | 0.0013 | 0.0005 | 0 |
| 0 | 0 | 0 | 0 | 0 | 0.0005 | 0 |
| 0 | 0 | 0 | 0 | 0 | 0 | 0 |
| 0 | 0.0002 | 0 | 0 | 4.41E-05 | 0 | 0.0002 |
| . | . | . | . | . | . | . |
| 0 | 0.001 | 0 | 0 | 0 | 0 | 0 |
| 0 | 0 | 0 | 0 | 0 | 0 | 0 |
| 0.0029 | 0 | 0 | 0 | 0.0034 | 0.0038 | 0 |
| 0.002 | 0.0002 | 0.0003 | 0 | 0.0033 | 0.0024 | 0 |
| 0 | 0 | 9.41E-05 | 0 | 0.0005 | 0 | 0 |
| 0.002 | 0.0002 | 0.0003 | 0 | 0.0033 | 0.0024 | 0 |
| 0 | 0 | 9.42E-05 | 0.0158 | 0.0013 | 0 | 0.0004 |
| 0 | 0 | 0 | 0 | 0 | 0 | 0 |
| . | . | . | . | . | . | . |
| 0 | 0 | 0 | 0 | 0.0006 | 0.0005 | 0.0023 |
| 0.0003 | 0 | 0.0023 | 0 | 0.0013 | 0.0005 | 0 |
| 0.006 | 0 | 0 | 0.0095 | 0.0007 | 0.0048 | 0.001 |
| . | . | . | . | . | . | . |
| 0.0003 | 0 | 9.43E-05 | 0 | 0.0001 | 0.0014 | 0 |
| 0 | 0 | 0 | 0 | 5.88E-05 | 0.001 | 0 |
| 0 | 0 | 0 | 0 | 0.0003 | 0.0014 | 0 |

| | | | | | | |
|---|---|---|---|---|---|---|
| 0 | 0 | 0 | 0 | 0 | 0 | 0 |
| 0.002 | 0.0002 | 0.0003 | 0 | 0.0033 | 0.0024 | 0 |
| 0.0003 | 0 | 0.0023 | 0 | 0.0013 | 0.0005 | 0 |
| 0 | 0 | 0 | 0 | 0.0003 | 0.0014 | 0 |
| 0 | 0 | 0 | 0 | 0 | 0 | 0 |
| 0 | 0 | 0 | 0 | 0.0003 | 0.0014 | 0 |
| . | . | . | . | . | . | . |
| . | . | . | . | . | . | . |
| 0 | 0 | 0 | 0 | 4.54E-05 | 0 | 0.0026 |
| 0.0075 | 0 | 0 | 0.0095 | 0.0005 | 0.0014 | 0 |
| 0 | 0 | 0 | 0 | 0.0006 | 0.0005 | 0.0023 |
| 0 | 0 | 0 | 0 | 0 | 0 | 0 |
| 0.0003 | 0 | 9.43E-05 | 0 | 0.0001 | 0.0014 | 0 |
| . | . | . | . | . | . | . |
| 0 | 0 | 0 | 0 | 0.0002 | 0 | 0 |
| . | . | . | . | . | . | . |
| 0 | 0.0033 | 0 | 0 | 5.88E-05 | 0 | 0.0002 |
| 0.0081 | 0.0006 | 0 | 0.0032 | 0.0011 | 0.0005 | 0.001 |
| 0 | 0.0002 | 0 | 0 | 0 | 0 | 0 |
| 0 | 0 | 0 | 0 | 0 | 0 | 0 |
| 0 | 0.0033 | 0 | 0 | 5.88E-05 | 0 | 0.0002 |
| 0 | 0.0035 | 0 | 0 | 0 | 0.0005 | 0 |
| 0 | 0.005 | 0 | 0 | 0 | 0 | 0.0002 |
| . | . | . | . | . | . | . |
| 0 | 0.0008 | 0 | 0 | 0 | 0 | 0 |
| 0 | 0.0008 | 0 | 0 | 0 | 0.0005 | 0 |
| . | . | . | . | . | . | . |
| . | . | . | . | . | . | . |
| 0 | 0 | 0 | 0 | 0 | 0 | 0.0012 |
| . | . | . | . | . | . | . |
| 0 | 0 | 0 | 0 | 0 | 0 | 0 |
| 0 | 0.0002 | 0 | 0 | 0 | 0 | 0 |
| 0 | 0.0002 | 0 | 0 | 0 | 0 | 0 |
| . | . | . | . | . | . | . |
| 0 | 0.0006 | 0 | 0 | 1.47E-05 | 0 | 0 |
| 0 | 0.0023 | 0 | 0 | 0 | 0.0005 | 0 |
| 0 | 0.0002 | 0 | 0 | 0 | 0 | 0 |
| . | . | . | . | . | . | . |
| 0 | 0.0008 | 0.0002 | 0 | 7.35E-05 | 0.0005 | 0 |
| 0 | 0.0042 | 0 | 0 | 0 | 0.0005 | 0 |
| 0 | 0.0002 | 0 | 0 | 0 | 0 | 0 |
| 0 | 0.0004 | 0 | 0 | 0.0005 | 0 | 0.0002 |
| 0 | 0.0002 | 0 | 0 | 0 | 0 | 0 |

| | | | | | | |
|---|---|---|---|---|---|---|
| 0 | 0.005 | 0 | 0 | 0 | 0 | 0.0002 |
| 0 | 0.0008 | 0 | 0 | 0 | 0.0005 | 0 |
| 0 | 0.0002 | 0 | 0 | 5.88E-05 | 0 | 0.0002 |
| 0 | 0.0006 | 0 | 0 | 0 | 0 | 0 |
| 0 | 0.0014 | 0 | 0 | 0 | 0 | 0 |
| . | . | . | . | . | . | . |
| 0 | 0 | 0 | 0 | 4.41E-05 | 0 | 0 |
| 0 | 0.0008 | 0 | 0 | 0 | 0 | 0 |
| 0 | 0.0006 | 0 | 0 | 1.47E-05 | 0 | 0 |
| . | . | . | . | . | . | . |
| 0 | 0.001 | 0 | 0 | 0 | 0 | 0 |
| . | . | . | . | . | . | . |
| . | . | . | . | . | . | . |
| 0 | 0.0002 | 0 | 0 | 1.47E-05 | 0.001 | 0 |
| . | . | . | . | . | . | . |
| 0 | 0.0004 | 0 | 0 | 0 | 0 | 0 |
| 0 | 0.0037 | 0 | 0 | 0 | 0.0005 | 0 |
| 0.0035 | 0.0031 | 0 | 0 | 0.0002 | 0 | 0.0008 |
| 0 | 0.0038 | 0 | 0 | 0 | 0 | 0 |
| 0 | 0.0037 | 0 | 0 | 0 | 0.0005 | 0 |
| 0 | 0.0006 | 0 | 0 | 5.91E-05 | 0 | 0 |
| 0 | 0.0002 | 0 | 0 | 0 | 0 | 0 |
| 0 | 0.0035 | 0 | 0 | 0 | 0.0005 | 0 |
| 0 | 0.0037 | 0 | 0 | 0 | 0.0005 | 0 |
| 0 | 0.0006 | 0 | 0 | 0 | 0 | 0 |
| 0 | 0.0017 | 0 | 0 | 0 | 0 | 0 |
| 0.0035 | 0.0031 | 0 | 0 | 0.0002 | 0 | 0.0008 |
| 0 | 0.0037 | 0 | 0 | 0.0004 | 0.0005 | 0.0019 |
| 0 | 0.0033 | 0 | 0 | 5.88E-05 | 0 | 0.0002 |
| 0 | 0.0029 | 0 | 0 | 0 | 0 | 0 |
| . | . | . | . | . | . | . |
| . | . | . | . | . | . | . |
| 0 | 0 | 0 | 0 | 4.41E-05 | 0 | 0 |
| 0 | 0.0037 | 0 | 0 | 0 | 0.0005 | 0 |
| 0 | 0.0037 | 0 | 0 | 0.0004 | 0.0005 | 0.0019 |
| 0 | 0.0008 | 0.0002 | 0 | 7.35E-05 | 0.0005 | 0 |
| . | . | . | . | . | . | . |
| 0.0003 | 0.0002 | 0 | 0 | 7.35E-05 | 0 | 0 |
| . | . | . | . | . | . | . |
| . | . | . | . | . | . | . |
| 0 | 0.0025 | 0 | 0 | 1.47E-05 | 0 | 0.0002 |
| 0 | 0.0006 | 0 | 0 | 0 | 0 | 0 |
| 0 | 0.0037 | 0 | 0 | 0 | 0.0005 | 0 |

| | | | | | | |
|---|---|---|---|---|---|---|
| 0 | 0.0044 | 0 | 0 | 1.47E-05 | 0 | 0.0002 |
| 0 | 0.0037 | 0 | 0 | 0.0004 | 0.0005 | 0.0019 |
| 0 | 0.0002 | 0 | 0 | 0 | 0 | 0 |

| REVEL | avsnp150 | CLNALLELEID | CLNDN | CLNDISDB | CLNREVSTAT | CLNSIG |
|---|---|---|---|---|---|---|
| . | . | . | . | . | . | . |
| . | rs578196082 | . | . | . | . | . |
| 0.143 | rs61762988 | . | . | . | . | . |
| 0.469 | rs777751998 | . | . | . | . | . |
| 0.321 | rs138539483 | . | . | . | . | . |
| 0.291 | . | 1691191 | Neurodevelo | MONDO:MOI | criteria_provi | Uncertain_sig |
| 0.693 | rs777268578 | . | . | . | . | . |
| 0.318 | rs756754424 | . | . | . | . | . |
| 0.676 | rs34329467 | . | . | . | . | . |
| 0.198 | rs112637700 | . | . | . | . | . |
| 0.233 | rs150119037 | . | . | . | . | . |
| 0.143 | rs61762988 | . | . | . | . | . |
| 0.317 | . | . | . | . | . | . |
| 0.496 | rs191475834 | . | . | . | . | . |
| . | rs954554112 | . | . | . | . | . |
| 0.379 | rs575451633 | 688823 | Camptomelic | MONDO:MOI | criteria_provi | Benign |
| 0.122 | . | . | . | . | . | . |
| 0.343 | . | . | . | . | . | . |
| 0.233 | rs150119037 | . | . | . | . | . |
| 0.279 | rs764049309 | . | . | . | . | . |
| 0.685 | rs745687224 | 1025143 | ADULT_syndr | MONDO:MOI | criteria_provi | Uncertain_sig |
| . | rs778139957 | 329509 | Orofacial_cle | MONDO:MOI | criteria_provi | Uncertain_sig |
| 0.51 | rs569425024 | . | . | . | . | . |
| 0.205 | rs371396397 | . | . | . | . | . |
| . | . | . | . | . | . | . |
| 0.221 | rs375998501 | . | . | . | . | . |
| 0.826 | rs771010789 | 1376869 | Autosomal_d | MONDO:MOI | criteria_provi | Uncertain_sig |
| 0.106 | rs752409607 | . | . | . | . | . |
| 0.295 | . | . | . | . | . | . |
| 0.465 | rs200109736 | . | . | . | . | . |
| 0.233 | rs150119037 | . | . | . | . | . |
| 0.574 | rs142362355 | . | . | . | . | . |
| 0.257 | rs761672999 | . | . | . | . | . |
| 0.198 | rs112637700 | . | . | . | . | . |
| 0.085 | rs569026862 | . | . | . | . | . |
| 0.198 | rs143987252 | . | . | . | . | . |
| 0.164 | . | . | . | . | . | . |
| 0.263 | . | . | . | . | . | . |
| 0.559 | rs192267587 | . | . | . | . | . |
| 0.597 | rs766583971 | 1442183 | ADULT_syndr | MONDO:MOI | criteria_provi | Uncertain_sig |
| 0.114 | rs141834826 | . | . | . | . | . |
| 0.273 | rs376651409 | . | . | . | . | . |

| | | | | | | |
|---|---|---|---|---|---|---|
| 0.143 | rs61762988 | . | . | . | . | . |
| 0.116 | . | . | . | . | . | . |
| 0.137 | rs142494121 | 173850 | Cardiomyopa | Human_Phen | criteria_provi | Conflicting_in |
| 0.64 | . | 910206 | Cardiomyopa | Human_Phen | criteria_provi | Uncertain_sig |
| 0.685 | rs745687224 | 1025143 | ADULT_syndr | MONDO:MOI | criteria_provi | Uncertain_sig |
| . | . | . | . | . | . | . |
| . | . | . | . | . | . | . |
| 0.181 | rs145963001 | . | . | . | . | . |
| 0.702 | rs121912767 | 32742 | Orofacial_cle | MONDO:MOI | criteria_provi | Conflicting_in |
| 0.139 | rs140945349 | 1522832 | not_provided | MedGen:CN5 | criteria_provi | Benign |
| 0.403 | rs759749431 | . | . | . | . | . |
| 0.27 | rs372507236 | . | . | . | . | . |
| 0.1 | rs192078390 | . | . | . | . | . |
| . | . | . | . | . | . | . |
| 0.061 | rs148104757 | 298238 | Vitreoretinop | Human_Phen | criteria_provi | Benign/Likely_ |
| 0.279 | rs748592687 | 1359181 | not_provided | MedGen:CN5 | criteria_provi | Uncertain_sig |
| 0.932 | rs780626687 | 197036 | Primary_dilat | EFO:EFO_000 | criteria_provi | Uncertain_sig |
| 0.125 | rs140803495 | . | . | . | . | . |
| 0.191 | . | . | . | . | . | . |
| 0.307 | rs142885240 | 54093 | Primary_dilat | EFO:EFO_000 | criteria_provi | Conflicting_in |
| 0.351 | rs142717240 | 54136 | Cardiomyopa | Human_Phen | criteria_provi | Conflicting_in |
| . | rs745803083 | . | . | . | . | . |
| 0.143 | rs61762988 | . | . | . | . | . |
| 0.523 | rs201675393 | . | . | . | . | . |
| 0.242 | rs367984272 | 1804409 | Cardiovascula | MedGen:CN2 | criteria_provi | Uncertain_sig |
| 0.233 | rs150119037 | . | . | . | . | . |
| 0.283 | rs531280433 | . | . | . | . | . |
| 0.087 | rs769755490 | 793786 | not_provided | MedGen:CN5 | criteria_provi | Likely_benign |
| 0.358 | rs140355865 | 722332 | not_provided | MedGen:CN5 | criteria_provi | Likely_benign |
| . | rs773853170 | . | . | . | . | . |
| 0.602 | . | 954262 | not_provided | MedGen:CN5 | criteria_provi | Uncertain_sig |
| 0.211 | rs370988754 | . | . | . | . | . |
| 0.548 | rs103988511 | 1461543 | Arrhythmoge | MONDO:MOI | criteria_provi | Uncertain_sig |
| 0.676 | rs34329467 | . | . | . | . | . |
| 0.143 | rs61762988 | . | . | . | . | . |
| 0.143 | rs61762988 | . | . | . | . | . |
| 0.409 | . | . | . | . | . | . |
| 0.145 | rs147773518 | . | . | . | . | . |
| 0.127 | . | 830929 | SHORT_syndr | MONDO:MOI | criteria_provi | Uncertain_sig |
| 0.651 | rs102611397 | . | . | . | . | . |
| 0.357 | rs149513743 | 178566 | Primary_dilat | EFO:EFO_000 | criteria_provi | Conflicting_in |
| 0.379 | rs575451633 | 688823 | Camptomelic | MONDO:MOI | criteria_provi | Benign |
| 0.417 | rs61753592 | . | . | . | . | . |

| | | | | | | | |
|---|---|---|---|---|---|---|---|
| | 0.137 | rs200234458 | . | . | . | . | . |
| . | | rs756517331 | 1415772 | not_provided | MedGen:CN5 | criteria_provi | Uncertain_sig |
| | 0.275 | rs148165391 | . | . | . | . | . |
| | 0.443 | . | . | . | . | . | . |
| | 0.702 | rs121912767 | 32742 | Orofacial_cle | MONDO:MOI | criteria_provi | Conflicting_in |
| | 0.346 | rs764585766 | . | . | . | . | . |
| | 0.089 | rs868002362 | 1015800 | Neurodevelo | MONDO:MOI | criteria_provi | Uncertain_sig |
| | 0.143 | rs61762988 | . | . | . | . | . |
| | 0.401 | rs371342595 | . | . | . | . | . |
| | 0.9 | . | . | . | . | . | . |
| . | | rs770947287 | 732616 | not_provided | MedGen:CN5 | criteria_provi | Likely_benign |
| | 0.321 | rs138539483 | . | . | . | . | . |
| | 0.673 | rs770008356 | . | . | . | . | . |
| . | | . | . | . | . | . | . |
| . | | rs578196082 | . | . | . | . | . |
| | 0.676 | rs34329467 | . | . | . | . | . |
| . | | rs758720302 | . | . | . | . | . |
| | 0.411 | . | . | . | . | . | . |
| | 0.031 | rs201932618 | . | . | . | . | . |
| . | | rs774278800 | . | . | . | . | . |
| | 0.557 | . | . | . | . | . | . |
| | 0.148 | . | . | . | . | . | . |
| . | | . | . | . | . | . | . |
| | 0.143 | rs61762988 | . | . | . | . | . |
| | 0.233 | rs150119037 | . | . | . | . | . |
| | 0.277 | rs150750800 | . | . | . | . | . |
| | 0.198 | rs112637700 | . | . | . | . | . |
| . | | . | . | . | . | . | . |
| | 0.078 | rs772364788 | . | . | . | . | . |
| | 0.631 | . | . | . | . | . | . |
| | 0.358 | rs74608225 | . | . | . | . | . |
| | 0.239 | rs150468995 | 1601390 | not_provided | MedGen:CN5 | criteria_provi | Likely_benign |
| | 0.325 | rs62620235 | 413445 | not_provided | MedGen:CN5 | criteria_provi | Uncertain_sig |
| | 0.189 | rs773532758 | . | . | . | . | . |
| | 0.307 | rs142885240 | 54093 | Primary_dilat | EFO:EFO_000 | criteria_provi | Conflicting_in |
| | 0.374 | . | 832447 | Arrhythmoge | MONDO:MOI | criteria_provi | Uncertain_sig |
| | 0.349 | rs779608764 | 1006928 | Arrhythmoge | MONDO:MOI | criteria_provi | Uncertain_sig |
| | 0.413 | rs767837787 | 428460 | SHORT_syndr | MONDO:MOI | criteria_provi | Uncertain_sig |
| | 0.508 | rs775080202 | 910358 | Cardiomyopa | Human_Phen | criteria_provi | Uncertain_sig |
| | 0.735 | rs376960358 | 38572 | Orofacial_cle | MONDO:MOI | criteria_provi | Conflicting_in |
| . | | rs727502780 | 33206 | Prostate_can | MedGen:C40 | criteria_provi | Conflicting_in |
| . | | rs764542298 | . | . | . | . | . |
| | 0.358 | rs74608225 | . | . | . | . | . |

| | | | | | | | |
|---|---|---|---|---|---|---|---|
| . | | . | . | . | . | . | . |
| | 0.143 | rs61762988 | . | . | . | . | . |
| | 0.307 | rs142885240 | 54093 | Primary_dilat | EFO:EFO_000 | criteria_provi | Conflicting_in |
| | 0.171 | rs773614862 | . | . | . | . | . |
| | 0.135 | rs367923318 | 1152169 | SHORT_syndr | MONDO:MOI | criteria_provi | Uncertain_sig |
| | 0.496 | . | . | . | . | . | . |
| | 0.72 | rs147000526 | 54110 | Cardiomyopa | Human_Phen | criteria_provi | Conflicting_in |
| | 0.38 | rs145471270 | . | . | . | . | . |
| | 0.637 | . | . | . | . | . | . |
| . | | . | . | . | . | . | . |
| . | | . | 368316 | not_provided | MedGen:CN5 | criteria_provi | Likely_pathog |
| | 0.276 | rs776782334 | . | . | . | . | . |
| | 0.363 | rs748630143 | . | . | . | . | . |
| . | | rs763973005 | . | . | . | . | . |
| | 0.103 | rs142860268 | 1499539 | not_provided | MedGen:CN5 | criteria_provi | Uncertain_sig |
| | 0.177 | . | . | . | . | . | . |
| | 0.616 | . | . | . | . | . | . |
| . | | rs763148343 | 1460855 | not_provided | MedGen:CN5 | criteria_provi | Uncertain_sig |
| | 0.198 | rs112637700 | . | . | . | . | . |
| | 0.064 | rs754282657 | 933048 | not_provided | MedGen:CN5 | criteria_provi | Uncertain_sig |
| | 0.847 | rs768407146 | . | . | . | . | . |
| | 0.72 | rs147000526 | 54110 | Cardiomyopa | Human_Phen | criteria_provi | Conflicting_in |
| . | | rs100173287 | . | . | . | . | . |
| | 0.239 | rs758237203 | . | . | . | . | . |
| | 0.261 | . | . | . | . | . | . |
| | 0.051 | rs144881278 | . | . | . | . | . |
| | 0.1 | . | . | . | . | . | . |
| | 0.057 | rs141074913 | 1407443 | not_provided | MedGen:CN5 | criteria_provi | Uncertain_sig |
| | 0.201 | rs147255921 | 298275 | Vitreoretinop | Human_Phen | criteria_provi | Benign/Likely |
| | 0.361 | rs749213773 | . | . | . | . | . |
| | 0.358 | rs140355865 | 722332 | not_provided | MedGen:CN5 | criteria_provi | Likely_benign |
| | 0.72 | rs147000526 | 54110 | Cardiomyopa | Human_Phen | criteria_provi | Conflicting_in |
| . | | . | . | . | . | . | . |
| | 0.352 | rs199916044 | 1418532 | not_provided | MedGen:CN5 | criteria_provi | Uncertain_sig |
| | 0.195 | rs201963528 | 738622 | not_provided | MedGen:CN5 | criteria_provi | Likely_benign |
| | 0.191 | rs105334990 | . | . | . | . | . |
| | 0.655 | rs139430946 | . | . | . | . | . |
| | 0.363 | rs191185509 | . | . | . | . | . |
| . | | . | . | . | . | . | . |
| | 0.358 | rs140355865 | 722332 | not_provided | MedGen:CN5 | criteria_provi | Likely_benign |
| | 0.825 | . | . | . | . | . | . |
| | 0.335 | rs61733441 | 1038783 | not_provided | MedGen:CN5 | no_assertion_ | Likely_benign |
| | 0.343 | rs183321849 | . | . | . | . | . |

| | | | | | | | |
|---|---|---|---|---|---|---|---|
| | 0.249 | rs763574306 | 1497275 | SHORT_syndr | MONDO:MOI | criteria_provi | Uncertain_sig |
| | 0.676 | rs371020228 | 511086 | Cardiomyopa | Human_Phen | criteria_provi | Uncertain_sig |
| . | | . | . | . | . | . | . |
| . | | rs774964135 | . | . | . | . | . |
| | 0.1 | . | . | . | . | . | . |
| | 0.219 | . | . | . | . | . | . |
| | 0.523 | rs200140155 | . | . | . | . | . |
| | 0.847 | rs768407146 | . | . | . | . | . |
| | 0.265 | rs150993445 | . | . | . | . | . |
| . | | rs778061502 | 707536 | not_provided | MedGen:CN5 | criteria_provi | Likely_benign |
| | 0.216 | rs747962162 | . | . | . | . | . |
| | 0.243 | rs62639993 | . | . | . | . | . |
| | 0.467 | rs552784311 | . | . | . | . | . |
| | 0.467 | rs552784311 | . | . | . | . | . |
| . | | . | . | . | . | . | . |
| . | | rs767001931 | . | . | . | . | . |
| | 0.818 | . | . | . | . | . | . |
| | 0.386 | . | . | . | . | . | . |
| | 0.419 | rs753241704 | . | . | . | . | . |
| . | | . | . | . | . | . | . |
| | 0.239 | rs758237203 | . | . | . | . | . |
| | 0.415 | rs139530177 | . | . | . | . | . |
| | 0.496 | rs191475834 | . | . | . | . | . |
| | 0.162 | rs532561475 | 1343449 | not_provided | MedGen:CN5 | criteria_provi | Uncertain_sig |
| | 0.522 | rs560943760 | . | . | . | . | . |
| | 0.204 | rs181722888 | . | . | . | . | . |
| . | | . | . | . | . | . | . |
| | 0.216 | rs747962162 | . | . | . | . | . |
| | 0.318 | rs145123264 | . | . | . | . | . |
| | 0.143 | rs61762988 | . | . | . | . | . |
| | 0.358 | rs140355865 | 722332 | not_provided | MedGen:CN5 | criteria_provi | Likely_benign |
| | 0.243 | rs62639993 | . | . | . | . | . |
| | 0.358 | rs140355865 | 722332 | not_provided | MedGen:CN5 | criteria_provi | Likely_benign |
| | 0.307 | rs142885240 | 54093 | Primary_dilat | EFO:EFO_000 | criteria_provi | Conflicting_in |
| | 0.214 | . | . | . | . | . | . |
| | 0.155 | rs768910172 | . | . | . | . | . |
| | 0.114 | rs141834826 | . | . | . | . | . |
| | 0.496 | rs191475834 | . | . | . | . | . |
| | 0.137 | rs142494121 | 173850 | Cardiomyopa | Human_Phen | criteria_provi | Conflicting_in |
| | 0.667 | rs754598854 | . | . | . | . | . |
| . | | rs778061502 | 707536 | not_provided | MedGen:CN5 | criteria_provi | Likely_benign |
| | 0.186 | rs748902535 | . | . | . | . | . |
| | 0.195 | rs201963528 | 738622 | not_provided | MedGen:CN5 | criteria_provi | Likely_benign |

| | | | | | | | |
|---|---|---|---|---|---|---|---|
| . | | rs770977933 | . | . | . | . | . |
| | 0.358 | rs140355865 | 722332 | not_provided | MedGen:CN5 | criteria_provi | Likely_benign |
| | 0.496 | rs191475834 | . | . | . | . | . |
| | 0.195 | rs201963528 | 738622 | not_provided | MedGen:CN5 | criteria_provi | Likely_benign |
| | 0.1 | . | . | . | . | . | . |
| | 0.195 | rs201963528 | 738622 | not_provided | MedGen:CN5 | criteria_provi | Likely_benign |
| | 0.04 | . | . | . | . | . | . |
| | 0.129 | . | . | . | . | . | . |
| . | | rs768349158 | 719105 | not_provided | MedGen:CN5 | criteria_provi | Likely_benign |
| | 0.655 | rs139430946 | . | . | . | . | . |
| | 0.114 | rs141834826 | . | . | . | . | . |
| | 0.067 | rs61733390 | 1094632 | not_provided | MedGen:CN5 | criteria_provi | Likely_benign |
| . | | rs778061502 | 707536 | not_provided | MedGen:CN5 | criteria_provi | Likely_benign |
| | 0.316 | . | . | . | . | . | . |
| | 0.352 | rs199916044 | 1418532 | not_provided | MedGen:CN5 | criteria_provi | Uncertain_sig |
| | 0.323 | . | . | . | . | . | . |
| | 0.312 | rs145101521 | . | . | . | . | . |
| . | | . | 1602378 | not_provided | MedGen:CN5 | criteria_provi | Likely_benign |
| | 0.678 | rs775388344 | . | . | . | . | . |
| | 0.289 | rs764820053 | . | . | . | . | . |
| | 0.312 | rs145101521 | . | . | . | . | . |
| | 0.522 | rs202087317 | . | . | . | . | . |
| . | | . | . | . | . | . | . |
| | 0.457 | rs772742406 | . | . | . | . | . |
| . | | rs747790677 | . | . | . | . | . |
| | 0.605 | rs770493925 | 1656379 | Orofacial_cle | MONDO:MOI | criteria_provi | Conflicting_in |
| | 0.097 | rs749204336 | . | . | . | . | . |
| . | | . | . | . | . | . | . |
| . | | rs766783747 | . | . | . | . | . |
| | 0.335 | rs751566392 | 301057 | Arrhythmoge | MONDO:MOI | criteria_provi | Uncertain_sig |
| | 0.621 | rs771101622 | . | . | . | . | . |
| | 0.438 | rs397516944 | 54090 | Cardiomyopa | Human_Phen | criteria_provi | Uncertain_sig |
| | 0.788 | rs767711596 | . | . | . | . | . |
| | 0.402 | rs762586316 | . | . | . | . | . |
| . | | rs761733639 | . | . | . | . | . |
| | 0.265 | rs144202228 | 1073021 | not_provided | MedGen:CN5 | criteria_provi | Likely_benign |
| | 0.438 | rs397516944 | 54090 | Cardiomyopa | Human_Phen | criteria_provi | Uncertain_sig |
| | 0.602 | . | . | . | . | . | . |
| | 0.127 | rs139283675 | . | . | . | . | . |
| | 0.037 | rs78429618 | . | . | . | . | . |
| | 0.645 | rs764276612 | . | . | . | . | . |
| | 0.78 | rs141683209 | . | . | . | . | . |
| | 0.367 | rs777401627 | . | . | . | . | . |

| | | | | | | | |
|---|---|---|---|---|---|---|---|
| . | | . | . | . | . | . | . |
| | 0.605 | rs770493925 | 1656379 | Orofacial_cle | MONDO:MOI | criteria_provi | Conflicting_in |
| | 0.117 | rs768307299 | . | . | . | . | . |
| | 0.577 | rs767906723 | 1493932 | ADULT_syndr | MONDO:MOI | criteria_provi | Uncertain_sig |
| | 0.204 | rs144392839 | 224330 | Primary_dilat | EFO:EFO_000 | criteria_provi | Conflicting_in |
| | 0.321 | rs776000695 | . | . | . | . | . |
| | 0.521 | rs745803587 | . | . | . | . | . |
| . | | rs747790677 | . | . | . | . | . |
| | 0.082 | rs749344552 | . | . | . | . | . |
| | 0.442 | . | . | . | . | . | . |
| | 0.363 | rs748630143 | . | . | . | . | . |
| | 0.201 | . | . | . | . | . | . |
| | 0.913 | . | . | . | . | . | . |
| | 0.292 | rs139501357 | . | . | . | . | . |
| | 0.593 | rs777407386 | . | . | . | . | . |
| | 0.234 | rs766465283 | . | . | . | . | . |
| | 0.256 | rs575030530 | . | . | . | . | . |
| | 0.499 | rs199987597 | . | . | . | . | . |
| | 0.083 | rs141586900 | . | . | . | . | . |
| | 0.256 | rs575030530 | . | . | . | . | . |
| | 0.295 | rs578203724 | . | . | . | . | . |
| | 0.356 | rs752941845 | 359753 | Ventricular_t | EFO:EFO_000 | criteria_provi | Uncertain_sig |
| | 0.522 | rs202087317 | . | . | . | . | . |
| | 0.256 | rs575030530 | . | . | . | . | . |
| | 0.177 | rs751303379 | . | . | . | . | . |
| . | | rs767476557 | 1137111 | not_provided | MedGen:CN5 | criteria_provi | Likely_benign |
| | 0.499 | rs199987597 | . | . | . | . | . |
| | 0.215 | rs200525338 | 1609502 | not_provided | MedGen:CN5 | criteria_provi | Benign |
| | 0.312 | rs145101521 | . | . | . | . | . |
| | 0.23 | rs200056605 | . | . | . | . | . |
| | 0.287 | . | . | . | . | . | . |
| | 0.541 | rs781454634 | . | . | . | . | . |
| | 0.521 | rs745803587 | . | . | . | . | . |
| | 0.256 | rs575030530 | . | . | . | . | . |
| | 0.215 | rs200525338 | 1609502 | not_provided | MedGen:CN5 | criteria_provi | Benign |
| | 0.127 | rs139283675 | . | . | . | . | . |
| | 0.275 | rs758433852 | . | . | . | . | . |
| . | | rs765195788 | . | . | . | . | . |
| | 0.515 | . | . | . | . | . | . |
| | 0.81 | . | . | . | . | . | . |
| | 0.516 | rs534215890 | 739236 | Orofacial_cle | MONDO:MOI | criteria_provi | Likely_benign |
| | 0.577 | rs767906723 | 1493932 | ADULT_syndr | MONDO:MOI | criteria_provi | Uncertain_sig |
| | 0.256 | rs575030530 | . | . | . | . | . |

| | | | | | | |
|---|---|---|---|---|---|---|
| 0.124 | rs201704836 | . | . | . | . | . |
| 0.215 | rs200525338 | 1609502 | not_provided | MedGen:CN5 | criteria_provi | Benign |
| 0.261 | . | . | . | . | . | . |

ıterpretations_of_pathogenicity

ıterpretations_of_pathogenicity

ıterpretations_of_pathogenicity
ıterpretations_of_pathogenicity

ıterpretations_of_pathogenicity

nterpretations_of_pathogenicity

nterpretations_of_pathogenicity

nterpretations_of_pathogenicity
nterpretations_of_pathogenicity

ıterpretations_of_pathogenicity

ıterpretations_of_pathogenicity

ıterpretations_of_pathogenicity

ıterpretations_of_pathogenicity

ıterpretations_of_pathogenicity

ıterpretations_of_pathogenicity

ıterpretations_of_pathogenicity

nterpretations_of_pathogenicity

nterpretations_of_pathogenicity
